# LLPS condensates of Fha initiate the inside-out assembly of the type VI secretion system

**DOI:** 10.1101/2023.12.21.572528

**Authors:** Tong-Tong Pei, Ying An, Xing-Yu Wang, Han Luo, Yumin Kan, Hao Li, Ming-Xuan Tang, Zi-Yan Ye, Jia-Xin Liang, Tao Jian, Hao-Yu Zheng, Zeng-Hang Wang, Xiaoye Liang, Mingjie Zhang, Xiaotian Liu, Tao Dong

## Abstract

The type VI secretion system (T6SS) is one of the most powerful nanomachines employed by Gram-negative pathogens for penetrating diverse cell envelopes, including bacteria and fungi, to deliver potent effectors into target cells. While the membrane-anchored contractile tubular structure of the T6SS is well characterized, the assembly process remains poorly understood. The prevailing model suggests that the assembly of T6SS initiates from its outer-membrane component. Here, we report a distinct model that the cytoplasmic protein Fha initiates T6SS assembly in *Acidovorax citrulli*, an important plant pathogen. Fha dictates the formation of the inner-membrane complex and the baseplate, and directly interacts with these key components. Importantly, imaging and biochemical assays reveal that Fha undergoes liquid-liquid phase separation (LLPS), forming condensates that selectively recruit essential T6SS proteins, which are otherwise dispersed in cells. Fha also exhibited conserved functions in human pathogens *Vibrio cholerae* and *Pseudomonas aeruginosa*. These findings unveil an inside-first LLPS-driven model for T6SS assembly and suggest LLPS might be broadly involved in mediating the assembly of bacterial macromolecular complexes and facilitating interspecies interactions and pathogenesis.

**Significance statement:** The T6SS plays a pivotal role in interspecies competition and host-microbe interactions by delivering toxins to various prokaryotes and eukaryotes. Its crucial function relies on a membrane-anchored macromolecular structure comprising at least 13 conserved components. However, the mechanisms governing the efficient assembly of its diverse cytosolic and membrane-bound components remain elusive. Here, we identify Fha, a conserved cytosolic protein, as a key initiator of T6SS assembly. Fha recruits multiple structural and effector components, forming LLPS condensates. Fha homologs of plant and human pathogens exhibit conserved functions. Our findings not only unveil an inside-first assembly model for the T6SS, initiating from inner-membrane and baseplate components, but also suggest LLPS may have a broader impact on bacterial physiology beyond intracellular activities.

## Introduction

Type VI secretion systems (T6SSs), commonly found in Gram-negative bacteria, play a pivotal role in delivering toxic effectors to various cell types, promoting survival in polymicrobial communities and host-microbe interactions (1–6). The T6SS conserved structure consists of a membrane complex, a baseplate, and a tail-tube complex (7–10). Current model suggests that T6SS assembly adopts an outside-first mechanism, whereby the outer-membrane (OM) protein TssJ initiates the process by recruiting two inner-membrane (IM) proteins TssM and TssL, with a 5-fold membrane complex characterized by a 3:2:2 stoichiometry (7, 11). The membrane complex then facilitates the assembly of the cytoplasmic baseplate, composed of TssE, TssF, TssG, and TssK (8, 12–14). The baseplate further supports the polymerization of the tail-tube complex, comprising stacked hexameric Hcp rings forming a needle-like tube, surrounded by a helical sheath composed of TssB/TssC subunits and tipped by a VgrG/PAAR spike (9, 12, 15). While the static structures of the T6SS have been characterized, the dynamic recruitment of these dispersed structural proteins for the assembly of the macromolecular membrane-anchored T6SS remains elusive.

Proteins containing intrinsically disordered regions frequently undergo liquid-liquid phase separation (LLPS) to form membrane-less organelles that play diverse roles in eukaryotic cells (16–18). Bacterial cells, despite their smaller size, also harbor LLPS-capable proteins, including ribonuclease E, RNA polymerase, cell division protein FtsZ, and the membrane-bound ABC transporter Rv1747 (19–22). These proteins form condensates that act as scaffolds, selectively recruiting one or a few client proteins and nucleic acids (19, 21). However, the role of LLPS in regulating more intricate macrostructural apparatuses, such as bacterial protein secretion systems (23), remains unknown.

*Acidovorax citrulli*, a bacterial pathogen that affects various cucurbit plants such as melons and watermelons, is responsible for causing bacterial fruit blotch (BFB), a disease with significant global economic burdens (24, 25). Notably, the T6SS of *A. citrulli* group Ⅱ model strain AAC00-1 displays constitutive killing activities against a wide range of bacterial and fungal species, including *Escherichia coli*, *Bacillus subtilis*, *Mycobacterium smegmatis*, and *Saccharomyces cerevisiae* (6, 26). This T6SS represents one of the most potent systems discovered thus far, providing a unique platform for understanding the capabilities of T6SS-mediated delivery.

In this study, we present an inside-first model for T6SS biogenesis, driven by the cytoplasmic protein Fha. Fha initiates the assembly of the inner-membrane complex and baseplate by interacting with multiple T6SS structural proteins. Remarkably, Fha undergoes LLPS both *in vitro* and *in vivo*, forming condensates that selectively recruit dispersed T6SS structural proteins within cells. We also demonstrate the functional conservation of Fha in two critical human pathogens, *Vibrio cholerae* and *Pseudomonas aeruginosa*. These findings not only unveil the critical role of Fha in T6SS initiation across diverse organisms but also suggest the broader significance of LLPS in facilitating bacterial macromolecular assembly for interspecies competition and pathogenesis.

## Results

### Fha promotes the assembly of T6SS membrane complex and baseplate

To visualize the T6SS assembly process in AAC00-1, we constructed chromosomal fusions of fluorescence proteins to the IM components TssL and TssM, baseplate component TssG, and a cytoplasmic T6SS protein Fha. Bacterial competition results showed potent killing activities of these fluorescence-labeled strains against *E. coli*, indicating the fluorescence-labeled strains are T6SS-active (Supplementary Figure 1 A and B). Through fluorescence microscopy, we examined sfGFP_TssL and sfGFP_TssG foci formation in a panel of mutants lacking crucial T6SS components (Figure 1A). We first confirmed the loss of T6SS secretion in these T6SS mutants using the Hcp secretion assay (Supplementary Figure 1C). Foci formation of TssL required Fha and TssM, while mutants lacking TssJ, TssF, TssB, Hcp, or an assembly chaperone TssA, displayed foci similar to the parental strain (Figure 1A). Similarly, there were comparable levels of TssM foci in both parental and the Δ*tssJ* samples but not in the Δ*fha* mutant (Supplementary Figure 1D), confirming the OM protein TssJ is dispensable for the formation of the IM complex TssM/L.

**Fig. 1.**
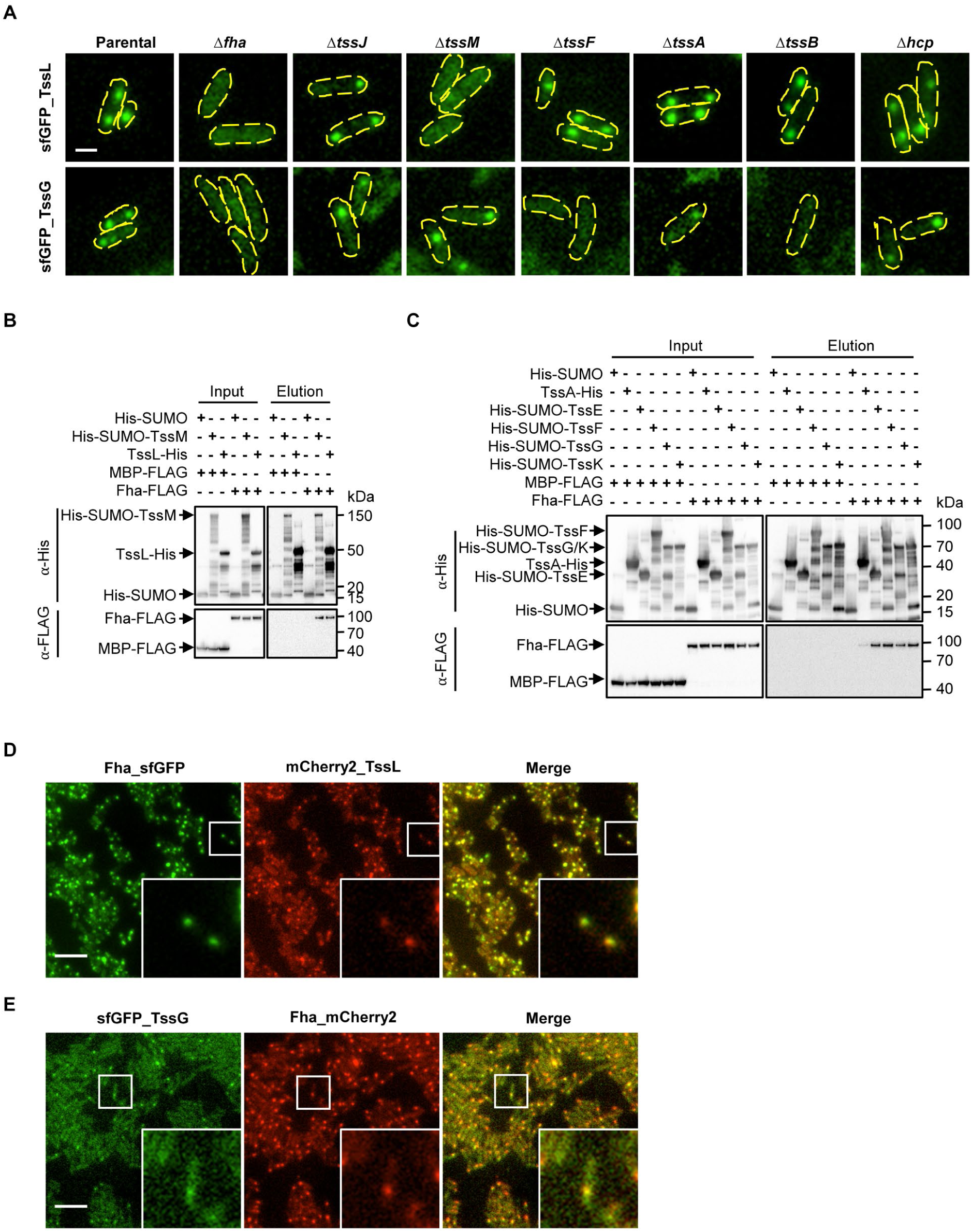
Fha facilitates the assembly of T6SS membrane complex and baseplate via direct interaction. **A**, Fluorescence microscopy images showing sfGFP_TssL and sfGFP_TssG localization in AAC00-1 Parental, Δ*fha*, Δ*tssJ*, Δ*tssM*, Δ*tssF*, Δ*tssA*, Δ*tssB*, and Δ*hcp*. A representative 5- ×5-μm field of cells is shown. Genotypes are indicated at the top. Yellow dashed lines depict the contour of bacterial cells. Scale bar: 1 μm. **B**, Interaction of Fha with TssM and TssL. Pull-down analysis was performed using His-tagged SUMO (control), SUMO-TssM, or TssL and FLAG-tagged MBP (control) or Fha. **C**, Interaction of Fha with TssA, TssE, TssF, TssG, and TssK. Pull-down analysis was performed using His-tagged SUMO (control), TssA, SUMO-TssE, SUMO-TssF, SUMO-TssG, SUMO-TssK, and FLAG-tagged MBP (control) or Fha. Fluorescence microscopy images showing co-localization between Fha_sfGFP and mCherry2_TssL (**D**) or sfGFP_TssG and Fha_mCherry2 (**E**) in AAC00-1. A representative 30- × 30-μm field of cells with a 3× magnified 5- ×5-μm inset (marked by box) is shown. Scale bar: 5 μm.

Furthermore, Fha, along with TssF and TssB, was required for TssG foci formation, whereas the absence of TssJ, TssM, TssA or Hcp had no effect (Figure 1A), suggesting the baseplate formation does not depend on the presence of the membrane complex or the tubular structure. Deleting *fha* abolished the T6SS-mediated killing ability of AAC00-1, which could be partially complemented by a plasmid-borne Fha (Supplementary Figure 1 E and F). Pull-down assays demonstrated the interaction of Fha with IM proteins TssL and TssM (Figure 1B), as well as with baseplate components TssE, TssF, TssG, and TssK, with a relatively weaker interaction observed with TssA (Figure 1C). Negative controls, His-SUMO and MBP-FLAG, did not show any interaction with T6SS proteins. To test whether the observed Fha interaction is specific, we examined the Fha homolog in *V. cholerae* V52 (Fha^VC^), which shares 32% sequence identity. Pull-down analysis showed that, while Fha^AC^ interacted with TssL and TssE, Fha^VC^ was found to interact with TssE but not TssL, suggesting some specificity of Fha interactions (Supplementary Figure 1G). Through a bacterial two-hybrid assay utilizing adenylate cyclase CyaA reconstitution (27), we confirmed the interaction between Fha with the cytoplasmic domain of TssL (Supplementary Figure 1H). However, there was no observed interaction between Fha and the cytoplasmic domain (66-458 aa) of TssM in the two-hybrid assay, which may be attributed to improper folding of the truncated TssM-T25 fusion (Supplementary Figure 1H). Fluorescence microscopy analysis further revealed co-localization of Fha foci with TssL and TssG in AAC00-1 (Figure 1 D and E).

These results collectively indicate that the cytoplasmic protein Fha promotes the assembly of both the membrane complex and the baseplate, presenting a distinct T6SS assembly pathway that differs from the previously established model initiated by the OM protein TssJ (7).

### Fha forms LLPS condensates *in vitro* and *in vivo*

We observed the formation of spherical droplets of purified Fha_sfGFP proteins but not sfGFP control in a dextran-70 supplemented buffer, which serves as a volume-excluding polymer to mimic intracellular environments (Figure 2A) (28). We also found droplet formation and dynamic fusion using purified untagged Fha proteins (Figure 2B, Supplementary Figure 2A). Individual Fha_sfGFP droplets subjected to photobleaching experiments exhibited partial fluorescence recovery within minutes (Figure 2 C and D), suggesting the rapid exchange of Fha_sfGFP between the droplets and the surrounding solution.

**Fig. 2.**
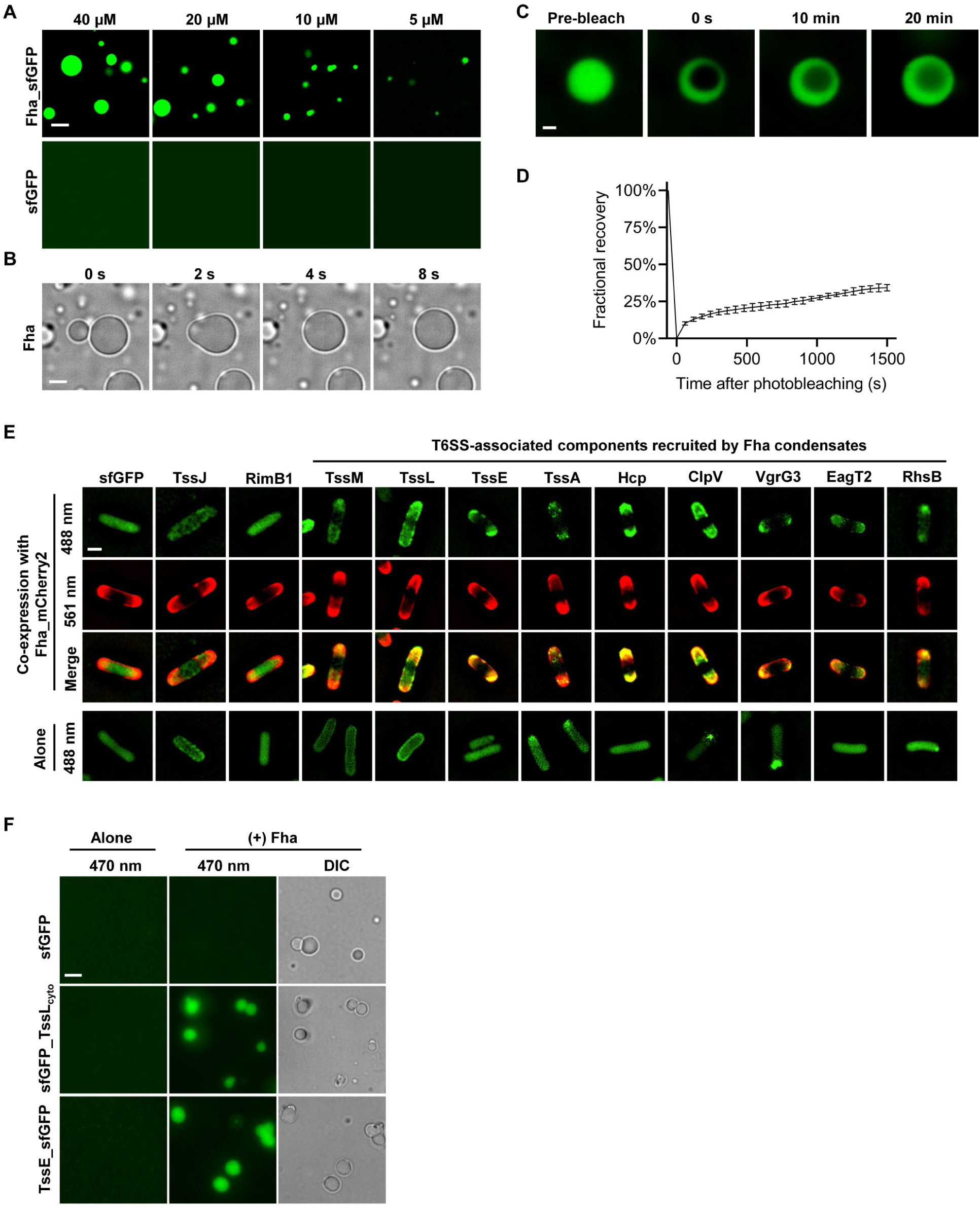
Fha undergoes LLPS both *in vitro* and *in vivo*. **A**, Fluorescence images showing Fha_sfGFP underwent LLPS at the indicated concentrations. The sfGFP was used as a negative control. **B**, DIC images showing time-lapse fusion of two individual Fha droplets. A final concentration of 40 µM Fha proteins in a 5% dextran-70 solution was employed. **C**, Representative fluorescence recovery of a photobleached Fha_sfGFP droplet. A representative 8- ×8-μm field is shown. Scale bar: 1 μm. **D**, Quantification of the FRAP analyses for 3 different Fha_sfGFP droplets. **E**, Fluorescence microscopy images showing Fha_mCherry2 condensates recruited a number of T6SS-associated client proteins in *E. coli* BL21(DE3). The sfGFP and sfGFP-labeled TssJ, RimB1, TssM, TssL, TssE, TssA, Hcp, ClpV, VgrG3, EagT2, and RhsB were cloned on a constitutive-expression vector pBBR1MCS2, respectively. To mitigate the impact of the C-terminal toxin domain and self-cleavage of RhsB, RhsB^NM^ ^D280A^ mutant were used here. The sfGFP and sfGFP-labeled T6SS-associated proteins were expressed alone or co-expressed with Fha_mCherry2 in *E. coli* BL21(DE3). A representative 5- × 5-μm field of cells is shown. Scale bar: 1 μm. **F**, Fluorescence microscopy images showing Fha recruited TssL cytoplasmic domain and TssE via LLPS *in vitro*. Fha proteins were mixed with sfGFP, sfGFP_TssL_cyto_ or TssE_sfGFP at a ratio of 10: 1 in a 5% dextran-70 solution before imaging. For **A**, **B**, and **F**, a representative 30- ×30-μm field is shown. Scale bar: 5 μm.

Consistent with the *in vitro* results, Fha_sfGFP formed polar condensates when expressed in *E. coli* BL21(DE3) (Supplementary Figure 2B). After photobleaching a single focus, the polar fluorescence signals of Fha_sfGFP were partially restored within minutes (Supplementary Figure 2 B and C). These findings collectively suggest that Fha undergoes LLPS both *in vitro* and *in vivo*.

### Fha condensates recruit T6SS-associated components

To test whether Fha condensates could recruit other T6SS proteins, we expressed sfGFP-labeled components alone or co-expressed them with Fha_mCherry2 in *E. coli* BL21(DE3). Dispersed distribution was observed for membrane proteins TssJ, TssM, and TssL, and cytoplasmic proteins TssE, TssA, Hcp, the chaperone protein EagT2 and immunity protein RimB1 of T6SS effector RhsB (Figure 2E, Supplementary Figure 2D). Interestingly, some cells formed foci when expressing sfGFP-labeled VgrG3 or ClpV (Figure 2E, Supplementary Figure 2D). Co-expression with Fha_mCherry2 led to co-localization of all tested proteins except TssJ, RimB1, and the negative control sfGFP (Figure 2E, Supplementary Figure 2D). Intriguingly, TssA exhibited small foci inside the Fha condensates, suggesting TssA proteins might self-polymerize and occupy separate regions within the condensates (Figure 2E, Supplementary Figure 2D).

We further investigated whether Fha condensates can recruit T6SS-associated proteins *in vitro*. Using purified sfGFP-labeled TssL cytoplasmic domain (sfGFP_TssL_cyto_) and TssE (TssE_sfGFP), we found that the otherwise dispersed sfGFP_TssL_cyto_ or TssE_sfGFP formed droplets when Fha was added while the GFP control did not (Figure 2F, Supplementary Figure 3A), suggesting that Fha could recruit these two T6SS proteins, independent of other proteins.

### FHA domain and IDR dictate LLPS of Fha

To investigate the mechanism of Fha-mediated LLPS, we employed AlphaFold2 to predict the Fha structure (Supplementary Figure 3B). Our analysis revealed three distinct regions: an N-terminal Forkhead-associated (FHA) domain, an intrinsically disordered region (IDR), and a C-terminal domain featured with five alpha helices (Figure 3A, Supplementary Figure 3B). To determine the contributions of these regions to LLPS, we first constructed Fha truncation mutants and expressed them in *E. coli* BL21(DE3). The sfGFP-labeled mutants, Fha^N^ (1-101 aa) and Fha^IDR^ (102-561 aa) formed polar localized condensates at cell poles, whereas the Fha^C^ mutant (562-777 aa) displayed a membrane-associated distribution (Figure 3A, Supplementary Figure 3C). Photobleaching experiments showed partial fluorescence recovery within minutes for both Fha^N^ and Fha^IDR^ (Figure 3B, Supplementary Figure 3 D and E). Notably, Fha^IDR^ exhibited a slower rate of fluorescence recovery compared to full-length Fha and Fha^N^ (Figure 3B). Besides, Fha^IDR^ and Fha^ΔC^ exhibited one condensate per cell, whereas Fha^N^, Fha^ΔN^, and Fha^ΔIDR^ formed two condensates per cell (Figure 3A, Supplementary Figure 3C). These findings suggest that both the N-terminal FHA domain and the IDR can mediate LLPS, but they exhibit divergent properties affecting condensate merging. The C-terminal domain preferentially localized to the membrane and modulated the effect of Fha^IDR^, resulting in two condensates in cells (Figure 3A, Supplementary Figure 3C). The membrane association might be attributed to an amphipathic helix (APH) found in one of the five alpha-helices within the C-terminal domain (Supplementary Figure 3B) (29).

**Fig. 3.**
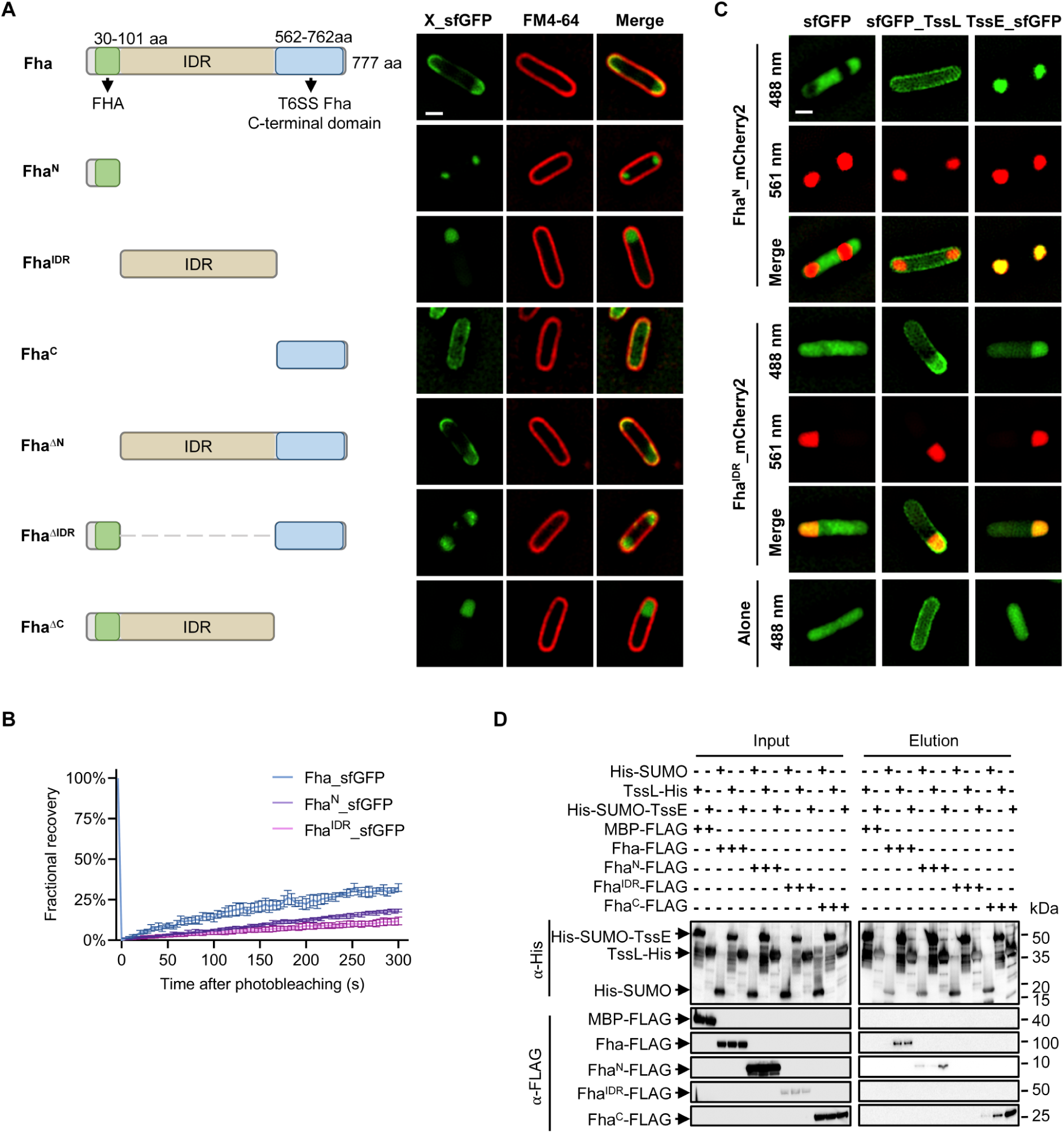
The FHA domain and IDR form distinct condensates. **A**, Fluorescence microscopy images showing the subcellular localization of Fha_sfGFP and its truncated mutants in *E. coli* BL21(DE3). Cells were stained with FM 4-64 before imaging. **B**, Quantification of the FRAP analyses for Fha_sfGFP, Fha^N^_sfGFP, or Fha^IDR^_sfGFP foci in *E. coli* BL21(DE3). **C**, Fluorescence microscopy images showing the co-localization of Fha^N^_mCherry2 or Fha^IDR^_mCherry2 with sfGFP, sfGFP_TssL, or TssE_sfGFP in *E. coli* BL21(DE3). The sfGFP and sfGFP-labeled TssL and TssE were expressed alone or co-expressed with Fha^N^_mCherry2 or Fha^IDR^_mCherry2 in *E. coli* BL21(DE3), respectively. For **A** and **C**, a representative 5- × 5-μm field of cells is shown. Scale bar: 1 μm. **D**, Interaction of Fha, Fha^N^, Fha^IDR^, and Fha^C^ with TssL, and TssE. Pull-down analysis was performed using His-tagged SUMO (control), TssL, or SUMO-TssE, and FLAG-tagged MBP (control), Fha, Fha^N^, Fha^IDR^, or Fha^C^.

To determine key residues that may control the LLPS formation of Fha, we performed sequence alignment of *A. citrulli* Fha with 45 additional Fha homologs (Supplementary Data 1). We mutated 42 conserved uncharged amino acids to serine, 3 conserved acidic, and 2 basic amino acids to asparagine. However, none of the tested single or combinatorial residue changes abolished condensate formation of Fha_sfGFP when they were expressed in *E. coli* BL21(DE3) (Supplementary Figure 3F).

### Fha^N^ and Fha^IDR^ condensates selectively recruit client proteins

We next tested the co-localization of mCherry2-labeled Fha^N^ and Fha^IDR^ variants with sfGFP-labeled IM component TssL and baseplate protein TssE, using sfGFP as control. Co-localization experiments showed that Fha^IDR^ and Fha^N^ condensates recruited TssE, while IM protein TssL only co-localized with Fha^IDR^ (Figure 3C, Supplementary Figure 4A). Moreover, Fha^N^ excluded sfGFP from the condensates whereas the Fha^IDR^ did not (Figure 3C, Supplementary Figure 4A). To corroborate the imaging results, we performed pull-down assays testing the interaction of Fha domains and found that Fha^N^ interacted with TssE but not TssL (Figure 3D), which is consistent with the imaging data. Fha^C^ showed interaction with both TssE and TssL (Figure 3D). Due to poor expression of Fha^IDR^, its pull-down result was inconclusive. These findings suggest that all three domains of Fha are involved in protein interactions and Fha^N^ and Fha^IDR^ condensates can selectively recruit and exclude specific client proteins.

### The Fha IDR is crucial for T6SS activities in AAC00-1

We next tested the contributions of the FHA domain and IDR to the assembly of T6SS membrane complex, baseplate and sheath structures in AAC00-1. We first verified that introducing chromosomal TssB_mCherry2 into the *sfGFP_tssL* strains did not affect T6SS-mediated killing abilities (Supplementary Figure 4 B and C). Fluorescence microscopy analyses revealed that the deletion of the FHA domain (*fha^ΔN^*) or the IDR (*fha^ΔIDR^*) resulted in a reduction in foci formation when compared to the full-length Fha (Figure 4 A and B). Additionally, the *fha^ΔN^* mutant displayed a decreased number of foci for TssB, TssL, and TssG, while the *fha^ΔIDR^* mutant exhibited a decreased number of TssG foci and lacked detectable TssB and TssL foci. (Figure 4 A, C, and D, Supplementary Figure 4D). Bacterial competition assay confirmed that, while the *fha^ΔN^*mutant exhibited impaired but not abolished T6SS-mediated killing against *E. coli*, the *fha^ΔIDR^* mutant failed to outcompete *E. coli* (Figure 4E, Supplementary Figure 4E). Plasmid-borne Fha could partially complement the T6SS-mediated killing abilities of both *fha^ΔN^* and *fha^ΔIDR^* mutants (Supplementary Figure 4 F and G). These results underscore the indispensable role of Fha IDR in controlling T6SS assemblies in AAC00-1.

**Fig. 4.**
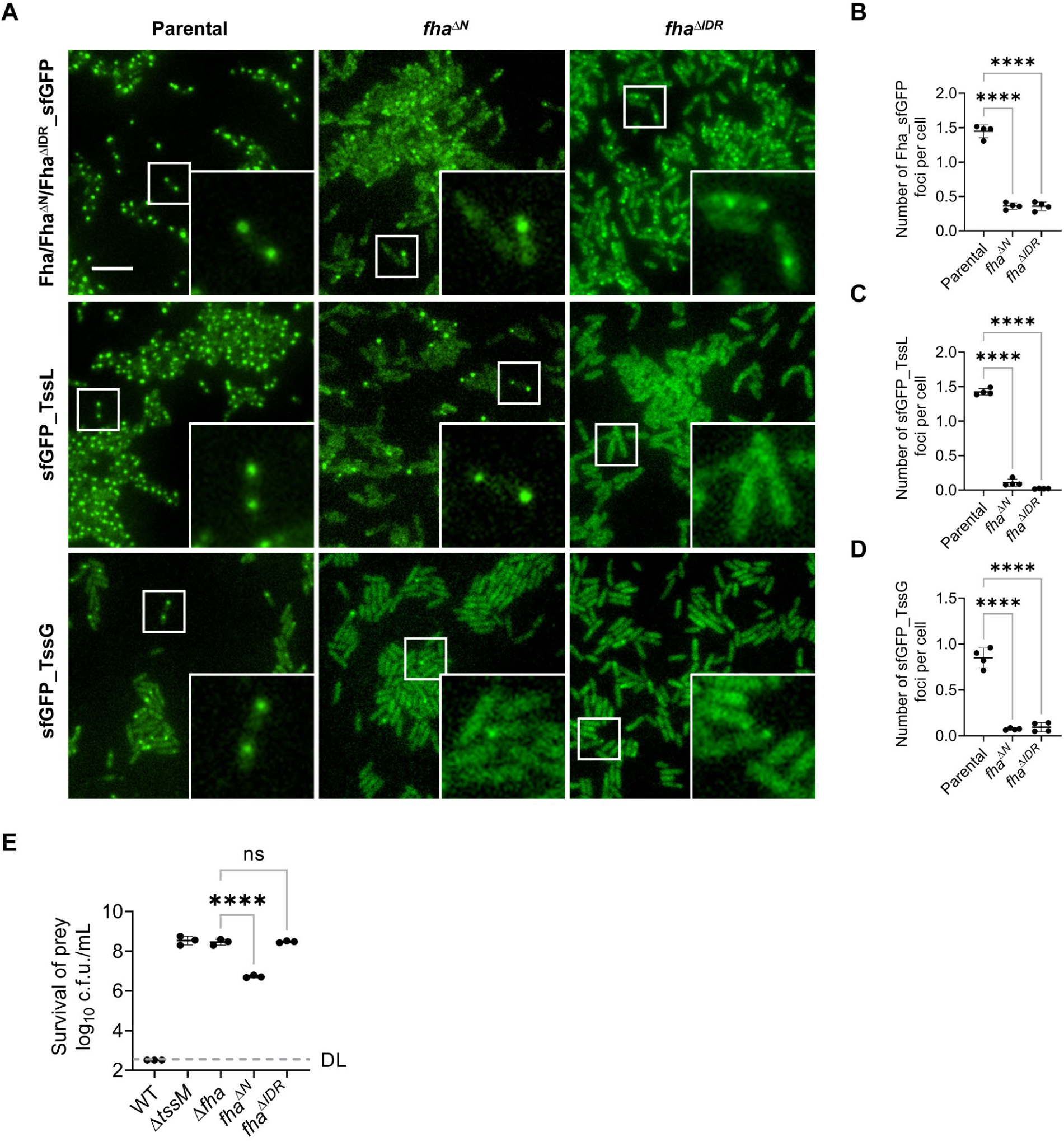
Fha IDR is indispensable for T6SS assembly in AAC00-1. **A**, Fluorescence microscopy images showing the signals of chromosomal sfGFP-labeled Fha (or its mutants), TssL, and TssG in AAC00-1 Parental, *fha*^Δ*N*^, and *fha*^Δ*IDR*^. A representative 30- × 30-μm field of cells with a 3× magnified 5- × 5-μm inset (marked by box) is shown. Genotypes are indicated at the top. Scale bar: 5 μm. Quantification of the formed Fha_sfGFP (**B**), sfGFP_TssL (**C**), sfGFP_TssG (**D**) foci in AAC00-1 Parental, *fha*^Δ*N*^, and *fha*^Δ*IDR*^. **E**, Competition analysis of the AAC00-1 wild type (WT), T6SS-null mutant Δ*tssM*, *fha*^Δ*N*^, and *fha*^Δ*IDR*^ mutants. Killer strains are indicated and the prey strain is the *E. coli* MG1655 carrying pPSV37-sfGFP plasmid. Cells of killer and prey were mixed at a ratio of 10:1 (killer: prey) and co-incubated for 3 h at 37 °C. Survival of killer strains during competition assays is depicted in Supplementary Figure 4E. For B to E, error bars indicate the mean +/- standard deviation of at least three biological replicates and statistical significance was calculated using One-way ANOVA test for each group. \*\*\*\**P* < 0.0001. ns, not significant. DL, detection limit.

### The C-terminal APH is indispensable for T6SS assemblies in AAC00-1

To determine the role of the C-terminal APH in Fha membrane-associated localization, we introduced specific amino acid substitutions within the APH region of the Fha^C^_sfGFP. We constructed three APH mutants with varying net charges and reduced hydrophilicities, while preserving the predicted α-helix (Figure 5A, Supplementary Figure 5A). Specifically, the APH mut^N^ retains a neutral net charge, similar to its parental, while the APH mut^-4^ and APH mut^+4^ exhibit net charges of -4 and +4, respectively (Figure 5A, Supplementary Figure 5A). By expressing these Fha^C^_sfGFP variants in *E. coli*, we found that all three mutants exhibited cytosolic distribution while the wild-type Fha^C^_sfGFP was membrane-associated, suggesting that the membrane association is mediated by the APH (Figure 5A, Supplementary Figure 5A). Next, we examined the effect of the APH on the T6SS activities of AAC00-1 by introducing the APH residue mutations into the chromosomal encoded Fha. Although the chromosomal APH mutations resulted in only a slight decrease in the number of Fha foci per cell, these mutations abolished the formation of T6SS membrane complexes and baseplates, sheath assemblies, and T6SS-mediated killing (Figure 5 B-E, Supplementary Figure 5 B-D). These mutants could be partially complemented through the introduction of a plasmid-borne Fha (Supplementary Figure 5 B and C). Importantly, the APH mutation in full-length Fha did not disrupt its interactions with TssE and TssL *in vitro*, which might be attributed to the complexity of the interactions involving Fha and its interacting proteins (Supplementary Figure 5E). However, when the APH mutation was introduced in the Fha^C^ truncation, a significant reduction in protein levels was observed, rendering the interaction results less definitive (Supplementary Figure 5E). The C-terminal domain of Fha homologs in *V. cholerae* V52 and *P. aeruginosa* PAO1 also contain predicted APHs, suggesting the APHs are conserved (Supplementary Figure 5F).

**Fig. 5.**
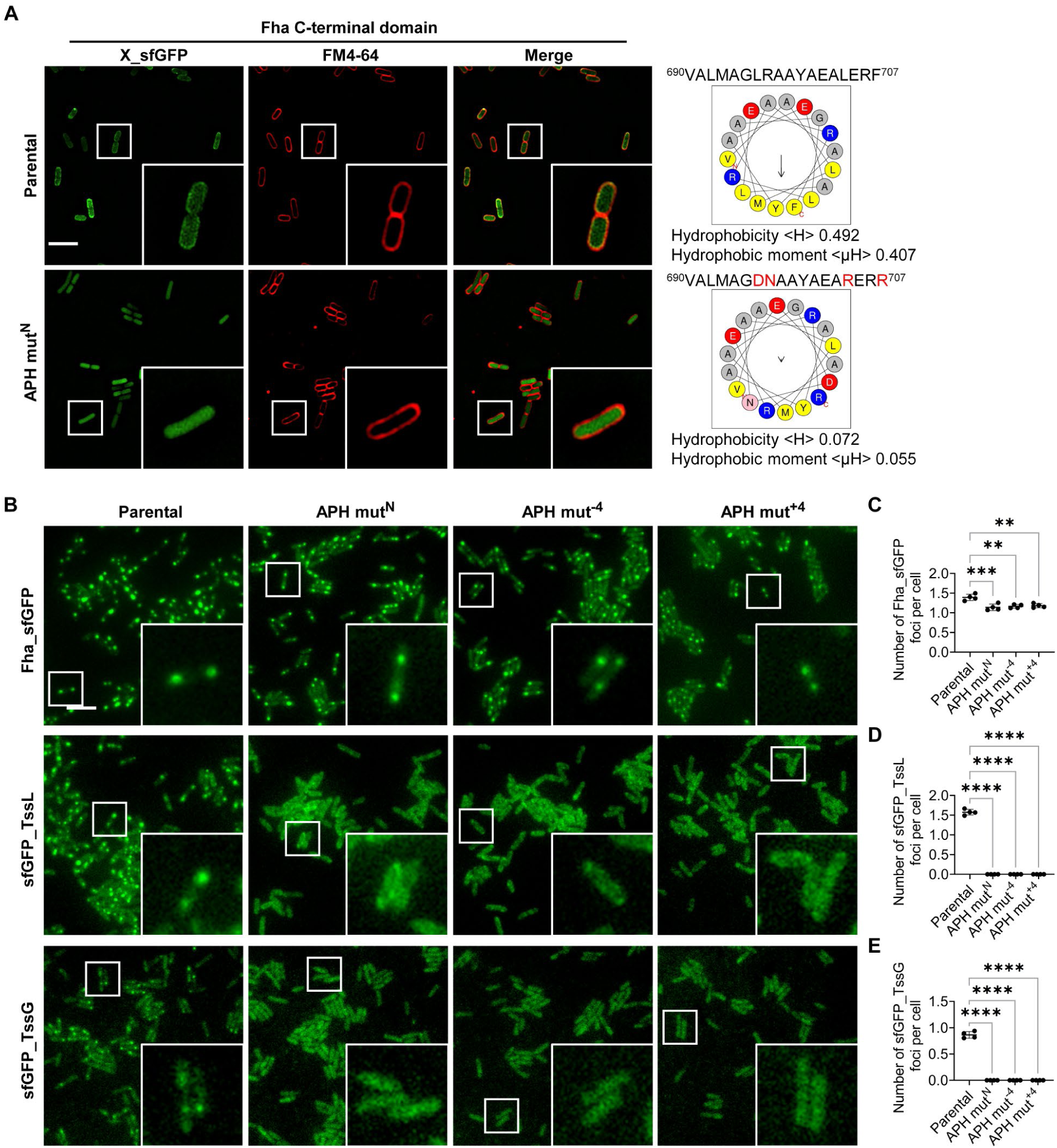
The Fha C-terminal APH is essential for T6SS activities in AAC00-1. **A**, Fluorescence microscopy images showing the subcellular localization of Fha^C^_sfGFP and its mutant in *E. coli* BL21(DE3). Cells were stained with FM 4-64 before imaging. The predicted Fha C-terminal APH (from 690 to 707 aa) and its mutant were subjected to hydrophobicity, hydrophobic moment, and amino acid composition analysis using HeliQuest. Yellow, hydrophobic residues; blue, basic; red, acidic; pink, asparagine and glutamine; gray, other residues. **B**, Fluorescence microscopy images showing the signals of chromosomal sfGFP-labeled Fha (or its mutants), TssL, and TssG in AAC00-1 Parental and its APH mutants. Genotypes are indicated at the top. Scale bar: 5 μm. Quantification of the formed Fha_sfGFP (**C**), sfGFP_TssL (**D**), and sfGFP_TssG (**E**) foci in AAC00-1 Parental and its APH mutants. For **A** and **B**, a representative 30- ×30-μm field of cells with a 3×magnified 5- ×5-μm inset (marked by box) is shown. Scale bar: 5 μm.

### Fha functions are conserved in other species

Through analyzing 167 distinct T6SS clusters extracted from the SecReT6 database (30), we identified Fha exclusively in T6SSi (Supplementary Data 2) (31). Phylogenetic analysis, based on the sequences of representative TssC homologs (Supplementary Data 3), revealed a widespread presence of Fha in T6SSi1, T6SSi3, and T6SSi5, but not in T6SSi2, T6SSi4a, and T6SSi4b (Figure 6A). To test the T6SS-initiating and LLPS functions of Fha homologs, we expressed C-terminal sfGFP-labeled Fha1 (Fha1^PA^_sfGFP) from *P. aeruginosa* PAO1 and Fha (Fha^VC^_sfGFP) from *V. cholerae* V52 in *E. coli* BL21(DE3). Both proteins formed polar condensates and their fluorescence signals were partially recovered within minutes after photobleaching, suggesting these two Fha homologs could also form LLPS condensates (Figure 6 B and C, Supplementary Figure 6 A and B).

**Fig. 6.**
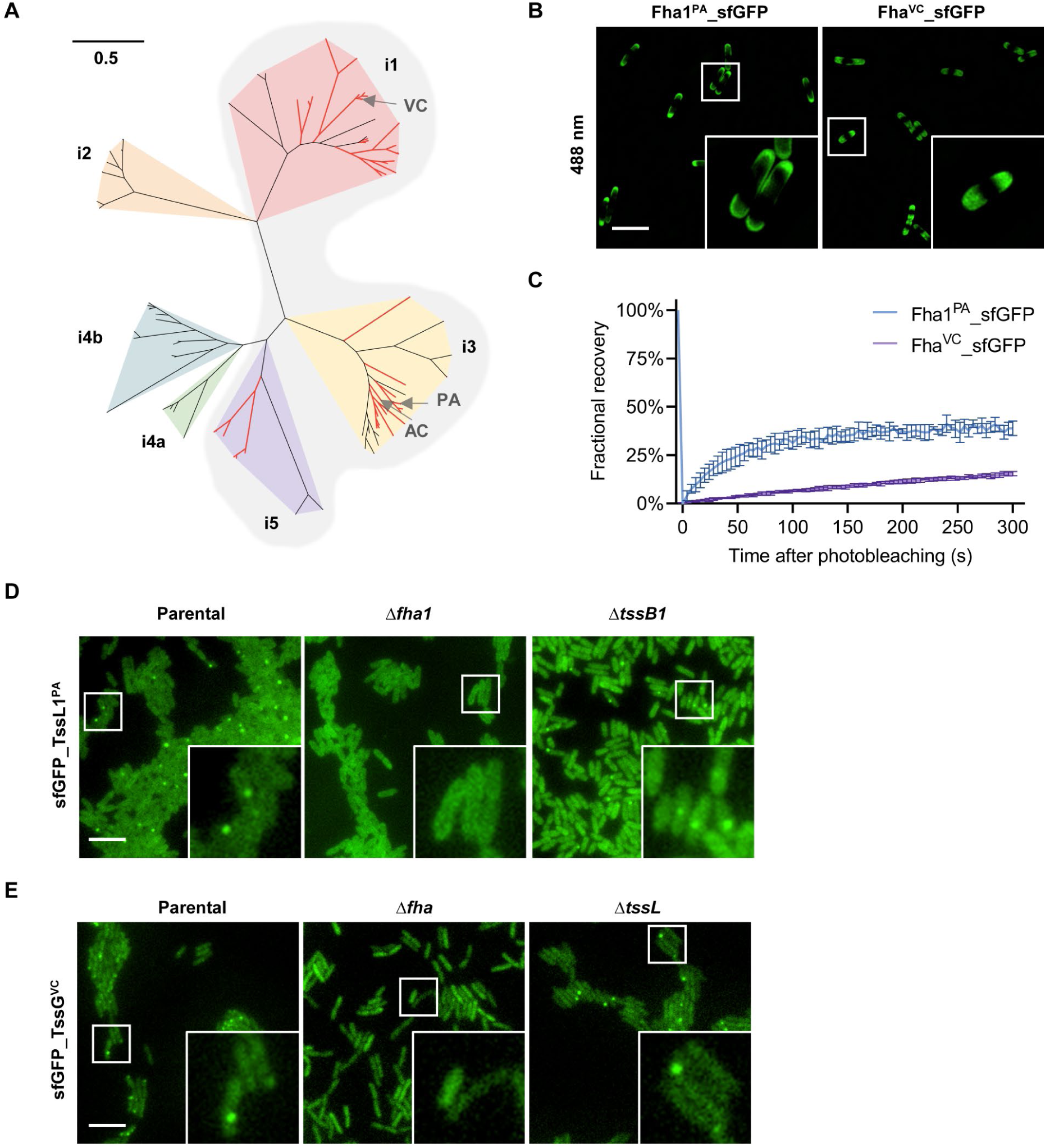
Fha homologs are functionally conserved. **A**, Maximum likelihood (ML) phylogenetic tree generated from a partial alignment of 154 representative TssC sequences spanning the diversity present in T6SSi1, T6SSi2, T6SSi3, T6SSi4a, T6SSi4b, and T6SSi5, as indicated. The corresponding T6SS clusters containing Fha homologs are highlighted with red branches, while the T6SS subtypes with Fha homologs are shaded in gray. The positions of T6SS in *A. citrulli* AAC00-1, *P. aeruginosa* (H1-T6SS), and *V. cholerae* are indicated by gray arrows, denoted as AC, PA, and VC, respectively. The scale bar represents the number of amino acid changes per site. **B**, Fluorescence microscopy images showing the subcellular localization of Fha1^PA^_sfGFP and Fha^VC^_sfGFP in *E. coli* BL21(DE3). **C**, Quantification of the FRAP analyses for Fha1^PA^_sfGFP or Fha^VC^_sfGFP in *E. coli* BL21(DE3). **D**, Fluorescence microscopy images showing sfGFP_TssL1 localization in PAO1 Parental, Δ*fha1*, and Δ*tssB1*. **E**, Fluorescence microscopy images showing sfGFP_TssG localization in V52 Parental, Δ*fha*, and Δ*tssL*. For **B**, **D** and **E**, a representative 30- ×30-μm field of cells with a 3× magnified 5- ×5-μm inset (marked by box) is shown. Genotypes are indicated at the top. Scale bar: 5 μm.

We next tested the effect of Fha on T6SS protein localization and assembly. In PAO1, deleting *fha1^PA^* led to a diffuse distribution of sfGFP_TssL1 and TssB1_mCherry2 while the Δ*tssB1* control sample exhibited sfGFP_TssL1 foci (Figure 6D, Supplementary Figure 6C), suggesting Fha1^PA^ is essential for H1-T6SS membrane complex assembly. In *V. cholerae* V52, we found that the sfGFP_TssL^VC^ labeling impaired T6SS activities in V52 (Supplementary Figure 6D). Therefore, we added the sfGFP tag to chromosomally encoded TssG^VC^ and confirmed that the sfGFP_TssG^VC^ strain could actively assemble T6SS structures (Supplementary Figure 6E). Imaging results show that deletion of the *fha^VC^* abolished TssG foci formation while deletion of the *tssL^VC^* did not, suggesting that Fha but not the intact inner-membrane complex was required for TssG foci formation (Figure 6E, Supplementary Figure 6E). These findings suggest the LLPS capability of Fha homologs and their role in initiating T6SS assembly might be conserved.

## Discussion

Our study unveils a distinct inside-first model for T6SS assembly in *A. citrulli*, initiated by the cytoplasmic protein Fha that directs the formation of IM complex and baseplate (Figure 7), in contrast to the OM TssJ-initiated model established in *E. coli* (7). Fha not only directly interacts with the IM complex and baseplate proteins but also forms LLPS condensates that recruit a priming factor TssA, secreted proteins Hcp, VgrG, effector protein RhsB and its cognate chaperone protein EagT2. Importantly, Fha-mediated LLPS formation and T6SS assembly are conserved in *P. aeruginosa* and *V. cholerae*, suggesting LLPS might be a general mechanism for macromolecular assembly. Considering the widespread distribution of T6SS among bacterial pathogens and its diverse functions in interspecies interactions and pathogenesis, our findings highlight the broad impact of LLPS on bacterial physiological processes far beyond intracellular activities including gene regulation and division.

**Fig. 7.**
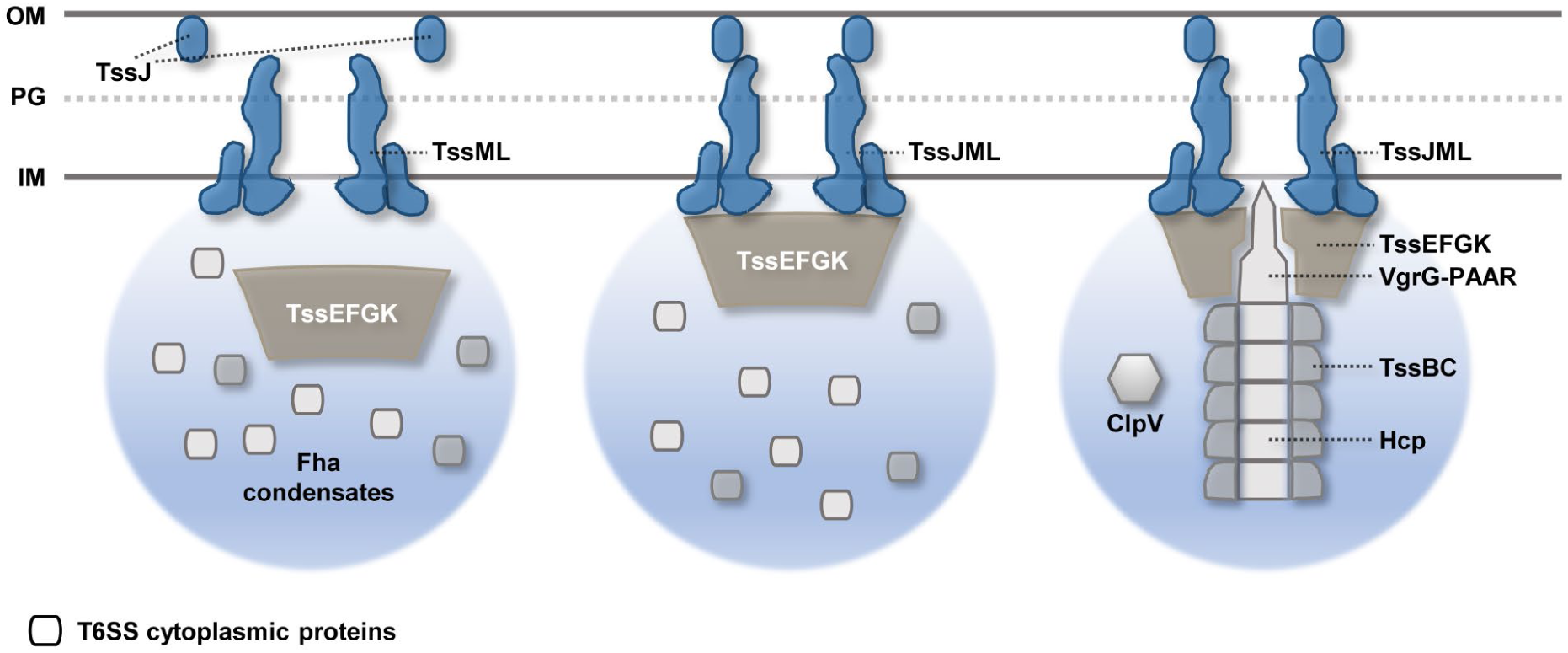
Schematic representation of the inside-first LLPS-driven model for T6SS assembly in AAC00-1. In *A. citrulli* AAC00-1, the cytoplasmic Fha protein undergoes LLPS, giving rise to condensates that selectively recruit T6SS-associated components. The assembly of IM TssML complex and baseplate likely occurs within these Fha condensates independently. The TssML complex then interacts with the baseplate and recruits the OM TssJ, forming an assembling platform that supports the elongation of the Hcp tube and TssBC sheath. IM, inner membrane. PG, peptidoglycan. OM, outer membrane.

T6SSs are grouped into distinct categories based on sequence divergence (31–34), but the conservation of their assembly processes remains unclear. The prevailing model, exemplified by the non-Fha-encoding T6SS in enteroaggregative *E. coli*, suggests the OM protein TssJ recruits and assembles the IM complex as the initial step of T6SS (7). However, TssJ is not required for T6SS functions in *Serratia marcescens* (35), and a TssJ homolog is missing in *Acinetobacter* species (36). Although these results raise questions about the TssJ-initiated model, the requirement of an OM anchor for all T6SSs, except for the noncanonical T6SSiv in *Amoebophilus asiaticus* (33), implies that TssJ-independent T6SSs may still follow an outside-first model using other OM anchor proteins. The discovery of Fha-mediated assembly provides a distinct model for T6SS initiation, which is likely conserved among Fha-containing T6SSs. Our data suggest that Fha may serve dual functions: acting as a key scaffold that brings T6SS components together and as a molecular sieve to prevent interference from other cytosolic proteins during T6SS assembly. The latter is supported by the observations that the Fha condensates recruited the effector RhsB but did not recruit its immunity protein RimB1 or the cytosolic protein sfGFP (Figure 2E).

Interestingly, membrane complex and baseplate could assemble independently of each other in AAC00-1. In *P. aeruginosa* PAO1, it was observed that the baseplate component TssK1 could form foci independently of the membrane protein TssM1 (37). Our previous study showed that *V. cholerae* V52 could form long tubular structures without the presence of membrane complex or baseplate components using a non-contractile model (38). From an evolutionary perspective, the IM complex shares homology with the type IV secretion apparatus, whereas the baseplate and the tubular structure share homology with contractile phage tails (1, 39, 40). It remains elusive how the structurally and evolutionarily independent membrane complex and baseplate are recruited to form the T6SS structure during evolution. Whether Fha acts as a structural component of the T6SS in AAC00-1 also warrants further investigation. However, the absence of Fha in the cryo-ET T6SS structure of *V. cholerae* suggests Fha is more likely to facilitate structure assembly rather than being a structural component itself (41).

FHA domains are phosphor-peptide recognition modules found in both bacteria and eukaryotes, exhibiting a high specificity for phosphorylated threonine residues (42). These domains have been identified in numerous proteins with crucial functions in DNA damage response, cell growth, signal transduction, cell cycle regulation, and pathogenesis (43, 44). However, LLPS formation of the FHA domain-containing protein has only been shown in a membrane-bound ABC transporter Rv1747 in *M. tuberculosis*, whose cytoplasmic fragment Rv1747^1–310^, containing the FHA-domain, displays LLPS properties and recruits a kinase and a phosphatase (21). The T6SS assembly mediated by Fha in AAC00-1 is much more complex, involving stepwise and orchestrated assembly of the membrane complex and the cytoplasmic structures. In *P. aeruginosa*, Fha1 is controlled by a TPP (threonine phosphorylation) signal transduction pathway, involving a kinase PpkA and a phosphatase PppA, for T6SS activation (45). Interestingly, in *Agrobacterium tumefaciens*, PpkA phosphorylates TssL but not Fha (46). While AAC00-1 possesses PpkA and PppA homologs, its T6SS is active during routine growth in LB media (6). Furthermore, LLPS of Fha in *E. coli* and *in vitro* suggests that the Fha forms condensates independent of the phosphorylation. The Fha in *V. cholerae* V52, which has a constitutively active T6SS that lacks TPP components, also exhibited LLPS condensates. Therefore, whether Fha functions and its LLPS are modulated by the TPP pathway in AAC00-1 is to be addressed by future research.

In summary, our findings reveal a distinct LLPS-mediated T6SS-assembly model. These findings will prompt further investigations into the role of LLPS in facilitating the assembly of other macromolecular structures in bacteria, as well as the evolutionary significance of Fha-mediated LLPS for assembly. Through T6SS-mediated divergent functions, our findings also demonstrate a broader impact of LLPS on bacterial physiology, such as pathogenesis and interspecies interactions. These results may also lead to future therapeutic interventions targeting LLPS to reduce T6SS-mediated functions and control infections as alternatives to antibiotic treatment.

## Materials and Methods

### Bacterial strains, plasmids, and growth conditions

Strains were grown in LB media following standard culturing conditions. Antibiotics were used at the following concentrations: kanamycin (50 µg/ml), gentamicin (10 µg/ml for AAC00-1, and 20 µg/ml for PAO1), streptomycin (100 µg/ml), irgasan (25 µg/ml) and carbenicillin (50 µg/ml). All plasmids were constructed by Gibson assembly and verified by Sanger sequencing. Precise deletion mutants were constructed by using standard homologous recombination method. For mutant construction, the pEXG2.0 suicide vector was used in AAC00-1 and PAO1 (47), and the pDS132 suicide vector in V52 (48). Plasmids, strains, and primers used in this study are listed in Supplementary Table 1.

### Bacterial two-hybrid assay

As previously described, the N-terminus or C-terminus of different proteins were fused with the T18 and T25 split domains of the *Bordetella* adenylate cyclase (47). The two plasmids encoding corresponding proteins were co-transformed into the reporter strain BTH101 Δ*cyaA*. The single colonies for each transformation were inoculated into 300 μl of liquid LB. After 5 to 6 h growth at 30 °C, 3 μl of each culture were spotted onto LB-agar plates with carbenicillin, kanamycin, IPTG (0.02 mM), and X-Gal (80 µg/ml) and incubated at room temperature for 16 h. The experiment was done two times and a representative result is shown.

### Fluorescence microscopy of T6SS assembly

AAC00-1 strains were grown overnight in liquid LB media to OD_600_ ∼ 2 at 28°C. The cultures were then centrifuged at 4,500 ×*g* for 2 min, resuspended in liquid LB media to OD_600_ ∼ 10, and spotted onto 1% agarose-0.5 × PBS pads. The Nikon Ti2-E inverted microscope was used for imaging, which equipped with the Perfect Focus System and a CFI Plan Apochromat DM 100X

Oil objective lens. ET-GFP (Chroma #49002) and ET-mCherry (Chroma #49008) filter sets were used for fluorescence excitation. All imaging experiments were done at least two times and a representative result is shown.

### *In vitro* liquid–liquid phase separation (LLPS) assay

The purified Fha and Fha_sfGFP proteins were dialyzed against LLPS buffer (10 mM Na_2_HPO_4_/NaH_2_PO_4_, 200 mM NaCl, pH 7.4) and subsequently concentrated using centrifugal filter units with Ultracel-30K membranes, resulting in a final concentration of approximately 80 µM. To prepare the samples for imaging, the concentrated proteins were diluted in LLPS buffer through a 2-fold serial dilution. Each sample was then mixed with an equal volume of 20% dextran-70 in LLPS buffer unless stated otherwise. The resulting mixtures were loaded onto glass. The ZEISS LSM 980 with Airyscan 2 microscope, equipped with a 63×/1.40 oil objective lens and a laser of 488 nm, and the Nikon Ti2-E inverted microscope, equipped a 100×/1.40 oil objective lens and an ET-GFP (Chroma #49002) filter set, were used for imaging. All imaging experiments were done at least two times and a representative result is shown.

### Fluorescence recovery after photobleaching (FRAP)

For the *in vitro* FRAP analysis, the Fha_sfGFP droplets formed with purified proteins and dextran-70 were loaded onto glass slides. Image acquisition was performed using the ZEISS LSM 980 with Airyscan 2 microscope, equipped with a 63×/1.40 oil objective lens and a laser of 488 nm. The photobleaching was initiated by irradiation with 100% laser power onto the selected regions for 10 ms. For the *in vivo* FRAP analysis, the *E. coli* BL21(DE3) strains were grown in liquid LB media to OD_600_ ∼ 0.6 and induced with 0.2 mM IPTG for 40 min at 37°C. The cultures were then centrifuged at 4,500 ×*g* for 2 min, resuspended in liquid LB media, and spotted onto 1% agarose-0.5 × PBS pads. Image acquisition was performed using the Nikon Ti-E A1R HD25+SIM S microscope, which was equipped with the Perfect Focus System and a CFI SR HP Apochromat TIRF 100XC objective lens, along with a laser of 488 nm. The photobleaching was initiated by irradiation with 20% laser power onto the selected regions for 1 s. Fluorescence recovery was assessed by normalizing the fluorescence intensity within the bleached region to an adjacent unbleached area using the “FRAP_profiler_V2” Fiji plugin. The pre-bleaching region served as the reference at 100%, and fluorescence recovery was quantified at each time point. All imaging experiments were done two times and a representative result is shown.

### Co-localization assay in *E. coli*

The N-terminus or C-terminus of different T6SS-associated proteins were fused with sfGFP based on pBBR1MCS2 vector, respectively. The sfGFP-labeled proteins were then transferred into *E. coli* BL21(DE3) alone or co-transferred with mCherry2-labeled Fha or it truncated mutants based on pET22b vector as indicated. *E. coli* BL21(DE3) strains carrying one or two plasmids were grown in liquid LB media to OD_600_ ∼ 0.6 and induced with 0.2 mM IPTG for 2 h at 37°C. The cultures were then centrifuged at 4,500 ×*g* for 2 min, resuspended in liquid LB media, and spotted onto 1% agarose-0.5 × PBS pads. Image acquisition was performed using the Nikon Ti-E A1R HD25+SIM S microscope, which was equipped with the Perfect Focus System and a CFI SR HP Apochromat TIRF 100XC objective lens, along with the 488 and 561 nm excitation lasers. The 3D-SIM imaging was conducted. All imaging experiments were done two times and a representative result is shown.

### Bacterial competition assay

Details are in Supplementary Materials and Methods.

### Western blotting analysis

Details are in Supplementary Materials and Methods.

### Protein secretion assay

Details are in Supplementary Materials and Methods.

### Protein pull-down assay

Details are in Supplementary Materials and Methods.

### Protein purification

Details are in Supplementary Materials and Methods.

### Bioinformatic analysis

Details are in Supplementary Materials and Methods.

## Acknowledgments

This work was supported by funding from National Key R&D Program of China (2020YFA0907200, 2018YFA0901200), National Natural Science Foundation of China (32030001). We thank Yang Fu, Kuo Zhang, and Jin-Sheng Liu for providing materials and helpful discussions. The funders had no role in study design, data collection and interpretation, or the decision to submit the work for publication.

## Author contributions

T.D. conceived the project. T.P., Y.A., X.W., H.Luo., Y.K., H.Li., M.T., Z.Y., J.L., H.Z., Z.W, and X.Liang. performed research. T.J., M.Z., and X.Liu. provided key reagents and materials. T.P. and T.D. wrote the manuscript.

---

**Supplementary Figure 1. Fha promotes the T6SS biogenesis through direct interactions. A**, Competition analysis of the fluorescence-labeled AAC00-1 strains. Killer strains are indicated and the prey strain is the *E. coli* MG1655 carrying pPSV37-sfGFP plasmid. The wild type (WT) and T6SS-null mutant Δ*tssM* serve as positive and negative control, respectively. Survival of killer strains during competition assays is depicted in **B**. **C**, Secretion analysis of Hcp in AAC00-1 wild type (WT), Δ*fha*, Δ*tssJ*, Δ*tssM*, Δ*tssF*, Δ*tssA*, Δ*tssB*, and Δ*hcp*. RpoB serves as a control for equal loading and autolysis. **D**, Fluorescence microscopy images showing sfGFP_TssM localization in AAC00-1 Parental, Δ*tssJ*, and Δ*fha*. A representative 30- × 30-μm field of cells with a 3× magnified 5- × 5-μm inset (marked by box) is shown. Genotypes are indicated at the top. Scale bar: 5 μm. **E**, Competition analysis of the AAC00-1 WT, Δ*tssM*, Δ*fha*, and Δ*fha* mutant complemented with a plasmid-borne Fha. Killer strains are indicated and the prey strain is the *E. coli* MG1655 carrying pPSV37-sfGFP plasmid. Cells of killer and prey were mixed at a ratio of 10:1 (killer: prey), and co-incubated for 3 h at 37 °C. Survival of killer strains during competition assays is depicted in **F**. For **A**, **B**, **E**, and **F**, error bars indicate the mean +/- standard deviation of at least three biological replicates and statistical significance was calculated using One-way ANOVA test for each group, \*\**P* < 0.01, \*\*\*\**P* < 0.0001. DL, detection limit. **G**, Interaction of Fha^AC^ and Fha^VC^ with TssL and TssE. Pull-down analysis was performed using His-tagged SUMO (control), TssL, or SUMO-TssE, and FLAG-tagged MBP (control), Fha^AC^, or Fha^VC^. **H**, Bacterial two-hybrid analysis evaluating the interaction of Fha with TssJ^ΔSP^, TssM_cyto_, TssM_peri_, TssL_cyto_, and TssM_peri_. TssJ^ΔSP^ deleted its natural Sec signal. Co-expression of T25 or T18 adenylate cyclase subunit-fused proteins was performed in BTH101 reporter strain as indicated. The control proteins Pal and TolB showed positive interaction (49).

**Supplementary Figure 2. Fha condensates selectively recruit T6SS-associated components. A**, SDS-PAGE analysis of purified Fha proteins. **B**, Representative fluorescence recovery of a photobleached Fha_sfGFP focus in *E. coli* BL21(DE3). A representative 8- ×8-μm field is shown. Scale bar: 1 μm. **C**, Quantification of the FRAP analyses for Fha_sfGFP in *E. coli* BL21(DE3). **D**, Fluorescence microscopy images showing Fha_mCherry2 condensates recruited a number of T6SS-associated client proteins in *E. coli* BL21(DE3). The sfGFP and sfGFP-labeled TssJ, RimB1, TssM, TssL, TssE, TssA, Hcp, ClpV, VgrG3, EagT2, and RhsB were cloned on a constitutive-expression vector pBBR1MCS2, respectively. To mitigate the impact of the C-terminal toxin domain and self-cleavage of RhsB, RhsB^NM^ ^D280A^ mutant were used here. The sfGFP and sfGFP-labeled T6SS-associated proteins were expressed alone or co-expressed with Fha_mCherry2 in *E. coli* BL21(DE3). A representative 30- × 30-μm field of cells is shown. Scale bar: 5 μm. White boxes indicate the selected areas (5 ×5 µm) shown in Figure 2E.

**Supplementary Figure 3. Characterization of the FHA domain, IDR, and Fha C-terminal domain in Fha. A**, SDS-PAGE analysis of purified sfGFP, sfGFP_TssL_cyto_, and sfGFP_TssE proteins. **B**, The predicted structure of Fha using AlphaFold2. The N-terminal region of Fha (1- 101 aa) is depicted in green, while the C-terminal region (562 to 777 aa) is highlighted in blue. The predicted amphipathic helix in the C-terminus of Fha is illustrated on the right. Yellow, hydrophobic residues; blue, basic; red, acidic; gray, other residues. **C**, Fluorescence microscopy images (30 ×30 µm) showing the subcellular localization of Fha_sfGFP and its truncated mutants in *E. coli* BL21(DE3). Cells were stained with FM 4-64 before imaging. Scale bar, 5 μm. White boxes indicate the selected areas (5 ×5 µm) shown in Figure 3A. D, Representative fluorescence recovery of a photobleached Fha^N^_sfGFP focus in *E. coli* BL21(DE3). **E**, Representative fluorescence recovery of a photobleached Fha^IDR^_sfGFP focus in *E. coli* BL21(DE3). For **D** and **E**, a representative 8- ×8-μm field is shown. Scale bar: 1 μm. **F**, Fluorescence microscopy images showing the subcellular localization of sfGFP-labeled Fha and its mutants in *E. coli* BL21(DE3). A representative 30- ×30-μm field of cells with a 3× magnified 5- ×5-μm inset (marked by box) is shown. Scale bar: 5 μm.

**Supplementary Figure 4. The Fha IDR is indispensable for T6SS activities in AAC00-1. A**, Fluorescence microscopy images (30 ×30 µm) showing the co-localization of Fha^N^_mCherry2 or Fha^IDR^_mCherry2 with sfGFP, sfGFP_TssL, or TssE_sfGFP in *E. coli* BL21(DE3). The sfGFP and sfGFP-labeled TssL and TssE were expressed alone or co-expressed with Fha^N^_mCherry2 or Fha^IDR^_mCherry2 in *E. coli* BL21(DE3), respectively. White boxes indicate the selected areas (5 ×5 µm) shown in Figure 3C. B, Competition analysis of the AAC00-1 wild type (WT), T6SS-null mutant Δ*tssM*, and the *sfGFP_tssL tssB_mCherry2* mutant. Killer strains are indicated and the prey strain is the *E. coli* MG1655 carrying pPSV37-sfGFP plasmid. Survival of killer strains during competition assays is depicted in **C**. **D**, Fluorescence microscopy images showing the co- localization between sfGFP_TssL and TssB_mCherry2 in AAC00-1 Parental, *fha*^Δ*N*^, and *fha*^Δ*IDR*^. A representative 30- ×30-μm field of cells with a 3× magnified 5- ×5-μm inset (marked by box) is shown. Genotypes are indicated at the top. Scale bar: 5 μm. The corresponding signals of sfGFP_TssL were shown in Figure 4A. E, Survival of killer strains during competition assays for which the survival of the prey is depicted in Figure 4E. F, Competition analysis of the AAC00-1 wild type (WT), T6SS-null mutant Δ*tssM*, *fha*^Δ*N*^, *fha*^Δ*IDR*^, and *fha*^Δ*N*^ or *fha*^Δ*IDR*^ mutant complemented with a plasmid-borne Fha. Killer strains are indicated and the prey strain is the *E. coli* MG1655 carrying pPSV37-sfGFP plasmid. Survival of killer strains during competition assays is depicted in **G**. For **B**, **C**, **E**, **F**, and **G**, error bars indicate the mean +/- standard deviation of at least three biological replicates and statistical significance was calculated using One-way ANOVA test for each group, \*\*\**P* < 0.001, \*\*\*\**P* < 0.0001. DL, detection limit.

**Supplementary Figure 5. The Fha IDR is indispensable for T6SS activities in AAC00-1. A,** Fluorescence microscopy images showing the subcellular localization of Fha^C^ ^APH^ ^mut-4^_sfGFP and Fha^C^ ^APH^ ^mut+4^_sfGFP mutant in *E. coli* BL21(DE3). Cells were stained with FM 4-64 before imaging. **B**, Competition analysis of the AAC00-1 wild type (WT), T6SS-null mutant Δ*tssM*, *fha*^APH^ ^mutN^, *fha*^APH^ ^mut-4^, and *fha*^APH^ ^mut+4^ mutants. Killer strains are indicated and the prey strain is the *E. coli* MG1655 carrying pPSV37-sfGFP plasmid. Cells of killer and prey were mixed at a ratio of 10:1 (killer: prey) and co-incubated for 3 h at 37 °C. Survival of killer strains during competition assays is depicted in **C**. Error bars indicate the mean +/- standard deviation of three biological replicates and statistical significance was calculated using One-way ANOVA test for each group. \*\*\*\**P* < 0.0001. DL, detection limit. **D**, Fluorescence microscopy images showing the co-localization between sfGFP_TssL and TssB_mCherry2 in AAC00-1 Parental and its APH mutants. The corresponding signals of sfGFP_TssL were shown in Figure 5B. For **A** and **D**, a representative 30- ×30-μm field of cells with a 3× magnified 5- ×5-μm inset (marked by box) is shown. Scale bar: 5 μm. **E**, Interaction of Fha, Fha^APH^ ^mutN^, Fha^C^, and Fha^C^ ^APH^ ^mutN^ with TssL and TssE. Pull-down analysis was performed using His-tagged SUMO (control), TssL, or SUMO-TssE, and FLAG-tagged MBP (control), Fha, Fha^APH^ ^mutN^, Fha^C^, or Fha^C^ ^APH^ ^mutN^. **F**, Predicted APH in Fha^PA1^ and Fha^VC^. For **A** and **F**, all the predicted APH were subjected to hydrophobicity, hydrophobic moment, and amino acid composition analysis using HeliQuest. Yellow, hydrophobic residues; blue, basic; red, acidic; purple, serine and threonine; pink, asparagine and glutamine; gray, other residues.

**Supplementary Figure 6. Fha homologs mediate the assembly of membrane complex and baseplate in other species. A**, Representative fluorescence recovery of a photobleached Fha1^PA^_sfGFP focus in *E. coli* BL21(DE3). **B**, Representative fluorescence recovery of a photobleached Fha^VC^_sfGFP focus in *E. coli* BL21(DE3). For **A** and **B**, a representative 8- × 8- μm field is shown. Scale bar: 1 μm. **C**, Fluorescence microscopy images showing the co- localization between sfGFP_TssL1^PA^ and TssB1^PA^_mCherry2 in PAO1 Parental and Δ*fha1*. The corresponding signals of sfGFP_TssL1^PA^ were shown in Figure 6D. **D**, Fluorescence microscopy images showing the signals of sfGFP_TssL^VC^ and TssB^VC^_mCherry2 in V52. **E**, Fluorescence microscopy images showing the co-localization between sfGFP_TssG^VC^ and TssB^VC^_mCherry2 in V52 Parental, Δ*fha*, and Δ*tssL*. The corresponding signals of sfGFP_TssL^VC^ were shown in Figure 6E. For **C** to **E**, a representative 30- ×30-μm field of cells with a 3× magnified 5- ×5-μm inset (marked by box) is shown. Scale bar: 5 μm.

**Supplementary Table 1. Plasmids, strains, and primers used in this study.**

**Supplementary Data 1. Alignment of Fha homologs.**

**Supplementary Data 2. Distribution of Fha homologs in 167 T6SS clusters.**

**Supplementary Data 3. Sequence file of TssC homologs for Figure 6A**.

