## Supplementary Information for "LLPS condensates of Fha initiate the inside-out assembly of the type VI secretion system"

**Running title: Fha primes T6SS assembly**

Tong-Tong Pei[1, 2], Ying An[2], Xing-Yu Wang[1, 2], Han Luo[1], Yumin Kan[1], Hao Li[1, 2], Ming-Xuan Tang[2], Zi-Yan Ye[2], Jia-Xin Liang[2], Tao Jian[2], Hao-Yu Zheng[1, 2], Zeng-Hang Wang[1, 2], Xiaoye Liang[1], Mingjie Zhang[2], Xiaotian Liu[2], Tao Dong[2] *

**Affiliations:**

[1] State Key Laboratory of Microbial Metabolism, Joint International Research Laboratory of Metabolic & Developmental Sciences, School of Life Sciences and Biotechnology, Shanghai Jiao Tong University, Shanghai, 200240, China

[2] School of Life Sciences, Southern University of Science and Technology, Shenzhen, Guangdong, 518055, China

* Corresponding

**Supplementary Materials and Methods**

**Bacterial competition assay**

AAC00-1 killer strains were grown in liquid LB media to $OD_{600}$ ~ 2 at 28°C, while prey cells were collected from overnight LB-agar plates. Cells of killer and prey were mixed at a ratio of 10:1 (killer: prey), spotted on LB-agar plates, and co-incubated for 1 h at 37 °C unless stated otherwise. Survival of killer and prey cells was quantified by 10-fold serial dilution on selective media, respectively. Error bars indicate the mean +/- standard deviation of at least three biological replicates. One-way ANOVA with Dunnett's multiple comparisons test was performed to determine *P*-values. Statistical analysis was performed using the GraphPad Prism software (9.3.0).

**Western blotting analysis**

The Western blotting analyses were performed as described previously (1). Briefly, proteins of interest were run on an SDS-PAGE gel (Yeasen Biotechnology) and transferred to a PVDF membrane (Bio-Rad). The membrane was treated with blocking buffer (5% [w/v] non-fat milk, 50 mM Tris, 150 mM NaCl, 0.1% [v/v] Tween-20, pH 7.6) for 1 h at room temperature, followed by sequential incubation with primary antibodies and secondary HRP-conjugated antibodies in antibody diluent buffer (1% [w/v] non-fat milk, 50 mM Tris, 150 mM NaCl, 0.1% [v/v] Tween-20, pH 7.6). The Clarity ECL solution (Bio-Rad) was used for detecting signals. Monoclonal antibodies to epitope tags were ordered from Smart-lifesciences (FLAG, Product # SLAB0102 and 6His, Product # SLAB2803) and Biolegend (RpoB, Product # 663905). The polyclonal antibody to Aave_1465 (Hcp) was custom-made by Shanghai Youlong Biotech. The HRP-linked secondary antibodies were purchased from ZSGB-Bio (Product # ZB-2305 [mouse] and # ZB-2301 [rabbit]).

**Protein pull-down assay**

Genes of interest were cloned with epitope tags into pET, pBAD and pBBR1MCS2 vectors as indicated. For pET vectors, *E. coli* BL21(DE3) cells carrying corresponding plasmids were grown in LB with appropriate antibiotics to $OD_{600}$ ~ 0.6 and induced with 1 mM IPTG overnight at 20 °C. For pBAD vectors, *E. coli* T-Fast cells carrying corresponding plasmids were grown in LB with appropriate antibiotics to $OD_{600}$ ~ 0.6, and induced with 0.1% arabinose for 3 h at 30 °C. For pBBR1MCS2 vectors, *E. coli* T-Fast cells carrying corresponding plasmids were grown in LB with appropriate antibiotics for about 6 h at 37 °C. Cells were then collected by centrifugation, resuspended in lysis buffer (20 mM Tris, 500 mM NaCl, 50 mM imidazole with 1 × protease inhibitor [Thermo Scientific], pH 8.0) as indicated, and lysed by sonication on ice. After centrifugation, the acquired supernatants were mixed as indicated and loaded to Ni-NTA beads (Smart-lifesciences). The mixtures were incubated for 1 h with rotation at 4 °C, before washing by wash buffer (20 mM Tris, 500 mM NaCl, 50 mM imidazole, pH 8.0) for four times, and eluted in elution buffer (20 mM Tris, 500 mM NaCl, 400 mM imidazole, pH 8.0). Input and eluted samples were analyzed by Western blotting analyses.

**Protein purification**

Strep-tagged or His-tagged proteins were expressed using the pET vectors in *E. coli* BL21(DE3), respectively. The cells were cultured in liquid LB media to exponential phase ($OD_{600}$ ~ 0.6) at 37 °C and induced with 1 mM IPTG at 16 °C for 16 h. The cells were subsequently collected by centrifugation at 4,500 × *g* for 15 min. For purifying strep-tagged proteins, the resulting pellets were resuspended in PBS buffer (10 mM $Na_2HPO_4$, 140 mM NaCl, 3 mM KCl, pH 7.4) and lysed using sonication. Lysates were centrifuged at 15,000 × *g* for 30 min and the supernatants were incubated with Streptactin beads (Smart-lifesciences). Proteins were eluted in PBS buffer supplemented with 2.5 mM biotin. For purifying His-tagged proteins, the resulting pellets were resuspended in Tris buffer (20 mM Tris-HCl, 150 mM NaCl, pH 7.4) and lysed using sonication. Lysates were centrifuged at 15,000 × *g* for 30 min and the supernatants were incubated with Ni-NTA beads (Smart-lifesciences). Proteins were eluted in Tris buffer supplemented with variable concentrations of imidazole.

**Protein secretion assay**

Cultures were grown aerobically in liquid LB media at 28 °C overnight to $OD_{600}$ ~ 2 and then normalized to $OD_{600}$ = 2. To make the “Culture” samples, 40 µL of the normalized cultures were taken and mixed with 10 µL 5 × SDS loading buffer (Epizyme Biotech). Next, the remaining cultures were centrifuged at 10,000 × *g* for 2 min to sperate the supernatants and pellets. Supernatants were used as the “Sec” samples, while pelleted cells were resuspended in fresh liquid LB media and used as the “Cell” samples. All of the “Culture”, “Cell” and “Sec” samples were analyzed by Western blotting analyses.

**Bioinformatic analysis**

All gene sequences of *A. citrulli* AAC00-1 were retrieved from the draft genome assembly (GenBank NC_008752.1), managed and analyzed by Benchling. The Fha sequence was analyzed with the CD-search tool of NCBI webserver. The sequences of 45 Fha homologs were retrieved from NCBI and aligned using Clustal Omega (2). Alignment view was generated using ESPript 3 with default settings (3). A total of 167 T6SS clusters were retrieved from the SecReT6 database (4). The classification of T6SS was based on the maximum likelihood phylogenetic tree of TssC homologs. The TssC protein sequences from 154 T6SSi clusters were aligned by ClustalΩ and phylogeny (bootstrap = 1000) was generated by the Interactive Tree Of Life (iTOL) server with default settings (5). The predicated Fha structure was generated with AlphaFold2 (6). The helical wheel plot was generated with Heliquest (7).

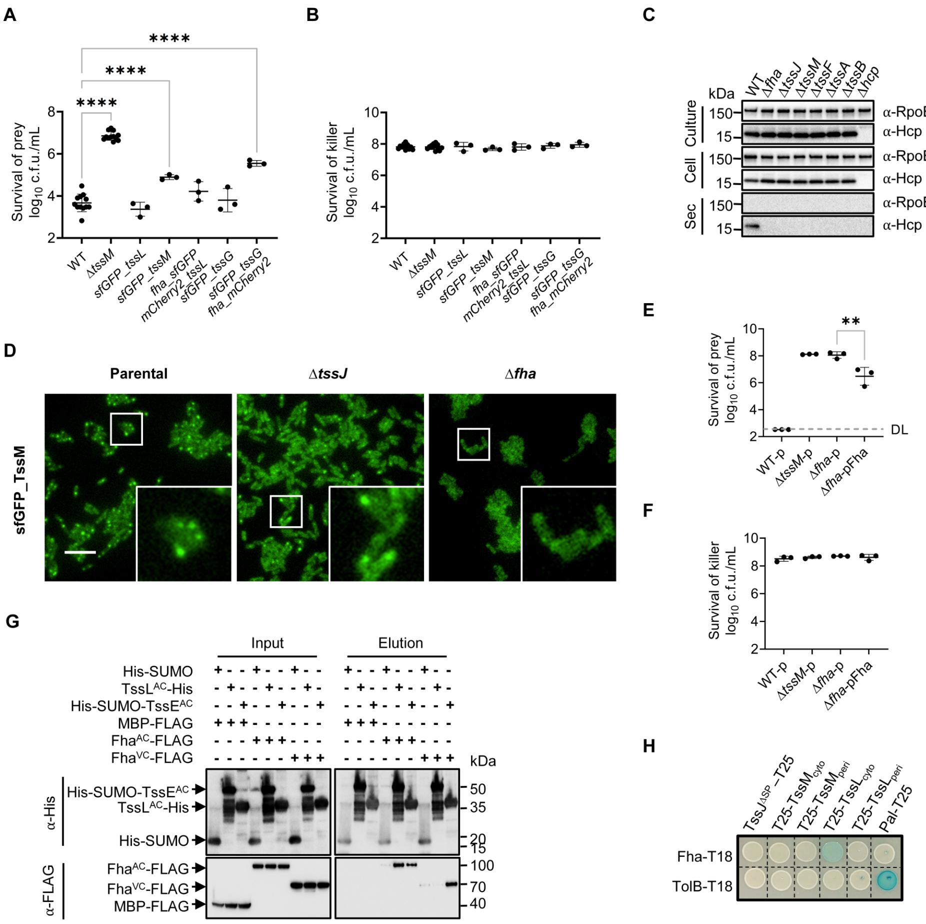


**Supplementary Figure 1. Fha promotes the T6SS biogenesis through direct interactions. A**, Competition analysis of the fluorescence-labeled AAC00-1 strains. Killer strains are indicated and the prey strain is the *E. coli* MG1655 carrying pPSV37-sfGFP plasmid. The wild type (WT) and T6SS-null mutant Δ*tssM* serve as positive and negative control, respectively. Survival of killer strains during competition assays is depicted in **B**. **C**, Secretion analysis of Hcp in AAC00-1 wild type (WT), Δ*fha*, Δ*tssJ*, Δ*tssM*, Δ*tssF*, Δ*tssA*, Δ*tssB*, and Δ*hcp*. RpoB serves as a control for equal loading and autolysis. **D**, Fluorescence microscopy images showing sfGFP_TssM localization in AAC00-1 Parental, Δ*tssJ*, and Δ*fha*. A representative 30- × 30-μm field of cells with a 3× magnified 5- × 5-μm inset (marked by box) is shown. Genotypes are indicated at the top. Scale bar: 5 μm. **E**, Competition analysis of the AAC00-1 WT, Δ*tssM*, Δ*fha*, and Δ*fha* mutant complemented with a plasmid-borne Fha. Killer strains are indicated and the prey strain is the *E. coli* MG1655 carrying pPSV37-sfGFP plasmid. Cells of killer and prey were mixed at a ratio of 10:1 (killer: prey), and co-incubated for 3 h at 37 °C. Survival of killer strains during competition assays is depicted in **F**. For **A**, **B**, **E**, and **F**, error bars indicate the mean +/- standard deviation of at least three biological replicates and statistical significance was calculated using One-way ANOVA test for each group, ***P* < 0.01, *****P* < 0.0001. DL, detection limit. **G**, Interaction of Fha$^{AC}$ and Fha$^{VC}$ with TssL and TssE. Pull-down analysis was performed using His-tagged SUMO (control), TssL, or SUMO-TssE, and FLAG-tagged MBP (control), Fha$^{AC}$, or Fha$^{VC}$. **H**, Bacterial two-hybrid analysis evaluating the interaction of Fha with TssJ$^{\Delta SP}$, TssM$_{cyto}$, TssM$_{peri}$, TssL$_{cyto}$, and TssM$_{peri}$. TssJ$^{\Delta SP}$ deleted its natural Sec signal. Co-expression of T25 or T18 adenylate cyclase subunit-fused proteins was performed in BTH101 reporter strain as indicated. The control proteins Pal and TolB showed positive interaction (8).

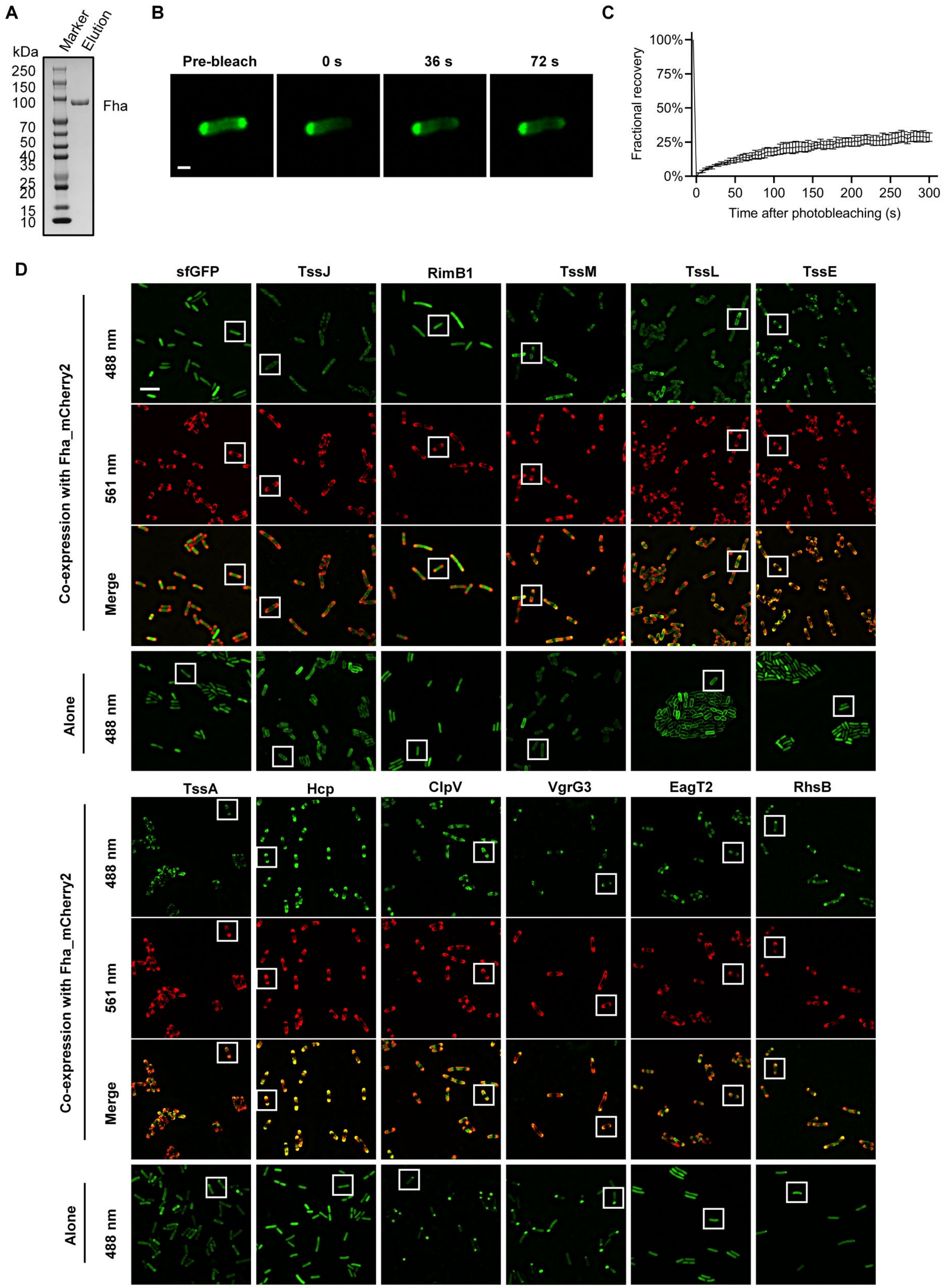
A
B
C
D
Marker
Elution
kDa
250
150
100
70
50
40
35
25
20
15
10
Fha
Pre-bleach
0 s
36 s
72 s
100%
75%
50%
25%
0%
Fractional recovery
0
50
100
150
200
250
300
Time after photobleaching (s)
sfGFP
TssJ
RimB1
TssM
TssL
TssE
Co-expression with Fha_mCherry2
488 nm
561 nm
Merge
Alone
488 nm
TssA
Hcp
ClpV
VgrG3
EagT2
RhsB
Co-expression with Fha_mCherry2
488 nm
561 nm
Merge
Alone
488 nm

**Supplementary Figure 2. Fha condensates selectively recruit T6SS-associated components. A**, SDS-PAGE analysis of purified Fha proteins. **B**, Representative fluorescence recovery of a photobleached Fha_sfGFP focus in *E. coli* BL21(DE3). A representative 8- × 8-μm field is shown. Scale bar: 1 μm. **C**, Quantification of the FRAP analyses for Fha_sfGFP in *E. coli* BL21(DE3). **D**, Fluorescence microscopy images showing Fha_mCherry2 condensates recruited a number of T6SS-associated client proteins in *E. coli* BL21(DE3). The sfGFP and sfGFP-labeled TssJ, RimB1, TssM, TssL, TssE, TssA, Hcp, ClpV, VgrG3, EagT2, and RhsB were cloned on a constitutive-expression vector pBBR1MCS2, respectively. To mitigate the impact of the C-terminal toxin domain and self-cleavage of RhsB, RhsB$^{NM\ D280A}$ mutant were used here. The sfGFP and sfGFP-labeled T6SS-associated proteins were expressed alone or co-expressed with Fha_mCherry2 in *E. coli* BL21(DE3). A representative 30- × 30-μm field of cells is shown. Scale bar: 5 μm. White boxes indicate the selected areas (5 × 5 μm) shown in Figure 2E.

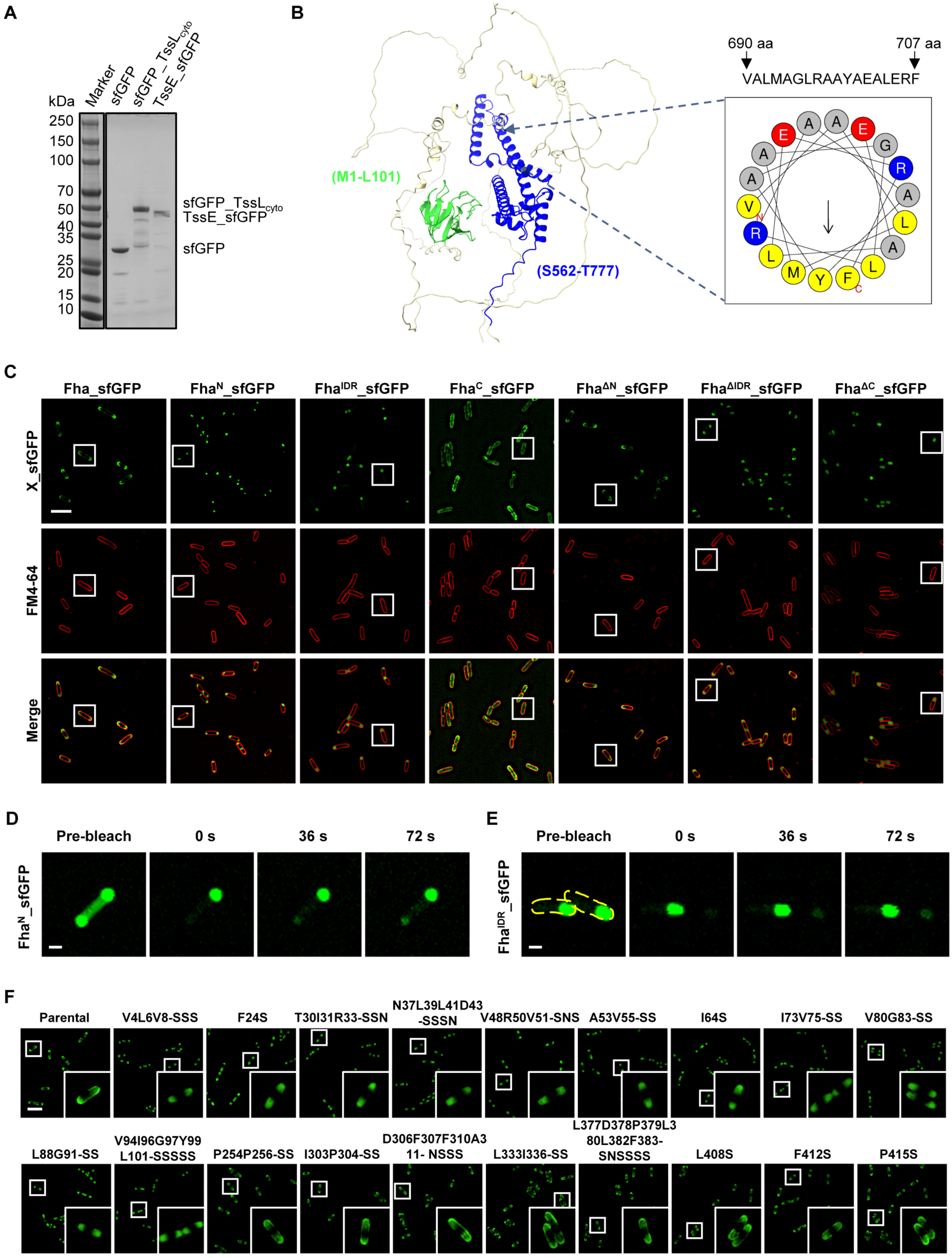
A
kDa
Marker
sfGFP
sfGFP_TssLcyto
TssE_sfGFP
250
150
100
70
50
40
35
25
20
15
10
sfGFP_TssLcyto
TssE_sfGFP
sfGFP
B
690 aa
707 aa
VALMAGLRAAYAEALERF
(M1-L101)
(S562-T777)
C
Fha_sfGFP
FhaN_sfGFP
FhaIDR_sfGFP
FhaC_sfGFP
FhaΔN_sfGFP
FhaΔIDR_sfGFP
FhaΔC_sfGFP
X_sfGFP
FM4-64
Merge
D
Pre-bleach
0 s
36 s
72 s
FhaN_sfGFP
E
Pre-bleach
0 s
36 s
72 s
FhaIDR_sfGFP
F
Parental
V4L6V8-SSS
F24S
T30I31R33-SSN
N37L39L41D43
-SSSN
V48R50V51-SNS
A53V55-SS
I64S
I73V75-SS
V80G83-SS
L88G91-SS
V94I96G97Y99
L101-SSSSS
P254P256-SS
I303P304-SS
D306F307F310A3
11- NSSS
L333I336-SS
L377D378P379L3
80L382F383-
SNSSSS
L408S
F412S
P415S

**Supplementary Figure 3. Characterization of the FHA domain, IDR, and Fha C-terminal domain in Fha.** **A**, SDS-PAGE analysis of purified sfGFP, sfGFP_TssL$_{cyto}$, and sfGFP_TssE proteins. **B**, The predicted structure of Fha using AlphaFold2. The N-terminal region of Fha (1-101 aa) is depicted in green, while the C-terminal region (562 to 777 aa) is highlighted in blue. The predicted amphipathic helix in the C-terminus of Fha is illustrated on the right. Yellow, hydrophobic residues; blue, basic; red, acidic; gray, other residues. **C**, Fluorescence microscopy images (30 × 30 μm) showing the subcellular localization of Fha_sfGFP and its truncated mutants in *E. coli* BL21(DE3). Cells were stained with FM 4-64 before imaging. Scale bar, 5 μm. White boxes indicate the selected areas (5 × 5 μm) shown in Figure 3A. **D**, Representative fluorescence recovery of a photobleached Fha$^{N}$_sfGFP focus in *E. coli* BL21(DE3). **E**, Representative fluorescence recovery of a photobleached Fha$^{IDR}$_sfGFP focus in *E. coli* BL21(DE3). For **D** and **E**, a representative 8- × 8-μm field is shown. Scale bar: 1 μm. **F**, Fluorescence microscopy images showing the subcellular localization of sfGFP-labeled Fha and its mutants in *E. coli* BL21(DE3). A representative 30- × 30-μm field of cells with a 3× magnified 5- × 5-μm inset (marked by box) is shown. Scale bar: 5 μm.

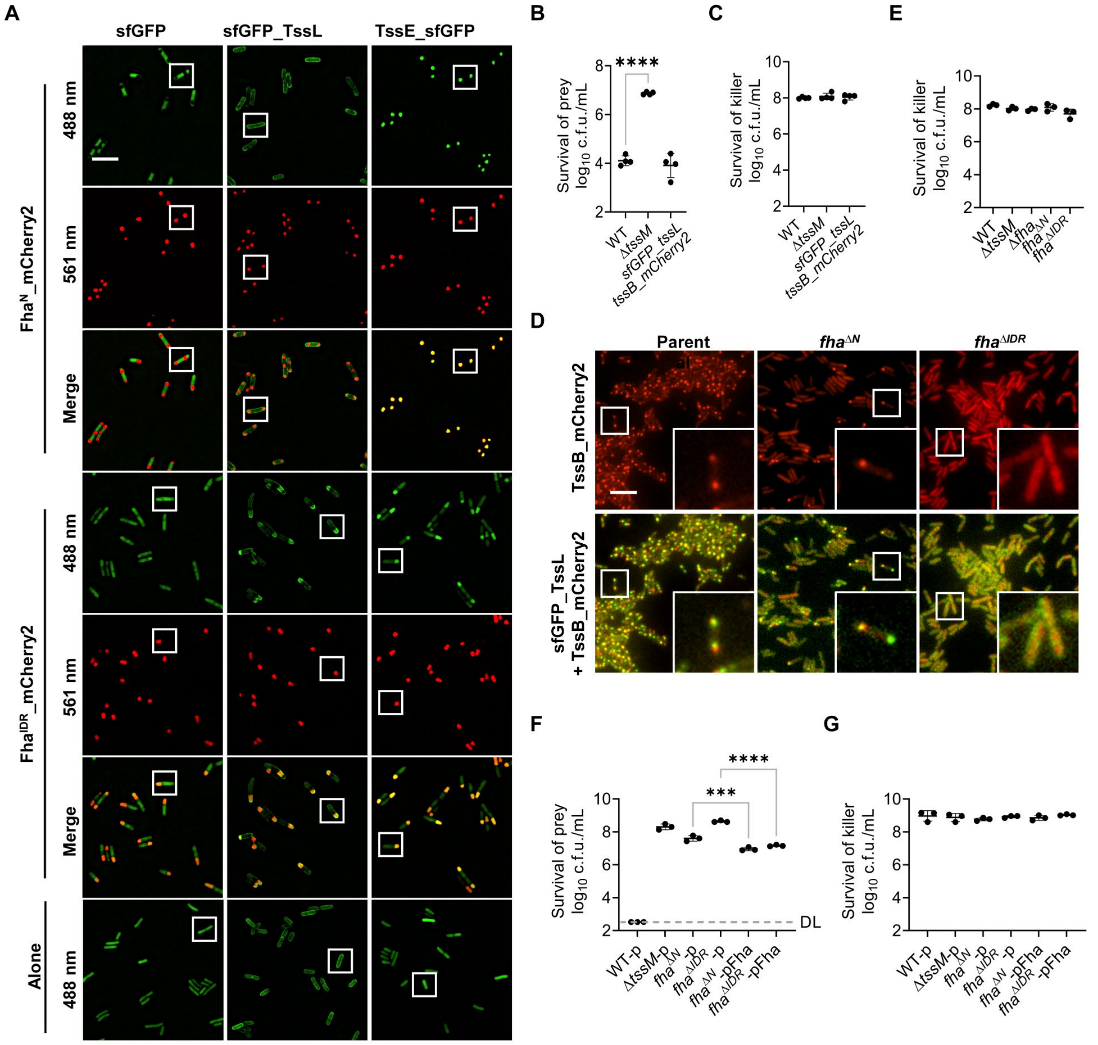


**Supplementary Figure 4. The Fha IDR is indispensable for T6SS activities in AAC00-1. A**, Fluorescence microscopy images (30 × 30 µm) showing the co-localization of $Fha^{N}$_mCherry2 or $Fha^{IDR}$_mCherry2 with sfGFP, sfGFP_TssL, or TssE_sfGFP in *E. coli* BL21(DE3). The sfGFP and sfGFP-labeled TssL and TssE were expressed alone or co-expressed with $Fha^{N}$_mCherry2 or $Fha^{IDR}$_mCherry2 in *E. coli* BL21(DE3), respectively. White boxes indicate the selected areas (5 × 5 µm) shown in Figure 3C. **B**, Competition analysis of the AAC00-1 wild type (WT), T6SS-null mutant Δ*tssM*, and the *sfGFP_tssL tssB_mCherry2* mutant. Killer strains are indicated and the prey strain is the *E. coli* MG1655 carrying pPSV37-sfGFP plasmid. Survival of killer strains during competition assays is depicted in **C**. **D**, Fluorescence microscopy images showing the co-localization between sfGFP_TssL and TssB_mCherry2 in AAC00-1 Parental, $fha^{\Delta N}$, and $fha^{\Delta IDR}$. A representative 30- × 30-µm field of cells with a 3× magnified 5- × 5-µm inset (marked by box) is shown. Genotypes are indicated at the top. Scale bar: 5 µm. The corresponding signals of sfGFP_TssL were shown in Figure 4A. **E**, Survival of killer strains during competition assays for which the survival of the prey is depicted in Figure 4E. **F**, Competition analysis of the AAC00-1 wild type (WT), T6SS-null mutant Δ*tssM*, $fha^{\Delta N}$, $fha^{\Delta IDR}$, and $fha^{\Delta N}$ or $fha^{\Delta IDR}$ mutant complemented with a plasmid-borne Fha. Killer strains are indicated and the prey strain is the *E. coli* MG1655 carrying pPSV37-sfGFP plasmid. Survival of killer strains during competition assays is depicted in **G**. For **B**, **C**, **E**, **F**, and **G**, error bars indicate the mean +/- standard deviation of at least three biological replicates and statistical significance was calculated using One-way ANOVA test for each group, ****P* < 0.001, *****P* < 0.0001. DL, detection limit.

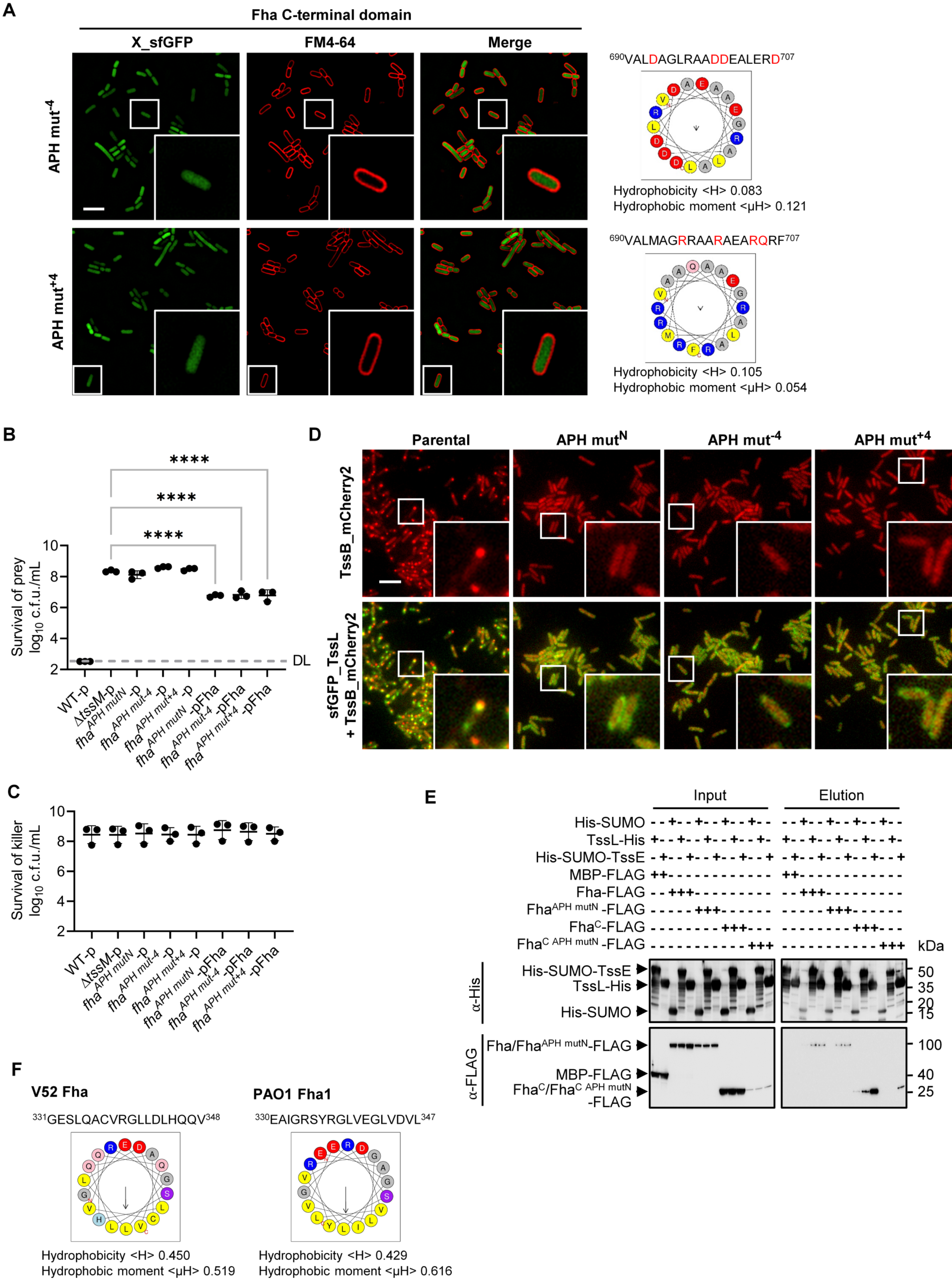
A
Fha C-terminal domain
X_sfGFP
FM4-64
Merge
APH mut-4
APH mut+4
690VALDAGLRAADDEALERD707
Hydrophobicity <H> 0.083
Hydrophobic moment <μH> 0.121
690VALMAGRRAARAEARQRF707
Hydrophobicity <H> 0.105
Hydrophobic moment <μH> 0.054
B
Survival of prey log10 c.f.u./mL
****
****
****
DL
WT-p
ΔtssM-p
fha APH mutN -p
fha APH mut-4 -p
fha APH mut+4 -p
fha APH mutN -pFha
fha APH mut-4 -pFha
fha APH mut+4 -pFha
C
Survival of killer log10 c.f.u./mL
WT-p
ΔtssM-p
fha APH mutN -p
fha APH mut-4 -p
fha APH mut+4 -p
fha APH mutN -pFha
fha APH mut-4 -pFha
fha APH mut+4 -pFha
D
Parental
APH mutN
APH mut-4
APH mut+4
TssB_mCherry2
sfGFP_TssL + TssB_mCherry2
E
Input
Elution
His-SUMO
TssL-His
His-SUMO-TssE
MBP-FLAG
Fha-FLAG
FhaAPH mutN -FLAG
FhaC-FLAG
FhaC APH mutN -FLAG
kDa
α-His
His-SUMO-TssE
TssL-His
His-SUMO
50
35
20
15
α-FLAG
Fha/FhaAPH mutN-FLAG
MBP-FLAG
FhaC/FhaC APH mutN -FLAG
100
40
25
F
V52 Fha
PAO1 Fha1
331GESLQACVRGLLDLHQQV348
330EAIGRSYRGLVEGLVDVL347
Hydrophobicity <H> 0.450
Hydrophobic moment <μH> 0.519
Hydrophobicity <H> 0.429
Hydrophobic moment <μH> 0.616

**Supplementary Figure 5. The Fha C-terminal APH is essential for T6SS activities in AAC00-1. A,** Fluorescence microscopy images showing the subcellular localization of Fha$^{C\ APH\ mut\text{-}4}$_sfGFP and Fha$^{C\ APH\ mut+4}$_sfGFP mutant in *E. coli* BL21(DE3). Cells were stained with FM 4-64 before imaging. **B**, Competition analysis of the AAC00-1 wild type (WT), T6SS-null mutant Δ*tssM*, *fha*$^{APH\ mutN}$, *fha*$^{APH\ mut\text{-}4}$, and *fha*$^{APH\ mut+4}$ mutants. Killer strains are indicated and the prey strain is the *E. coli* MG1655 carrying pPSV37-sfGFP plasmid. Cells of killer and prey were mixed at a ratio of 10:1 (killer: prey) and co-incubated for 3 h at 37 °C. Survival of killer strains during competition assays is depicted in **C**. Error bars indicate the mean +/- standard deviation of three biological replicates and statistical significance was calculated using One-way ANOVA test for each group. ****$P$ < 0.0001. DL, detection limit. **D**, Fluorescence microscopy images showing the co-localization between sfGFP_TssL and TssB_mCherry2 in AAC00-1 Parental and its APH mutants. The corresponding signals of sfGFP_TssL were shown in Figure 5B. For **A** and **D**, a representative 30- × 30-μm field of cells with a 3× magnified 5- × 5-μm inset (marked by box) is shown. Scale bar: 5 μm. **E**, Interaction of Fha, Fha$^{APH\ mutN}$, Fha$^{C}$, and Fha$^{C\ APH\ mutN}$ with TssL and TssE. Pull-down analysis was performed using His-tagged SUMO (control), TssL, or SUMO-TssE, and FLAG-tagged MBP (control), Fha, Fha$^{APH\ mutN}$, Fha$^{C}$, or Fha$^{C\ APH\ mutN}$. **F**, Predicted APH in Fha$^{PA1}$ and Fha$^{VC}$. For **A** and **F**, all the predicted APH were subjected to hydrophobicity, hydrophobic moment, and amino acid composition analysis using HeliQuest. Yellow, hydrophobic residues; blue, basic; red, acidic; purple, serine and threonine; pink, asparagine and glutamine; gray, other residues.

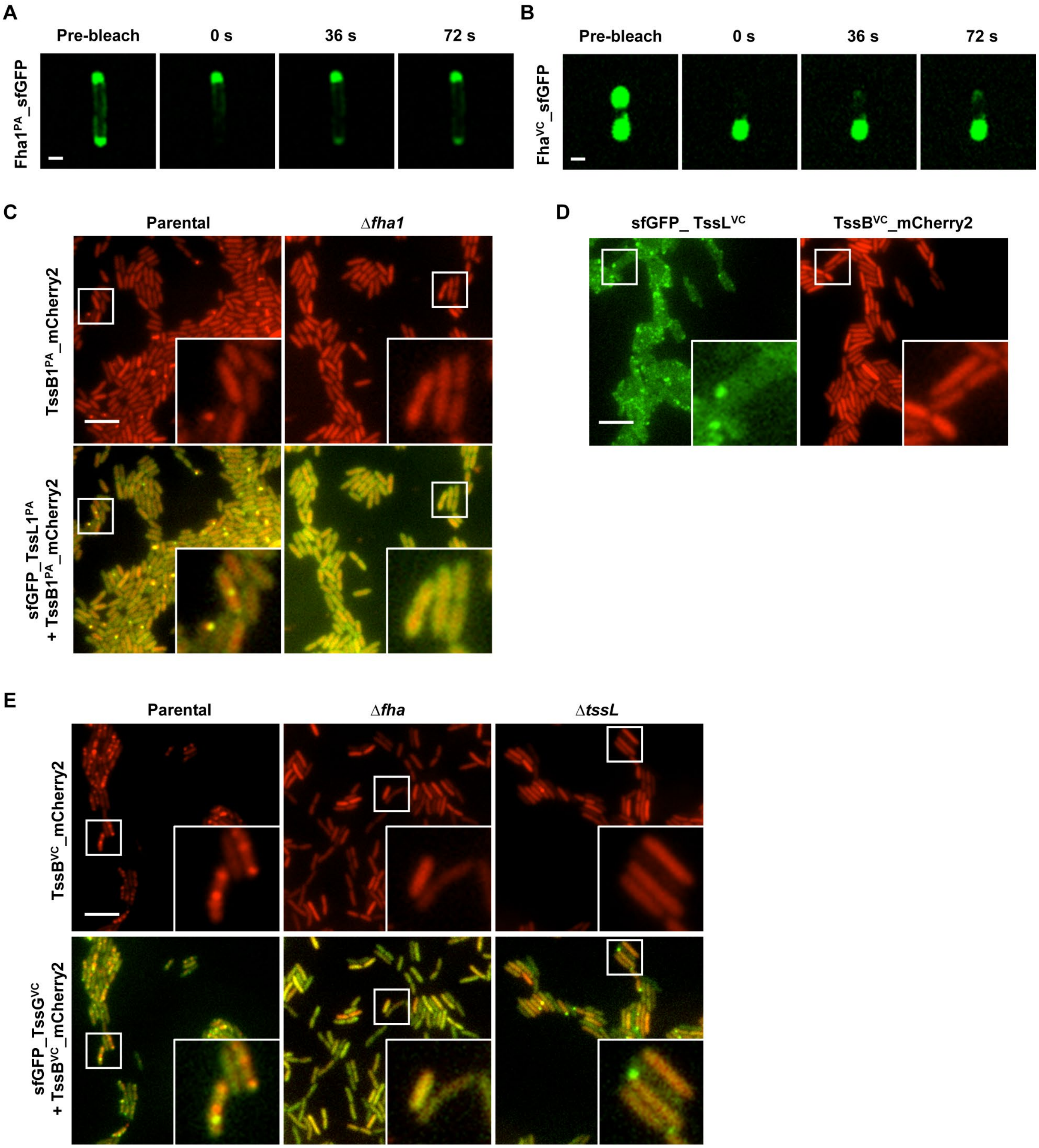


**Supplementary Figure 6. Fha homologs mediate the assembly of membrane complex and baseplate in other species.** **A**, Representative fluorescence recovery of a photobleached $Fha1^{PA}$_sfGFP focus in *E. coli* BL21(DE3). **B**, Representative fluorescence recovery of a photobleached $Fha^{VC}$_sfGFP focus in *E. coli* BL21(DE3). For **A** and **B**, a representative 8- × 8-μm field is shown. Scale bar: 1 μm. **C**, Fluorescence microscopy images showing the co-localization between sfGFP_$TssL1^{PA}$ and $TssB1^{PA}$_mCherry2 in PAO1 Parental and Δ*fha1*. The corresponding signals of sfGFP_$TssL1^{PA}$ were shown in Figure 6D. **D**, Fluorescence microscopy images showing the signals of sfGFP_$TssL^{VC}$ and $TssB^{VC}$_mCherry2 in V52. **E**, Fluorescence microscopy images showing the co-localization between sfGFP_$TssG^{VC}$ and $TssB^{VC}$_mCherry2 in V52 Parental, Δ*fha*, and Δ*tssL*. The corresponding signals of sfGFP_$TssL^{VC}$ were shown in Figure 6E. For **C** to **E**, a representative 30- × 30-μm field of cells with a 3× magnified 5- × 5-μm inset (marked by box) is shown. Scale bar: 5 μm.

### Supplementary Table 1. Plasmids, strains, and primers

| Plasmid | Description | Reference |
|---|---|---|
| pEXG2.0 | Suicidal conjugation vector for chromosomal allelic changes in AAC00-1 and PAO1 | (9) |
| pEXG2.0-tssJ | Suicidal vector to construct the AAC00-1 in-frame deletion mutant of *tssJ* | This study |
| pEXG2.0-tssL | Suicidal vector to construct the AAC00-1 in-frame deletion mutant of *tssL* | This study |
| pEXG2.0-tssM | Suicidal vector to construct the AAC00-1 in-frame deletion mutant of *tssM* | (1) |
| pEXG2.0-tssF | Suicidal vector to construct the AAC00-1 in-frame deletion mutant of *tssF* | This study |
| pEXG2.0-tssA | Suicidal vector to construct the AAC00-1 in-frame deletion mutant of *tssA* | This study |
| pEXG2.0-tssB | Suicidal vector to construct the AAC00-1 in-frame deletion mutant of *tssB* | This study |
| pEXG2.0-hcp | Suicidal vector to construct the AAC00-1 in-frame deletion mutant of *hcp* | This study |
| pEXG2.0-fha | Suicidal vector to construct the AAC00-1 in-frame deletion mutant of *fha* | This study |
| pEXG2.0-fha$^{\Delta N}$ | Suicidal vector to construct the AAC00-1 in-frame deletion of N-terminal FHA domain in Fha | This study |
| pEXG2.0-fha$^{\Delta IDR}$ | Suicidal vector to construct the AAC00-1 in-frame deletion of IDR in Fha | This study |
| pEXG2.0-fha$^{APH\ mut1}$ | Suicidal vector to construct the AAC00-1 mutant with chromosomal mutation of Fha C-terminal APH | This study |
| pEXG2.0-fha$^{APH\ mut2}$ | Suicidal vector to construct the AAC00-1 mutant with chromosomal mutation of Fha C-terminal APH | This study |
| pEXG2.0-fha$^{APH\ mut3}$ | Suicidal vector to construct the AAC00-1 mutant with chromosomal mutation of Fha C-terminal APH | This study |
| pEXG2.0-sfGFP-TssL | Suicidal vector to construct the AAC00-1 mutant with chromosomal insertion sfGFP_TssL | This study |
| pEXG2.0-mCherry2-TssL | Suicidal vector to construct the AAC00-1 mutant with chromosomal insertion mCherry2_TssL | This study |
| pEXG2.0-TssB-mCherry2 | Suicidal vector to construct the AAC00-1 mutant with chromosomal insertion TssB_mCherry2 | This study |
| pEXG2.0-Fha-sfGFP | Suicidal vector to construct the AAC00-1 mutant with chromosomal insertion Fha_sfGFP | This study |
| pEXG2.0-Fha-mCherry2 | Suicidal vector to construct the AAC00-1 mutant with chromosomal insertion Fha_mCherry2 | This study |
| pEXG2.0-sfGFP-TssG | Suicidal vector to construct the AAC00-1 mutant with chromosomal insertion sfGFP_TssG | This study |
| pEXG2.0-sfGFP-TssL1 | Suicidal vector to construct the PAO1 mutant with chromosomal insertion sfGFP_TssL1 | This study |
| pEXG2.0-fha1 | Suicidal vector to construct the PAO1 in-frame deletion mutant of *fha1* | This study |
| pEXG2.0-tssB1 | Suicidal vector to construct the PAO1 in-frame deletion mutant of *tssB1* | This study |
| pDS132 | Suicidal conjugation vector for chromosomal allelic changes in V52 | (10) |
| pDS132-sfGFP-TssG | Suicidal vector to construct the V52 mutant with chromosomal insertion sfGFP_TssG | This study |
| pDS132-sfGFP-TssL | Suicidal vector to construct the V52 mutant with chromosomal insertion sfGFP_TssL | This study |
| pDS132-fha | Suicidal vector to construct the V52 in-frame deletion mutant of *fha* | This study |
| pDS132-tssL | Suicidal vector to construct the V52 in-frame deletion mutant of *tssL* | This study |
| pPSV37-sfGFP | IPTG inducible expression of sfGFP | Lab stock |

| | | |
|---|---|---|
| pBBR1MCS2 | A broad-host-range cloning vector, kanamycin resistance | Lab stock |
| pBBR1MCS2-Fha-3V5 | Constitutive expression of Fha with a C-terminal 3V5 tag | This study |
| pBBR1MCS2-Fha-FLAG | Constitutive expression of Fha with a C-terminal FLAG tag | This study |
| pBAD24Kan-MBP-FLAG | Arabinose inducible expression of MBP with a C-terminal FLAG tag | This study |
| pETSUMO | IPTG inducible expression of His-SUMO | Lab stock |
| pETSUMO-TssM | IPTG inducible expression of TssM with an N-terminal His-SUMO tag | This study |
| pETSUMO-TssE | IPTG inducible expression of TssE with an N-terminal His-SUMO tag | This study |
| pETSUMO-TssF | IPTG inducible expression of TssF with an N-terminal His-SUMO tag | This study |
| pETSUMO-TssG | IPTG inducible expression of TssG with an N-terminal His-SUMO tag | This study |
| pETSUMO-TssK | IPTG inducible expression of TssK with an N-terminal His-SUMO tag | This study |
| pET22b-Fha-sfGFP-2strep | IPTG inducible expression of Fha_sfGFP with a C-terminal 2Strep tag | This study |
| pET22b-Fha-sfGFP-His | IPTG inducible expression of Fha_sfGFP with a C-terminal His tag | This study |
| pET22b-TssJ-His | IPTG inducible expression of TssJ with a C-terminal His tag | This study |
| pET22b-TssL-His | IPTG inducible expression of TssL with a C-terminal His tag | This study |
| pET22b-TssA-His | IPTG inducible expression of TssA with a C-terminal His tag | This study |
| pET21a-Fha-2strep | IPTG inducible expression of Fha with a C-terminal 2Strep tag | This study |
| pKNT25-Pal | IPTG inducible expression of Pal for bacterial two-hybrid analysis | Lab stock |
| pKNT25-TssJ$^{\Delta SP}$ | IPTG inducible expression of TssJ$^{\Delta SP}$ for bacterial two-hybrid analysis | This study |
| pKNT25N-TssM$_{cyto}$ | IPTG inducible expression of TssM$_{cyto}$ for bacterial two-hybrid analysis | This study |
| pKNT25-TssM$_{cyto}$ | IPTG inducible expression of TssM$_{cyto}$ for bacterial two-hybrid analysis | This study |
| pKNT25N-TssM$_{peri}$ | IPTG inducible expression of TssM$_{peri}$ for bacterial two-hybrid analysis | This study |
| pKNT25N-TssL$_{cyto}$ | IPTG inducible expression of TssL$_{cyto}$ for bacterial two-hybrid analysis | This study |
| pKNT25N- TssL$_{peri}$ | IPTG inducible expression of TssL$_{peri}$ for bacterial two-hybrid analysis | This study |
| pCH363-TolB | IPTG inducible expression of TolB for bacterial two-hybrid analysis | Lab stock |
| pCH363-Fha | IPTG inducible expression of Fha for bacterial two-hybrid analysis | This study |
| pET22b-Fha$^{N}$-sfGFP-His | IPTG inducible expression of Fha$^{N}$_sfGFP with a C-terminal His tag | This study |
| pET22b-Fha$^{IDR}$-sfGFP-His | IPTG inducible expression of Fha$^{IDR}$_sfGFP with a C-terminal His tag | This study |
| pET22b-Fha$^{C}$-sfGFP-His | IPTG inducible expression of Fha$^{C}$_sfGFP with a C-terminal His tag | This study |
| pET22b-Fha$^{\Delta N}$-sfGFP-His | IPTG inducible expression of Fha$^{\Delta N}$_sfGFP with a C-terminal His tag | This study |
| pET22b-Fha$^{\Delta IDR}$-sfGFP-His | IPTG inducible expression of Fha$^{\Delta IDR}$_sfGFP with a C-terminal His tag | This study |
| pET22b-Fha$^{\Delta C}$-sfGFP-His | IPTG inducible expression of Fha$^{\Delta C}$_sfGFP with a C-terminal His tag | This study |
| pET22b-Fha$^{N}$-FLAG | IPTG inducible expression of Fha$^{N}$ with a C-terminal FLAG tag | This study |

| | | |
|---|---|---|
| pET22b-Fha$^{IDR}$-FLAG | IPTG inducible expression of Fha$^{IDR}$ with a C-terminal FLAG tag | This study |
| pET22b-Fha$^{C}$-FLAG | IPTG inducible expression of Fha$^{C}$ with a C-terminal FLAG tag | This study |
| pET22b-Fha-mCherry2-His | IPTG inducible expression of Fha_mCherry2 with a C-terminal His tag | This study |
| pET22b-Fha$^{N}$-mCherry2-His | IPTG inducible expression of Fha$^{N}$_mCherry2 with a C-terminal His tag | This study |
| pET22b-Fha$^{IDR}$-mCherry2-His | IPTG inducible expression of Fha$^{IDR}$_mCherry2 with a C-terminal His tag | This study |
| pET22b-Fha$^{C}$-mCherry2-His | IPTG inducible expression of Fha$^{C}$_mCherry2 with a C-terminal His tag | This study |
| pET22b-Fha$^{C\ APH\ mut1}$-sfGFP-2strep | IPTG inducible expression of Fha$^{C\ APH\ mut1}$_sfGFP with a C-terminal 2strep tag | This study |
| pET22b-Fha$^{C\ APH\ mut2}$-sfGFP-2strep | IPTG inducible expression of Fha$^{C\ APH\ mut2}$_sfGFP with a C-terminal 2strep tag | This study |
| pET22b-Fha$^{C\ APH\ mut3}$-sfGFP-2strep | IPTG inducible expression of Fha$^{C\ APH\ mut3}$_sfGFP with a C-terminal 2strep tag | This study |
| pet22b-Fha-sfGFP-V4L6V8-SSS | IPTG inducible expression of Fha-sfGFP-V4L6V8-SSS with a C-terminal 2strep tag | This study |
| pet22b-Fha-sfGFP-F24S | IPTG inducible expression of Fha-sfGFP-V4L6V8-SSS with a C-terminal 2strep tag | This study |
| pet22b-Fha-sfGFP-T30-I31-R33-SSN | IPTG inducible expression of Fha-sfGFP-T30-I31-R33-SSN with a C-terminal 2strep tag | This study |
| pet22b-Fha-sfGFP-N37-L39-L41-D43-SSSN | IPTG inducible expression of Fha-sfGFP-N37-L39-L41-D43-SSSN with a C-terminal 2strep tag | This study |
| pet22b-Fha-sfGFP-V48-R50-V51-SNS | IPTG inducible expression of Fha-sfGFP-V48-R50-V51-SNS with a C-terminal 2strep tag | This study |
| pet22b-Fha-sfGFP-A53-V55-SS | IPTG inducible expression of Fha-sfGFP-A53-V55-SS with a C-terminal 2strep tag | This study |
| pet22b-Fha-sfGFP-I64S | IPTG inducible expression of Fha-sfGFP-I64S with a C-terminal 2strep tag | This study |
| pet22b-Fha-sfGFP-I73-V75-SS | IPTG inducible expression of Fha-sfGFP-I73-V75-SS with a C-terminal 2strep tag | This study |
| pet22b-Fha-sfGFP-L88-G91-SS | IPTG inducible expression of Fha-sfGFP-L88-G91-SS with a C-terminal 2strep tag | This study |
| pet22b-Fha-sfGFP-V94-I96-G97-Y99-L101-SSSSS | IPTG inducible expression of Fha-sfGFP-V94-I96-G97-Y99-L101-SSSSS with a C-terminal 2strep tag | This study |
| pet22b-Fha-sfGFP-P254-P256-SS | IPTG inducible expression of Fha-sfGFP-P254-P256-SS with a C-terminal 2strep tag | This study |
| pet22b-Fha-sfGFP-I303-P304-SS | IPTG inducible expression of Fha-sfGFP-I303-P304-SS with a C-terminal 2strep tag | This study |
| pet22b-Fha-sfGFP-D306-F307-F310-A311- NSSS | IPTG inducible expression of Fha-sfGFP-D306-F307-F310-A311- NSSS with a C-terminal 2strep tag | This study |
| pet22b-Fha-sfGFP-L333-I336-SS | IPTG inducible expression of Fha-sfGFP-L333-I336-SS with a C-terminal 2strep tag | This study |
| pet22b-Fha-sfGFP-L377-D378-P379-L380-L382-F383-SNSSSS | IPTG inducible expression of Fha-sfGFP-L377-D378-P379-L380-L382-F383-SNSSSS with a C-terminal 2strep tag | This study |
| pet22b-Fha-sfGFP-L408S | IPTG inducible expression of Fha-sfGFP-L408S with a C-terminal 2strep tag | This study |
| pet22b-Fha-sfGFP-F412S | IPTG inducible expression of Fha-sfGFP-F412S with a C-terminal 2strep tag | This study |
| pet22b-Fha-sfGFP-P415S | IPTG inducible expression of Fha-sfGFP-P415S with a C-terminal 2strep tag | This study |
| pBBR1MCS2-sfGFP | Constitutive expression of sfGFP | This study |

| | | |
|---|---|---|
| pBBR1MCS2-TssJ-sfGFP | Constitutive expression of TssJ_sfGFP | This study |
| pBBR1MCS2-sfGFP-TssM | Constitutive expression of sfGFP_TssM | This study |
| pBBR1MCS2-sfGFP-TssL | Constitutive expression of sfGFP_TssL | This study |
| pBBR1MCS2-sfGFP-TssA | Constitutive expression of sfGFP_TssA | This study |
| pBBR1MCS2-Hcp-sfGFP | Constitutive expression of Hcp_sfGFP | This study |
| pBBR1MCS2-VgrG3-sfGFP | Constitutive expression of VgrG3_sfGFP | This study |
| pBBR1MCS2-ClpV-sfGFP | Constitutive expression of ClpV_sfGFP | This study |
| pBBR1MCS2-EagT2-sfGFP | Constitutive expression of EagT2_sfGFP | This study |
| pBBR1MCS2-RhsB$^{NM\ D280A}$-sfGFP | Constitutive expression of RhsB$^{NM\ D280A}$_sfGFP | This study |
| pBBR1MCS2-RimB1-sfGFP | Constitutive expression of RimB1_sfGFP | This study |
| pET22b-Fha1$^{PA}$-sfGFP-His | IPTG inducible expression of Fha1$^{PA}$_sfGFP with a C-terminal His tag | This study |
| pET22b-Fha$^{vc}$-sfGFP-His | IPTG inducible expression of Fha1$^{VC}$_sfGFP with a C-terminal His tag | This study |

| **Strain** | **Genotype** | **Description** | **Reference** |
|---|---|---|---|
| *Acidovorax citrulli* AAC00-1 | AAC00-1 | Reference strain of Group II strain | (1) |
| | Δ*tssM* | In-frame deletion of *tssM* | (1) |
| | Δ*fha* | In-frame deletion of *fha* | This study |
| | *sfGFP_tssL* | Chromosomal fusion of the sfGFP_TssL | This study |
| | *sfGFP_tssL tssB_mCherry2* | Chromosomal fusion of the sfGFP_TssL and TssB_mCherry2 | This study |
| | *sfGFP_tssM* | Chromosomal fusion of the sfGFP_TssM | This study |
| | *fha_sfGFP* | Chromosomal fusion of the Fha_sfGFP | This study |
| | *mCherry2_tssL fha_sfGFP* | Chromosomal fusion of the mCherry2_TssL and Fha_sfGFP | This study |
| | *fha_mCherry2* | Chromosomal fusion of the Fha_mCherry2 | This study |
| | *sfGFP_tssG* | Chromosomal fusion of the sfGFP_TssG | This study |
| | *fha_mCherry2 sfGFP_tssG* | Chromosomal fusion of the Fha_mCherry2 and sfGFP_TssG | This study |
| | Δ*fha sfGFP_tssL* | *sfGFP_tssL* with in-frame deletion of *fha* | This study |
| | Δ*tssJ sfGFP_tssL* | *sfGFP_tssL* with in-frame deletion of *tssJ* | This study |
| | Δ*tssM sfGFP_tssL* | *sfGFP_tssL* with in-frame deletion of *tssM* | This study |
| | Δ*tssF sfGFP_tssL* | *sfGFP_tssL* with in-frame deletion of *tssF* | This study |
| | Δ*tssA sfGFP_tssL* | *sfGFP_tssL* with in-frame deletion of *tssA* | This study |
| | Δ*tssB sfGFP_tssL* | *sfGFP_tssL* with in-frame deletion of *tssB* | This study |
| | Δ*hcp sfGFP_tssL* | *sfGFP_tssL* with in-frame deletion of *Hcp* | This study |

| | | | |
|---|---|---|---|
| | *ΔtssJ sfGFP_tssM* | *sfGFP_tssM* with in-frame deletion of *tssJ* | This study |
| | *Δfha sfGFP_tssM* | *sfGFP_tssM* with in-frame deletion of *fha* | This study |
| | *Δfha sfGFP_tssG* | *sfGFP_tssG* with in-frame deletion of *fha* | This study |
| | *ΔtssJ sfGFP_tssG* | *sfGFP_tssG* with in-frame deletion of *tssJ* | This study |
| | *ΔtssM sfGFP_tssG* | *sfGFP_tssG* with in-frame deletion of *tssM* | This study |
| | *ΔtssF sfGFP_tssG* | *sfGFP_tssG* with in-frame deletion of *tssF* | This study |
| | *ΔtssA sfGFP_tssG* | *sfGFP_tssG* with in-frame deletion of *tssA* | This study |
| | *ΔtssB sfGFP_tssG* | *sfGFP_tssG* with in-frame deletion of *tssB* | This study |
| | *Δhcp sfGFP_tssG* | *sfGFP_tssG* with in-frame deletion of *Hcp* | This study |
| | *fha*$^{\Delta N}$ | In-frame deletion of N-terminal FHA domain in Fha | This study |
| | *fha*$^{\Delta IDR}$ | In-frame deletion of IDR in Fha | This study |
| | *fha*$^{\Delta N}$*_sfGFP* | Chromosomal mutation of Fha$^{\Delta N}$_sfGFP | This study |
| | *fha*$^{\Delta IDR}$*_sfGFP* | Chromosomal mutation of Fha$^{\Delta IDR}$_sfGFP | This study |
| | *fha*$^{\Delta N}$ *sfGFP_tssL tssB_mCherry2* | *sfGFP_tssL tssB_mCherry2* with in-frame deletion of N-terminal FHA domain in Fha | This study |
| | *fha*$^{\Delta IDR}$ *sfGFP_tssL tssB_mCherry2* | *sfGFP_tssL tssB_mCherry2* with in-frame deletion of IDR in Fha | This study |
| | *fha*$^{\Delta N}$ *sfGFP_tssG* | *sfGFP_tssG* with in-frame deletion of N-terminal FHA domain in Fha | This study |
| | *fha*$^{\Delta IDR}$ *sfGFP_tssG* | *sfGFP_tssG* with in-frame deletion of IDR in Fha | This study |
| | *fha*$^{APH\ mutN}$ | Chromosomal mutation of Fha$^{APH\ mutN}$ | This study |
| | *fha*$^{APH\ mut-4}$ | Chromosomal mutation of Fha$^{APH\ mut-4}$ | This study |
| | *fha*$^{APH\ mut+4}$ | Chromosomal mutation of Fha$^{APH\ mut+4}$ | This study |
| | *fha*$^{APH\ mutN}$*_sfGFP* | Chromosomal mutation of Fha$^{APH\ mutN}$_sfGFP | This study |
| | *fha*$^{APH\ mut-4}$*_sfGFP* | Chromosomal mutation of Fha$^{APH\ mut-4}$_sfGFP | This study |
| | *fha*$^{APH\ mut+4}$*_sfGFP* | Chromosomal mutation of Fha$^{APH\ mut+4}$_sfGFP | This study |
| | *fha*$^{APH\ mutN}$ *sfGFP_tssG* | *sfGFP_tssG* with chromosomal mutation of Fha$^{APH\ mutN}$ | This study |
| | *fha*$^{APH\ mut-4}$ *sfGFP_tssG* | *sfGFP_tssG* with chromosomal mutation of Fha$^{APH\ mut-4}$ | This study |
| | *fha*$^{APH\ mut+4}$ *sfGFP_tssG* | *sfGFP_tssG* with chromosomal mutation of Fha$^{APH\ mut+4}$ | This study |
| | *fha*$^{APH\ mutN}$ *sfGFP_tssL tssB_mCherry2* | *sfGFP_tssL tssB_mCherry2* with chromosomal mutation of Fha$^{APH\ mutN}$ | This study |
| | *fha*$^{APH\ mut-4}$ *sfGFP_tssL tssB_mCherry2* | *sfGFP_tssL tssB_mCherry2* with chromosomal mutation of Fha$^{APH\ mut-4}$ | This study |
| | *fha*$^{APH\ mut+4}$ *sfGFP_tssL tssB_mCherry2* | *sfGFP_tssL tssB_mCherry2* with chromosomal mutation of Fha$^{APH\ mut+4}$ | This study |
| *Pseudomonas aeruginosa* PAO1 | *ΔretS ΔtssB2 ΔtssB3 tssB1_mCherry2 sfGFP_tssL1* | Parental strain | This study |
| | *ΔretS ΔtssB2 ΔtssB3 tssB1_mCherry2 sfGFP_tssL1 Δfha1* | PAO1 parental strain with in-frame deletion of *fha1* | This study |

| | | | |
|---|---|---|---|
| | *ΔretS ΔtssB2 ΔtssB3 sfGFP_tssL1 ΔtssB1* | PAO1 parental strain with in-frame deletion of *tssB1* | This study |
| *Vibrio cholerae* V52 | *rhh tssB_mCherry2 sfGFP_tssL* | Chromosomal fusion of the sfGFP_TssL and TssB_mCherry2 | This study |
| | *rhh tssB_mCherry2 sfGFP_tssG* | Parental strain | This study |
| | *rhh tssB_mCherry2 sfGFP_tssG Δfha* | V52 parental strain with in-frame deletion of *fha* | This study |
| | *rhh tssB_mCherry2 sfGFP_tssG ΔtssL* | V52 parental strain with in-frame deletion of *tssL* | This study |
| *E. coli* | | | |
| T-Fast | | Strain used for cloning and gene expression | TIANGEN |
| WM6026 | | Strain used for conjugation | Lab stock |
| BTH101 | | Host strain used for two-hybrid analysis | (11) |
| BL21(DE3) | | Strain used for protein expression | Lab stock |
| MG1655 | | Strain used for competition assay | Lab stock |

| **Primer** | **Sequence (5'-3')** | **Description** |
|---|---|---|
| pDS132-hifi-f | cgatcctttttaacccatcac | Forward primer to amplify pDS132 vector |
| pDS132-hifi-r | cttctagaggtaccgcatgc | Reverse primer to amplify pDS132 vector |
| pDS132-f | tgttgcatgggcataaagttgc | Forward confirmation primer of pDS132 vector |
| pDS132-r | acggctgacatgggaattcc | Reverse confirmation primer of pDS132 vector |
| pEXG2.0-hifi-f | agatctcagagtcgacctgcagaa | Forward primer to amplify pEXG2.0 vector |
| pEXG2.0-hifi-r | gctcgagctcgaattcggta | Reverse primer to amplify pEXG2.0 vector |
| pEXG2.0-f | ctgttgcatgggcataaagttgc | Forward confirmation primer of pEXG2.0 vector |
| pEXG2.0-r | cttcacgttcgctcgcgtat | Reverse confirmation primer of pEXG2.0 vector |
| Hcp-KO1 | taccgaattcgagctcgagcccgggaagcaccgtcttgaccgtatg | Forward primer to amplify the upstream of *Aave_1465* for constructing in-frame deletion of *Aave_1465* |
| Hcp-KO2 | ttgaaggtgccagcggcgaaatcaaggaaaacaaagaagcctga | Reverse primer to amplify the upstream of *Aave_1465* for constructing in-frame deletion of *Aave_1465* |
| Hcp-KO3 | gcttctttgttttccttgatttcgccgctggcacctt | Forward primer to amplify the downstream of *Aave_1465* for constructing in-frame deletion of *Aave_1465* |
| Hcp-KO4 | ctgcaggtcgactctgagatctgggaccgcacatgcacct | Reverse primer to amplify the downstream of *Aave_1465* for constructing in-frame deletion of *Aave_1465* |
| Hcp-KO5 | cttgggcatgcgaggct | Forward primer to confirm the in-frame deletion of *Aave_1465* |
| Hcp-KO6 | cggatgggaaaacttggcc | Reverse primer to confirm the in-frame deletion of *Aave_1465* |
| Fha-KO2 | ttcataggcctccaggaagggttcgactttgtccatgtccg | Reverse primer to amplify the upstream of *Aave_1468* for constructing in-frame deletion of *Aave_1468* |

| | | |
|---|---|---|
| Fha-KO3 | ggacatggacaaagtcgaacccttcctggaggcctatgaaaac | Forward primer to amplify the downstream of *Aave_1468* for constructing in-frame deletion of *Aave_1468* |
| Fha-KO4 | ctgcaggtcgactctgagatctcgccctgatgaatcccttc | Reverse primer to amplify the downstream of *Aave_1468* for constructing in-frame deletion of *Aave_1468* |
| Fha-KO5 | gaatgcaggccgaagaggc | Forward primer to confirm the in-frame deletion of *Aave_1468* |
| Fha-KO6 | tgcgaacacggcgaaggt | Reverse primer to confirm the in-frame deletion of *Aave_1468* |
| TssJ-KO1 | taccgaattcgagctcgagcatatcgcaggcgaggagttcc | Forward primer to amplify the upstream of *Aave_1470* for constructing in-frame deletion of *Aave_1470* |
| TssJ-KO2 | ccgcatcgatgaatcccttccgtgccat | Reverse primer to amplify the upstream of *Aave_1470* for constructing in-frame deletion of *Aave_1470* |
| TssJ-KO3 | gaagggattcatcgatgcggtgaagatccagacc | Forward primer to amplify the downstream of *Aave_1470* for constructing in-frame deletion of *Aave_1470* |
| TssJ-KO4 | ctgcaggtcgactctgagatctatcttcagcagatcgtcgaacag | Reverse primer to amplify the downstream of *Aave_1470* for constructing in-frame deletion of *Aave_1470* |
| TssJ-KO5 | gctgtgggacgagtacaagcg | Forward primer to confirm the in-frame deletion of *Aave_1470* |
| TssJ-KO6 | cgcagatcgtcgtggtcgtag | Reverse primer to confirm the in-frame deletion of *Aave_1470* |
| TssL-KO1 | taccgaattcgagctcgagcccttcattcccacggtggtc | Forward primer to amplify the upstream of *Aave_1472* for constructing in-frame deletion of *Aave_1472* |
| TssL-KO2 | cctcgaagcctcactatcaaacgttgacggagttctgcatg | Reverse primer to amplify the upstream of *Aave_1472* for constructing in-frame deletion of *Aave_1472* |
| TssL-KO3 | cgtttgatagtgaggcttcgagggacgctatcac | Forward primer to amplify the downstream of *Aave_1472* for constructing in-frame deletion of *Aave_1472* |
| TssL-KO4 | ctgcaggtcgactctgagatctggaccatcagcacgatctccac | Reverse primer to amplify the downstream of *Aave_1472* for constructing in-frame deletion of *Aave_1472* |
| TssL-KO5 | aggacgtcaccgacggatg | Forward primer to confirm the in-frame deletion of *Aave_1472* |
| TssL-KO6 | tcagctgcagctcggcatc | Reverse primer to confirm the in-frame deletion of *Aave_1472* |
| TssA-KO1 | taccgaattcgagctcgagccaccgtttcttcgacaaggc | Forward primer to amplify the upstream of *Aave_1475* for constructing in-frame deletion of *Aave_1475* |
| TssA-KO2 | ggattttgtccctcttcgatgtcgatcatgatgg | Reverse primer to amplify the upstream of *Aave_1475* for constructing in-frame deletion of *Aave_1475* |
| TssA-KO3 | gacatcgaagagggacaaaatccgccatcctg | Forward primer to amplify the downstream of *Aave_1475* for constructing in-frame deletion of *Aave_1475* |
| TssA-KO4 | ctgcaggtcgactctgagatctctggtatgcatcgtccgacag | Reverse primer to amplify the downstream of *Aave_1475* for constructing in-frame deletion of *Aave_1475* |
| TssA-KO5 | cctccatttcgcagatggtg | Forward primer to confirm the in-frame deletion of *Aave_1475* |
| TssA-KO6 | tgatctgctcggagagcttgc | Reverse primer to confirm the in-frame deletion of *Aave_1475* |
| TssB-KO1 | taccgaattcgagctcgagctggatgttcgcccgttactg | Forward primer to amplify the upstream of *Aave_1476* for constructing in-frame deletion of *Aave_1476* |

| | | |
|---|---|---|
| TssB-KO2 | tgagcgtcgtccttgcttaaggtcaccatagtgaaatcttc | Reverse primer to amplify the upstream of *Aave_1476* for constructing in-frame deletion of *Aave_1476* |
| TssB-KO3 | ccttaagcaaggacgacgctcagtgaaagggac | Forward primer to amplify the downstream of *Aave_1476* for constructing in-frame deletion of *Aave_1476* |
| TssB-KO4 | ctgcaggtcgactctgagatctaggccgatgtacttcgagtcg | Reverse primer to amplify the downstream of *Aave_1476* for constructing in-frame deletion of *Aave_1476* |
| TssB-KO5 | caagccggagcagcagttc | Forward primer to confirm the in-frame deletion of *Aave_1476* |
| TssB-KO6 | gttcgacttcgaggaggaaacg | Reverse primer to confirm the in-frame deletion of *Aave_1476* |
| TssF-KO1 | taccgaattcgagctcgagccaacggcagctattactgggtg | Forward primer to amplify the upstream of *Aave_1480* for constructing in-frame deletion of *Aave_1480* |
| TssF-KO2 | cgctaccagccgcggattcatgtca | Reverse primer to amplify the upstream of *Aave_1480* for constructing in-frame deletion of *Aave_1480* |
| TssF-KO3 | aatccgcggctggtagcgcgctggaaggtg | Forward primer to amplify the downstream of *Aave_1480* for constructing in-frame deletion of *Aave_1480* |
| TssF-KO4 | ctgcaggtcgactctgagatctcaggatcagctcgtaatcccatg | Reverse primer to amplify the downstream of *Aave_1480* for constructing in-frame deletion of *Aave_1480* |
| TssF-KO5 | gaggcgttcgagtggatcg | Forward primer to confirm the in-frame deletion of *Aave_1480* |
| TssF-KO6 | accacgtcgtccatcccag | Reverse primer to confirm the in-frame deletion of *Aave_1480* |
| sfGFP-X-hifif | tctaaaggtgaagaactgttcaccg | Forward primer to amplify *sfGFP* |
| sfGFP-X-hifir | tcctgcggccgctttgtagag | Reverse primer to amplify *sfGFP* |
| sfGFP-TssL-KI1 | taccgaattcgagctcgagctcgggaacatccagctgtcg | Forward primer to amplify the upstream of *Aave_1472* for constructing chromosomal *sfGFP_tssL* |
| sfGFP-TssL-KI2 | cggtgaacagttcttcacctttagacatggacatgtccgctttcag | Reverse primer to amplify the upstream of *Aave_1472* for constructing chromosomal *sfGFP_tssL* |
| sfGFP-TssL-KI3 | ctacaaagcggccgcaggaggaggacagaactccgtcaacgttttcg | Forward primer to amplify *Aave_1472* for constructing chromosomal *sfGFP_tssL* |
| sfGFP-TssL-KI4 | ctgcaggtcgactctgagatctctggatcaccgcgacatagc | Reverse primer to amplify *Aave_1472* for constructing chromosomal *sfGFP_tssL* |
| sfGFP-TssL-KI5 | cggacgtcaacgacatggg | Forward primer to confirm the chromosomal *sfGFP_tssL* |
| sfGFP-TssL-KI6 | ccttgccggacacttcgttg | Reverse primer to confirm the chromosomal *sfGFP_tssL* |
| sfGFP-TssM-KI1 | taccgaattcgagctcgagcaagttcttccagctgctgtccaag | Forward primer to amplify the upstream of *Aave_1473* for constructing chromosomal *sfGFP_tssM* |
| sfGFP-TssM-KI2 | cggtgaacagttcttcacctttagacatgttttagttcttcggattgcc | Reverse primer to amplify the upstream of *Aave_1473* for constructing chromosomal *sfGFP_tssM* |
| sfGFP-TssM-KI3 | ctacaaagcggccgcaggaggaggactgcgcaagatcttttcattcttcttc | Forward primer to amplify *Aave_1473* for constructing chromosomal *sfGFP_tssM* |
| sfGFP-TssM-KI4 | ctgcaggtcgactctgagatcttgccgaagaactcgttgaagc | Reverse primer to amplify *Aave_1473* for constructing chromosomal *sfGFP_tssM* |
| sfGFP-TssM-KI5 | gcgctgcacaacgaaacctg | Forward primer to confirm the chromosomal *sfGFP_tssM* |
| sfGFP-TssM-KI6 | gggattggtgaagccccagac | Reverse primer to confirm the chromosomal *sfGFP_tssM* |

| | | |
|---|---|---|
| sfGFP-TssG-KI1 | taccgaattcgagctcgagcggaccgcaacaaatttctcctg | Forward primer to amplify the upstream of *Aave_1481* for constructing chromosomal *sfGFP_tssG* |
| sfGFP-TssG-KI2 | acctttagacattcatgccgcgggcctctg | Reverse primer to amplify the upstream of *Aave_1481* for constructing chromosomal *sfGFP_tssG* |
| sfGFP-TssG-KI3 | gtacaaggcggccgcaggaggaggaatgagcgaccgtgtgcc | Forward primer to amplify *Aave_1481* for constructing chromosomal *sfGFP_tssG* |
| sfGFP-TssG-KI4 | ctgcaggtcgactctgagatctgtccagccggaaatcgtc | Reverse primer to amplify *Aave_1481* for constructing chromosomal *sfGFP_tssG* |
| sfGFP-TssG-KI5 | ctgcacgagtatttcgcctttc | Forward primer to confirm the chromosomal *sfGFP_tssG* |
| sfGFP-TssG-KI6 | gacctcgcggatggaaatg | Reverse primer to confirm the chromosomal *sfGFP_tssG* |
| Fha-sfGFP-KI1 | taccgaattcgagctcgagctggaaggactcgaccttggtg | Forward primer to amplify *Aave_1468* for constructing chromosomal *fha_sfGFP* |
| Fha-sfGFP-KI2 | ctgcggccgccgtcctagccttagccgcag | Reverse primer to amplify *Aave_1468* for constructing chromosomal *fha_sfGFP* |
| Fha-sfGFP-KI3 | gctgtacaagtaatcttccatatcgcaggcgag | Forward primer to amplify the downstream of *Aave_1468* for constructing chromosomal *fha_sfGFP* |
| Fha-sfGFP-KI4 | ctgcaggtcgactctgagatctgttgctgatcatgccgcag | Reverse primer to amplify the downstream of *Aave_1468* for constructing chromosomal *fha_sfGFP* |
| Fha-sfGFP-KI5 | cgacctgtctcagtcgttca | Forward primer to confirm the chromosomal *fha_sfGFP* |
| Fha-sfGFP-KI6 | tagcttgcccggtcgaaag | Reverse primer to confirm the chromosomal *fha_sfGFP* |
| Fha-sfGFP-hifif | ctaggacggcggccgcaggaggag | Forward primer to amplify *sfGFP* |
| Fha-sfGFP-hifir | ctgcgatatggaagattacttgtacagctcgtccatgcc | Reverse primer to amplify *sfGFP* |
| TssB-mCherry2-KI1 | taccgaattcgagctcgagccacaactgctgctggtggc | Forward primer to amplify the upstream of *Aave_1476* and *Aave_1476* for constructing chromosomal *tssB_mCherry2* |
| TssB-mCherry2-KI2 | tgcggccgcctgagcgtcgtcggtcttctc | Reverse primer to amplify the upstream of *Aave_1476* and *Aave_1476* for constructing chromosomal *tssB_mCherry2* |
| TssB-mCherry2-KI3 | ctgtacaagtaaaagggacagccgcatgaac | Forward primer to amplify the downstream of *Aave_1476* for constructing chromosomal *tssB_mCherry2* |
| TssB-mCherry2-KI4 | ctgcaggtcgactctgagatctccatcgcataggcggagttc | Reverse primer to amplify the downstream of *Aave_1476* for constructing chromosomal *tssB_mCherry2* |
| TssB-mCherry2-KI5 | tgctggtggtcctgaacga | Forward primer to confirm the chromosomal *tssB_mCherry2* |
| TssB-mCherry2-KI6 | tagtattcggccggcttgtg | Reverse primer to confirm the chromosomal *tssB_mCherry2* |
| TssB-mCherry2-hifif | acgctcaggcggccgcaggaggag | Forward primer to amplify *mCherry2* |
| TssB-mCherry2-hifir | tgcggccgcctgagcgtcgtcggtcttctc | Reverse primer to amplify *mCherry2* |
| mCherry2-TssL hifif | atggtgagcaagggcgagga | Forward primer to amplify *mCherry2* |
| mCherry2-TssL hifir | aaaacgttgacggagttctgtcctcctcctgcggccgccttgtacagctcgtcca | Reverse primer to amplify *mCherry2* |
| mCherry2-TssL-KI2 | tcctcgcccttgctcaccatggacatgtccgctttcag | Reverse primer to amplify the upstream of *Aave_1472* for constructing chromosomal *mCherry2_tssL* |
| mCherry2-TssL-KI3 | cagaactccgtcaacgttttcg | Forward primer to amplify *Aave_1472* for constructing chromosomal *mCherry2_tssL* |

| | | |
|---|---|---|
| Fha-mCherry2-hifif | taaggctaggacggcggccgcaggaggag | Forward primer to amplify *mCherry2* |
| Fha-mCherry2-hifir | gatatggaagattacttgtacagctcgtccatgccg | Reverse primer to amplify *mCherry2* |
| Fha-mCherry2-KI2 | tgcggccgccgtcctagccttagccgcag | Reverse primer to amplify *Aave_1468* for constructing chromosomal *fha_mCherry2* |
| Fha-mCherry2-KI3 | gagctgtacaagtaatcttccatatcgcaggcgag | Forward primer to amplify the downstream of *Aave_1468* for constructing chromosomal *fha_mCherry2* |
| pBBR2-hifiF | agctgtttcctgtgtgaaattg | Forward primer to amplify pBBRMCS2 vector |
| pBBR2-3V5-hifiR | ggtaaacctattcctaatcctctcct | Reverse primer to amplify pBBRMCS2 vector |
| pBBR2-f | agcgcaacgcaattaatgtgag | Forward confirmation primer of pBBRMCS2 vector |
| pBBR2-r | catcgcagtacggcctattgg | Reverse confirmation primer of pBBRMCS2 vector |
| pBBR-Fha-3V5-F | ggattaggaataggtttacccgtcctggccttcgccg | Forward primer to amplify *Aave_1468* |
| pBBR-Fha-3V5-R | atttcacacaggaaacagctatggacaaagtcgaactcagggtcg | Reverse primer to amplify *Aave_1468* |
| 1468-flag-hifir | cacttgtcatcgtcgtccttgtaatccgtcctggccttcgccg | Reverse primer to amplify *Aave_1468* |
| pETDuet-f | cacgatgcgtccggcgtagagg | Forward confirmation primer of pET vector |
| pETDuet-r | ggttatgctagttattgctcagcggt | Reverse confirmation primer of pET vector |
| pETSUMO-hifi-f | aagcttaggtatttattcggcgcaaagtg | Forward primer to amplify pETSUMO vector |
| pETSUMO-hifi-r | ggtaccggatccaccaccaatctg | Reverse primer to amplify pETSUMO vector |
| pETSUMO-TssM-f | ttggtggtggatccggtaccatgctgcgcaagatcttttcattcttc | Forward primer to amplify *Aave_1473* |
| pETSUMO-TssM-r | cgccgaataaatacctaagctttcagagattgccaggacacgcga | Reverse primer to amplify *Aave_1473* |
| pETSUMO-TssE-f | cagattggtggtggatccggtaccatggcgacgcagagcaggga | Forward primer to amplify *Aave_1479* |
| pETSUMO-TssE-r | cgccgaataaatacctaagcttccgctgctcggacacggac | Reverse primer to amplify *Aave_1479* |
| pETSUMO-TssF-f | ttggtggtggatccggtaccatgaatccgcggctggtcga | Forward primer to amplify *Aave_1480* |
| pETSUMO-TssF-r | cgccgaataaatacctaagctttgccgcgggcctctggcc | Reverse primer to amplify *Aave_1480* |
| pETSUMO-TssG-f | ttggtggtggatccggtaccatgagcgaccgtgtgccc | Forward primer to amplify *Aave_1481* |
| pETSUMO-TssG-r | cgccgaataaatacctaagcttcgacgcttcggccgccggga | Reverse primer to amplify *Aave_1481* |
| pETSUMO-TssK-f | ttggtggtggatccggtaccatggtttggcagcgcaaggt | Forward primer to amplify *Aave_1471* |
| pETSUMO-TssK-r | cgccgaataaatacctaagcttgcgccgtatggcccacatttc | Reverse primer to amplify *Aave_1471* |
| pET22b-hifi-f | aagcttgcggccgcactcga | Forward primer to amplify pET22b vector |
| pET22b-hifi-r | catatgtatatctccttcttaaagttaaacaaaattatttctagagg | Reverse primer to amplify pET22b vector |
| 22b-TssJ-f | ctttaagaaggagatatacatatgcatcgggaagcgatgg | Forward primer to amplify *Aave_1470* |
| 22b-TssJ-r | tcgagtgcggccgcaagcttaggagcggtctggatcttcacc | Reverse primer to amplify *Aave_1470* |
| 22b-TssL-f | ctttaagaaggagatatacatatgcagaactccgtcaacgt | Forward primer to amplify *Aave_1472* |
| 22b-TssL-r | tcgagtgcggccgcaagcttgttcttcggattgccgg | Reverse primer to amplify *Aave_1472* |
| 22b-Fha-f | ttaagaaggagatatacatatggacaaagtcgaactcagggt | Forward primer to amplify *Aave_1468* |

| | | |
|---|---|---|
| 22b-Fha-r | gagtgcggccgcaagcttcgtcctggccttcgccg | Reverse primer to amplify *Aave_1468* |
| 22b-TssA-f | ctttaagaaggagatatacatatgatcgacatcgaagagcttcttga | Forward primer to amplify *Aave_1475* |
| 22b-TssA-r | tcgagtgcggccgcaagcttggatggcggattttgtcctcc | Reverse primer to amplify *Aave_1475* |
| pKNT/CH-hifi-f | actagtgcggtaccgagctcgaattc | Forward primer to amplify pKNT25 and pCH363 vector |
| pKN/CH-hifi-r | catagctgtttcctgtgtgaaattgttatcc | Reverse primer to amplify pKNT25 and pCH363 vector |
| pKNT/CH N-hifi-f | gcgagagagggtaccgaattc | Forward primer to amplify pKNT25N and pCH363N vector |
| pKNT/CH N-hifi-r | tctagagcatgcaagctttgcg | Reverse primer to amplify pKNT25N and pCH363N vector |
| pCH363N-f | cgaagttctcgccggatgtac | Forward confirmation primer of pCH363N vector |
| pCH363N-r | cgaaatacgggcagacatgg | Reverse confirmation primer of pCH363N vector |
| pKNT25N-f | cattatgccgcatctgtccaac | Forward confirmation primer of pKNT25N vector |
| pKNT25N-r | tggcttaactatgcggcatcag | Reverse confirmation primer of pKNT25N vector |
| pCH363-f | ggaattgtgagcggataaca | Forward confirmation primer of pCH363 vector |
| pCH363-r | cgattttccacaacaagtcg | Reverse confirmation primer of pCH363 vector |
| pKNT25f | gcggataacaatttcacacagg | Forward confirmation primer of pKNT25 vector |
| pKNT25r | gtttgcgtaaccagcctgat | Reverse confirmation primer of pKNT25 vector |
| TssJ-dsp-hifi-f | ggaaacagctatgtgcggcatgatcagcaacct | Forward primer to amplify *Aave_1470* |
| TssJ-hifi-r | gagctcggtaccgcactagtaggagcggtctggatcttcacc | Reverse primer to amplify *Aave_1470* |
| TssM cyto2-pck-hifif | caaagcttgcatgctctagaagatggtggagagctaaggagatgaatgcacgcct g | Forward primer to amplify *Aave_1473* |
| TssMcyto2-pck-hifir | aattcggtaccctctctcgctcaccgcaactggcgctccttc | Reverse primer to amplify *Aave_1473* |
| TssM-peri-hifif | catgctctagaagccacggcaacaaccagaaatac | Forward primer to amplify *Aave_1473* |
| TssM-peri-hifir | aattcggtaccctctctcgctcagagattgccaggacacg | Reverse primer to amplify *Aave_1473* |
| pCH363N-TssL-hifif | caaagcttgcatgctctagacagaactccgtcaacgt | Forward primer to amplify *Aave_1472* |
| pCH363N-TssL-cyto-r | cctctctcgcttacaggctccagggcggcac | Reverse primer to amplify *Aave_1472* |
| pCH363N-TssL -peri-f | ctctagaagacttgcttctcgctcggacgagctgttcg | Forward primer to amplify *Aave_1472* |
| pCH363N-TssL-hifir | aattcggtaccctctctcgcttagttcttcggattgcc | Reverse primer to amplify *Aave_1472* |
| Fha-pC/K-hifi-f | tcacacaggaaacagctatggacaaagtcgaactcagggtcgt | Forward primer to amplify *Aave_1468* |
| Fha-pC/K-hifi-r | gagctcggtaccgcactagtccttgccttagcagccgcctcggcgttttcatag | Reverse primer to amplify *Aave_1468* |
| Fha-dN-fd | cggacatgcattccgtcgtgccgggct | Forward primer to amplify *Aave_1468* |
| Fha-dN-rv | acgacggaatgcatgtccgctacgacccg | Reverse primer to amplify *Aave_1468* |
| Fha-dIDR-fd | gtcctgagtcctctggcaccggtgctgccggaactgg | Forward primer to amplify *Aave_1468* |

| | | |
|---|---|---|
| Fha-dIDR-rv | ggtgccagaggactcaggacgtagggcccgatg | Reverse primer to amplify *Aave_1468* |
| sfgfp-pbbr-fd | ggaaacagctatggcaggaggaggatctaaaggtgaagaac | Forward primer to amplify *sfGFP* |
| TssM-pbbr-hifir | attaggaataggtttacctcagagattgccaggacacgc | Reverse primer to amplify *Aave_1473* |
| TssL-pbbr-hifir | ggattaggaataggtttaccgttttagttcttcggattgccg | Reverse primer to amplify *Aave_1472* |
| TssA-pbbr-hifir | attaggaataggtttaccttatcaggatggcggattttgtcc | Reverse primer to amplify *Aave_1475* |
| TssE-sfgfp-hifi-f | atttcacacaggaaacagctatggcgacgcagagcaggga | Forward primer to amplify *Aave_1479* |
| TssE-sfgfp-hifi-r | gatcctcctcctgcggccgcccgctgctcggacacggac | Reverse primer to amplify *Aave_1479* |
| Hcp-pbbr-hifif | atttcacacaggaaacagctatgtccgtcgatatgttcatgaagg | Forward primer to amplify *Aave_1465* |
| Hcp-sfGFP-hifir | agatcctcctcctgcagcggcttctttgttttccttgatgttcc | Reverse primer to amplify *Aave_1465* |
| ClpV-pbbr-hifif | atttcacacaggaaacagctatgtctgaaatcagccggcaagc | Forward primer to amplify *Aave_1483* |
| ClpV-sfgfp-hifir | gatcctcctcctgcagcggcctcttcgtactcgatggc | Reverse primer to amplify *Aave_1483* |
| TssB-sfgfp-hifi-f | atttcacacaggaaacagctatggtgaccttaagcaagaccg | Forward primer to amplify *Aave_1476* |
| TssB-sfgfp-hifi-r | agatcctcctcctgcggccgcctgagcgtcgtcggtcttctcg | Reverse primer to amplify *Aave_1476* |
| EagT2-sfgfp-hifi-f | atttcacacaggaaacagctatgcagtaccacatccaggaagcc | Forward primer to amplify *Aave_0498* |
| EagT2-sfgfp-hifi-r | agatcctcctcctgcagcggcgcgcaggcgcaggctgtgca | Reverse primer to amplify *Aave_0498* |
| RhsB-sfgfp-hifi-f | atttcacacaggaaacagctatgcccaaggccgcccgactg | Forward primer to amplify *Aave_0499* |
| RhsB-sfgfp-hifi-r | agatcctcctcctgcagcggcccaaccccagggatcgatccaggaca | Reverse primer to amplify *Aave_0499* |
| RimB1-sfgfp-hifi-f | tcacacaggaaacagctatgcaggatttttggtatccccgagtc | Forward primer to amplify *rimB1* |
| RimB1-sfgfp-hifi-r | agatcctcctcctgcagcggcgccgttttccaatatcttcttcagg | Reverse primer to amplify *rimB1* |
| helix-mu1-fd | cgcagaagctcgtgaacgtcgagaccctgcagctctggagcagcgcaccgacgt | Forward primer to amplify *Aave_1468* |
| helix-mu1-rv | gacgttcacgagcttctgcgtaagctgcgttgtcacctgccatcagggccacctgg tg | Reverse primer to amplify *Aave_1468* |
| helix-mu2-fd | gcagatgacgaggctctggagcgtgacgaccctgcagctctggagcagcgcac cgacgt | Forward primer to amplify *Aave_1468* |
| helix-mu2-rv | ctccagagcctcgtcatctgcagcacgcagtccagcgtccagggccacctggtg gacc | Reverse primer to amplify *Aave_1468* |
| helix-mu3-fd | acgtgcagaggctcgtcagcgcttcgaccctgcagccctggagcagcgcacc | Forward primer to amplify *Aave_1468* |
| helix-mu3-rv | gctgacgagcctctgcacgtgcagctcgacgtccagccatcagggccacctggt gga | Reverse primer to amplify *Aave_1468* |
| V4L6V8-SSS-fd | ggacaaatccgaatccaggagcgtccgccatgcggacgaa | Forward primer to amplify *Aave_1468* |
| V4L6V8-SSS-rv | tcctggattcggatttgtccatatgtatatctccttcttaaagttaaac | Reverse primer to amplify *Aave_1468* |
| F24S-fd | cggtgtagcaggtggtactattggtcgcggagggcagaacaagc | Forward primer to amplify *Aave_1468* |
| F24S-rv | ccaatagtaccacctgctacaccggataccgccgtccataagcggt | Reverse primer to amplify *Aave_1468* |
| T30-I31-R33-SSN-fd | ggtggatccagcggtaacggagggcagaacaagctcgtg | Forward primer to amplify *Aave_1468* |

| | | |
|---|---|---|
| T30-I31-R33-SSN-rv | ccgttaccgctggatccacccgccacgccgaatacc | Reverse primer to amplify *Aave_1468* |
| N37-L39-L41-D43-SSSN-fd | agcaagagcgtgtcctccaacagcgatgcgggtgtcg | Forward primer to amplify *Aave_1468* |
| N37-L39-L41-D43-SSSN-rv | gttggaggacacgctcttgctctgccctccgcggccgat | Reverse primer to amplify *Aave_1468* |
| V48-R50-V51-SNS-fd | ggtagcgccaacagccatgcgatggtgcgcatcg | Forward primer to amplify *Aave_1468* |
| V48-R50-V51-SNS-rv | catggctgttggcgctacccgcatcgctgtcggac | Reverse primer to amplify *Aave_1468* |
| A53-V55-SS-fd | tgcattccatgtcccgcatcgaatccgaggcc | Forward primer to amplify *Aave_1468* |
| A53-V55-SS-rv | atgcgggacatggaatgcacgcgggcgacac | Reverse primer to amplify *Aave_1468* |
| I64S-fd | gaggcagctttcagcgccaatctctgcgagcg | Forward primer to amplify *Aave_1468* |
| I64S-rv | tggcgctgaaagctgcctcggattcgatgcgca | Reverse primer to amplify *Aave_1468* |
| I73-V75-SS-fd | aggtcgtcgttctcggctggtgccgaagtgcgctcg | Forward primer to amplify *Aave_1468* |
| I73-V75-SS-rv | cagccgagaacgacgacctgcgctcgcagagattggc | Reverse primer to amplify *Aave_1468* |
| V80-G83-SS-fd | tcttcgttgctggagctgagtcgaggtcgtccgaagaggtgcgcctggca | Forward primer to amplify *Aave_1468* |
| V80-G83-SS-rv | ctcagctccagcaacgaagatgctcctgcgctcgcagagattggcgat | Reverse primer to amplify *Aave_1468* |
| L88-G91-SS-fd | tgaggtcggctgtctcggacgaggtccgcatcgg | Forward primer to amplify *Aave_1468* |
| L88-G91-SS-rv | gtccgagacagccgacctcacctcttcgcccgagcg | Reverse primer to amplify *Aave_1468* |
| V94-I96-G97-Y99-L101-fd | gaggtccggtccttcggtctcgcattccgtcgtgccggg | Forward primer to amplify *Aave_1468* |
| V94-I96-G97-Y99-L101-rv | gaccgaaggaccggacctcgactcgtcccccactgccag | Reverse primer to amplify *Aave_1468* |
| P254-P256-SS-fd | tccatgtccgcttcggacggggtccatggtttc | Forward primer to amplify *Aave_1468* |
| P254-P256-SS-rv | cgtccgaagcggacatggacgcggaccgggat | Reverse primer to amplify *Aave_1468* |
| I303-P304-SS-fd | ccctcgtctcgtccgaagacttcgatcttttcgcggtg | Forward primer to amplify *Aave_1468* |
| I303-P304-SS-rv | gtcttcggacgagacgagggcccgggggc | Reverse primer to amplify *Aave_1468* |
| D306-F307-F310-A311-NSSS-fd | gaattccgacctttcctcggtggaccccccgcaagg | Forward primer to amplify *Aave_1468* |
| D306-F307-F310-A311-NSSS-rv | ccgaggaaaggtcggaattctccggaatgacgagggc | Reverse primer to amplify *Aave_1468* |
| L333-I336-SS-fd | agctcgtccgagtcggccaacatgcggcacgac | Forward primer to amplify *Aave_1468* |
| L333-I336-SS-rv | ggccgactcggacgagctcttagcctgcaaaccgc | Reverse primer to amplify *Aave_1468* |
| L377-D378-P379-L380-L382-F383-fd | gcaacagctcgagctcttcgacgggcggctcggatgc | Forward primer to amplify *Aave_1468* |

| | | |
|---|---|---|
| L377-D378-P379-L380-L382-F383-rv | cgaagagctcgagctgttgctttcgtcgcgcgggtccag | Reverse primer to amplify *Aave_1468* |
| L408S-fd | ccgactcttcgcagtcgttcagccttccgagg | Forward primer to amplify *Aave_1468* |
| L408S-rv | gaacgactgcgaagagtcggaccccgaaccgagca | Reverse primer to amplify *Aave_1468* |
| F412S-fd | ctcagagctcgtcgcttccgaggggcgtgcag | Forward primer to amplify *Aave_1468* |
| F412S-rv | cggaagcgacgagctctgagacaggtcggaccccgaa | Reverse primer to amplify *Aave_1468* |
| P415S-fd | ttctccctcacccgtggtgtgcaggcgccggtggg | Forward primer to amplify *Aave_1468* |
| P415S-rv | acaccacgggtgagggagaacgactgagacaggtcggacc | Reverse primer to amplify *Aave_1468* |
| PA Fha-22b-hifif | ctttaagaaggagatatacatatgccgctgcgattgacca | Forward primer to amplify *PA0081* |
| PA Fha-sfGFP-hifir | agatcctcctcctgcagctgcggaacgccgtagtcgagcg | Reverse primer to amplify *PA0081* |
| VC Fha-22b-hifif | ctttaagaaggagatatacatatgaactcagtgacattaccttcgttgac | Forward primer to amplify *VCA0112* |
| VC Fha-sfGFP-hifir | gatcctcctcctgcagctgctagctccagttgcttctcgcgaattttgc | Reverse primer to amplify *VCA0112* |

**Note for references**
