## Supplementary Data 1 for "LLPS condensates of Fha initiate the inside-out assembly of the type VI secretion system"

```
                       β1      β2       β3     β4      β5         β6
ABM32059.1          ─────────→TT─→   ──────→TT ───→    TT  ─→       ───
                         1        10        20        30        40        50
ABM32059.1         .....MDKVELRVVRHADEAVDRLWTAVFGVAGGTIGRGGQNKLVLSDSDAGVARVHAMV
NP_248771.1        ......MPLRLTITSYHKLTPGQCSEKVLDQGQLTIGRGPDNDWVLPDPERLVSSRHCTI
WP_001089704.1     MNSVTLPSLTLLVTNAQHLESGLSAQHTWTSSGGVIGSSPNAQWRLVDAQGSVKPMHCEV
WP_200016399.1     ............................................................
WP_063326054.1     .....MDKIELRVVDHDGVPVDRHWHAVFGMAGGTIGRSGQNKLIISDPDASVARVHAMV
WP_012205666.1     .....MDKIELRVVDHDGVPVDRHWHAVFGMAGGTIGRSGQNKLIISDPDASVARVHAMV
MBK9235436.1       .....MDTIELHVAQHAGAALGRSWHATFGMAGGTIGRSGQNKLVLPDVDAGVARVHAMV
MCA0243819.1       .....MDKIEIRVESFNGEAVTKPWSAVFGLAGGTIGRGGQNKLVLPDPDAHVARVHAMV
MCZ2441668.1       .....MDKIEIRVEGQHGQPVTKPWAALFGLAGGTIGCGSQNKLVLPDIDTQVDRVHAMV
WP_202760356.1     ............................................................
WP_252978133.1     ............................................................
WP_065346974.1     .....MDKIELRVVDHDGVPVDRHWHAVFGMAGGTIGRSGQNKLIISDPDASVARVHAMV
WP_230355409.1     .....MDKIELRVVDHDGVPVDRHWHAVFGMAGGTIGRSGQNKLIISDPDASVARVHAMV
WP_267179154.1     .....MDKIELRVVDHDGVAVDRHWYAVFGMAGGTIGRSGQNKLIISDPDASVARVHAMV
WP_088164707.1     .....MDKIELRVVDHDGVPVDRHWHAVFGMAGGTIGRSGQNKLIISDPDASVARVHAMV
WP_133073230.1     .....MDKIELRVVDHDGVAVDRHWHAVFGMAGGTIGRSGQNKLIISDPDASVARVHAMV
MCC7098932.1       .....MDTIEIRVEAHDGVPVAKPWAAVFALSGGTIGRGSQNKLVLPEADAQVDPVHAMV
WP_238004152.1     .....MDKIELRVVDHDGVPVDRHWHAVFGMAGGTIGRSGQNKLIISDPDASVARVHAMV
MBV8620386.1       .....MEKIEIRVERALGVPVDRHWHAVFGVAGGTIGRGGQNKLVLPDADAGVARVHAMV
ODU08341.1         .....MEQIEIRVEGHHGQPVTKPWAALFGLAGGTIGCGGQNKLVLPDAEAQVDRVHAMV
MBC7710584.1       .....MAALTLQVVSIDAAPVGQPTMHAFEATGGSIGRDETNLLALPDKHRRVSRLHATI
MBL8310054.1       .....MSSITLTVMSIDASAPLQRIAQTFSGTGGTIGRDDSNTLALPDKHRRVSRLHGTV
WP_238363320.1     ............................................................
MCX8099153.1       .....MNSLTLTVVTIDAVAPSVPISATFGEQGGTIGRDENSTLQLADKHRRVSRLHGTI
MCM2252316.1       .....MPQVTLDLVSLDDKPV.AGASCRFEGAGGTIGRDEGNTLALPDPHRRVSRLHATI
RYF42966.1         .....MPHIILDLISLDGKPL.NGLSRRFEGEGGTIGRDEGNTLSLPDSHRRVSRLHAAI
WP_281607349.1     .......MLTIKAVSYNGQATEVPLAGEFGELGGVIGRADGNALVLPDPDRYVSRMHAMV
MDB5976135.1       ............................................................
WP_201212573.1     .......MITIRVVSCNGEAPARPLAVEFDELGGTIGRAEMNQLVLPDPERHISRLQARV
MBP8267915.1       ........MRISVISFNGAPPAQPTGFDFDSRGGAIGREEGNELTLPDPERHISRVQARI
MCZ2135183.1       .........................MALFDARGGTIGRDEGNSMALPDRHRRVSRLHATV
WP_131145415.1     .......MLTIKAVSYNGQATDVPLAGEFGELGGVIGRADGNALVLPDPDRYVSRMHAMV
WP_166302793.1     ......MQLRLKAQAAYANETPPPIEGLFDTAGGTIGRDPRCHMVLPDPMRRISRIQVQI
WP_267727965.1     ........MRLTVTSFKGSSAPQPIQAEFGPLGGSIGREPGNALVLPDPERHISRLHAQV
WP_299903544.1     ........MRISVISFNGAPPARPLAYDFDSRGGAIGREEGNELTLPDPERHISRVQARI
WP_065341524.1     .......MLRISVIQVNGFPPATPLAGEFDERGGTIGRDDSNALTLPDPGRHISRMQAQV
MBL8494843.1       .......MMRIRVISLHGQPLAQEIAHDFDEAGGTLGRAETNQLALPDAQRHISRLQGRI
WP_058641008.1     .......MIRIRVTGHQGRMAPG.AMATFGAAGGTIGRAPQNALVLDDPDRTVSRVHAQV
WP_183632487.1     ........MRLIVTSFKGLPPLQPIEAEFGQSGGSIGREPGNQLVLPDPERHISRLHAQV
RYY83951.1         ............................................................
MCU0967987.1       ............................................................
WP_106557035.1     .......MIRIAVISHQGAPAVRKLAAQFGADGGTIGRAPTNALVLDDPDRTVSRVHAQV
MBK7663138.1       .......MMKIRVITFQGQAVPHAISHDFDELGGTIGRAETNQLALPDPQRHISRLQARV
WP_203203692.1     ........MRWTVIEHAGRPVAAGPSALLAAPGGTIGRSPDNHLVLPDEQRQISRLQATV
WP_295989070.1     ............................................................
MBT9459293.1       .......MITLTVTSFNGAPSDIPLSASFDELGGTIGRADTNQLVLPDPDRAISRVHAQI


                       β7               β8     β9        β10       β11     β12
ABM32059.1         ──→T.T─────→    TT   ──→  TT──→  TT  ───→   TT  ───→TT───→    TT
                         60        70        80        90        100       110
ABM32059.1         RIES.EAAFIANLCERRSIFV.AGAEVRSGEEVRLAVGDEVRIGPYVLHSVVPGSPFVPV
NP_248771.1        LNRD.GVYYLTDTSTNGVLLVNAGHRLRRGNSEPLQDGETVRLGEYDILVQLGHDIALPG
WP_001089704.1     MMVD.GAYCLKDSCGSTYVNG.ADMPLGVGQMARLTHKDEVSVGPYRLRVLFGNAIEVEE
WP_200016399.1     ............................................................
WP_063326054.1     RLDS.EGAFVANLSERDPIWV.AEQPVRSGQEVGLPLGARIRIGAYTLAAVAPGTPFEAP
WP_012205666.1     RLDS.EGAFVANLSERDPIWV.AEQPVRSGQEVGLPLGARIRIGAYTLAAVAPGTPFEAP
MBK9235436.1       RLDS.ERAYIANLCERRAMWV.DGLEVRSGQEVTLPLGAQVQIGPYVLQAGKLGATRLTD
MCA0243819.1       RLET.DKALIANLCERRAIVV.GGQELPSGQETVLPIGTQVHVGPYRLRALRPGTPWQAA
MCZ2441668.1       RLQA.GKVYVVNLCERRPLRV.DDEDLAPGQETLLPIGARMQVGPYSLRAALPGSPWAPS
WP_202760356.1     ............................................................
WP_252978133.1     ............................................................
WP_065346974.1     RLDS.EGAFVANLSERDPIWV.AEQPVRSGQEVGLPLGARIRIGAYTLAAVAPGTPFEAP
WP_230355409.1     RLDS.EGAFVANLSERDPIWV.AEQPVRSGQEVGLPLGARIRIGAYTLAAVAPGTPFEAP
WP_267179154.1     RLDS.EGAFVANLSERDPIWV.ADQPVRSGQEVALTLGARIRIGAYTLAAVAPGTPFEAP
WP_088164707.1     RLDS.EGAFVANLSERDPIWV.AEQPVRSGQEVGLPLGARIRIGAYTLAAVAPGTPFEAP
WP_133073230.1     RLDS.EGAFVANLSERDPIWV.ADQPVRSGQEVALTLGARIRIGAYTLAAVAPGTPFEAP
MCC7098932.1       RLHG.GKAFLTNLCERRNLRV.GEAELPPGQETPLAIGSRIQVGPYTLRAALPGGAAADD
WP_238004152.1     RLDS.EGAFVANLSERDPIWV.AEQPVRSGQEVGLPLGARIRIGAYTLAAVAPGTPFEAP
MBV8620386.1       RLES.EGAFIANLCERRSMQV.NDSELRSGQELRLPLGATIVIGPYTLVAVAPGTPVPDF
ODU08341.1         RLQA.GKVYVVNLCERRSLPV.DNEALPPGQETLLSIGARIQIGPYILRAAMPGTAWGPP
MBC7710584.1       SFPG.GVPTITNASTSLPLAV.GEQQLDGGDSMPITQGALIEIGPYILQAHVGNTRTDTL
MBL8310054.1       SFVE.GVATLTNSSTSLPVMV.GEQQLDCGQSTALRHGEQIEIGPYVLVATSDAGAVPRT
WP_238363320.1     ............................................................
MCX8099153.1       RFDG.PVAILTNASMSLPIIV.GDRQLDYGEAVRIEDGDYIEFGPFVVVAHYNSPSAQAA
MCM2252316.1       SFPH.DVPTITNSSTVLTIDA.GGAQLKSGQAMPLAQGMRLGIGPYVLLVSSCEAPAGAE
RYF42966.1         GFPG.GVATITNSSAVLPISV.GGMQLEGGQAMPLLPGMQLEIGPYVLGVRSTEASALAA
WP_281607349.1     LFRN.GRHVIRDLSNASPVFV.NDRPLGHGNEATLADGDELRIGGYTLAVADARVSPAVP
MDB5976135.1       ............................................................
WP_201212573.1     AWRN.GSFEIANV.GANPLDV.AGQSIANGASTTIEPGDRITVGGYLLEVSAGGSGPAPS
MBP8267915.1       EFDG.TRFVLVDQ.GGNPSSV.NDRPVGKGNRVVLAGGDRIAIGGYLLEVEFAREAPVFS
MCZ2135183.1       SFPG.GIPTLTNSSTSLPILV.GDQQLDSGESMLIVNGTQIEIGPYVLLATLSAASLAGL
WP_131145415.1     LFRN.GRHVIRDLSNASPVFV.NDRPLGHGNEATLADGDELRIGGYTLAVADARVSPVVP
WP_166302793.1     LWEN.NAFHLLNASTSNPIYV.NGRELGPGGRSVMGAKEEWRTGNYLIEVEQLADTTVAA
WP_267727965.1     LCEN.GVFIIEDH.GGNPTLI.NGRPVGRGNRMAVADGDTLVIGSYELRAHIDRPAPALS
WP_299903544.1     EFDG.ARFVFVDM.GANPSSV.NDRPVGKGNRVVLAGGERLAIGGYVLEAEFDHAAPVFS
WP_065341524.1     QYDG.SRYVLTDH.GGNPTHL.NGRPLGKGASATLRDGDEIGIADYTLRVEIVRTVSGFG
MBL8494843.1       VCRA.GRFELINQ.GANPINV.DGRPVANGSAAPIGPGTRIEIGGYLMEASGPPAAPPQA
WP_058641008.1     NCRG.GRFFLVDR.GSNPLQC.DGRPVGAGNEVELRGGEQLVIGTFALAVEAAEALPPGP
WP_183632487.1     IWER.GVFLIEDH.GGNPTLI.NGRPVGRGNRTALANGDLLAIGGYELRAEVERAGVNSH
RYY83951.1         .......................................RLVIGSFELDVRAVAASGMPE
MCU0967987.1       ............................................................
WP_106557035.1     LCRD.GRFFIVDR.GSNPMLC.NGHPLGAGHEAALSQGDRLVIGGFELVVEMLSSQHAPA
MBK7663138.1       VCRD.GHYEIINQ.GANPITV.DGHAIGNGSSIPIAAGARIEIGGYLLEALAPPQS....
WP_203203692.1     RFDDEGRAVLRNMSAVLPISV.NGQTLRHDQESPIAEGDRVTIGSYVLEATPANAGTSRA
WP_295989070.1     ............................................................
MBT9459293.1       VFRA.GGFAVVDR.GSNPILV.NGKAVGSGREAPVADGDQLQIGGYVLAIKRSATTAKST
```

*ABM32059.1*

```
                          120          130                 140
ABM32059.1          AAVPAP.VPAPAPEP..VSA...PA....PVANPSSP....A..........SAA...GV
NP_248771.1         ............................................................
WP_001089704.1      QYGSLD.......AL..F..........................................
WP_200016399.1      ............................................................
WP_063326054.1      PSAAMQ.NAQATQAT..QAT...QA....TQAAQATQ....A..........AAPPATAA
WP_012205666.1      PSAAMQ.NAQAAQ.....................ATQ....A..........AAPPATAA
MBK9235436.1        AQASAG.D...GPN................SKTPTSP....A..................
MCA0243819.1        GEALPA.A...........................AP....P..................
MCZ2441668.1        GSGAAE.A.........G.................DP....F..................
WP_202760356.1      ............................................................
WP_252978133.1      ............................................................
WP_065346974.1      PSAAMQ.NAQNAKAA..................QATQ....A..........AAPPATAA
WP_230355409.1      PSAAMQ.NAQA...A..................QATQ....T..........AAPAATAA
WP_267179154.1      ASPGAA.VPAPASAA..FAA..............AVT....P..........SAPSAPSA
WP_088164707.1      PSAAMQ.NAQATQAA..QET.............QAAA....P..........AAPPATAA
WP_133073230.1      ASPGAA.VPAPASAA..FAA..............AVT....P..........SAPSAPSA
MCC7098932.1        .............DG..F.................DA....L..................
WP_238004152.1      PSAAMQ.NAQATQAA..Q..................................AAPPATAA
MBV8620386.1        PAAAEP.DLVQRPP.....................SP....V..........VAPAVSPA
ODU08341.1          ALAAAE.ASAAGERL..A.................DP....L..................
MBC7710584.1        PPQ......PPEPPA..VRR...DA..LPTVLPQAARTD..AIATTPA....ANIDLLAA
MBL8310054.1        PVTTV....VHRPAAQHSTA...PARSAPNPAVPAFRPPSPV.DVPPA............
WP_238363320.1      ............................................................
MCX8099153.1        RAALS....ATTIPV.............................................
MCM2252316.1        PQEVL....EPTQN..............................................
RYF42966.1          PAPAQ....GAASPL............HPPVAAPVSAPHAP..WAPPPGAGVANLDSLFA
WP_281607349.1      PAAPHA.GLA.........................AR....SPAAPPAGSPVLSPVLSPV
MDB5976135.1        ............................................................
WP_201212573.1      P...........................................................
MBP8267915.1        ............................................................
MCZ2135183.1        TPP......VAIPPVEASRV...EA..RPNVAPPTIAPAPPSLIAPPLGATTGEGSFAAQ
WP_131145415.1      PAAPHA.GLA.........................PK....SPAVLPAAPPAGSPVLSPV
WP_166302793.1      APLRETPAQASNPLLQGFTPAAA........IAQASTPRTPIPMDPLATFEPMQAAAIPV
WP_267727965.1      ............................................................
WP_299903544.1      ............................................................
WP_065341524.1      ............................................................
MBL8494843.1        RQVHPS.QEVTISPE.AFA.....AARR....AAAQR....PATPPSLGAGMAQPVPAAR
WP_058641008.1      ............................................................
WP_183632487.1      ............................................................
RYY83951.1          ............................................................
MCU0967987.1        ............................................................
WP_106557035.1      ............................................................
MBK7663138.1        ..VPPA.QEVTIAPE.AFR.....APPR....PMAVP....SPMAASPQPEIQRPASMPQ
WP_203203692.1      ASAIAP.APAAVEPTVAWPPAVESASSRLAAAPVSSR....LASTPDFGAQA..ASLPTS
WP_295989070.1      ............................................................
MBT9459293.1        ............................................................
```

*ABM32059.1*

```
                                                                              TT
                      150        160           170        180        190        200
ABM32059.1          PVPVSVPVAIPVANVPQGE..GLTVVPSPPVAMPAAGTGEGLAAAAASAEEDDNPFA.ML
NP_248771.1         ...............................................SGNPQTDPFT.SF
WP_001089704.1      ................................................ASKDEDLLADLT
WP_200016399.1      PQPQIVPP..PVAAQPP................................LPEATPLA...
WP_063326054.1      PQPQIVPP..PVAAQPP................................LPEPAPLA...
WP_012205666.1      PQPQIVPP..PVAVQPP................................LQE........
MBK9235436.1        ................................................REDTPRPLE.L.
MCA0243819.1        ...........................................PAAAAAALDENPFA.FI
MCZ2441668.1        ...........................................GPEAGIAADDNPFA.FI
WP_202760356.1      ............................................................
WP_252978133.1      ............................................................
WP_065346974.1      PQPQIVPP..PVAVQPP................................LPEATPLA...
WP_230355409.1      PQPQIAPP..PVAAQPP................................LPEAAPLA.EA
WP_267179154.1      PSTPSVPS..PVTVIPQ................................PPIQQPLA.VD
WP_088164707.1      PQPQIVPP..PVAAQPP................................LPEAAPPA...
WP_133073230.1      PSTPSVPS..PVTAIPQ................................PPIQQPLA.VD
MCC7098932.1        ...........................................AAPGAMAADDNPFA.FI
WP_238004152.1      PQPQIVPP..PVAAQPP................................LPEAAPPA...
MBV8620386.1        VSPSVSPP..ALAAADP...............................GTDPANPFA.FL
ODU08341.1          ...........................................AAPIAEAADDNPFA.FI
MBC7710584.1        .........................DPL.............AH...WPEASVGAAYVSLL
MBL8310054.1        .........................VPL.............D.............LDALF
WP_238363320.1      ............................................................
MCX8099153.1        .........................NPF.............EAGG...............
MCM2252316.1        ........................L...................A.AVAPAGADPFAGLS
RYF42966.1          PA..........GAPAAAAFQDLGIAPI.............APGS.SAQRTGSDPFGDLL
WP_281607349.1      MSPSL.PPNLPAGPPSAMP......AMR.............DTEPRLAGVPKDDPLG.LF
MDB5976135.1        MA..............EQT......QTP.............LRAPPTAMAPRDDPLE.IF
WP_201212573.1      .........................A................AEGAGPSAAKDDPLG.LF
MBP8267915.1        ...........................................RPA.EGGGGADPLG.LF
MCZ2135183.1        .........................NPH.............RQRTQAPTTPVDDPLRALL
WP_131145415.1      MSPSL.PPNLPAGPPSAMP......AMR.............VMEPRPAGVPKDDPLG.LF
WP_166302793.1      PETPLPEPLIPAPSIAPAI....AATPT.............YTDLPSPASVMKGPFDDLL
WP_267727965.1      ..........................................TAPA.SAASGNDPLG.LF
WP_299903544.1      ...........................................TPT.VVASGADPLG.LF
WP_065341524.1      ..........................................NETLPNIEQRGDPLG.LD
MBL8494843.1        PVPV....QAPQPPMAPAP......LPP.............QAAPSLAAGPVDDPLG.LF
WP_058641008.1      ..........................................AA............T.AG
WP_183632487.1      ..........................................AAPT.TPTVASDPLG.LF
RYY83951.1          ..........................................GA.........DV....F
MCU0967987.1        ............................................................
WP_106557035.1      ..........................................AA.........APAP.AH
MBK7663138.1        PAP.....RAPEAPRAPPA......PAP.............AVASSLPAGPVDDPLG.MF
WP_203203692.1      M.....PPAYP...PAPPP......AAA.............PLDALLPQAPVSDVFSDLF
WP_295989070.1      ............................................................
MBT9459293.1        ............................................PD...DPFADLLG.PA
```

*ABM32059.1*

```
                       210         220       230                   240
ABM32059.1      GRV..EGPAIAASAP..SRAPAPAALPTASTAP............IPAFGPVPADAP..V
NP_248771.1     DAL..MSRQAAG................................................
WP_001089704.1  LDI...EPEKH.................................................
WP_200016399.1  .............................EAAP............L........ADV..L
WP_063326054.1  .............................EAAP............L........ADV..L
WP_012205666.1  ..............................ATP............L........ADI..L
MBK9235436.1    .AN..SGP....................................................
MCA0243819.1    GRV..EASPSPPA.......AA........PSP...........................
MCZ2441668.1    GRV..EAEMPARA.......GA........ARA...........................
WP_202760356.1  ....................PL......AEAAP............L........ADV..L
WP_252978133.1  ............................................................
WP_065346974.1  .............................EAAP............L........ADV..L
WP_230355409.1  A.......PLAEA......APL......AEAAP............L........ADV..L
WP_267179154.1  S......PLQPQA......TGP......AEAAP............L........ADV..L
WP_088164707.1  .............................EAAP............L........ADV..L
WP_133073230.1  S......PLQPQA......TGP......AEAAP............L........ADV..L
MCC7098932.1    GRV..EAGAPARV.......PA........AA............................
WP_238004152.1  .............................EAAP............L........ADV..L
MBV8620386.1    G......PALDSRTP..RPVPV......SQTAP............V........D.A..L
ODU08341.1      GRI..ETDAPARS.......AG........ASP............V..............
MBC7710584.1    A.....TSEPVAN...QPPDQFSAGLSAFDFAVG..SNAGTASDPLASLGMPPLA.QPPI
MBL8310054.1    APT..ASSKGGVG..GTLGELFPPSG....VASSAASLLAP.AEPLA.VETPPVASSVAA
WP_238363320.1  ..................................................MIPAA.P...
MCX8099153.1    ..G..VASAPPAA....LPTPAPPA...............................APIL
MCM2252316.1    APD.....AP.......APG..P......SAVPMPGAPLHPVDDPLAGLLGSAAS.TPR.
RYF42966.1      AGL..SPAAPPPAPAAPMPAPPPPHVPVAISAPAPVAAALPVDDPLAGLLGPGTA.LSSP
WP_281607349.1  DSA..AAPSSGG................................................
MDB5976135.1    GAA..PAP....................................................
WP_201212573.1  GGL..AQGG...................................................
MBP8267915.1    GGA..SASADPFADLLGTP.........................................
MCZ2135183.1    GDA..AANVPAAA..GAPDALFAADLPVVSPLLSAVDSVSPGGDVAGNFGVIPPA.P...
WP_131145415.1  DSP..ASPPSGG................................................
WP_166302793.1  GAP..VAPDAAGE.....PLTYPS..........QTAHEHLPDDPFA.MASLPTPQPVRA
WP_267727965.1  GGA..PSTDPFGAPASFAPAAFAPAAPPAAAKP......................TVPPS
WP_299903544.1  GGA..PSASPLFADPFAAPA....................................GTPS
WP_065341524.1  LDL..AL...............PPDLPAGLGAP.....................AAAAPP
MBL8494843.1    GGA..PSGT...................................................
WP_058641008.1  P.......................................................PAPA
WP_183632487.1  GSA..PSSDPFGAPAMSAAAT.PPIMPPRAAAP......................M....
RYY83951.1      A.......................................................PARS
MCU0967987.1    ............................................................
WP_106557035.1  G.......................................................HVPA
MBK7663138.1    GGA..PRTG...................................................
WP_203203692.1  GAGALPIGDTHAAPPREAPLQ..................................QPVPQ
WP_295989070.1  ............................................................
MBT9459293.1    AGA..RGPAAAQAPAAR...........................................
```

*ABM32059.1*

```
                   250       260                                     270
ABM32059.1      PPSRSASMPAPDGVHGFAASPASA..............................IPVPSD
NP_248771.1     ............................................................
WP_001089704.1  ..ELPTNTDPLQALDNLMGAAGSSTS..................................
WP_200016399.1  PAAAPGIADPWEGLTALSLDAIQT..............................PAVPSI
WP_063326054.1  PAAAPGIADPWEGLTALSLDAIQT..............................PAVPSI
WP_012205666.1  PAAAPGIADPWEGLTALSLDAIQT..............................PAVPSI
MBK9235436.1    ..........................................................SK
MCA0243819.1    ............................................................
MCZ2441668.1    ............................................................
WP_202760356.1  PAAAPGIADPWEGLTALSLDAIQT..............................PAVPSI
WP_252978133.1  ............................................................
WP_065346974.1  PAAAPGIADPWEGLTALSLDAIQT..............................PAVPSI
WP_230355409.1  PAAAPGIADPWEGLTALSLDAIQT..............................PAVPSI
WP_267179154.1  PPAAAGIADPWEGLTALSLDDIQT..............................PPAPSA
WP_088164707.1  PAAAPGIADPWEGLTALSLDAIQT..............................PAVPSI
WP_133073230.1  PPAAAGIADPWEGLTALSLDDIQT..............................PPAPSA
MCC7098932.1    ............................................................
WP_238004152.1  PAAGPGIADPWEGLTALSLDAIQT..............................PAVPSI
MBV8620386.1    PPV....AHPM.................................................
ODU08341.1      ............................................................
MBC7710584.1    ASAIPVDVDPFADLLAGIGAAPGAPPSNGFAVN................PSSRPDAAQAG
MBL8310054.1    PSASPSGGDPFADLLSGIGSAAPTSA..APPAG................GRPPVLGLPAA
WP_238363320.1  EPGPSAGGDPFANLLVGIGSPSA....SSQAM...................PAIGHVPAG
MCX8099153.1    PAAPDLADDPFAIYLEGIGTPTPPSVPLAPSTAPFVEAPTPTPTAATPSVLSPPASHIGG
MCM2252316.1    ..APAAGADPLAAFGGA.........................................PA
RYF42966.1      AFPPASASDPFAGLGIGGAGAVSAPDPFAGAAA................SASAPEAIAPG
WP_281607349.1  ..........F...........................SPAPPS................
MDB5976135.1    ............................................................
WP_201212573.1  .....PDEDPFASILAGAVGAAVPPM...........A.TEAPA.......RSPVPAP..
MBP8267915.1    APAASGGDDPFAVFAAP.....................LK.PVA.......GPSP...E.
MCZ2135183.1    S...PASVDPFAGFLEGIGTPTP....QGGAFT................GRNAGGATPVG
WP_131145415.1  ..........F...........................SPAPPS................
WP_166302793.1  PVHSSSVGDPFGDLMGA...........................................
WP_267727965.1  PSAFDNADDPFAVFAVA.....................SPPPTP.......APSH...A.
WP_299903544.1  TPPPAAGDDPFAVFLAS.....................AR.PAP.......AVAP.....
WP_065341524.1  PRAAALDDDPFAVFAAA.....................TPTPPA.......PPAAAPPA.
MBL8494843.1    .......ADPFADILNGVTASPSPQP...........QVRPAPA.......ASAPPQPAA
WP_058641008.1  PQPVAAGDDPFADLLAG.....................LGPGPA.......APAAASPAR
WP_183632487.1  ...PAPVDDPFAVFAAA.....................VPTPP.........AAT...N.
RYY83951.1      ATPAPADDDPFADLLAG.....................LATP...........SASAPA.
MCU0967987.1    ........................................................APA.
WP_106557035.1  QAALPTDDDPFADLLAG.....................LGPAAP.......AAAPAA...
MBK7663138.1    .......ADPFADILSGVTAASPLPP..............PEPA.......ASP......
WP_203203692.1  PVPPP.AQRPPAPVAAPVFAPASPAFAP..PPAAFASPVPPPPA.......APAP.APAA
WP_295989070.1  ................G.....................GAPPAA.......PAAPSRPA.
MBT9459293.1    ...TPGLVDPLAAFGFEPTPQTSPGI...........YSRPPVA.......APA.ASPS.
```

```
ABM32059.1                                                                     TT   TT
                      280         290                             300              310
                        .           .                               .                .
ABM32059.1         Q..GRQDNAWSPAVPQD...............RPAAAPAPRALVIP........EDFDLF
NP_248771.1        ...........................................SA............PAF
WP_001089704.1     ............LL...............DDPKSEQPLSAQPSLVPQDNLVLQNADFTPQ
WP_200016399.1     E..S.......A.IP.P.......EQAALAMPQP.AKPQHRPLLIP........NDFDPF
WP_063326054.1     E..S.......A.IP.P.......EQAALAMPQP.AKPQHRPLLIP........NDFDPF
WP_012205666.1     E..S.......A.IP.P.......EQAALAMPQP.AKPQHRPLLIP........NDFDPF
MBK9235436.1       L..AATPNPWADIEPQV.......REVYSLLPKSAPVDAHRPLLIP........ADFDPF
MCA0243819.1       ........................R......AAPAPA.PPGALLIP........DDFDPF
MCZ2441668.1       ................P...........APGTPPSPSRAANAPMIP........DDFDPF
WP_202760356.1     E..S.......A.IP.P.......EQAALAMPQH.AKPQHRPLLIP........NDFDPF
WP_252978133.1     ............................MAMPQP.ARPQHRPLLIP........NDFDPF
WP_065346974.1     E..S.......A.IP.P.......EQAALAMPQP.AKPQHRPLLIP........NDFDPF
WP_230355409.1     E..S.......A.IP.P.......EQAALAMPQP.AKPQHRPLLIP........NDFDPF
WP_267179154.1     Q..S.......A.IP.P.......EQAAMAMPQP.ARPQHRPLLIP........NDFDPF
WP_088164707.1     E..S.......A.IP.P.......EQAALAMPQP.AKPQHRPLLIP........NDFDPF
WP_133073230.1     Q..S.......A.IP.P.......EQAAMAMPQP.ARPQHRPLLIP........NDFDPF
MCC7098932.1       .............................RASAPASAPAGHGSLIP........EDFDPF
WP_238004152.1     E..S.......A.IP.P.......EQAALAMPQP.AKPQHRPLLIP........NDFDPF
MBV8620386.1       ..............S.A........AGLAGSPRPEGPTPPRPLIIP........ADFDTF
ODU08341.1         ................A.......CA..GTAAQPSGSGAASAPTIP........DDFDPF
MBC7710584.1       LSADPMAAFGGSMSFTQGEPLS.E....RSPLYGSGTAGAANGLLP........DDFNAF
MBL8310054.1       TSADPFASLLMP...GGGTPIS.APVAFAATPVA..PPPPTPSVIP........DDFNPF
WP_238363320.1     LVADPLAALGGSLSTTGYS..........SPPVASSGANISGALIP........DDFNPF
MCX8099153.1       VPLDPLAALGGALPEARPSP.......S.......WRDGLAEGDAR........SAENPF
MCM2252316.1       AGADPFAALGQLPRAAL.DPLA.APAPSADLGGAPARTPPPALLIP........EDFNPF
RYF42966.1         SQLDPFAALGLGPGAG........APNALATGMMAPSPPPPAVVIP........EDFNPF
WP_281607349.1     .................................GSFAHAPGAGGIP........DDFSF.
MDB5976135.1       .......................................AGQGGIP........ADYDPF
WP_201212573.1     .............AERPAS.........AP...SVAGASASAPVIP........EDFDPF
MBP8267915.1       .........................................APALR........DPFAAA
MCZ2135183.1       LSVDPLAALGSSPGNFSG...........APPIAPV..PPTNTGMA........EDFNAF
WP_131145415.1     .................................GGFAHAPGAGGIP........DDFSF.
WP_166302793.1     ............PIDTHMS.........SAPAHSIGHASAMPRVIP........DDFNPF
WP_267727965.1     .........................................APPPP........DPFAMP
WP_299903544.1     .........................................AEAAH........DPFAPP
WP_065341524.1     .........................................ASVSD........DPFAAF
MBL8494843.1       A..DPLAGVWSPPARTPTESESLSELLAAPLPAGPAMPNNRAPLIP........DDFDPF
WP_058641008.1     R..............SDAAAAAPALPAAFADPLGAVPGPAATAPAD........DPFAGL
WP_183632487.1     .........................................TAPLP........DPFAAP
RYY83951.1         ................ARAPAKPADPLLFPNPMETQPR.NARAPEV........DPFADL
MCU0967987.1       .........................................APVIP........DDWNPF
WP_106557035.1     ...................MQAPAPAGLFPDPLGAGR..APSASAD........DPFADL
MBK7663138.1       V..DDPLAGWTAPPSTPSPEQSLADLLAQPSASSQSMPSARPGMIP........DDFDPF
WP_203203692.1     RSSMPTAADWDAILAN............AP.....RRADTTPEPMP........DPFAHE
WP_295989070.1     ................AAAAAALPDSLLFPDPMESGSR.NVQAAQV........DPFANL
MBT9459293.1       .........................................LGGIP........DDWDPF
```

```
                          η1                                                     α1
ABM32059.1               ℓℓℓ          TT                                        ℓℓℓℓ
                                320                                     330      340
                                  .                                       .        .
ABM32059.1         AVDPRKGEEK.KD..AWGSG.......L....................QAKSLSEIANMR
NP_248771.1        AEPAPTP.HP.........................................AVTAHFQGG
WP_001089704.1     ADSEY...E.....................................MTSSIRLKKILGFA
WP_200016399.1     APPPQQARQQ.EE..GWAGL.......PS.................QGT....GELIQQR
WP_063326054.1     APPPQQARQQ.EE..GWAGL.......PS.................QGT....GDLIQQR
WP_012205666.1     APPPQQARQQ.EE..GWAGL.......PS.................QGT....GELIQQR
MBK9235436.1       ARDLKRDAQA.QD..PWAGG.......L....................QATSLAEVANLK
MCA0243819.1       ARDIEQERRD.KD..PWAGG.......L....................PMHNLAEMAPL.
MCZ2441668.1       AQDAERERRE.KD..PWADG.......L....................PMQNLAEMAAV.
WP_202760356.1     APPPQQARQQ.EE..GWAGL.......PS.................QGT....GDLIQQR
WP_252978133.1     APPPEQARQQ.EE..GWASL.......PS.................QGK....GELIQQR
WP_065346974.1     APPPQQARQQ.EE..GWAGL.......PS.................QGT....GELIQQR
WP_230355409.1     APPPQQARQQ.EE..GWAGL.......PS.................QGT....GELIQQR
WP_267179154.1     APPPEQARQQ.EE..GWAGL.......PS.................HGK....GELIQLR
WP_088164707.1     APPPQQARQQ.EE..GWAGL.......PS.................QGT....GDLIQQR
WP_133073230.1     APPPEQARQQ.EE..GWAGL.......PS.................HGK....GELIQLR
MCC7098932.1       ARDIERERRE.KD..PWAEG.......L....................PMQNLAEMAAV.
WP_238004152.1     APPPQQARQQ.EE..GWAGL.......PS.................QGT....GDLIQQR
MBV8620386.1       APRPATAASAGRA..PWGEA.......AR.................DGLSGSLVEIAGVR
ODU08341.1         ARDAERERRE.KD..PWAEG.......L....................PMQSLAEMAAV.
MBC7710584.1       HMPSSSSRNA.AD..PLAGL.......MGSGAI...A.DTSRVGAVSDGTPSIDSLFQTA
MBL8310054.1       ELPTVTSRNT.SD..PLAGL.......TGTAPI...GLSDTGTGGGLHAAPSIDSLFATP
WP_238363320.1     AMPSETSRNA.AD..PLSGL.......LGTAPG...QPDGAAAASGPAVAPPIDSLFQNG
MCX8099153.1       ALPSAAVRNT.AD..PLSLF.......APPPS....PAENA..LAGGAVAPPVESLFPPV
MCM2252316.1       DLPSASARNS.AD..PLAAM.......SGGSGA...GAPHA....LVQAEESIDALFAPG
RYF42966.1         ELPSAAARNT.AD..PLSSL.......MDPGGA...PTGEA....LPGNEQSIDVLFGAT
WP_281607349.1     SELVP..RAP.AP...DP..RQPLPDDFDLGLG.............QAPRTDINQFYDLG
MDB5976135.1       ADMFAPETRK.AP...HA..GAAPPAELRLGTG.............DTSGQSLDQLFGLG
WP_201212573.1     ADPFAQAPAA.KL.......ASDNLDDLGFG...........ASGESVSGPGISTIFGLE
MBP8267915.1       S.P...PPAA...P.........RADLLGLG...........G...VRGEQSVDALFGLG
MCZ2135183.1       ALPSDAQRNT.AD..PLAGL.......LGQAPG...GADHVA...VSATAPPIDSLFRVL
WP_131145415.1     SELVP..RAP.AP...DP..RQPLPDDFDLGLG.............QAARTDINQFYDLG
WP_166302793.1     ALGGVSRRNM.DD..PLSDL.......LRPGN...........VKDMFPERSLDAIFQPT
WP_267727965.1     A.P...APRP...A.........APLGMDLN...............LAPAASVDDLFGLG
WP_299903544.1     P.P...APPP...A.........RVDVLGLG...........A....GCESSVDALFGLD
WP_065341524.1     A.PARPAPPP.APAA......SRAGDPLGIG...........ALAQPAEASSLDALFGLD
MBL8494843.1       ADPFAVPPKT.AE.......PEALDDDLGFG...........LAGKSGSGSSIDTAYGLA
WP_058641008.1     LGPAPAAP............AL.PDDFSDLGLP........AASHPTANAQRIDDLFGMG
WP_183632487.1     P.P...APKP...S.........APLGMDLN...............LAPAASVDDLFGLG
RYY83951.1         LGPAPAS.......APAAGFGS.LDDFSDLGVA........KGGGK...SNGIDDLFGGM
MCU0967987.1       A.PEPQSSAR.PV......MGPAAGL..............GLPIAAPPRAESIDDLFGLG
WP_106557035.1     LAPSAPASGG.A.AAPARG.LP.LDDFSDLGLS........A.SPK...SPGIDELFG..
MBK7663138.1       ADPLAAPPKD.EE.......VNP.FDDMEVG...........LGEATGKGANFDSMYELG
WP_203203692.1     ..PFELPS......................................QARRNPADPLAQLN
WP_295989070.1     LGPGPSS.......KPAAGLGA.LDDFSDLGAP........PANGK...AASIDELFGGM
MBT9459293.1       A.PDPVAARP.AAQDLARSLGSGGNFGLDAGAAAPSALIPDLPASGPGNDNSLDNLFGLK
```

```
                                  α2                      η2                                η3
ABM32059.1          QQ.............QQQQ              QQ....Q           TT          TT     QQQQ
                                           350                   360         370          380
ABM32059.1          HD.............ELLQSLPASGA....KFAHDMDNPAHAGLPKALDPRDELDPLRL
NP_248771.1         ..SPL.......DTKPDLF.......................................D
WP_001089704.1      KA..............T.........................................
WP_200016399.1      HD.............GMVHTLPLKG.....LSTEALHDASHTGLPSCFEQSQKLDPLAL
WP_063326054.1      HD.............GMVHTLPLKG.....LSTEALHDASHTGLPSCFEQSQKLDPLAL
WP_012205666.1      HD.............GMVHTLPLKG.....LSTEALHDASHTGLPSCFEQSQKLDPLAL
MBK9235436.1        SD.............GLLQSLALTG.....HFESAIDNPAHPGLPKRLEPTNVVDPMVL
MCA0243819.1        RD.............ELLRSLPAIDR...VDQPSPLERSAVKGLPASLDPQMELDPLRL
MCZ2441668.1        HD.............DLLRTLPPADR...LAQASSLGQSAVRGLPASLDPNGELDPLRL
WP_202760356.1      HD.............GMVHTLPLKG.....LSTEALHDASHTGLPSCFEQSQKLDPLAL
WP_252978133.1      HD.............GMVQTLPLKG.....LSTEALHDASHTGLPSCFEASQKLDPLAL
WP_065346974.1      HD.............GMVHTLPLKG.....LSTEALHDASHTGLPSCFEQSQKLDPLAL
WP_230355409.1      HD.............GMVHTLPLKG.....LSTEALHDASHTGLPSCFEQSQKLDPLAL
WP_267179154.1      HD.............GMVQTLPLKG.....LSTEALHDASHTGLPSCFEASQKLDPLAL
WP_088164707.1      HD.............GMVHTLPLKG.....LSTEALHDASHTGLPSCFEQSQKLDPLAL
WP_133073230.1      HD.............GMVQTLPLKG.....LSTEALHDASHTGLPSCFEASQKLDPLAL
MCC7098932.1        HD.............DLLRTLPPIGR...LAQPSVLDAEAVRGLPASLDPHGERDPLRL
WP_238004152.1      HD.............GMVHTLPLKG.....LSTEALHDASHTGLPSCFEQSQKLDPLAL
MBV8620386.1        ED.............GLLQALPRLQ.....ELAQRLDNPAHSGLPASLERSAELDPLRL
ODU08341.1          HD.............DLLRTLPPVER...LGQGSALHEAAVRGLPASLDPRGELDPLRL
MBC7710584.1        GHDSSY..........ASL.LGLAPAARG....GDGSIEGLIRS...D....TANDPMSL
MBL8310054.1        APAA...........VDALLAPSAAGHG....QADGWGGLLQA...D....ANSDPLAL
WP_238363320.1      SGFDRV..........GDSLHAPLSQARD....GNSDLSALLGP...AD...GNSDPLAL
MCX8099153.1        APPLAD..........PVLEAFARPGERATPGLGDGSLDSLLRP...N....EPTDPLEV
MCM2252316.1        GAASGD..........PFLAAV.GAGAGVPGSASGAASSLLAQ...P....DNGDPLAL
RYF42966.1          GASHAL..........TIKGAP.G..SDFLG..GPGGGSLLEP...L....ENSNPMAM
WP_281607349.1      PAA...........GSELFP...........DGHPLAPGGANG...SGAA..SVDPLVA
MDB5976135.1        GGA...........APELFP...........NGNPLSDPGQRP...DARD..ERDPLVA
WP_201212573.1      RGARA.......GTAADLL.............AGTPLAAPAPGK.D.EKGA..LLDPLAA
MBP8267915.1        GG..........A..ADPF.............AGTPLGDAGVVP.S.NAGS..VVDPLAL
MCZ2135183.1        PGEHA..........DPFLATPVTP.SS....QGGDLGGLLDP...RDG..GSTDPLAL
WP_131145415.1      PAA...........GSELFP...........DGHPLAPGGANG...SGAA..SVDPLVA
WP_166302793.1      EGSID.......SMTVDPLQ.......................AEHHQS...FMDASTHVDPLAL
WP_267727965.1      TA..........SSAADPF.............AGTPLSSHTAPP.A.AQPN..AADPLAL
WP_299903544.1      GG..........A.APDPF.............FGTPLGEPAPSAAP.AAPE..VVDPLAL
WP_065341524.1      GA..........STGGDPF.............AGSPLGNPAHAPAP.FDGG..GEDPLAL
MBL8494843.1        APPPV.......GNGIDPF.............LGSSLSDPLAGR.G.GD.A..PIDPLAA
WP_058641008.1      A...G.......LPGSDPL.............ALSPLADPLLQP.N.TAGS..AADPLAA
WP_183632487.1      GG..........KGAADPF.............AGTALSSHTDAP.A.SAPA..AIDPLAL
RYY83951.1          GGL.........GMGSDPL.............ALSPLADPLLQP.N.TA.S..HADPLAA
MCU0967987.1        PM..........TPGTDPL.............GAAPAQALRLQP...GTAG..DADPLRA
WP_106557035.1      ..IGG.......SGGGDPL.............ASSPLGDPLRQP.N.TA.S..AADPLAA
MBK7663138.1        PSMPA.......DTRDPF..............VGSSLGESPVEG.P.VNET..PLDPLAA
WP_203203692.1      PGATTDVSDFALKRGIDPLSLFAAD..RD...APSPLSDPRPTALT.HAPALEDVHPAMD
WP_295989070.1      GGSGG.......GIGGDPL.............ALSPLADPLLQP.N.TA.S..NADPLAA
MBT9459293.1        PG..........MGG.DPL.............ANSALDAPMAKP...NMAA..SHDPLAS
```

```
ABM32059.1
                            390                     400              410          420              430
ABM32059.1          FTGGSDAASALV.......PERSAMVLGSGSDLSQSFSLPRGVQAPVG..PGLPGAA...
NP_248771.1         FL..TP.P...........PPGAAPRPDHVPAEQHDFRPPEPVIPPPPATTPA..PPPAG
WP_001089704.1      FAKQTKSEAPNV..T..SATPTISQYKTSESNVSEGFTMDEK..................
WP_200016399.1      FDQGGLPPDAQP.......L...QQASGRGSELGQAFNLPRMQSHAQPAEPVPPQEAAAP
WP_063326054.1      FDQGGLPPDAQP.......L...QQASGRGSELGQAFNLPRMQSHAQPAEPVPPQEAAAP
WP_012205666.1      FDQGGLPPDAQP.......L...QQASGRGSELGQAFNLPRMQSHAQPAEPVPPQEAAAP
MBK9235436.1        FRDDHEDVTPAM.......AP..CPAPTRGNDLVQLFSMPRESGSAPA..PA........
MCA0243819.1        FESAPTLPGLPP.......D...AARLLHGSELGQALRLPREAAMPAAEAPVIAPVI...
MCZ2441668.1        FDDAPSAPGPAF.......A...SAHLLRGSGMHQAVRLPQAMDAAAVP..AAAPV....
WP_202760356.1      FDQGGLPPDAQP.......L...QQASGRGSELGQAFNLPRMQSHAQPAEPVPPQEAAAP
WP_252978133.1      FDQGGLPGDAAP.......L...QQASGRGSELGQAFNLPRMQPPAQPAAQG....AAAP
WP_065346974.1      FDQGGLPPDAQP.......L...QQASGRGSELGQAFNLPRMQSHAQPAEPVPPPEAAAP
WP_230355409.1      FDQGGLPPDAQP.......L...QQASGRGSELGQAFNLPRMQSHAQPAGPVPPPEAAAP
WP_267179154.1      FDQGGLPGDAAP.......L...QQASGRGSELGQAFNLPRMQPPAQPAAQG....AAAP
WP_088164707.1      FDQGGLPPDAQP.......L...QQASGRGSELGQAFNLPRMQSHAQPAEPVPPQEAAAP
WP_133073230.1      FDQGGLPGDAAP.......L...QQASGRGSELGQAFNLPRMQPPAQPAAQG....AAAP
MCC7098932.1        FGDEPTQPGLAP.......E...SARLLRGSEMHQALRLPQLASADPLA..ALASA....
WP_238004152.1      FDQGGLPPDAQP.......L...QQASGRGSELGQAFNLPRMQSHAQPAEPVPPQEAAAP
MBV8620386.1        FELQTEAPEAAV.......G...TTPGIGGDRLGQVFTLPRRADAAG...AEPPKEVARP
ODU08341.1          FDDVPTQPGLAP.......E...SARLLRGSEMHQAMRLPRAAPTEPMP..TEALS....
MBC7710584.1        FDIAESAP...........RQSAQPMRNDSMEVGSAFAPPLARLAERQSHAPLHPNP...
MBL8310054.1        FDAPAPAP.............ALQPMRDTTPEIGSAFVPPVAAPLARPAVSA..PTP...
WP_238363320.1      FDSPAAAP.............THRPMRDDSLELGAAFAPPLPLHADPLAAFQ.QAAP...
MCX8099153.1        LQGQSAQT...........PLAPVADHLPEFRAAFEPPRAVEASADLSMARRSET...
MCM2252316.1        FGDA.ARP...........ESLGVPMRDDLAEIGGAYQPPRPLQPQPASPPAPQA.....
RYF42966.1          FGDA.PAP...........DGMSRAMRDDLAEVGGAYQPPRALDPGGPFGMPGQS.....
WP_281607349.1      IGAVPA.P...........KPAPASQRDDAPEIRSAFTPPAAQHAEPSHP.........
MDB5976135.1        LGGAPQ.P...........RY.PDTARDDTPQIRAAFIPPTATFGAREQARPDAAAAPSC
WP_201212573.1      LSGAAP.A...........EPKQAAVPDQVPEIRAAYRPPAAK................
MBP8267915.1        FGGAAP.A...........AATPEPQRDDAPVLNQAFAPPRPEPA.EPEVTHV..LA...
MCZ2135183.1        LGSAEPAP.............ALRPMRDDGKEVGAAFAPPLAQSSASGA.TG.ASAA...
WP_131145415.1      IGAVPA.P...........KPAPASQRDDAPEIRSAFTPPAAQRIEPVQP.........
WP_166302793.1      FSEDRKEDALNPEMLFGNTAIREKTQSDHTSELGSYFRAPRALQDPAPPGTIDPLQAFSQ
WP_267727965.1      FGGTPA.P...........APAAAPGRDDAPMLGESFTPPRPIGPAPTLA.....PAPNP
WP_299903544.1      FGGPAA.P...........APGAAPQRDDAPLLNQAFTPPRAEIPAGAELTQV..VAPRP
WP_065341524.1      LGGMAA.............TPSSAPVRNDSPLLNDAFAPPRLIDDSPPPP.LA..PAPAS
MBL8494843.1        FSGDLGAP...........IPGFQSMPDQVPELHGAFVPPRVEPFAAPEPGAFAADSFAP
WP_058641008.1      FAQPAP.A...........SQA..PRSDHVPIGRFGFTPPREAEGGGAGGDAAA.EPIRI
WP_183632487.1      FGAAPA.V...........APSAAPLRDDAPMLGEAFTPPRPLAPQSAEP.....PAA.M
RYY83951.1          FERAAP.V...........TPV..PRGDGLPMGQFAYQPPREVQAPTP............
MCU0967987.1        LAQGPR.P...........APP.ATIGDEGSELHTPMPL.................PPAAA
WP_106557035.1      LQGTAA.S...........AAK..PRSDHLPVGQFAYTPPREVAPPPPVA..........
MBK7663138.1        FSET.PLP...........GGGYQSMPDQVPELHGAFVPPRVEPFAAPAPGAFAGPTPAR
WP_203203692.1      LLEKRP.A...........VP.GYTQPNHAHEVSGQFRPPTPMPLPVAQPPAR......M
WP_295989070.1      LQQAAP.A...........APV..PRPDHLPIDQFGFTPPKAVEPPRPPA.....PPPVM
MBT9459293.1        LNSAPK.A...........SAA..SASDAFSDLNRPFVPPPAIGAKPVAP..P..PPPVE
```

*ABM32059.1*

```
                                  440
                                    .
ABM32059.1        .........EVPGSSPPVVQG.....................................AG
NP_248771.1       G..................APLIPADWDPFAELLGNTP.......APSATPVA.......
WP_001089704.1    ............................................................
WP_200016399.1    AQHAL...PPVIDTVIPAVQEQLPVQVEARA.................QA..QD...IPV
WP_063326054.1    AQHAL...PPVIDTVIPAVQEQLPVQVEARA.....QV..........QA..QD...IPV
WP_012205666.1    AQPAL...PPVIDTVIPAVQEQLPAQVEARA.................QA..QD...IPV
MBK9235436.1      ............................................................
MCA0243819.1      ...............APMITPA....................................IP
MCZ2441668.1      ............................................................
WP_202760356.1    AQPAL...PPVIDTVIPAVQEQLPVQVEARA.....QVQ....AQVPTQA..QA...IPV
WP_252978133.1    AQPA.....GLEQSLAPAAQTQQPVQVQAAPA....AVPVAVPAPLPAAAPVAA...PMA
WP_065346974.1    AQPAL...PPVIDTVIPAVQEQLPVQVEARA.....QVPTQVQTQAPAQA..QD...IPV
WP_230355409.1    AQPAL...PPVIDTVIPAVQEQLPVQVEARA.....QVPTQVQTQAPAQA..QD...IPV
WP_267179154.1    AQPA.....GLEQSLAPAAQTQQPVQVQAAPA....AVPVAVPAPLPAAAPVAA...PMA
WP_088164707.1    AQHAL...PPVIDTVIPAVQEQLPVQVEARA.....QVQTQVQT....QA..QD...IPV
WP_133073230.1    AQPA.....GLEQSLAPAAQTQQPVQVQAAPA....AVPVAVPAPLPAAAPVAA...PMA
MCC7098932.1      ....................AA....................................AP
WP_238004152.1    AQPAL...PPVIDTVIPAVQEQLPVQVEARA.....QV..............QD...IPV
MBV8620386.1      LPTQAPTGPTRSQSVEPAPHAD.....TA................SPGERPLYK...LTG
ODU08341.1        ............................................................
MBC7710584.1      .............SMVA...........................................
MBL8310054.1      .............TATA...........................................
WP_238363320.1    .............TPVV...........................................
MCX8099153.1      ............................................................
MCM2252316.1      ............................................................
RYF42966.1        ............................................................
WP_281607349.1    ..............DGPADGGNMVVSWNAPDELPGGDGIKTMIVQSPARQARGRQAETPT
MDB5976135.1      GATPAP.......SISKAAEQEMVFSWDRRDGAAADGPIRAVVVPAQ.............
WP_201212573.1    ..................LPDDAFLSWEQAGAGKAAESAEPAEAPSP..RPRA.......
MBP8267915.1      ...............GGPAPAGMVFSWDDAASAPAPRS.......VAAE...........
MCZ2135183.1      .............SLIP...........................................
WP_131145415.1    ..............AGPADGGNMVVSWNAPDELPGGDGIKTMIVQSPARKARAPETETSS
WP_166302793.1    APSLPS.......NLEPLAPIDQGLSS........PPTAQPAIQQSPVHQS.........
WP_267727965.1    LPPPAP.......TTDAGASSGIVVSWTETPLVQSA.P.......APSDINELF......
WP_299903544.1    AGPRGD.......DAAPAAPGAMVFSWDKAPPAASVPA.......AVAE...........
WP_065341524.1    AAAPAP.......QRSPAAPQGGVVSWASTPAAPAAPA.......APAAPAAPA......
MBL8494843.1      PQPQPS.......TRVDAVGNGQFVSWETQSQSARPMAPPPASRQSAAADPLLSMFDA..
WP_058641008.1    G..RAP.......APRPAAPAAAMPPSAPRAG................AAPVQT......
WP_183632487.1    APPPQP.......APAPEPAPGIVFSWSEPATPSPATP.......SSADTAERV......
RYY83951.1        ...........................APE..............................
MCU0967987.1      VDVPVT..........VIKPAGAVFSWDEAAPVEPH............AAP.RT......
WP_106557035.1    ............................................................
MBK7663138.1      AA..............PPSEKEVFMSWEKELPRQG...GRRSSAPSPAEDPLLAMFDG..
WP_203203692.1    V........................EPEPEREAAPATP........VAAPPTASMQSA..
WP_295989070.1    A..PAP.......APMVP.......AQQPL..............................
MBT9459293.1      LIPPLT.......DIKPALPSGAVLSWDAPSGDGHTVI.......RPAARA.DA......
```

*ABM32059.1*

```
                        450       460       470       480                 490
                          .         .         .         .                   .
ABM32059.1        TNPG...MASAAPPAPPLEGLQRVEGLDLGAFGSGSASADPLGWPP.......V...ES.
NP_248771.1       .......Q......PLPTAEP..............TPLAMPFADPG......IT......
WP_001089704.1    .........................................VLDLL...EEEVA....K.
WP_200016399.1    AAPSPAPVPQIPAEPAALDGLLPTNGLDLDLFKSGLAHVAPIARAA.......VA.AASA
WP_063326054.1    AAPAPAPVPQIPAEPAALDGLLPTNGLDLDLFKSGLAHVAPVARAA.......VA.PTSA
WP_012205666.1    AAPAPAPVPQIPAEPAALDGLLPTNGLDLDLFKSGLAHAAPVARAT.......VA.AASA
MBK9235436.1      ..........PAQQVTEALGLRRTQGLDLSLFGADTNPTAALNFGL.......PA.DAQS
MCA0243819.1      PAIPSSITPPPPPAPLQAAGLHRVDGLDLTMFDRAPGALPAGGGEP.......PV.APEA
MCZ2441668.1      .....AALEGTAQAPLHSAGLQRIEGLDLAMFDRALGD...GGAES.......AV.APEA
WP_202760356.1    AAPAPAPVPQIPAEPAALDGLLPTNGLDLDLFKSGLAHVAPVARAA.......VA.PTSA
WP_252978133.1    T......APEIPAEPVVLDGLLPTNGLDLDIFQSGAGQAAPAIVSA.......AS.AATA
WP_065346974.1    AAPAPAPVPQIPAEPAALDGLLPTNRLDLDLFKSGLAHVAPVARAA.......VA.PTSA
WP_230355409.1    AAPAPAPVPQIPAEPAALDGLLPTNGLDLDLFKSGLAHVAPVARAA.......VA.PTSA
WP_267179154.1    TAPAAATAPEIHAEPVVLDGLLPTNGLDLDIFQSGAGQAAPAIVSA.......AS.AATA
WP_088164707.1    AAPAPAPVPQIPAEPAALDGLLPTNGLDLDLFKSGLAHVAPVARAA.......VA.AASA
WP_133073230.1    TAPAAATAPEIPAEPVVLDGLLPTNGLDLDIFQSGAGQAAPAIVSA.......AS.AATA
MCC7098932.1      AATAPAETPRPAAPPLQAAGLQRMEGLDLAMFDRAPAPAP.MSADE.......VI.APEA
WP_238004152.1    AAPAPAPVPQIPAEPAALDGLLPTNGLDLDLFKSGLAHVAPVARAA.......VA.PTSA
MBV8620386.1      ......TVELVPQE..........................PQARPA.......GA.WPTV
ODU08341.1        .....PPAQEALQQPADAAGLQRLEGLDLAMFDRPASA...GSPEP.......VA.APQA
MBC7710584.1      ......................................ALPINAFAAQ...PVPA..APA
MBL8310054.1      ......................................SRPMAD.............PFA
WP_238363320.1    ......................................VDPFAGLMAS...PVAQ..PAA
MCX8099153.1      ......................................GLPDLG...L...PAPA..ASA
MCM2252316.1      ............................................................
RYF42966.1        ...................................................P........
WP_281607349.1    TQPGPVLA......PEPTARA..AEPAA..T.AQPAATGQPAAGAR....AATSAAPVAH
MDB5976135.1      ........................................RGGDA...........DVA.
WP_201212573.1    ................................DVEIAPS.SGAETAKGS...........
MBP8267915.1      ....................................APSTPL..........ARAEISVE
MCZ2135183.1      ......................................GDPFLDLLGS...PAPQ..AHS
WP_131145415.1    TGPAPALA......PEPPALA..AGPAA..T.A......QPSGGAR....AAVPAQPLAH
WP_166302793.1    ..............ASPAASSNAVNLDA..FFDL...SGAPDNGLMGL.D..........
WP_267727965.1    ...............VPPAS....KG..........A.AAVS..........VRIEQAPA
WP_299903544.1    ....................................APPAPA..........VRVELSAE
WP_065341524.1    ...............APAAPA..APA..........APAAPVAGDD.G..REAPTRIVPA
MBL8494843.1      ..............PSAGA.....SQSD..VLGLGVAPV.PGDDMLAAFDAVPSAAPLAS
WP_058641008.1    ...............PPGAGA..A.G..........AVPDPFADLL.G..DEPFG.....
WP_183632487.1    ...............VPAAG....TG..........ADPGVS..........VRMESTPF
RYY83951.1        .......................................VQFDDMT.G..LPVDA.....
MCU0967987.1      ...............VPGVR....RA..........SP..................EPPA
WP_106557035.1    .......................................PRFDDMT.G..EAIRA.....
MBK7663138.1      ..............ASPAAGA..PAGAD..VLGLGVTPAMPGDDLMAGLGTPARV...E.
WP_203203692.1    ........................................PVEAVLAA............
WP_295989070.1    .......................................AQFDDMT.G..QPIRI.....
MBT9459293.1      ...............APAA.................A...................PMPA
```

*ABM32059.1*

```
                             500                   510          520       530
ABM32059.1          .GPAPAAVDVLPQAT..L..........SGLAAGDAKAVPIE...EVALAHVPAAAGTPA
NP_248771.1         QQPQPQP.....QPQ..PQPQP......................................
WP_001089704.1      ...S.......FQ...............................................
WP_200016399.1      PP.....LDAGVNVY..IPAA.......SSILGQGDAAIPIG...AL.....PS...TAA
WP_063326054.1      .P.....LDAGVDVN..IPSA.......SSILGQGDAAIPIG...AL.....PS...TAA
WP_012205666.1      PP.....LDAGINVN..IPAT.......SSILGQGDAAIPIG...AL.....PS...TAA
MBK9235436.1        AAPVPPAVPTSPA...................PSARRVIELS...ES..ATWPVAVAAPT
MCA0243819.1        ..TIFSNIDANLTSTQLLPMRAPVPLPQVPTLAVVAPGLDLDLDG........PA.....
MCZ2441668.1        ..TVFARFDADLAPTQLQALRPPPETA.......AAAPLDVRLDLDLAFAASPEAAISGA
WP_202760356.1      .P.....LDAGINVN..IPAA.......SSILGQGDTAIPIG...AL.....PS...TAA
WP_252978133.1      PANTSSSSSSSSSVH..FNAA.......SSILGQGAAAIPVG...TL.....PC...TAA
WP_065346974.1      .P.....LDTGVDVN..IPSA.......SSILGQGDAAIPIG...AL.....PS...TAA
WP_230355409.1      .P.....LDTGINVN..IPAA.......SSILGQGDAAIPIG...AL.....PS...TAA
WP_267179154.1      PANTS......SSAH..FNAA.......SSILGQGAAAIPVG...TL.....PC...TAA
WP_088164707.1      PL.....LDTGINVN..IPAA.......SSILGQGDAAIPIG...AL.....PS...TAA
WP_133073230.1      PANTS......SSAH..FNAA.......SSILGQGAAAIPVG...TL.....PC...TAA
MCC7098932.1        ..TVFARFDADLAPTRIQPLRLTPAAQEGG....GAPALDLDLDLAPGLSREAQSPPLPS
WP_238004152.1      P......LDTGINVN..IPAA.......SSILGQGDAAIPIG...AL.....PS...TAA
MBV8620386.1        PPSVPAA....LDL.......................ELPL................AAA
ODU08341.1          ..TVFARFDADLAPTQLQPLRSAATPA.......AAAPLDLSLDLEPALAAAAPA.....
MBC7710584.1        VAPH.................................................P......
MBL8310054.1        L...........................................................
WP_238363320.1      VSVV.................................................P..GSNT
MCX8099153.1        SSPL.................................................IFEGRAA
MCM2252316.1        ............................................................
RYF42966.1          ............................................................
WP_281607349.1      PPPVPSP.....QPN..VQPQADPHPH...............................GA
MDB5976135.1        ............................................................
WP_201212573.1      ....................................ERLPGA....EA............
MBP8267915.1        ............................................................
MCZ2135183.1        LPPA.................................................P......
WP_131145415.1      ......P.....QPI..VHPRADPHPH...............................AA
WP_166302793.1      ..........................................................ST
WP_267727965.1      IPPV....................................VA....PT.....I.....T
WP_299903544.1      ............................................................
WP_065341524.1      PPPP....................................EA....PR.....ANVGIER
MBL8494843.1        LPPMAP......A....MPPQAPLPPAAAPLVDFNAPTLPGD....DT.....FFAGIAQ
WP_058641008.1      .PAV....................................A.....SG.....GF....V
WP_183632487.1      ATPV....................................AA....PA.....Q.....Q
RYY83951.1          .PAP....................................PP....IA.....P......
MCU0967987.1        RPAV....................................A...................
WP_106557035.1      .APV....................................AP....VA.....Q......
MBK7663138.1        APPP..................PAPPADKP.AASAPPTLPGD....TQ......FAGIGN
WP_203203692.1      .................MPTHPPATPAEAPLPAALPT.................MAPVTT
WP_295989070.1      .SGP....................................EG....SG.....KP....L
MBT9459293.1        RPPV....................................AA....K.............
```

*ABM32059.1*

```
                        540                            550         560       570
ABM32059.1          PVPAPAPAP.....................APAPTASHG..PAGEEGTPKSPLAPVLPEL
NP_248771.1         .....QPQ....................................PQPQP..ASVAAPTPP
WP_001089704.1      .........................................PQLE.......KSAYSTMH
WP_200016399.1      PVDIPVQ.P.....................Q...........................PA
WP_063326054.1      PVDIPVQ.P.....................Q...........................PA
WP_012205666.1      PVDIPVQ.P.....................Q...........................PA
MBK9235436.1        PTPTPAPAP.....................AVAPMAIPR..IDEPA..........SAPA
MCA0243819.1        ..PTPFDPP.....................A..........AAAA...PPVPVDP.APAP
MCZ2441668.1        P....TAAP.....................E..........AADA...STAHTGT.APAS
WP_202760356.1      PVDIPVQ.P.....................Q...........................PA
WP_252978133.1      PVDAPVQ.P.....................Q...........................PA
WP_065346974.1      PVDIPVQ.P.....................Q...........................PA
WP_230355409.1      PVDIPVQ.P.....................Q...........................PA
WP_267179154.1      PVDAPVQ.P.....................Q...........................PA
WP_088164707.1      PVDIPVQ.P.....................Q...........................PA
WP_133073230.1      PVDAPVQ.P.....................Q...........................PA
MCC7098932.1        PPTAPLPPP.....................A..........PQVP...TPSPPDP.STVP
WP_238004152.1      PVDIPVQ.P.....................Q...........................PA
MBV8620386.1        PASAPLPAS.....................A...........................PP
ODU08341.1          ...................................................PTNP.STLP
MBC7710584.1        .......LAQ......................STPAA....PQR....I..APSAPSVTS
MBL8310054.1        .........................................PQ.........HSSAPRSA
WP_238363320.1      VAASPLPAMA......................PVPAA....PS...ARP..VAAFEPVAA
MCX8099153.1        PPAEPWAISS......................SKPAV....SG...AG...LAPPPPAST
MCM2252316.1        .............................................APRG..AAPAQAFQP
RYF42966.1          .................................PP..........AQPH..FSQAQHGQA
WP_281607349.1      PNLQPAPPPRAPHGAPPAHAMP.....PGPAMRSSQAVQPAPAMPSE........SGAQP
MDB5976135.1        .....LPPPEAPAAIAP..........................LPPKL.........PAA
WP_201212573.1      V.PEPPPTPVA.................................................
MBP8267915.1        .R..................................MA...PD........PVAAV....
MCZ2135183.1        ...PPLPPVV......................ASPAR....PQSPVPTS..VPQAPAGGA
WP_131145415.1      PNLQQAPHPQAGQGVPPADAMP.....PGPAMRPAPAVPPSAAVPSEYV..AQPAAGAQP
WP_166302793.1      PVAEPTPAPTIQA.......HE....VPIAALP........TQTPAATI..AAEPSPPPT
WP_267727965.1      PPVAPAPAPAA.....................ATPQPT...PAP.VSAP..PLATTPPVH
WP_299903544.1      .R..................................IE...AA........APE.S....
WP_065341524.1      APLTPAPTPAA..........................S...PVPPASAP..AAAAQPAAA
MBL8494843.1        APLSPAPVPVQPAAPLVARAFEEV....................PAAAP..ARAA..QAP
WP_058641008.1      DLLAPASPP................................................MPA
WP_183632487.1      APA.................................PT...ATP.PSAP.....APAPLP
RYY83951.1          .......PP...........................PP...PPPPVIAP....VAAAPTA
MCU0967987.1        ...........................................PSVQA....PAPARPA.
WP_106557035.1      .....A..A...........................PP...AVTPATTP....VAQAAPP
MBK7663138.1        IKSA........AEPMIARAFAEPGVVPAPAARPPAVA...SAPPATAP..KPAP..AVA
WP_203203692.1      PVVAPAPAA...........IEVPVEVPAEVVAPKASPEPVA...VTAP...SSPATDTA
WP_295989070.1      EPLEPLKPP...........................AP...APSPAARP....VQQQP..
MBT9459293.1        .PVDYAPT............................GM...ARPPAAA....ASPARPVA
```

```
                           α3            η4    η5                    α4
ABM32059.1
                       580       590       600       610       620       630
ABM32059.1          EAASPAELQSLIAAFLEGAGVLQKKVEPQKLSPEFMRSFGEAFRVAVQGTIDLLAARSEI
NP_248771.1         ASAAASAGGDLLQAFLRGAGMTQLKVD.PAGAEAQMEAIGRSYRGLVEGLVDVLRARASL
WP_001089704.1      TSSTSQANHLVTGPILEGLGVDLSDENDMNRMHLLSQELGESLQACVRGLLDLHQQVSES
WP_200016399.1      A...SADLQALAAAFMEGAGLNPAKTH.FEVTPEFMRTFGEALRIAVQGTIDLLAARSEI
WP_063326054.1      A...NADLQALAAAFMEGAGLNPAKTH.FEVTPEFMRTFGEALRIAVQGTIDLLAARSEI
WP_012205666.1      A...NADLQALAAAFMEGAGLNPAKTH.FEVTPEFMRTFGEALRIAVQGTIDLLAARSEI
MBK9235436.1        TRSAPGNPDELARAFLEGAGLSHVRVD.LTLSPEFMRAFGEAFRTAIEGTIDLLAARSEI
MCA0243819.1        AMTSAADLQALMQAFIEGIGLPPERVK.SELTPEFMQRVGQTFRTAIQGTMDLLHARSEI
MCZ2441668.1        .TPSADDVQALLQAFVQGLGLPPERAQ.FDLDPELMRRFGEVLRTAVQGTMELLQARAGI
WP_202760356.1      A...NADLQALAAAFMEGAGLNPAKTH.FEVTPEFMRTFGEALRIAVQGTIDLLAARSEI
WP_252978133.1      ASAAHADLQALATAFMEGAGLNPAKTH.FEVTPEFMRTFGEALRIAVQGTIDLLAARSEI
WP_065346974.1      A...NADLQALAAAFMEGAGLNPAKTH.FEVTPEFMRTFGEALRIAVQGTIDLLAARSEI
WP_230355409.1      A...NADLQALAAAFMEGAGLNPAKTH.FEVTPEFMRTFGEALRIAVQGTIDLLAARSEI
WP_267179154.1      ASAAHADLQALATAFMEGAGLNPAKTH.FEVTPEFMRTFGEALRIAVQGTIDLLAARSEI
WP_088164707.1      A...HADLQALAAAFMEGAGLNPAKTH.FEVTPEFMRTFGEALRIAVQGTIDLLAARSEI
WP_133073230.1      ASAAHADLQALATAFMEGAGLNPAKTH.FEVTPEFMRTFGEALRIAVQGTIDLLAARSEI
MCC7098932.1        .HPAAADEHALMQALLEGLGLPPERAP.FEMTPEFMRRFGELLRTAVQGTMDLLHARAEV
WP_238004152.1      A...HADLQALAVAFMEGAGLNPAKTH.FEVTPEFMRTFGEALRIAVQGTIDLLAARSEI
MBV8620386.1        PAAPTLDEQALIAAFLEGAGLAPDRVE.MRLTPEFMRHFGEAFKVAVQGTIDLIAARSEI
ODU08341.1          .APSPDDVQALLQAFVEGLGLPPARAQ.LELTPEFMRRFGEVLRTAVQGTMDLLQARSEV
MBC7710584.1        APITPASTDALSNAFLLGAGLAPSALP.HGLTPEIMTIVGSLLRSATAGAVDMLAARAAT
MBL8310054.1        TAPPAASQDALTEAFLKGAGLTPSALP.QGLTPEVMQLVGSVLRSATAGAVDMLAARAAT
WP_238363320.1      PQRPTASPDALTEAFLRGAGLPPSALP.QGLTPEVMMMVGSVLRSATAGAVDMLAARAAT
MCX8099153.1        AEALPNGVAVLTQAFLEGAELPASALP.NGLTAEQMRLIGAVLRAALAGAIDLLAARATT
MCM2252316.1        PPMPVGSADALARAFLTGARLPPAALP.HGLTPEVMALIGALLRSATAGAIDMLAVRANI
RYF42966.1          QPMPSYPADELTRAFLAGANLQPSALP.QGLTPEIMAVVGSVLHAATTGAIGMLAIRANI
WP_281607349.1      AGNAQADSAELLRAFLAGAGVPDLDVR.GPLTPQMMHTFGQLLREATQGTLDLLLSRAVI
MDB5976135.1        PDRAPGADSELLRAFLQGAGVPDLAMP.AELTPEMMNQFGQLLREAARGTLDLLLSRALV
WP_201212573.1      ..PPAGESDALLQAFLQGLGLPGLKLE.QGLTPEAMQTIGQLLREATQGTLDLLLARATT
MBP8267915.1        QPTLGGVEGALLEAFLRGLGVPV.RLP.AGLTPELMEQIGVVLREATQGTLDLLTARALT
MCZ2135183.1        PLATATNPDALTEAFLRGAGLPPSALP.NGLTPEVLAMVGSVLRSATAGAVDMLAARAAT
WP_131145415.1      AGDAQADSAELLRAFLAGAGVPDLDVR.GPLTPQMMHTFGQLLREATQGTLDLLLSRAVI
WP_166302793.1      PAALDDNAQQLLDAFKRGAGLNDCRYP.QQLTPEMMFMIGQMLGNSVQGCMDLLGSRAAA
WP_267727965.1      QSATSASPDALLAAFQRGLGFPL.QPS.DGLSPERMEQIGRVLRESVAGTMALLTARALT
WP_299903544.1      PPVAPAGHDVLMEAFLRGLGVPV.RLP.AGLSPELMEQIGVMLREATQGTLELLTARALT
WP_065341524.1      SAGSSADADALLAAFARGIGLPRLAPA.GGLTPELMEHIGRMLREAVQGTVELLVARAAT
MBL8494843.1        APVGDASAEALRRAFLAGLGTPTLPLD..ALTPEMMERVGRLLREATQGTLDLLLARATT
WP_058641008.1      VADASGDAGELLAAFLRGLGGMH.QPP.AQLTPALMERVGVMLRSATEGTLQLLLTRQEL
WP_183632487.1      AGNEATPPDVLLAAFQRGLGFPL.QPA.DGLSPERMEQIGRVLRESVAGTMALLTARALT
RYY83951.1          APSAPVSDDVLLAAFLRGLGSQH.QPP.EALTPGLMERIGVILRASTEGTLQLLARQEF
MCU0967987.1        .PAAAAAPDELLAALLDGLGAPGL.RI.EALTPARMHLVGELLRESVRGAVELLVARAAL
WP_106557035.1      AATPAAPPDALLAAFLRGLGSTH.QAP.QQLTPELMERIGVLLRSATEGTLQLLLTRQEF
MBK7663138.1        PASSAAFSEALRRAFLAGLGTPNLPLP.EAMTPEMMERVGRLLREATQGTLDLLLARAMT
WP_203203692.1      PALPAASCEALFRAFLDGAGVPDVAGQ.QQLDAEMMRRLGRLMRAFTDGTIELLSSRAML
WP_295989070.1      SHERMASDDELLAAFLRGLASTH.QPP.EMLTPGLMERIGSMLRSATEGTLQLLLTRQEF
MBT9459293.1        SSVGSADTEALLAAFREGLATPTV.QV.DALTPELMKLIGQLIHESAKGTVDLLVARAAL
```

```
                                α5       α6                              α7
ABM32059.1                                          TT
                           640       650       660       670       680       690
ABM32059.1          KREFRAGVTVISSGANNPLKFLPTPEGVVMQMIGQN.FPGFMKPIPAMQEAYDDLRVHQV
NP_248771.1         KGEFRMAQTMIQPVQNNPLKFAPNVDEAMLLLLRRD.NQAFMAPDRAVADSFEDLKAHQL
WP_001089704.1      R..FGTLNRNLQPIEDNPLRLGLSYEETIRTLYDADKSVVHLSAPAAIAESLKTVRDHNE
WP_200016399.1      KREFRAGVTIIATGANNPLKFLPNSEGVLMQMVHQT.FPGFMKPLPAMQDAFQDLHVHQL
WP_063326054.1      KREFRAGVTIIATGANNPLKFLPNSEGVLMQMVHQT.FPGFMKPLPAMQDAFQDLHVHQL
WP_012205666.1      KREFRAGVTIIATGANNPLKFLPNSEGVLMQMVHQT.FPGFMKPLPAMQDAFQDLHVHQL
MBK9235436.1        KREFRAGVTIMASGANNPLKFLPNADGVIMQLAGQG.FPGFMKPVPAMKEAYRDLQVHQL
MCA0243819.1        KREFRADVTIMANRANNPLKFLPDADGVMMQMFGQT.FPGFMRPVPAMQEAYRDLRVHQL
MCZ2441668.1        KREFRADVTIIGNRANNPLKFLPDADGVLLQMLGQT.FPGFMRPVPAMHEAYRDLRVHQS
WP_202760356.1      KREFRAGVTIIATGANNPLKFLPNSEGVLMQMVHQT.FPGFMKPLPAMQDAFQDLHVHQL
WP_252978133.1      KREFRAGVTIIATGANNPLKFLPNSEGVLMQMVHQT.FPGFMKPLPAMQDAFQDLHVHQL
WP_065346974.1      KREFRAGVTIIATGANNPLKFLPNSEGVLMQMVHQT.FPGFMKPLPAMQDAFQDLHVHQL
WP_230355409.1      KREFRAGVTIIATGANNPLKFLPNSEGVLMQMVHQT.FPGFMKPLPAMQDAFQDLHVHQL
WP_267179154.1      KREFRAGVTIIATGANNPLKFLPNSEGVLMQMVHQT.FPGFMKPLPAMQDAFQDLHVHQL
WP_088164707.1      KREFRAGVTIIATGANNPLKFLPNSEGVLMQMVHQT.FPGFMKPLPAMQDAFQDLHVHQL
WP_133073230.1      KREFRAGVTIIATGANNPLKFLPNSEGVLMQMVHQT.FPGFMKPLPAMQDAFQDLHVHQL
MCC7098932.1        KREFRADVTIMANRANNPLKFLPDAEGVLLQMLGQT.FPGFTRPVPAMREAFRDLRVHQL
WP_238004152.1      KREFRAGVTIIATGANNPLKFLPNSEGVLMQMVHQT.FPGFMKPLPAMQDAFQDLHVHQL
MBV8620386.1        KREFRADVTVIASAANNPLKFLPTAEGVIMQLTGRS.FPGFMAPVPAMKEAYHDLQVHQL
ODU08341.1          KREFRADVTIMAQRANNPLKFLPDADGVLLQMFGQT.FPGFMRPVPAMREAYRDLRVHQI
MBC7710584.1        KREVQASVTIISVQANNPLKFLPNADAALLQLLGKK.MPGFMRADVAMRDAFDDLRAHEV
MBL8310054.1        KREVQASVTIISVQANNPLKFLPNADAALMQLLGKK.MPGFMRADVAMRDAFDDLRAHEV
WP_238363320.1      KREVQASVTIISVQANNPLKFLPNADAAMLQLLGKK.MPGFMRADVAMRDAFDDLRAHEV
MCX8099153.1        KRELQASVTIISVEANNPLKFLPNADAALTQLLGRK.MPGFMRADVAMRDAFEDLRAHEV
MCM2252316.1        KREVQASVTIISTEANNPLKFLPNAEAVLLQFLGRA.MPGFMRPDEAMDDAFNDLRAHEL
RYF42966.1          KREVQANVTIISSEANNPLKFLPDAETVLQQLLGKK.IPGFMGPDEAMKDAFNDLRAHEA
WP_281607349.1      KREIHAEMTMIVTRENNPLKFSPNVEVALTHLLAPR.GQGFMPPLVAMKDAHDDLRAHQF
MDB5976135.1        KQEVRADVTIIRPRENNPLKFSPSVEVALAHLLAPR.GGGFMAPVPAMKDAFDDLRAHQF
WP_201212573.1      KREVRAEVTMIVSRDNNPLKFSPSVEVALMHLLAPQ.GRGFLPPLAAMRDAWDDLRAHQL
MBP8267915.1        KREVRAEVTMIVSKGNNPLKFSPDVGFALSQLLAPQ.GRGFMPPEEAMRDAYDDLRAHQF
MCZ2135183.1        KREVQASVTIISVQANNPLKFLPNADAALLQLLSKK.MPGFMRADVAMRDAFDDLRAHEV
WP_131145415.1      KREIHAEMTMIVTRENNPLKFSPNVEVALTHLLAPR.GQGFMPPLVAMKDAHDDLRAHQF
WP_166302793.1      KQEVRIAVTLINAEANNPLKFLPTGAAALAQIFGPR.MPGFLSGPAAIESAHLDLRAHEV
WP_267727965.1      KREIRAEVTMIVSKENNPLKFAPDVEFALVQLLAPQ.GRGFMEPVAAMRDAYDDLRSHQF
WP_299903544.1      KREVRAEVTMIVSKGNNPLKFSPDVGFALSQLLAPQ.GRGFMPPKEAMRDAYDDLRAHQF
WP_065341524.1      KREVRADVTMIVSKNNNPLKFSPDVDFALMQLLLPQ.GSGFMKPDEAMQDAYDDLRAHQL
MBL8494843.1        KREVRADVTMIVSSDNNPLKFSPDVAAALLHLLVPQ.GPGFLSPVAAMQDAYDDLRSHQF
WP_058641008.1      KREVRAEVTMIAAQANNPLKFSPTVEVALGHLLGPP.LRGFMPAEAAMRDAYDDLRAHQF
WP_183632487.1      KREIRAEVTMIVSKENNPLKFAPDVEFALVQLLAPQ.GRGFMEPVKAMRDAYDDLRSHQF
RYY83951.1          KREVRAEVTMIASQANNPLKFSPTVEVALAHLLGPG.VRGFMPPEAAMRDAYNDLRAHQF
MCU0967987.1        KREMRAQMTMIVARENNPLKFSPSVEVALQHLLGPP.APGFMAPAPAVRDAFDDLRAHQL
WP_106557035.1      KRELRAEVTMIAAQANNPLKFSPTAEVALAHLLGPG.MRGFLPAEAAMRDAFDDLRSHQF
MBK7663138.1        KREVRADVTMIVSTDNNPLKFSPDVGAALLHLLVPQ.GPGFLTPVAAMRDAYDDLRSHQF
WP_203203692.1      KREVRAEITMIVDEENNPFKILPNGRAVLMQMFGAR.MPGFLPPEAAVHDALGDLQSHQL
WP_295989070.1      KREVRAEVTMIAAQANNPLKFSPTVEVALAHLLGPG.VRGFMPPEAAMRDAFNDLRAHQF
MBT9459293.1        KREVRAEATMIVARENNPLKFSPSVEAALQHLLSPP.TRGFMPAAPAMRDAYDDLRAHQF
```

```
                                     α8                              α9
ABM32059.1
                        700          710        720        730          740
                         .            .          .          .            .
ABM32059.1       ALMAGLRAAYAEALERFDPAALEQR.TDVTGGVFGKLSATSRKAALWDEYKRNFAEIRRG
NP_248771.1      AVMAGVQAAIRHLLARFEPAALEAR.FGKPAGL.SGLLPGARQAQNWDSFTELYAKILRE
WP_001089704.1   AMQFATSEALSQILNAFSPQVMLRR.FHHYKRN.SDTTQTSTDAWAWNMYCSYYQELTSN
WP_200016399.1   ALMAGIRAAYAEALTRFDPTELERR.TETNPGLLDKLLTNGRKAALWDDYKHNFDVLRRH
WP_063326054.1   ALMAGIRAAYAEALTRFDPTELERR.TETNPGLLDKLLTNGRKAALWDDYKHNFDVLRRH
WP_012205666.1   ALMAGIRAAYAEALTRFDPTELERR.TETNPGLLDKLLTNGRKAALWDDYKHNFDVLRRH
MBK9235436.1     GLMAGIRAAYTAALTRFDPLELESQ.APSRPGLLRGFSSLKRKAALWDHYKQGYTRIRQH
MCA0243819.1     ALMAGIRAAYAEALTRFDPAELERQ.AGDPGGLLARLGRNRHKAALWDDYKQRYAEIRRN
MCZ2441668.1     ALMAGLRAAFSEALSRLDPAALEKQ.VGASGSLLARLGRTHRRAALWDDYCERYAAIRRQ
WP_202760356.1   ALMAGIRAAYAEALTRFDPTELERR.TETNPGLLDKLLTNGRKAALWDDYKHNFDVLRRH
WP_252978133.1   ALMAGIRAAYAEALTRFDPTELERR.TETNPGLLDKLLTNGRKAALWDDYKHNFDVLRRH
WP_065346974.1   ALMAGIRAAYAEALTRFDPTELERR.TETNPGLLDKLLTNGRKAALWDDYKHNFDVLRRH
WP_230355409.1   ALMAGIRAAYAEALTRFDPTELERR.TETNPGLLDKLLTNGRKAALWDDYKHNFDVLRRH
WP_267179154.1   ALMAGIRAAYAEALTRFDPTELERR.TETNPGLLDKLLTNGRKAALWDDYKHNFDVLRRH
WP_088164707.1   ALMAGIRAAYAEALTRFDPTELERR.TETNPGLLDKLLTNGRKAALWDDYKHNFDVLRRH
WP_133073230.1   ALMAGIRAAYAEALTRFDPTELERR.TETNPGLLDKLLTNGRKAALWDDYKHNFDVLRRH
MCC7098932.1     ALMAGIRAAYAEALSRLDPAELEKQ.GGAGEGLLARLGRTHRKAALWDDYHARYDGLRRQ
WP_238004152.1   ALMAGIRAAYAEALTRFDPTELERR.TETNPGLLDKLLTNGRKAALWDDYKHNFDVLRRH
MBV8620386.1     ALMAGIRAAYAEALTRFDPVELEKQ.FPASGSLIGRWSAARRKAGLWDGYRQSYLDIRRH
ODU08341.1       ALMAGMRAALAEALSRLDPAELEQQQPGASDGLFARLGRVHRKAALWDDYRERYAGIRRH
MBC7710584.1     GVIAGTRAALTEVLGKFDPAVLGDK.LAKGSVL.ESLMPSARKAKLWDQYLERYLQIRRE
MBL8310054.1     GVIAGTRAALTEVLGKFSPDILGDK.LTRGSVL.ESLLPSARKAKLWDTYLDRYQQIRRE
WP_238363320.1   GVIAGTRAALTEVLGKFDPAVLGEK.LAKGSVL.ESVMPSARKAKLWDMYLERYLQIRRE
MCX8099153.1     GVIAGTRAALAEVLNKFDPECLERR.LSKGSVL.EALLPAARKARLWEQYVKLFSQVRRE
MCM2252316.1     GVIAGTRAALDEVLAKFDPAILGER.LIRGSLL.EKALPAMRKTRLWDLYLERYAQLRRE
RYF42966.1       GVIAGTRAALGEVLNKFDPAVLGER.LVGGSLL.ERALPAVRKTKLWDLYLERYAHIRRE
WP_281607349.1   AFMAGMRAALAGVLQRFDPAQLERR.LSGKSLV.DTLLPANRKAKLWDLYAEMYSELTRE
MDB5976135.1     GFMAGMRAALEGVLERFDPAQLELR.LTEKSVF.DSLLPASRKAKLWDRFSELYTEIGKE
WP_201212573.1   GFMAGMRAALAGVLQRFDPATLEQR.LSAKSLL.DSLLPMNRRAHLWELYISLYREIARE
MBP8267915.1     GFMAGMRAALAGVLKRFEPASLEQR.VVARSLL.DNLLPGSRKARLWDLFEQMYGEISRE
MCZ2135183.1     GVIAGTRAALTEVLGKFDPTVLGDK.LTRGSVL.ESLLPSARKAKLWDMYLERYLQIRRE
WP_131145415.1   AFMAGMRAALAGVLQRFDPAQLERR.LSGKSLV.DTLLPANRKAKLWDLYAEMYSELTRE
WP_166302793.1   GMMAGTQAAVQRLFERFDPQEIEQQ.LDAQGRQ.RTLFAAQRNARLWEMYCSRFQWLKEE
WP_267727965.1   GFMAGMRAALSGVLDRFKPEQLEGR.ISERGFL.DSVLPGARKAKLWDLYEQHFADISRE
WP_299903544.1   GFMAGMRAALAGVLKRFEPAALEQR.VVARSLL.DNLLPGSRKARLWDLFEQMYGEISQE
WP_065341524.1   GFMAGMRAALASILGRFTPAELEAR.LTTKSFL.DNVLPANRKAKLWDLYEQRYGDISRE
MBL8494843.1     GFMAGLRAALAGVLKRFDPAMLEQR.MAQKSML.DNVLPMNRRAKLWDLYIQLYRDIANE
WP_058641008.1   GVMVGLRAALASLIERFGPEALEKK.IAARSAL.DSLFAANRKARLWDQFTALYGTIAAE
WP_183632487.1   GFMAGMRAALTGVLARFKPEQLEGR.ISARGFL.DNVLPGSRKARLWDLYEQHYADISRE
RYY83951.1       GVMVGMRAALAHVIARFAPDELEKK.IAARSAL.DSLFAANRKAKLWDQFMALYGGIATE
MCU0967987.1     GVMAGMRAALEGVLQRFDPAQLEGQ.LTRGSAL.ASLIPSTRKARLWERFQELYAQLNRE
WP_106557035.1   GVMVGMRAALAQVIERFAPAELEKK.IAARSAL.DSLFAANRKAKLWDQFIALYGGIASE
MBK7663138.1     GFMAGLRAALAGVLKRFDPAVLEQR.LAQKSVL.DSLLPINRRAKLWDLYISLYRDIAVE
WP_203203692.1   GMVAGMRAALLMVLQRFDPEALKKD.TPHDGGLGEKLLPGGRDARLWRQLQQLHAETTAA
WP_295989070.1   GVMVGMRAALAHVIARFEPAELEKK.ISAKTAL.DALFSANRKAKLWDQFVSLYGGIASE
MBT9459293.1     GFIAGMRAALEGVLARFDPAELEGR.LSQRSGL.MSILPGSRKARMWEVFIEHYAQIRSD
```

```
                           α10
ABM32059.1
                750        760        770
                 .          .          .
ABM32059.1       AEDDLAAFSGQAFLEAYENAEAAAKART.............
NP_248771.1      AEDDFQELFGREFSRAYEEHSARLRRS..............
WP_001089704.1   RQRGFEKLFWEIFEQAYDRKIREKQLEL.............
WP_200016399.1   AEDDLMAFSGRTFVEAYENAEQSVKGQP.............
WP_063326054.1   AEDDLMAFSGRTFVEAYENAEQSVKGQP.............
WP_012205666.1   AEDDLMAFSGRTFVEAYENAEQSVKGQP.............
MBK9235436.1     AEDDLKAFSGQTFVNAYEAAADAAKDAL.............
MCA0243819.1     AEDDLTAFSGRTFVKAYDAAADAALDEDELLLRKKT.....
MCZ2441668.1     AEDELTAFSGRSFVKAYDAAADAALADDELPPGKPIRA...
WP_202760356.1   AEDDLMAFSGRTFVEAYENAEQSVKGQP.............
WP_252978133.1   AEDDLMAFSGRTFVEAYENAEQSVKGQP.............
WP_065346974.1   AEDDLMAFSGRTFVEAYENAEQSVKGQP.............
WP_230355409.1   AEDDLMAFSGRTFVEAYENAEQSVKGQP.............
WP_267179154.1   AEDDLMAFSGRTFVEAYENAEQSVKGQP.............
WP_088164707.1   AEDDLMAFSGRTFVEAYENAEQSVKGQP.............
WP_133073230.1   AEDDLMAFSGRTFVEAYENAEQSVKGQP.............
MCC7098932.1     AEDELSAFSGRSFVKAYDAAADAALCDDELPVARPPRTPAS
WP_238004152.1   AEDDLMAFSGRTFVEAYENAEQSVKGQP.............
MBV8620386.1     AEDDLTAFSGRTFVQAYEAAAEAAGLAP.............
ODU08341.1       AEDELTAFSGRSFVKAYDATADAAFADDELPPGKPIRA...
MBC7710584.1     AEDDFQSIFGRAFVQAYERETARIKGESPTAGGE.......
MBL8310054.1     AEDDFQSIFGRAFVQAYERETARIKGDTSGDTGKSSGGT..
WP_238363320.1   AEDDFQSIFGRAFVQAYERETARIKGEVPASGGDR......
MCX8099153.1     AEDDFHNVFGRAFVEAYERESRRIKGGTP............
MCM2252316.1     VEDDFQSVFGKAFVAAYESETARMKARSTDRGP........
RYF42966.1       AEDDFQSIFGRAFVEAYERETARMKKAGR............
WP_281607349.1   AEDDFHAWFGKAFLNAYEAQLDKLARGDEGGR.........
MDB5976135.1     AGDDFHALFGKAFLRAYEAQLAQLEQAGRADTR........
WP_201212573.1   AEDDFHALFGREFVRAYEEQVRRLHAADE............
MBP8267915.1     AEDDFHALFGREFLRAYEEQVARLQAEREPDR.........
MCZ2135183.1     AEDDFQSIFGRAFVQAYERETARIRGDGAVSGGDRS.....
WP_131145415.1   AEDDFHAWFGKAFLNAYEAQLDKLARGDEGGR.........
WP_166302793.1   MKNQTPASWGTEFHSAYQNETDSIDRKGPRK..........
WP_267727965.1   AEDDFQTLFGRAFLRAYEEQIDRLASDPGRE..........
WP_299903544.1   AEDDFHALFGREFLRAYEEQVERLQAEREPDR.........
WP_065341524.1   AEDDFHSLFGREFLKAYEAQIDQLQRERG............
MBL8494843.1     AEDDFHTLFGREFLRAYQEQVDRLEKSQ.............
WP_058641008.1   AEDDFHSLFGQAFLAAYEAQMARLKQGAAEAPPPAA.....
WP_183632487.1   AEDDFQTLFGHAFLKAYEEQIDRLASDPGRE..........
RYY83951.1       AEDDFHSLFGKAFLEAYEEQMARLKSADQ............
MCU0967987.1     AQDDFDELFGKAFLRAYEEQLERLQHEPRG...........
WP_106557035.1   AEDDFHALFGKAFLEAYEAQMARLKSGGS............
MBK7663138.1     AEDDFHTLFGREFLRAYQEQVERIGQGKP............
WP_203203692.1   VEDDFQAVFGRAFQQAYDREMERLKEARRA...........
WP_295989070.1   AEDDFHSLFGKAFLEAYEEQMARLKSDA.............
MBT9459293.1     ASDDFHTLFGKAFLEAYEEHIDQLKGDSP............
```
