## Supplementary Data 2 for "LLPS condensates of Fha initiate the inside-out assembly of the type VI secretion system"

| T6SS ID | Type | Organism | Fha | Location | Accession |
|---|---|---|---|---|---|
| T6SS00164 | i3 | Acidovorax citrulli AAC00-1 | 1 | 1617808..1647773 | chromosome (NC_008752) |
| T6SS00312 | i4b | Acinetobacter baumannii AB307-0294 | 0 | 2367193..2386603 | chromosome (NC_011595) |
| T6SS01164 | i4b | Acinetobacter baumannii AbCAN2 | 0 | 1213019..1232429 | chromosome (NZ_CP045428) |
| T6SS01046 | i4b | Acinetobacter baumannii ATCC 17978 | 0 | 2433628..2461317 | chromosome (NZ_CP012004) |
| T6SS01275 | i4b | Acinetobacter baumannii DSM 30011 | 0 | 536657..556068 | chromosome (NZ_JJOC02000004) |
| T6SS01119 | i4b | Acinetobacter baumannii strain UPAB1 | 0 | 2933094..2965138 | chromosome (NZ_CP032215) |
| T6SS00061 | i4b | Acinetobacter sp. ADP1 | 0 | 2633143..2661510 | chromosome (NC_005966) |
| T6SS00788 | i1 | Aeromonas dhakensis strain SSU | 1 | 1017876..1061054 | chromosome (NZ_JH815591) |
| T6SS01036 | i1 | Aeromonas hydrophila J-1 | 1 | 2866359..2898257 | chromosome (NZ_CP006883) |
| T6SS00948 | i1 | Aeromonas hydrophila NJ-35 | 1 | 2935924..2976020 | chromosome (CP006870) |
| T6SS17503 | i1 | Aeromonas hydrophila strain NF1 | 1 | 42575..80489 | chromosome (NZ_JDWB01000021) |
| T6SS00157 | i1 | Aeromonas hydrophila subsp. hydrophila ATCC 7966 | 1 | 1984799..2036268 | chromosome (NC_008570) |
| T6SS01208 | i1 | Aeromonas veronii strain TH0426 | 1 | 2455395..2478287 | chromosome (NZ_CP012504) |
| T6SS00016 | i5 | Agrobacterium fabrum str. C58 | 1 | 1460756..1488734 | chromosome linear (NC_003063) |
| T6SS01243 | i5 | Agrobacterium fabrum strain 12D13 | 1 | 1420086..1444587 | chromosome (NZ_CP033035) |
| T6SS01240 | i5 | Agrobacterium fabrum strain 1D132 | 1 | 1429744..1454431 | chromosome (NZ_CP033023) |
| T6SS01241 | i5 | Agrobacterium tumefaciens strain 12D1 | 1 | 1717106..1741674 | chromosome (NZ_CP033032) |
| T6SS01245 | i5 | Agrobacterium tumefaciens strain 15955 | 1 | 1999377..2023762 | chromosome (NZ_CP032918) |
| T6SS01242 | i5 | Agrobacterium tumefaciens strain 1D1108 | 1 | 2042910..2067248 | chromosome (NZ_CP032922) |
| T6SS01244 | i5 | Agrobacterium tumefaciens strain 1D1460 | 1 | 1440614..1465080 | chromosome (NZ_CP032927) |
| T6SS01103 | i5 | Agrobacterium tumefaciens strain 1D1609 | 1 | 1612910..1635897 | chromosome (NZ_CP026925) |
| T6SS01246 | i5 | Agrobacterium tumefaciens strain A6 | 1 | 2000609..2024994 | chromosome (NZ_CP033028) |

| | | | | | |
|---|---|---|---|---|---|
| T6SS00084 | i1 | Aliivibrio fischeri ES114 | 1 | 1087244..1118010 | chromosome I (NC_006840) |
| T6SS17506 | i1 | Aliivibrio fischeri strain ES401 | 1 | 345059..392876 | chromosome (NZ_SRJG01000005) |
| T6SS01278 | i1 | Aliivibrio fischeri strain FQ-A001 | 1 | 316834..358016 | chromosome (NZ_SJSX01000004) |
| T6SS01277 | i1 | Aliivibrio fischeri strain FQ-A001 | 1 | 747470..766109 | chromosome (NZ_SJSX01000002) |
| T6SS00161 | i3 | Azoarcus olearius | 1 | 4242264..4289066 | chromosome (NC_008702) |
| T6SS00009 | i3 | Bordetella bronchiseptica RB50 | 1 | 825621..868006 | chromosome (NC_002927) |
| T6SS01063 | i3 | Bradyrhizobium japonicum strain J5 | 1 | 3553506..3574647 | chromosome (NZ_CP017637) |
| T6SS00126 | i4b | Burkholderia cenocepacia AU 1054 | 0 | 2880426..2919827 | chromosome I (NC_008060) |
| T6SS01197 | i2 | Burkholderia cenocepacia H111 | 0 | 172983..201438 | chromosome (NZ_HG938370) |
| T6SS01196 | i4b | Burkholderia cenocepacia H111 | 0 | 347990..367838 | chromosome (NZ_HG938371) |
| T6SS00292 | i4b | Burkholderia cenocepacia J2315 | 0 | 363015..382880 | chromosome I (NC_011000) |
| T6SS01276 | i4b | Burkholderia cenocepacia strain K56-2 | 0 | 63074..82939 | chromosome (NZ_LAUA01000004) |
| T6SS01301 | i3 | Burkholderia gladioli strain NGJ1 | 1 | 55199..85982 | chromosome (LEKY01000012) |
| T6SS01300 | i4b | Burkholderia gladioli strain NGJ1 | 0 | 91265..111362 | chromosome (LEKY01000074) |
| T6SS01265 | i4b | Burkholderia glumae 336gr-1 | 0 | 2..8161 | chromosome (AYPW01000191) |
| T6SS17509 | i2 | Burkholderia glumae BGR1 | 0 | 1382281..1422638 | chromosome I (NC_012724) |
| T6SS00343 | i4b | Burkholderia glumae BGR1 | 0 | 397352..417437 | chromosome II (NC_012721) |
| T6SS00070 | i1 | Burkholderia mallei ATCC 23344 | 0 | 735641..772695 | chromosome II (NC_006349) |
| T6SS01295 | i3 | Burkholderia pseudomallei K96243 | 0 | 2499073..2525393 | chromosome II (CP009537) |
| T6SS01294 | i3 | Burkholderia pseudomallei K96243 | 0 | 2018346..2040803 | chromosome II (CP009537) |
| T6SS01293 | i3 | Burkholderia pseudomallei K96243 | 0 | 1914678..1943867 | chromosome II (CP009537) |
| T6SS01292 | i3 | Burkholderia pseudomallei K96243 | 1 | 1455143..1478378 | chromosome I (CP009538) |
| T6SS01006 | i4b | Burkholderia pseudomallei K96243 | 0 | 2062195..2082181 | chromosome II (CP009537) |
| T6SS00115 | i3 | Burkholderia thailandensis E264 | 0 | 3396298..3416317 | chromosome II (NC_007651) |
| T6SS00114 | i4b | Burkholderia thailandensis E264 | 0 | 2282208..2308569 | chromosome I (NC_007650) |

| | | | | | |
|---|---|---|---|---|---|
| T6SS00113 | i1 | Burkholderia thailandensis E264 | 0 | 996276..1025886 | chromosome II (NC_007650) |
| T6SS01220 | i1 | Campylobacter jejuni strain WP2202 | 1 | 13397..30371 | plasmid pCJDM202 (CP014743) |
| T6SS01013 | i3 | Citrobacter rodentium DBS100 | 0 | 2124567..2151602 | chromosome (CP038008) |
| T6SS00346 | i4b | Edwardsiella ictaluri 93-146 | 0 | 2631752..2651730 | chromosome (NC_012779) |
| T6SS00372 | i4b | Edwardsiella tarda EIB202 | 0 | 2555437..2575336 | chromosome (NC_013508) |
| T6SS00791 | i4b | Edwardsiella tarda PPD130/91 | 0 | 3625..24399 | chromosome (AY424360) |
| T6SS14098 | i2 | Enterobacter cloacae strain DSM 30054 | 0 | 4297267..4345922 | chromosome (NZ_CP056776) |
| T6SS14097 | i3 | Enterobacter cloacae strain DSM 30054 | 1 | 4037435..4081387 | chromosome (NZ_CP056776) |
| T6SS00396 | i2 | Enterobacter cloacae subsp. cloacae ATCC 13047 | 0 | 1819466..1844686 | chromosome (NC_014121) |
| T6SS00395 | i3 | Enterobacter cloacae subsp. cloacae ATCC 13047 | 1 | 1551980..1588894 | chromosome (NC_014121) |
| T6SS13923 | i3 | Enterobacter sp. DSM 30060 | 1 | 4465150..4496796 | chromosome (NZ_CP056118) |
| T6SS00464 | i4b | Escherichia coli 042 | 0 | 4893076..4921378 | chromosome (NC_017626) |
| T6SS00463 | i2 | Escherichia coli 042 | 0 | 4852875..4885437 | chromosome (NC_017626) |
| T6SS00935 | i2 | Escherichia coli 17-2 | 0 | 909481..942047 | chromosome (NZ_JACEFV010000001) |
| T6SS00931 | i4b | Escherichia coli 17-2 | 0 | 883479..901846 | chromosome (NZ_JACEFV010000001) |
| T6SS00037 | i2 | Escherichia coli CFT073 | 0 | 3217751..3253963 | chromosome (NC_004431) |
| T6SS00937 | i1 | Escherichia coli DE719 | 1 | 1..24039 | chromosome (KM251715) |
| T6SS01200 | i2 | Escherichia coli KTE165 | 0 | 180921..202372 | chromosome (NZ_KB733141) |
| T6SS01034 | i1 | Escherichia coli PCN033 | 1 | 246100..267921 | chromosome (NZ_CP006632) |
| T6SS00787 | i1 | Escherichia coli RS218 | 1 | 1..23615 | chromosome (JN837480) |
| T6SS00930 | i1 | Escherichia coli SEPT362 | 1 | 11200..34631 | chromosome (NZ_AOGL01000004) |
| T6SS00047 | i1 | Helicobacter hepaticus ATCC 51449 | 1 | 227242..253067 | chromosome (NC_004917) |
| T6SS01215 | i2 | Klebsiella pneumoniae strain Kp52.145 | 0 | 2564540..2589406 | chromosome (NZ_FO834906) |
| T6SS00548 | i2 | Klebsiella pneumoniae subsp. pneumoniae HS11286 | 0 | 2326237..2353260 | chromosome (NC_016845) |

| | | | | | |
|---|---|---|---|---|---|
| T6SS00344 | i2 | Klebsiella pneumoniae subsp. pneumoniae NTUH-K2044 | 0 | 2298791..2328477 | chromosome (NC_012731) |
| T6SS01283 | i2 | Kosakonia oryzae strain KO348 | 0 | 107454..120808 | chromosome (NZ_JZLI01000006) |
| T6SS01282 | i3 | Kosakonia oryzae strain KO348 | 1 | 149587..193032 | chromosome (NZ_JZLI01000015) |
| T6SS00443 | i3 | Methylomonas methanica MC09 | 0 | 915765..926524 | chromosome (NC_015572) |
| T6SS00127 | i1 | Myxococcus xanthus DK 1622 | 0 | 5996770..6055445 | chromosome (NC_008095) |
| T6SS01222 | i3 | Pantoea ananatis LMG 2665 | 1 | 9267..43295 | chromosome (KF590027) |
| T6SS00275 | i3 | Paraburkholderia phymatum STM815 | 0 | 654398..678783 | plasmid pBPHY01 (NC_010625) |
| T6SS00274 | i4a | Paraburkholderia phymatum STM815 | 0 | 510771..535444 | plasmid pBPHY01 (NC_010625) |
| T6SS00040 | i1 | Pectobacterium atrosepticum SCRI1043 | 1 | 3829218..3869582 | chromosome (NC_004547) |
| T6SS01170 | i1 | Pectobacterium brasiliense 1692 | 1 | 2749401..2774372 | chromosome (NZ_CP047495) |
| T6SS00050 | i1 | Photorhabdus laumondii subsp. laumondii TTO1 | 1 | 372357..415137 | chromosome (NC_005126) |
| T6SS00838 | i1 | Proteus mirabilis BB2000 | 1 | 916585..943300 | chromosome (NC_022000) |
| T6SS00272 | i1 | Proteus mirabilis HI4320 | 1 | 807789..834462 | chromosome (NC_010554) |
| T6SS01213 | i3 | Pseudomonas aeruginosa PAK | 1 | 5670794..5696508 | chromosome (NZ_CP020659) |
| T6SS01212 | i4a | Pseudomonas aeruginosa PAK | 0 | 1913764..1933758 | chromosome (NZ_CP020659) |
| T6SS01211 | i1 | Pseudomonas aeruginosa PAK | 0 | 1118829..1137804 | chromosome (NZ_CP020659) |
| T6SS00005 | i4a | Pseudomonas aeruginosa PAO1 | 0 | 2599893..2632231 | chromosome (H3-T6SS) (NC_002516) |
| T6SS00004 | i1 | Pseudomonas aeruginosa PAO1 | 1 | 1797958..1827821 | chromosome (H2-T6SS) (NC_002516) |
| T6SS00003 | i3 | Pseudomonas aeruginosa PAO1 | 1 | 83380..123495 | chromosome (H1-T6SS) (NC_002516) |
| T6SS00152 | i1 | Pseudomonas aeruginosa UCBPP-PA14 | 1 | 3806993..3840213 | chromosome (NC_008463) |
| T6SS00151 | i4a | Pseudomonas aeruginosa UCBPP-PA14 | 0 | 3011563..3044708 | chromosome (NC_008463) |
| T6SS00150 | i3 | Pseudomonas aeruginosa UCBPP-PA14 | 1 | 82730..121410 | chromosome (NC_008463) |
| T6SS00540 | i3 | Pseudomonas fluorescens F113 | 1 | 6672386..6694009 | chromosome (NC_016830) |
| T6SS00539 | i4a | Pseudomonas fluorescens F113 | 0 | 2807673..2826705 | chromosome (NC_016830) |

| | | | | | |
|---|---|---|---|---|---|
| T6SS00764 | i3 | Pseudomonas protegens CHA0 | 1 | 6697981..6718631 | chromosome (NC_021237) |
| T6SS00035 | i3 | Pseudomonas protegens Pf-5 | 1 | 6897920..6930515 | chromosome (NC_004129) |
| T6SS00015 | i1 | Pseudomonas putida KT2440 | 0 | 4590927..4618219 | chromosome (NC_002947) |
| T6SS00014 | i4b | Pseudomonas putida KT2440 | 0 | 3470069..3499780 | chromosome (NC_002947) |
| T6SS00013 | i1 | Pseudomonas putida KT2440 | 1 | 2984253..3012550 | chromosome (NC_002947) |
| T6SS01297 | i4b | Pseudomonas sp. JY-Q | 0 | 2910641..2930528 | chromosome (NZ_CP011525) |
| T6SS01219 | i4a | Pseudomonas syringae pv. actinidiae str. Shaanxi_M228 | 0 | 3616254..3638573 | chromosome (NZ_CP032631) |
| T6SS00086 | i4b | Pseudomonas syringae pv. syringae B728a | 0 | 5866174..5915475 | chromosome (NC_007005) |
| T6SS00042 | i1 | Pseudomonas syringae pv. tomato str. DC3000 | 1 | 6149242..6189243 | chromosome (NC_004578) |
| T6SS00026 | i4b | Ralstonia solanacearum GMI1000 | 0 | 926494..971950 | plasmid pGMI1000MP (NC_003296) |
| T6SS01261 | i5 | Rhizobium etli bv. mimosae str. Mim1 | 1 | 489772..515517 | plasmid pRetMIM1f (NC_021911) |
| T6SS00897 | i5 | Rhizobium leguminosarum bv. trifolii | 1 | 744..17137 | chromosome (AF361470) |
| T6SS00301 | i3 | Salmonella enterica subsp. enterica serovar Dublin str. CT_02021853 | 0 | 305501..331344 | chromosome (NC_011205) |
| T6SS00303 | i1 | Salmonella enterica subsp. enterica serovar Gallinarum str. 287/91 | 1 | 1114451..1137704 | chromosome (NC_011274) |
| T6SS00896 | i3 | Salmonella enterica subsp. enterica serovar Typhimurium | 0 | 322..27706 | chromosome (AJ320483) |
| T6SS00551 | i3 | Salmonella enterica subsp. enterica serovar Typhimurium str. 14028S | 0 | 305365..332747 | chromosome (NC_016856) |
| T6SS00023 | i3 | Salmonella enterica subsp. enterica serovar Typhimurium str. LT2 | 0 | 304665..339275 | chromosome (NC_003197) |
| T6SS00535 | i3 | Salmonella enterica subsp. enterica serovar Typhimurium str. SL1344 | 0 | 304656..332038 | chromosome (NC_016810) |

| | | | | | |
|---|---|---|---|---|---|
| T6SS01291 | i3 | Salmonella enterica subsp. enterica serovar Typhimurium strain ATCC 14028 | 0 | 10531..37913 | chromosome (NZ_MTFW01000011) |
| T6SS01264 | i3 | Serratia marcescens RM66262 | 1 | 563798..601203 | chromosome (NZ_JWLO01000001) |
| T6SS17505 | i3 | Serratia marcescens strain KZ11 | 1 | 580953..617916 | chromosome (NZ_PQGJ01000003) |
| T6SS17504 | i3 | Serratia marcescens strain KZ11 | 0 | 455631..479047 | chromosome (NZ_PQGJ01000003) |
| T6SS00929 | i3 | Serratia marcescens subsp. marcescens Db11 | 1 | 2365124..2402670 | chromosome I (HG326223) |
| T6SS01033 | i3 | Serratia sp. FS14 | 0 | 2830818..2854234 | chromosome (NZ_CP005927) |
| T6SS01224 | i1 | Vibrio alginolyticus 12G01 | 1 | 100481..131233 | chromosome (NZ_CH902589) |
| T6SS00939 | i5 | Vibrio alginolyticus 12G01 | 1 | 1440993..1467006 | chromosome (NZ_CH902589) |
| T6SS01088 | i1 | Vibrio anguillarum strain MHK3 | 1 | 244348..263491 | chromosome (NZ_CP022469) |
| T6SS01061 | i1 | Vibrio cholerae 2740-80 | 1 | 116326..135495 | chromosome (NZ_CP016325) |
| T6SS01168 | i1 | Vibrio cholerae C6706 | 1 | 296505..315674 | chromosome (NZ_CP046845) |
| T6SS01289 | i1 | Vibrio cholerae DL4211 | 1 | 54064..77732 | chromosome (NZ_MOLL01000005) |
| T6SS01290 | i1 | Vibrio cholerae DL4215 | 1 | 74535..98377 | chromosome (NZ_MOLM01000011) |
| T6SS00002 | i1 | Vibrio cholerae O1 biovar El Tor str. N16961 | 1 | 110871..145577 | chromosome II (Vas) (NC_002506) |
| T6SS01112 | i1 | Vibrio cholerae strain A1552 | 1 | 116945..140733 | chromosome (NZ_CP024868) |
| T6SS00790 | i1 | Vibrio cholerae V52 | 1 | 202021..225793 | chromosome (NZ_KQ410497) |
| T6SS01281 | i1 | Vibrio coralliilyticus OCN008 | 1 | 1627613..1656598 | chromosome I (NZ_CP048693) |
| T6SS01280 | i5 | Vibrio coralliilyticus OCN008 | 0 | 2761384..2787511 | chromosome II (NZ_CP048694) |
| T6SS01029 | i5 | Vibrio fluvialis 85003 | 0 | 1341..24915 | chromosome (KY319184) |
| T6SS01028 | i1 | Vibrio fluvialis 85003 | 0 | 2525..21696 | chromosome (KY319183) |
| T6SS00720 | i5 | Vibrio parahaemolyticus BB22OP | 1 | 1048650..1070709 | chromosome II (NC_019971) |
| T6SS00719 | i1 | Vibrio parahaemolyticus BB22OP | 1 | 1464178..1493343 | chromosome I (NC_019955) |
| T6SS00044 | i5 | Vibrio parahaemolyticus RIMD 2210633 | 1 | 1074208..1108114 | chromosome I (NC_004603) |
| T6SS00043 | i1 | Vibrio parahaemolyticus RIMD 2210633 | 1 | 1486402..1526996 | chromosome II (NC_004605) |

| | | | | | |
|---|---|---|---|---|---|
| T6SS01254 | i1 | Vibrio parahaemolyticus strain 12-009A/1335 | 1 | 10921..33508 | chromosome (NZ_MYFF01000494) |
| T6SS01253 | i5 | Vibrio parahaemolyticus strain 12-009A/1335 | 1 | 7384..30649 | chromosome (NZ_MYFF01000478) |
| T6SS01262 | i1 | Vibrio parahaemolyticus strain 12-297/B | 1 | 22753..40086 | chromosome (NZ_MYFG01000475) |
| T6SS01251 | i1 | Vibrio parahaemolyticus strain D4 | 1 | 1500..20954 | chromosome (NZ_NNJE01000049) |
| T6SS01250 | i5 | Vibrio parahaemolyticus strain D4 | 1 | 1..20649 | chromosome (NZ_NNJE01000075) |
| T6SS01247 | i1 | Vibrio parahaemolyticus strain T9109 | 1 | 206703..237535 | chromosome (NZ_JTGR01000038) |
| T6SS00945 | i1 | Vibrio proteolyticus NBRC 13287 | 1 | 17401..46675 | chromosome II (NZ_BATJ01000006) |
| T6SS01236 | i5 | Vibrio vulnificus 106-2A | 1 | 431921..457308 | chromosome (NZ_LMTD01000002) |
| T6SS01235 | i1 | Vibrio vulnificus 106-2A | 1 | 158416..182116 | chromosome (NZ_LMTD01000002) |
| T6SS00287 | i4b | Xanthomonas oryzae pv. oryzae PXO99A | 0 | 3845766..3891788 | chromosome (NC_010717) |
| T6SS01218 | i4b | Xanthomonas oryzae pv. oryzicola strain GX01 | 0 | 3500494..3550582 | chromosome (NZ_CP043403) |
| T6SS00382 | i3 | Xenorhabdus bovienii SS-2004 | 1 | 2053604..2079234 | chromosome (NC_013892) |
| T6SS00381 | i1 | Xenorhabdus bovienii SS-2004 | 1 | 268621..293062 | chromosome (NC_013892) |
| T6SS00017 | i3 | Yersinia pestis CO92 | 1 | 526127..562682 | chromosome (NC_003143) |
| T6SS00225 | i3 | Yersinia pseudotuberculosis IP 31758 | 1 | 3850633..3874248 | chromosome (NC_009708) |
| T6SS00260 | i3 | Yersinia pseudotuberculosis YPIII | 0 | 3905709..3929297 | chromosome (NC_010465) |
| T6SS00258 | i3 | Yersinia pseudotuberculosis YPIII | 0 | 1619849..1653371 | chromosome (NC_010465) |
| T6SS00466 | ii | Francisella tularensis subsp. holarctica LVS | 0 | 102157..124300 | chromosome (NC_007880) |
| T6SS00467 | ii | Francisella tularensis subsp. holarctica LVS | 0 | 1096796..1118939 | chromosome (NC_007880) |
| T6SS00158 | ii | Francisella tularensis subsp. novicida U112 | 0 | 1378294..1404786 | chromosome (NC_008601) |
| T6SS00081 | ii | Francisella tularensis subsp. tularensis SCHU S4 | 0 | 1379925..1405040 | chromosome (NC_006570) |
| T6SS00080 | ii | Francisella tularensis subsp. tularensis SCHU S4 | 0 | 1773260..1798375 | chromosome (NC_006570) |
| T6SS01296 | iii | Bacteroides dorei DSM 17855 | 0 | 7229..28879 | chromosome (NZ_DS995532) |
| T6SS00926 | iii | Bacteroides fragilis 638R | 0 | 2330059..2352459 | chromosome (NC_016776) |
| T6SS00907 | iii | Bacteroides fragilis NCTC 9343 | 0 | 2346972..2370693 | chromosome (NC_003228) |

| | | | | | |
|---|---|---|---|---|---|
| T6SS00908 | iii | Bacteroides fragilis YCH46 | 0 | 2277219..2299620 | chromosome (NC_006347) |
| T6SS00909 | iii | Bacteroides fragilis YCH46 | 0 | 3238391..3255873 | chromosome (NC_006347) |
| T6SS00910 | iii | Flavobacterium johnsoniae UW101 | 0 | 3892562..3923100 | chromosome (NC_009441) |
| T6SS01206 | iv | Candidatus Amoebophilus asiaticus 5a2 | 0 | 713229..728788 | chromosome (NC_010830) |
| T6SS01207 | iv | Candidatus Amoebophilus asiaticus 5a2 | 0 | 1380571..1401622 | chromosome (NC_010830) |
