## Supplementary Data 3 for "LLPS condensates of Fha initiate the inside-out assembly of the type VI secretion system"

>Acidovorax_citrulli_AAC00-1

MNTTTSLDAVQGTQTLEGSEFTSLLQKEFKPKTDQAREAVESAVQTLAQQALAASATLSDDAYQTVQAIIAEIDRKLSEQINLIIHHSDFQQLEGAWRGLHHLV

HNTEVDELLKIRVMNISKKELHRNMRRFKGVGWDQSPIFKKVYEEEYGQFGGEPFGCLVGDYYFDHSPPDVELLGEAKVSAAAHCPFIAGASPAMMQMDSW

QELANPRDLTKIFTNTEYAAWRSLRDSDDSKYIGLAMPRFLARLPYGAKTNPVDEFDFEEETAGGDHSKYAWANSAYAMAVNINRSFKYYGWCTSIRGVESGG

AVDNLPAHTFPTDDGGVDMKCPTEIAISDRREAELAKNGFMPLVHRKNSDVAAFIGAQSLHKPAEYYDPDATANANLSARLPYMFACCRFAHYLKCIVRDKIG

SFKERSDMERWLNDWIMNYVDGDPANSSQETKARKPLAAAEVQVAEVEGNPGYYTSKFFLRPHYQLEGLTVSLRLVSKLPSVKQAGN

>Acinetobacter_baumannii_AB307-0294

MNNTQSAAMPLVENEEVNLLDSIVEQSRIARNEEEHSRAKSLIGELAKEVMAGTITVSENMTLSIDKRIAEIDALISKQLSQIMHNEQFQKIESTWRGLYYFCQE

TPSNPLIKIRMLNTTKKELVKDFQGATDFDQSTLFKKIYEEEYGSFGGAPYSALIGDFEFDRTPSDMYLLEQISHVAAAAHAPFISAASPSILGLESFTDIDRPRDV

SKIFETAEYVQWRSFRDSEDSRYVALTLPHVLGRLPYHPKEGTATEGFNFIEDVSGENHNEYLWMNAAYAFGTRLTNAFDMHGWCAAIRGVEGGGLVEGLPV

HTFKTQDGEVVFKCPTEIAITDRREKELSDLGFIPLVHCKNTDYAAFFGAQSTQKPKKYDNDTANANSALSSQIQYIMAVSRIAHYLKAMMRDKVGSFASAGN

VEAFLNEWLSQYVLLDDGASQEAKAQYPLREASVKVVEDPAQPGHYKSVVFLRPHFQLDELSVSLRLVTELPQSSN

>Acinetobacter_baumannii_AbCAN2

MNNTQSAAMPLVENEEVNLLDSIVEQSRIARNEEEHSRAKSLIGELAKEVMAGTITVSENMTLSIDKRIAEIDALISKQLSQIMHNEQFQKIESTWRGLYYFCQE

TPSNPLIKIRMLNTTKKELVKDFQGATDFDQSTLFKKIYEEEYGSFGGAPYSALIGDFEFDRTPSDMYLLEQISHVAAAAHAPFISAASPSILGLESFTDIDRPRDV

SKIFETAEYVQWRSFRDSEDSRYVALTLPHVLGRLPYHPKEGAATEGFNFIEDVSGENHNEYLWMNAAYAFGTRLTNAFDMHGWCAAIRGVEGGGLVEGLPV

HTFKTQDGEVVFKCPTEIAITDRREKELSDLGFIPLVHCKNTDYAAFFGAQSTQKPKKYDNDTANANSALSSQIQYIMAVSRIAHYLKAMMRDKVGSFTSAGN

VEAFLNEWLSQYVLLDDGASQEAKAQYPLREASVKVVEDPAQPGHYKSVVFLRPHFQLDELSVSLRLVTELPQSSN

>Acinetobacter_baumannii_ATCC_17978

MNNTQSAAMPLVENEEVNLLDSIVEQSRIARNEEEHSRAKSLIGELAKEVMAGTITVSENMTLSIDKRIAEIDALISKQLSQIMHNEQFQKIESTWRGLYYFCQE

TPSNPLIKIRMLNTTKKELVKDFQGATDFDQSTLFKKIYEEEYGSFGGAPYSALIGDFEFDRTPSDMYLLEQISHVSAAAHAPFISAASPSILGLESFTDIDRPRDV

SKIFETAEYVQWRSFRDSEDSRYVALTLPHVLGRLPYHPKEGTATEGFNFIEDVSGENHNEYLWMNAAYAFGTRLTNAFDMHGWCAAIRGVEGGGLVEGLPV

HTFKTQDGEVVFKCPTEIAITDRREKELSDLGFIPLVHCKNTDYAAFFGAQSTQKPKKYDNDTANANSALSSQIQYIMAVSRIAHYLKAMMRDKVGSFASAGN

VEAFLNEWLSQYVLLDDGASQEAKAQYPLREASVKVVEDPAQPGHYKSVVFLRPHFQLDELSVSLRLVTELPQSSN

>Acinetobacter_baumannii_DSM_30011

MNNTQSAAMPLVENEEVNLLDSIVEQSRIARNEEEHSRAKSLIGELAKEVMAGTITVSENMTLSIDKRIAEIDALISKQLSQIMHNEQFQKIESTWRGLYYFCQE

TPSNPLIKIRMLNTTKKELVKDFQGATDFDQSTLFKKIYEEEYGSFGGAPYSALIGDFEFDRTPSDMYLLEQISHVAAAAHAPFISAASPSILGLESFTDIDRPRDV

SKIFETAEYVQWRSFRDSEDSRYVALTLPHVLGRLPYHPKEGTATEGFNFIEDVSGENHNEYLWMNAAYAFGTRLTNAFDMHGWCAAIRGVEGGGLVEGLPV

HTFKTQDGEVVFKCPTEIAITDRREKELSDLGFIPLVHCKNTDYAAFFGAQSTQKPKKYDNDTANANSALSSQIQYIMAVSRIAHYLKAMMRDKVGSFASAGN

VEAFLNEWLSQYVLLDDGASQEAKAQYPLREASVKVVEDPAQPGHYKSVVFLRPHFQLDELSVSLRLVTELPQSSN

>Acinetobacter_baumannii_strain_UPAB1

MNNTQSAAMPLVENEEVNLLDSIVEQSRIARNEEEHSRAKSLIGELAKEVMAGTITVSENMTLSIDKRIAEIDALISKQLSQIMHNEQFQKIESTWRGLYYFCQE

TPSNPLIKIRMLNTTKKELVKDFQGATDFDQSTLFKKIYEEEYGSFGGAPYSALIGDFEFDRTPSDMYLLEQISHVAAAAHAPFISAASPSILGLESFTDIDRPRDV

SKIFETAEYVQWRSFRDSEDSRYVALTLPHVLGRLPYHPKEGTATEGFNFIEDVSGENHNEYLWMNAAYAFGTRLTNAFDMHGWCAAIRGVEGGGLVEGLPV

HTFKTQDGEVVFKCPTEIAITDRREKELSDLGFIPLVHCKNTDYAAFFGAQSTQKPKKYDNDTANANSALSSQIQYIMAVSRIAHYLKAMMRDKVGSFASAGN

VEAFLNEWLSQYVLLDDGASQEAKAQYPLREASVKVVEDPAQPGHYKSVVFLRPHFQLDELSVSLRLVTELPQSSN

>Acinetobacter_sp.ADP1

MNNPQLANNTALEPENTTLLDSIVEQSRIARNDEEHVRAKNLIGELAKEVMAGTITVSENMTLSIDKRIAEIDALISAQLSQIMHHEKFQKIESTWRGLYYFCQE

TPSNPLIKIRMLNTTKKELVKDFQGATDFDQSTLFKKIYEEEYGSFGGAPYSALIGDFEFDRTPSDMYLLEQISHVAAAAHAPFISAADPSMFGLESYTDIDRPRD
VSKIFETAEYVQWRSFRDSEDARYVALTMPRVLGRLPYHPKEGTATEGFNFIEDVTGQDHNEYLWMNAAYAFGTRLTNAFDLHGWCAAIRGVEGGGLVEGLP
VHTFKTHDGEVAFKCPTEIAITDRREKELSDLGFIPLVHCKNTDYAAFFGAQSTQKPKKYDSDTANANSALSSQIQYIMAVSRIAHYLKAMMRDKVGSFASAG
NVEAFLNEWLSQYVLLDDGASQEAKAQYPLREASVKVVEDPAQPGHYKSVVFLRPHFQLDELSVSLRLVTELPQSSN

>Aeromonas_dhakensis_strain_SSU
MSLVEEQVQAGASAASSSLLDEIMAQARITPVDEGYSVAKQGVAALIANILDSGTTSEPVNKALVDSMIVELDKKLSKQMDVILHAKELQEMESSWRSLKLLV
DRTDFRENIKIQVLHATKEELLEDFEFSPEITQSGLYKHVYSTGYGQFGGQPVGAVIGDYAFTHSSPDIKLMQYVSAVGAMAHAPFISSVAPAFFGVDSFTDLPSI
KDLKSVFEGPAYTKWRSLRESEDARYLGLTAPRFLARLPYDPTENPIKGFNYQEDISSDHDHYLWGNTAYLMGTSLTDSFAKYRWCPNIIGPQSGGAIHDLPVH
VYEAMGQLQAKIPTEVLITDRREYELSEEGFITLTMRKDSDNAAFFSANSVQKPKVFPNTKEGKEAETNYKLGTQLPYMFIINRLAHYIKVLQREQIGSWKERQ
DLERELNGWIKQYVADQENPPADVRSRRPLRAAQIKVLDVEGEPGWYQVAMAVRPHFKYMGASFELSLVGRLDKE

>Aeromonas_hydrophila_J-1
MSLVEEQVQAGASAASSSLLDEIMAQARITPVDEGYNVAKQGVAALIANILDSGTTSDPVNKALVDSMIVELDKKLSKQMDVILHAKELQEMESSWRSLKLLV
DRTDFRENIKIQVLHATKEELLEDFEFSPEITQSGLYKHVYSSGYGQFGGQPVGAVIGDYAFTHSAPDIKLMQYVSAVGAMAHAPFISSVAPSFFGVDSFTDLSSI
KDLKSVFEGPAYTKWRALRESEDARYLGLTAPRFLARLPYDPTENPIKGFNYQEDISSDHDHYLWGNTAYLMGTSLTDSFAKYRWCPNIIGPQSGGAIHDLPVH
VYEAMGQLQAKIPTEVLITDRREYELSEEGFITLTMRKDSDNAAFFSANSVQKPKVFPNTKEGREAETNYKLGTQLPYMFIINRLAHYIKVLQREQIGSWKERQ
DLERELNSWIRQFVADQENPPAEVRSRRPLRAAQIKVLDVEGEPGWYQVAMAVRPHFKYMGASFELSLVGRLDKE

>Aeromonas_hydrophila_NJ-35
MSLVEEQVQAGASAASSSLLDEIMAQARITPVDEGYNVAKQGVAALIANILDSGTTSDPVNKALVDSMIVELDKKLSKQMDVILHAKELQEMESSWRSLKLLV
DRTDFRENIKIQVLHATKEELLEDFEFSPEITQSGLYKHVYSSGYGQFGGQPVGAVIGDYAFTHSAPDIKLMQYVSAVGAMAHAPFISSVAPSFFGVDSFTDLSSI
KDLKSVFEGPAYTKWRALRESEDARYLGLTAPRFLARLPYDPTENPIKGFNYQEDISSDHDHYLWGNTAYLMGTSLTDSFAKYRWCPNIIGPQSGGAIHDLPVH
VYEAMGQLQAKIPTEVLITDRREYELSEEGFITLTMRKDSDNAAFFSANSVQKPKVFPNTKEGREAETNYKLGTQLPYMFIINRLAHYIKVLQREQIGSWKERQ
DLERELNSWIRQFVADQENPPAEVRSRRPLRAAQIKVLDVEGEPGWYQVAMAVRPHFKYMGASFELSLVGRLDKE

>Aeromonas_hydrophila_strain_NF1
MSLVEEQVQAGASAASSSLLDEIMAQARITPVDEGYSVAKQGVAALIANILDSGTTSEPVNKALVDSMIVELDKKLSKQMDVILHAKELQEMESSWRSLKLLV
DRTDFRENIKIQVLHATKEELLEDFEFSPEITQSGLYKHVYSTGYGQFGGQPVGAVIGDYAFTHSSPDIKLMQYVSAVGAMAHAPFISSVAPAFFGVDSFTDLPSI
KDLKSVFEGPAYTKWRSLRESEDARYLGLTAPRFLARLPYDPTENPIKGFNYQEDISSDHDHYLWGNTAYLMGTSLTDSFAKYRWCPNIIGPQSGGAIHDLPVH
VYEAMGQLQAKIPTEVLITDRREYELSEEGFITLTMRKDSDNAAFFSANSVQKPKVFPNTKEGKEAETNYKLGTQLPYMFIINRLAHYIKVLQREQIGSWKERQ
DLERELNGWIKQYVADQENPPADVRSRRPLRAAQIKVLDVEGEPGWYQVAMAVRPHFKYMGASFELSLVGRLDKE

>Aeromonas_hydrophila_subsp._hydrophila_ATCC_7966
MSLVEEQVQAGASAASSSLLDEIMAQARITPVDEGYSVAKQGVAALIANILDSGTTSDPVNKALVDSMIVELDKKLSKQMDVILHAKELQEMESSWRSLKLLV
DRTDFRENIKIQVLHATKEELLEDFEFSPEITQSGLYKHVYSSGYGQFGGQPVGAVIGDYAFTHSAPDIKLMQYVSAVGAMAHAPFISSVAPSFFGVDSFTDLSSI
KDLKSVFEGPAYTKWRALRESEDARYLGLTAPRFLARLPYDPTENPIKGFNYQEDISSDHDHYLWGNTAYLMGTSLTDSFAKYRWCPNIIGPQSGGAIHDLPVH
VYEAMGQLQAKIPTEVLITDRREYELSEEGFITLTMRKDSDNAAFFSANSVQKPKVFPNTKEGREAETNYKLGTQLPYMFIINRLAHYIKVLQREQIGSWKERQ
DLERELNSWIRQFVADQENPPAEVRSRRPLRAAQIKVLDVEGEPGWYQVAMAVRPHFKYMGASFELSLVGRLDKE

>Aeromonas_veronii_strain_TH0426
MSLVEEQVQAGAGAASSSSLLDEIMAQARITPVDEGYSVAKQGVAALIANILDSGTTSEPVNKALVDSMIVELDKKLSKQMDVILHAKELQEMESSWRSLKLL
VDRTDFRENIKIQVLHATKEELLEDFEFSPEITQSGLYKHVYSTGYGQFGGQPIGAVIGDYAFTHSSPDIKLMQYVSAVGAMAHAPFISSVAPAFFGVDSFTDLPSI
KDLKSVFEGPAYTKWRSLRESEDSRYLGLTAPRFLARLPYDPTENPIKGFNYKEDISSDHDHYLWGNTAYLMGTSLTDSFAKYRWCPNIIGPQSGGAIHDLPVH
VYEAMGQLQAKIPTEVLITDRREYELSEEGFITLTMRKDSDNAAFFSANSVQKPKVFPNTKEGKEAETNYKLGTQLPYMFIINRLAHYIKVLQREQIGSWKERQ
DLERELNGWIKQYVADQENPPADVRSRRPLRAAQIKVLDVEGEPGWYQVAMAVRPHFKYMGASFELSLVGRLDKE

>Agrobacterium_fabrum_str._C58

MSAESLLKSEAQATTAENQGLLSKVVSATRQTEPDRAQNLLRTLTDQALKGTVKYDRNLTVTLNHAIAELDRTISEQLAAIMQAPEFAKLEGTWRGLNYLVKN
SETSVNLKIRVMNAGKRELARDLEKAVEFDQSRLFKAIYEDEFGTPGGEPLGAIIGDYEFGNSFDDVQLLQGVSSIAAAAFAPFISAASPHMFGFEDYRDLARPR
DLEKIFDTVEYAKWRSFRESDDSRFVTLALPRVLARMPYGPKTNPVDDFAYDETRGAVNGDLQHDEYCWMNAAYVMGTKLTEAFAKSGWCTAIRGAENGG
RVENLPMHVFSSDDGDLDLKCPTEVGITDRRDAELGKLGFLPLCHYKNTDYAVFFGAQTAHKPKLYDKPEATANAAVSARLPYIMATSRFAHYLKVMGRDKI
GSFMEASDCEVWLNRWIANYVNANDEAGEESRAKYPLRDAKVTVQEVPGKPGAYNAVAWMRPWLQMEELTTSLRMVARIPSKN

>Agrobacterium_fabrum_strain_12D13

MSAESLLKSEAQATTAEDQGLLSKVVSATRQTEPDRAQNLLRTLTDQALKGTVKYDRNLTVTLNHAIAELDRTISEQLAAIMQAPEFAKLEGTWRGLNYLVKN
SETSVNLKIRVMNAGKRELARDLEKAVEFDQSRLFKAIYEDEFGTPGGEPLGAIIGDYEFDNSFDDVQLLQGVSSIAAAAFAPFISAASPHMFGFEDYRDLARPR
DLEKIFDTVEYAKWRSFRESDDSRFVTLALPRVLARMPYGPKTNPVDDFAYDETRGAVNGDLQHDEYCWMNAAYVMGTKLTEAFAKSGWCTAIRGAENGG
RVENLPMHVFSSDDGDLDLKCPTEVGITDRRDAELGKLGFLPLCHYKNTDYAVFFGAQTAHKPKLYDKPEATANAAVSARLPYIMATSRFAHYLKVMGRDKI
GSFMEASDCEVWLNRWIANYVNANDEAGEESRAKYPLRDAKVTVQEVPGKPGAYNAVAWMRPWLQMEELTTSLRMVARIPSKN

>Agrobacterium_fabrum_strain_1D132

MSAESLLKSEAQATTAENQGLLSKVVSATRQTEPDRAQNLLRTLTDQALKGTVKYDRNLTVTLNHAIAELDRTISEQLAAIMQAPEFAKLEGTWRGLNYLVKN
SETSVNLKIRVMNAGKRELARDLEKAVEFDQSRLFKAIYEDEFGTPGGEPLGAIIGDYEFGNSFDDVQLLQGVSSIAAAAFAPFISAASPHMFGFEDYRDLARPR
DLEKIFDTVEYAKWRSFRESDDSRFVTLALPRVLARMPYGPKTNPVDDFAYDETRGAVNGDLQHDEYCWMNAAYVMGTKLTEAFAKSGWCTAIRGAENGG
RVENLPMHVFSSDDGDLDLKCPTEVGITDRRDAELGKLGFLPLCHYKNTDYAVFFGAQTAHKPKLYDKPEATANAAVSARLPYIMATSRFAHYLKVMGRDKI
GSFMEASDCEVWLNRWIANYVNANDEAGEESRAKYPLRDAKVTVQEVPGKPGAYNAVAWMRPWLQMEELTTSLRMVARIPSKN

>Agrobacterium_tumefaciens_strain_12D1

MSAESLLKNEAQATTAEDQGLLSKVVAATRQTEPDRAQNLLRTLTDQALKGTVKYDRNLTVTLNHAIAELDRTISEQLAAIMQAPEFAKLEGTWRGLNYLVK
NSETSVNLKIRVMNSGKRELARDLEKAVEFDQSRLFKAIYEDEFGTPGGEPLGAIIGDYEFDNSFDDVQLLQGVSSIAAAAFAPFISAASPHMFGFEDYRDLARP
RDLEKIFDTVEYAKWRSFRESDDSRFVTLALPRVLARMPYGPKTNPIDDFAYDETKGAVNGDLQHDEYCWMNAAYVMGTKLTDAFAKSGWCTAIRGAENGG
RVENLPMHVFSSDDGDLDLKCPTEVGITDRRDAELGKLGFLPLCHYKNTDYAVFFGAQTAHKPKLYDKPEATANAAVSARLPYIMATSRFAHYLKVMGRDKI
GSFMEASDCEVWLNRWIANYVNANDEAGEESRAKYPLRDAKVTVQEVPGKPGAYNAVAWMRPWLQMEELTTSLRMVARIPSKN

>Agrobacterium_tumefaciens_strain_15955

MSAESLLKNEAQAMTAEDQGLLSKVVAATRQTEPDRAQNLLRTLTDQALKGTVKYDRNLTVTLNHAIAELDRTISEQLAAIMQAPEFAKLEGTWRGLNYLVK
NSETSVNLKIRVMNAGKRELARDLEKAVEFDQSRLFKAIYEDEFGTPGGEPLGAIIGDYEFDNSFDDVQLLQGVSSIAAAAFAPFISAASPHMFGFEDYRDLARP
RDLEKIFDTVEYAKWRSFRESDDSRFVTLALPRVLARMPYGPKTNPIDDFAYDETKGAVNGDLEHDEYCWMNAAYVMGTKLTDAFAKSGWCTAIRGAENGG
KVENLPMHVFSSDDGDLDLKCPTEVGITDRRDAELGKLGFLPLCHYKNTDYAVFFGAQTAHKPKLYDKPEATANAAVSARLPYIMATSRFAHYLKVMGRDKI
GSFMEASDCEVWLNRWIANYVNANDEAGEESRAKYPLRDAKVTVQEVPGKPGAYNAVAWMRPWLQMEELTTSLRMVARIPSKN

>Agrobacterium_tumefaciens_strain_1D1108

MSAESLLKNEAQATTAEDQGLLSKVVAATRQTEPDRAQNLLRTLTDQALKGTVKYDRNLTVTLNHAIAELDRTISEQLAAIMQAPEFAKLEGTWRGLNYLVK
NSETSVNLKIRVMNAGKRELARDLEKAVEFDQSRLFKAIYEDEFGTPGGEPLGAIIGDYEFDNSFDDVQLLQGVSSIAAAAFAPFISAASPHMFGFEDYRDLARP
RDLEKIFDTVEYAKWRSFRESDDSRFVTLALPRVLARMPYGPKTNPIDDFAYDETKGAVNGDLEHDEYCWMNAAYVMGTKLTDAFAKSGWCTAIRGAENGG
KVENLPMHVFSSDDGDLDLKCPTEVGITDRRDAELGKLGFLPLCHYKNTDYAVFFGAQTAHKPKLYDKPEATANAAVSARLPYIMATSRFAHYLKVMGRDKI
GSFMEASDCEVWLNRWIANYVNANDEAGEESRAKYPLRDAKVTVQEVPGKPGAYNAVAWMRPWLQMEELTTSLRMVARIPSKN

>Agrobacterium_tumefaciens_strain_1D1460

MSAESLLKNEAQATTAEDQGLLSKVVAATRQTEPDRAQNLLRTLADQALKGTVKYDRNLTVTLNHAIAELDRTISEQLAAIMQAPEFAKLEGTWRGLNYLVK
NSETSVNLKIRVMNSGKRELARDLEKAVEFDQSRLFKAIYEDEFGTPGGEPLGAIIGDYEFDNSFDDVQLLQGVSSIAAAAFAPFISAASPHMFGFEDYRDLARP

RDLEKIFDTVEYAKWRSFRESDDSRFVTLALPRVLARMPYGPKTNPIDDFAYDETKGAVNGDLQHDEYCWMNAAYVMGTKLTDAFAKSGWCTAIRGAENGG
RVENLPMHVFSSDDGDLDLKCPTEVGITDRRDAELGKLGFLPLCHYKNTDYAVFFGAQTAHKPKLYDKPEATANAAVSARLPYIMATSRFAHYLKVMGRDKI
GSFMEASDCEVWLNRWIANYVNANDEAGEESRAKYPLRDAKVTVQEVPGKPGAYNAVAWMRPWLQMEELTTSLRMVARIPSKN

>Agrobacterium_tumefaciens_strain_1D1609
MSAESLLKNEAQATTAEDQGLLSKVVAATRQTEPDRAQNLLRTLTDQALKGTVKYDRNLTVTLNHAIAELDRTISEQLAAIMQAPEFAKLEGTWRGLNYLVK
NSETSVNLKIRVMNAGKRELARDLEKAVEFDQSRLFKAIYEDEFGTPGGEPLGAIIGDYEFDNSFDDVQLLQGVSSIAAAAFAPFISAASPHMFGFEDYRDLARP
RDLEKIFDTVEYAKWRSFRESDDSRFVTLALPRVLARMPYGPKTNPIDDFAYDETRGAVNGDLQHDEYCWMNAAYVMGTKLTDAFAKSGWCTAIRGAENGG
RVENLPMHVFSSDDGDLDLKCPTEVGITDRRDAELGKLGFLPLCHYKNTDYAVFFGAQTAHKPKLYDKPEATANAAVSARLPYIMATSRFAHYLKVMGRDKI
GSFMEASDCEVWLNRWIANYVNANDEAGEESRAKYPLRDAKVTVQEVPGKPGAYNAVAWMRPWLQMEELTTSLRMVARIPSKN

>Agrobacterium_tumefaciens_strain_A6
MSAESLLKNEAQAMTAEDQGLLSKVVAATRQTEPDRAQNLLRTLTDQALKGTVKYDRNLTVTLNHAIAELDRTISEQLAAIMQAPEFAKLEGTWRGLNYLVK
NSETSVNLKIRVMNAGKRELARDLEKAVEFDQSRLFKAIYEDEFGTPGGEPLGAIIGDYEFDNSFDDVQLLQGVSSIAAAAFAPFISAASPHMFGFEDYRDLARP
RDLEKIFDTVEYAKWRSFRESDDSRFVTLALPRVLARMPYGPKTNPIDDFAYDETKGAVNGDLEHDEYCWMNAAYVMGTKLTDAFAKSGWCTAIRGAENGG
KVENLPMHVFSSDDGDLDLKCPTEVGITDRRDAELGKLGFLPLCHYKNTDYAVFFGAQTAHKPKLYDKPEATANAAVSARLPYIMATSRFAHYLKVMGRDKI
GSFMEASDCEVWLNRWIANYVNANDEAGEESRAKYPLRDAKVTVQEVPGKPGAYNAVAWMRPWLQMEELTTSLRMVARIPSKN

>Aliivibrio_fischeri_ES114
MSDTSTAQELDNQVTVGLLDQIVAQTNLTPEDETYGIAKRGVSAFIEELLKPQNQEEPVKKALIDKMITEIDQRLSKQVDEILHHSSFQQLESAWRGLKLLVDRT
DFRENIKIEVINVSKTDLLEDFEDASEITQSGLYKHVYTNEFGTFGGQPVGAIIGNYDFGPSAPDIKNLQNLASVAAMSHAPFISAAGPKFFGLESFEGLPDLKDL
NDHFESPQYAKWQSFREQEDSRYVGLTLPRFLLRQPYDPEENPVKAFNYHENVSATHEDYLWGNTAFTMATRITESFANFRWCPNIIGPQSGGAVEDLPLHHFE
SMGDIETKIPTEVLVSDRREYELAEEGFIALTMRKGSDNAAFFSANSVQKAKFFGNTEEGKNAELNFRLGTQLPYLFIISRLAHYIKVLQREQIGSWKERTDLEQ
ELNTWIRQYIADQENPPAEVRSRRPLRAANIEVMDVEGNPGWYKVNLSVRPHFKYMGSDFTLSLVGKLDQE

>Aliivibrio_fischeri_strain_ES401
MTAEAQAPEQEAALVESGSLLDSILNETRLKPSDEGFDVAKRGVEAFIGELLSTNTSDKVDQSLVDLMISEIDQKLSKQVDAILHNKEVQAIESTWRGLKYLVD
HTDFRENIQIELISAKKDEVLDDFEDAPEVVKSGLYKQIYTREYGQFGGKPVGAVICDFQMSASSPDIKLMEYMANVGAMSHAPFITSASAKFFGLDSYEELPN
LKDLKSVFEGPQYTKWRGFREHEDARYVGLCTSRMMLRTPYSVEDNPIKAFDYNEHVKDGHDNYLWGNSAYAMASKISESFAKYRWCPNIIGPQSGGTVSD
LPVYNYESMGQIETKIPTEILISDRREYELAEEGFIALTMRKGSDNAAFFSANSVQKAKVFPNTPEGQQAEMNYKLGIQLPYMFIINRLAHYIKVLQREQIGSWK
ERSDLEIELNKWIRQYVSDQENPPAEVRGRRPLRAAKVEVSEVEGDPGWYKVSMSVRPHFKYMGASFDLSLVGKLDQN

>Aliivibrio_fischeri_strain_FQ-A001_T6SS-1
MSDTSTAQELDNQVTVGLLDQIVAQTNLTPEDETYGIAKRGVSAFIEELLKPQNQEEPVKKALIDKMITEIDQRLSKQVDEILHHSSFQQLESAWRGLKLLVDRT
DFRENIKIEVINVSKTDLLEDFEDASEITQSGLYKHVYTNEFGTFGGQPVGAIIGNYDFGPSAPDIKNLQNLASVAAMSHAPFISAAGPKFFGLESFEGLPDLKDL
NDHFESPQYAKWQSFREQEDSRYVGLTLPRFLLRQPYDPEENPVKAFNYHENVSATHEDYLWGNTAFTMATRITESFANFRWCPNIIGPQSGGAVEDLPLHHFE
SMGDIETKIPTEVLVSDRREYELAEEGFIALTMRKGSDNAAFFSANSVQKAKFFGNTEEGKNAELNFRLGTQLPYLFIISRLAHYIKVLQREQIGSWKERTDLEQ
ELNTWIRQYIADQENPPAEVRSRRPLRAANIEVMDVEGNPGWYKVNLSVRPHFKYMGSDFTLSLVGKLDQE

>Aliivibrio_fischeri_strain_FQ-A001_T6SS-2
MTAEAQAPEQEAALVESGSLLDSILNETRLKPSDEGFDVAKRGVEAFIGELLSTNTSDKVDQSLVDLMISEIDQKLSKQVDAILHNKEVQAIESTWRGLKYLVD
HTDFRENIQIELISAKKDEVLDDFEDAPEVVKSGLYKQIYTREYGQFGGKPVGAVICDFQMSASSPDIKLMEYMANVGAMSHAPFITSASAKFFGLDSYEELPN
LKDLKSVFEGPQYTKWRGFREHEDARYVGLCTSRMMLRTPYSVEDNPIKAFDYNEHVKDGHDNYLWGNSAYAMASKISESFAKYRWCPNIIGPQSGGTVSD
LPVYNYESMGQIETKIPTEILISDRREYELAEEGFIALTMRKGSDNAAFFSANSVQKAKVFPNTPEGQQAEMNYKLGIQLPYMFIINRLAHYIKVLQREQIGSWK
ERSDLEIELNKWIRQYVSDQENPPAEVRGRRPLRAAKVEVSEVEGDPGWYKVSMSVRPHFKYMGASFDLSLVGKLDQN

>Azoarcus_olearius

MADTQAQNEAQAGGLAIEGGDFESLLRQEFKPKTDEARSAVENAVRTLAEQALSQTTLIGTDVVKSIEAIIAALDQKLTEQVNQIIHHEDFQKLEGAWRGLHY
LVNNTETDEMLKIRVFNITKNELGKTLKRYKGTAWDQSPIFKRVYEEEYGQFGGEPFGCIVGDYHFDQSPPDVELLGEMAKVSAAAHTPFITGASPTIMQMDS
WQELANPRDLTKIFSTPEYAAWRSLRESDDAKYIGLAMPRFLSRVPYGARTNPVEEFDFEEDTGAADHSKYTWANAAYAMAVNINRSFKEYGWCSRIRGIESG
GAVENLPVHSFPTDDGGVDMKCPTEIAISDRREAELAKNGFMPLVHKKNSDFAAFIGAQSLHKPAEYDDPDATANANLGARLPYLFATCRFAHYLKCIVRDKI
GSFKERDDMQRWLQDWIMNYVDGDPSHSSEATKARRPLAAAEVVVEEVEGNPGFYSSKFFLRPHYQLEGLTVSLRLVSKLPSAKGG

>Bordetella_bronchiseptica_RB50

MTAVARRERAQAESSLAADDLSALLKKEFKPKTEQAREAVEHAVRTLAHQALENSGLGLSSDAYRTIQAIIAEIDRKLSEQINLVMHHPDFQQLEGAWRGLHYL
VTNTETDELLKIRFMCLSKNELGRTLKRYKGVGWDQSPIFKRVYEEEYGQFGGEPIGCLVGDYSFDHSPPDVELLGEIARISAAAHCPFIAGASPNVMQMDSW
QELSNPRDLTKIFTNTEYAAWRSLRESEDARYVGLAMPRFLARLPYGARTNPVDEFDFEEETDGANHDRYTWANSAYAMAANINRSFKLYGWCTSIRGVESG
GAVENLPCHTFPTDDGGVDMKCPTEIAISDRREAELAKNGFMPLVHRKNSDFAAFIGAQSLQRPQEYHDADASANAKLAARLPYLFACCRFAHYLKCIVRDKI
GSFRERSDMERWLNDWIMNYVDGDPANSSQETKARKPLAAAEVQVTEIEDNPGYYAAKFFLRPHYQLEGLTVSLRLVSKLPSLKQNEG

>Bradyrhizobium_japonicum_strain_J5

MAKEAVHKESTTQVKTVEADEFATLLKQSFKPRTERAATEVDNAVSTLVQEALKDSSVIKSDVLDTIEEMIARIDQKLTAQMNEIIHAPEFQAIESTWRGLHYLV
FNSETDANLKIRVMNVSKTELYRNLRLYPDARWDQSPLFKQIYEYEFGQLGGEPFGCLIGDYYFSHLPTDVQLLRDLSKIAGAAHAPFFSGAEPTLMGMDSWT
ELSNPRDIGKVFDTPEYAAWKGLRDQDDSRYLGLCMPRALGRLPYGAKSEPVEEFAFEEETDGHTGDKYGWINAAYAMAVNINRAFKEFGWCTRIRGVQSGG
EVINLPTHTFPTDDGGVDLKCPTEIAISDRREHELAKAGLIPLIHRKNTDKAAFIGAQSLYKPKKYFGEKGVDATASDNLSSRLPYMFAVSRFAHYLKCMVRDKI
GSMKEKDELTVWLQTWINEYVDANPALSSEAQKARKPLAAAKVEVIANEENPGYYNARFFLRPHFQLEAMDVGLSLVSRLPGPSS

>Burkholderia_cenocepacia_AU_1054

MNQQTAAAQASGAEYAAGTSLLDDIVEKSKVAKSDSEHARAKDLIGELVHQVLDGTVIVSDNLSATIDARVAELDRLISTQLSAVMHAPEFQRLESTWRGMDY
LVKESNTGQTIKIKALHAPKRDLVRDFKGASEFDQSALFKKVYEEEFGTFGGSPFGALIGDYEISRQPEDMYFIEQMSHVAAAAHAPFIASASPELLGLESFSDL
GKPRDLGKVFDTVEYAKWKSFRDAEDSRYVGLTLPRFLGRLPFNPKDGQTAENFNFVEDVDGTDHDKYLWCNAAWAFAARLTAAFDDFGWCAAIRGVEGG
GLVEDLPTHTFKTDDGEVALKCPTEIAITDRREKELSDLGFIPLVHCKNSDYAAFFAAQSVQKPKKYSTDSANANAVLSAQLQYIFSVSRVAHYLKAMMRDKIG
SFASAQNVETFLNRWISQYVLLDDNASQEQKAQFPLREASIQVSEIPGKPGSYRSVAFLRPHFQLDELSISLRLVADLPKPANS

>Burkholderia_cenocepacia_H111_T6SS-2

MNQQTAAAQASGAEYAAGTSLLDDIVEKSKVAKSDSEHARAKDLIGELVHQVLDGTVIVSDNLSATIDARVAELDRLISTQLSAVMHAPEFQRLESTWRGLDY
LVKESNTGQTIKIKALHAPKRDLVRDFKGASEFDQSALFKKVYEEEFGTFGGSPFGALIGDYEISRQPEDMYFIEQMSHVAAAAHAPFIASASPELLGLESFSDL
GKPRDLGKVFDTVEYAKWKSFRDAEDSRYVGLTLPRFLGRLPFNPKDGQTAENFNFVEDVDGTDHDKYLWCNAAWAFAARLTAAFDDFGWCAAIRGVEGG
GLVEDLPTHTFKTDDGEVALKCPTEIAITDRREKELSDLGFIPLVHCKNSDYAAFFAAQSVQKPKKYSTDSANANAVLSAQLQYIFSVSRVAHYLKAMMRDKIG
SFASAQNVETFLNRWISQYVLLDDNASQEQKAQFPLREASIQVSEIPGKPGSYRSVAFLRPHFQLDELSISLRLVADLPKPANS

>Burkholderia_cenocepacia_H111_T6SS-1

MNQNESRQRASETIALENDSVYAALCNKINLKPVAEARPLEAFRDNDALSEASADERVARGMDALLNLIAKESKPVDRLDKSLLDFYIGQLDRQISRQLDAVM
HAPDFQALEGRWRGLKMLVSRTDFRKNARIEVLDVSKEALQRDFEDSPELIQSGLYRLTYIEEYDTPGGQPISAMISDFEFANSPMDVALLRNISKVAAAAHMP
FIGSVGAAFFGKKSMEEVASIQDIGNYFDRAEYIKWKSFRDTDDARYVGLTMPRVLGRLPYGKDTTPVRAFNYEEAVKGPDHDKYLWVNASFAFAANMVRSF
VSNGWCVQIRGPQAGGKVEDLPVHLYDLGTGVQPKIPTEVLIPETREFEFANLGFIPLSFYKNHDFACFFSASSTQKPALYETKEATANSRINARLPYIFLLSRIAH
YLKLIQRENIGTTKDRRLLELELNNWIKGLVTEMKDPGDELQASHPLREAKVTVEDIEDNPGFFRIKLFIIPHFQVEGMDIGLSLVSQMPKAKG

>Burkholderia_cenocepacia_J2315

MNQQTAAAQASGAEYAAGTSLLDDIVEKSKVAKSDSEHARAKDLIGELVHQVLDGTVIVSDNLSATIDARVAELDRLISTQLSAVMHAPEFQRLESTWRGLDY
LVKESNTGQTIKIKALHAPKRDLVRDFKGASEFDQSALFKKVYEEEFGTFGGSPFGALIGDYEISRQPEDMYFIEQMSHVAAAAHAPFIASASPELLGLESFSDL
GKPRDLGKVFDTVEYAKWKSFRDAEDSRYVGLTLPRFLGRLPFNPKDGQTAENFNFVEDVDGTDHDKYLWCNAAWAFAARLTAAFDDFGWCAAIRGVEGG

GLVEDLPTHTFKTDDGEVALKCPTEIAITDRREKELSDLGFIPLVHCKNSDYAAFFAAQSVQKPKKYSTDSANANAVLSAQLQYIFSVSRVAHYLKAMMRDKIG
SFASAQNVETFLNRWISQYVLLDDNASQEQKAQFPLREASIQVSEIPGKPGSYRSVAFLRPHFQLDELSISLRLVADLPKPANS

>Burkholderia_cenocepacia_strain_K56-2
MNQQTAAAQASGAEYAAGTSLLDDIVEKSKVAKSDSEHARAKDLIGELVHQVLDGTVIVSDNLSATIDARVAELDRLISTQLSAVMHAPEFQRLESTWRGLDY
LVKESNTGQTIKIKALHAPKRDLVRDFKGASEFDQSALFKKVYEEEFGTFGGSPFGALIGDYEISRQPEDMYFIEQMSHVAAAAHAPFIASASPELLGLESFSDL
GKPRDLGKVFDTVEYAKWKSFRDAEDSRYVGLTLPRFLGRLPFNPKDGQTAENFNFVEDVDGTDHDKYLWCNAAWAFAARLTAAFDDFGWCAAIRGVEGG
GLVEDLPTHTFKTDDGEVALKCPTEIAITDRREKELSDLGFIPLVHCKNSDYAAFFAAQSVQKPKKYSTDSANANAVLSAQLQYIFSVSRVAHYLKAMMRDKIG
SFASAQNVETFLNRWISQYVLLDDNASQEQKAQFPLREASIQVSEIPGKPGSYRSVAFLRPHFQLDELSISLRLVADLPKPANS

>Burkholderia_gladioli_strain_NGJ1_T6SS-2
MESQKQFEGAPQGAVAEANDFAALLNKEFKPKNERAKEEVEAAVRTLAEQVLRDSNVVSDDVTQTINAYVAEIDRKLSEQLNHVLHHAEFQKLEGAWRGLH
HLVTHSETDPMLKIRVMNISKKELAKNLKRFKGVAWDQSPVFKRVYEEEYGQLGGEPYGCLIGDYYFDHSPQDVELLRGISQIAAASHAPFISAADSSLLGME
NWNELANPRDLSMIFTTPDYAGWRSLREMDDARYLALTMPRTLARLPYGAKTDPVEEFDFEEDTEGADSGKYTWQNAAYSMAVNINRSFKLYGWCTRIRGV
ESGGAVENLPVHTFPSDDGGVDMKCPTEIAITDRRSAELDKMGLMPLVHRKNSDVAAFIGAQSVAKPEEYDDPAATANSNLSARLPYMFACCRFAHYLKCIVR
DKIGSFSTREQMENWLSNWVMNYVDGDPVNSSEETKARKPLAAAQVVVEEVEDNPGYYTSKFFLKPHYQLEGLTVSLRLVSRLPSARAA

>Burkholderia_gladioli_strain_NGJ1-T6SS-1
MNQQTAAAQQAGALEGAETSLLDEIVEKSKVAKSQSEHARAKDLIGELVHQVLDGTVIVSDNLSATIDARVAELDRLISAQMSAVMHAPEFQKLEGTWRGLD
YLVSESNTGSTIKIKALHAPKRELVRDFKAATEFDQSALFKKVYEEEFGTFGGAPFGALVGDYEISRQPEDMYFIEQMSHVAAAAHAPFIASAAPELIGLESFAD
LGKPRDLGKVFDTVEYAKWKSFRDAEDSRYVGLTLPRFLGRLPFHPKDGQTAESFNFVEDVDGTDHSKYLWCNAAWAFAARLTAAFDDFGWCAAIRGVEGG
GLVEDLPTHTFKTDDGEIALKCPTEIAITDRREKELSDLGFIPLVHCKNSDYAAFFAAQSVQKPKKYNTDSANANAVLSAQLQYIFSVSRIAHYLKAMMRDKIG
SFASAQNVESFLNRWISQYVLLDDNATQEQKAQFPLREASVQVSEIPGKPGSYRSVAFLRPHFQLDELSISLRLVADLPKSANS

>Burkholderia_glumae_336gr-1
MNQQTAAAQQSGALEGAETSLLDEIVEKSKVAKSQSEHARAKDLIGELVHQVLDGTVIVSDNLSATIDARVAELDRLISAQMSAVMHAPEFQRLESTWRGLDY
LVSESNTGSTIKIKAMHAPKRELVRDFKAATEFDQSALFKKVYEEEFGTFGGAPFGALIGDYEISRQPEDMYFIEQMSHVAAAAHAPFIASAAPELIGLESFADL
GKPRDLGKVFDTVEYAKWKSFRDSEDSRYVGLTLPRFLGRLPFHPKDGQIAESFNFVEDVDGTDHSKYLWCNAAWAFAARLTAAFDDFGWCAAIRGVEGGG
LVEDLPTHTFKTDDGEIALKCPTEIAITDRREKELSDLGFIPLVHCKNSDYAAFFAAQSVQKPKKYNTDSANANAVLSAQLQYIFSVSRIAHYLKAMMRDKIGSF
ASAQNVESFLNRWISQYVLLDDNATQEQKAQFPLREASVQVAEIPGKPGSYRSVAFLRPHFQLDELSISLRLVADLPKSANS

>Burkholderia_glumae_BGR1_T6SS-4
MNETVVLENDSVYASLCSKINLTPVAEARPLEAFRDNDMLSEASADERIARGMSAFLDLIAQSSQPVERLDKSLLDFHIGQLDRQISRQLDAVMHTPAFQALEG
RWRGLKMLVARTDFRKNAKIEVLDVSKEALQRDFEDTPELIQSGLYRLTYIEEYDTPGGQPISAMISDFEFTNSPMDVALLRNISKVAAAAHMPFIGSVGAAFFG
KQSMEEVAAIQDIGNYFDRAEYIKWKSFRETDDARYVGLTMPRVLGRLPYGKDTTPVRAFNYEEAVKGPDHDRYLWVNASFAFAANMVRSFVNNGWCVQIR
GPQAGGKVEDLPVHLYDLGTGVQPKIPTEVLIPETREFEFANLGFIPLSFYKNHDFACFFSASSTQKPALYETKEATANSRINARLPYIFLLSRIAHYLKLIQRENIG
TTKDRRLLELELNNWIKGLVTEMKDPDDELQASHPLRDAKVTVEDIDDNPGFFRIKLFIIPHFQVEGMDIGLSLVSQMPKAKS

>Burkholderia_glumae_BGR1_T6SS-1
MNQQTAAAQQSGALEGAETSLLDEIVEKSKVAKSQSEHARAKDLIGELVHQVLDGTVIVSDNLSATIDARVAELDRLISAQMSAVMHAPEFQRLESTWRGLDY
LVSESNTGSTIKIKAMHAPKRELVRDFKAATEFDQSALFKKVYEEEFGTFGGAPFGALIGDYEISRQPEDMYFIEQMSHVAAAAHAPFIASAAPELIGLESFADL
GKPRDLGKVFDTVEYAKWKSFRDSEDSRYVGLTLPRFLGRLPFHPKDGQIAESFNFVEDVDGTDHSKYLWCNAAWAFAARLTAAFDDFGWCAAIRGVEGGG
LVEDLPTHTFKTDDGEIALKCPTEIAITDRREKELSDLGFIPLVHCKNSDYAAFFAAQSVQKPKKYNTDSANANAVLSAQLQYIFSVSRIAHYLKAMMRDKIGSF
ASAQNVESFLNRWISQYVLLDDNATQEQKAQFPLREASVQVAEIPGKPGSYRSVAFLRPHFQLDELSISLRLVADLPKSANS

>Burkholderia_mallei_ATCC_23344

MEGEHLQSPKHDDAPDATPPESPASLLDELIEAARVKRDEDAYPITRHGIQAFVAHLAKPKRPIETVSQATIDDMIAEIDRKLCRQIDAILHDPAFQQLESTWRSL
KFLVDRTDFRENVKVQILDVGKTALFDDFEDSPDITKSGLYQKVYTAEYGQFGGQPIGAIVANYTFGPGAQDVKLLQYVASTSAMAHTPFIAAAGPAFFGIDSF
GKLPNVKDLASLFEGPQYAKWNAFRESEDARYVGLTLPRFLLRLPYGANTTPVKRFNYEERVDGGDAHFLWGNAAFAFATRLTASFADYRWCANVIGPKGGG
TVTDLPLYAYESMGEIQNKIPTDVLISERREFELAEQGFIALTMRKNSDNAAFFSANSTQKPKFFGISKEGKEAELNYRLSTQLPYIFVVNRLAHYIKVIQRENIGS
WKERGDLEQELNQWIRQYVVDMDNPSQSVRSRRPLRQAQIVVSDVEGEPGWYRVDMKVRPH FKYMGAFFTLSLVGKLEKR

>Burkholderia_pseudomallei_K96243_T6SS-1
MNERAQTQADTRAAAQPVVARDEFAALLQKEFKPKTAEARESVERAVRTLAQQALEHTVGMTTDAYGSVKQIIAEIDRKLSEQINLILHHQEFQTLEGAWRGL
HYLVTNTETDELLKIKALPASRNELARTLKRYKGVAWDQSPLFRKVYEEEYGQFGGEPFGCLVGDFHFNHSPPDVEMLGELSKIAAAAHAPFIAGASPELMQM
DSWQELANPRDLTKIFQNTEYAAWRSLRQSEDSRYVGLAMPRFLARLPYGARTNPVDEFDFEEDTGAASHDRYTWANSAYAMAANINRSFKLYGWCSSIRGV
ESGGAVQGLPCHTFPTDDGGVDQKCPTEIAISDRREAELAKNGFMPFVHRKNSDFAAFIGAQSLYQPAEYHDPDATANARLSGRLPYLFACCRFAHYLKCIVRD
KIGSFRERDDMERWLNDWIMNYVDGDPANSSQETKARKPLAAAQVVVEEIDDNPGYYASKFFLRPHYQLEGLTVSLRLISKLPSAKAAGE

>Burkholderia_pseudomallei_K96243_T6SS-2
MAARESQARSADAQLATHSDFNALLSREFKPKTEQAREAVEHAVKTLAEQALANSVTLSDDAYKSIEAIIGEIDRKLSEQINLILHHDDFQQLESAWRGLHHLV
TNTETDEKLKIRFMDVSKDDLRRTMKRYKGVAWDQSPFFKQIYEEEYGQLGGEPYGCLVADYYFDHTPPDVDLLSSIGKVAAAAHAPFITGASPSVLQMDSW
QELANPRDLTKIFTQNLEYAPWNSLRNSEDARYIGLAMPRFLARLPYGIRTNPVDEFDFEESTDGSDHRKYVWANAAYAMAVNINRSFKHYGWCTLIRGVESG
GVVENLPCHTFPTDDGGIDMKCPTEIAISDRREAELAKNGFIPLIHRKNTDYAAFIGAQSLQKPAEYYDPDATANANLSARLPYLFACSRFAHYLKCIVRDKIGS
FKEREDMQQWLNEWIMNYVDADPANSSQETKARRPLAAAEVVVEDVEGNPGYYQAKFFLRPHFQLEGLTVSLRLVAKLPSVKEAA

>Burkholderia_pseudomallei_K96243_T6SS-3
MKKQQAQTAAAGVQPQADSDFAQLLAQEFKPKTEQAREAVEYAVRTLAEQALAQSATISDDAYKSIAAIIAQIDHKLSEQINLILHHADFQKLESAWRGLHHLV
SNTETDERLKIRFMDISKEELRRTMRRYKGQSWDQSPLFKQIYEEEYGQLGGEPYGCLVADYYFDHTPPDVDLLGSISKVAASAHTPFLSGASPSVLQMESWQE
LANPRDLTKIFTQNLEYASWNALRNMDDARYIGLAMPRFLSRLPYGVLTNPVDEFDFEEDTNGADHRRYAWTNAAYAMGVNINRSFRLYGWCSLIRGVESGG
TVENLPCHTFPTDDGGIDIKCPTEIAISDRREAELSKNGFIPLVHRKNTDHATFIGAQSLHKPAEYDDSDATANANLSARLPYLFACSRFAHYLKCIVRDKVGAFK
EREDMQRWLNEWIMNYVDADPANSSQDTKARRPLAAAEVVVEQAQGNPGYYQAKFFLRPHFQLEGLTVSLRLVAKLPSIKEAA

>Burkholderia_pseudomallei_K96243_T6SS-4
MSMQQLESSAEKVVVDQNNVNEDLKDILRRSFRPRTNEAAEAVQNAVETLLTYARRSRVVVREDVAQTIEQLVAELDKKISEQLTLVLHNKRFQSLEGAWRGL
HYLVSNTDTSENLKIRYLNISKADLGKTLRRFKGVVWDQSPIFKMIYEQEYGQFGGEPFGCLIGDFYFDHSMQDVSILTEMSKISAAAHAPFIAAAAPGLLQMD
DWSELSNPRDVSKIFTATEYAFWRRLRESNDSRYLALTLPRFLARVPYGPKTQPVEEFGFEEKVDPNRAEDFCWANSAYAMGANITRAFKTYGWCTKIRGVES
GGAVEVLPKFVLPSQDREVDLHCPTEIAISDRREHELSESGLMPLVYRKNSDTAAFIGAKTVHRPAIYEDDDATANSNLSSRLPYIFATCRFAHYLKCIVRDKIGS
FKSAEDTQRWLNDWLMNYVDGDPSISSEVTKSQRPLSAAEVVVDEIPENPGYYRAQFFLRPHFQLEGLTVSLRLVSKLPSTKHEVTT

>Burkholderia_pseudomallei_K96243_T6SS_5
MNQQTAAAQSTGVQAGTESSLLDEIVEKSKVAKSESEHARAKDLIGELVSQVLDGTVVVSDNLSATIDARVAELDRLISSQLSAVMHAAEFQRLESTWRGLDY
LVKESNTGSTIKIKALHAPKRDLVRDFKNATEFDQSALFKKVYEEEFGTFGGSPFGVLVGDYEISRQPEDQYFIEQMSHVAAAAHAPFIASAAPELLGLESFADL
GKPRDLGKVFDTVEYAKWKSFRDSEDSRYVGLTLPRFLGRLPFNPKDGAIAESFNFVEDVDGTDHDKYLWCNASWAFAARLTAAFDDFGWCAAIRGVEGGG
LVEDLPTHTFKTDDGEIALKCPTEIAITDRREKELSDLGFIPLVHCKNSDYAAFFAAQSVQKPKKYSTDSANANAVLSAQLQYIFSVSRIAHYLKAMMRDKIGSF
ASAQNVETFLNRWISQYVLLDDDATQEQKAQFPLREASVQVSEIPGKPGAYRSVAFLRPHFQLDELSISLRLVADLPKPANS

>Burkholderia_thailandensis_E264_T6SS-5
MEGEHLYSPKNDDAPPAPAPDSPASLLDELIEAARVKRDEEAYPITRHGIEAFVAHLARPKRPIETVSQATIDDMIAEIDRKLCRQVDAILHHPDFQQLESTWRSL
KFLVERTDFRENIKIQFLDVGKAALLDDFDDSPDITKSGLYQKVYAAEYGQFGGQPIGAIVANYTFGPGAQDVKLLQYVASTSAMAHTPFIAAAGPAFFGIDSFG
KLPNVKDLASLFEGPQFAKWNAFRESEDARYVGLTLPRFLLRLPYGANTTPVKRFNYDERVDGGDADFLWGNAAFAFATRLTASFADYRWCANVIGPKGGGT
VADLPLYAYEAMGEIQNKIPTDVLISERREFELAEQGFIALTMRKHSDNAAFFSANSTQKPKFFGISKEGKDAELNYRLGTQLPYIFVVNRLAHYIKVIQRENIGT

WKERGDLEQELNQWIRQYVADMDNPTEGVRSRRPLRQAEIFVSDVEGEPGWYRVDMKVRPHFKYMGASFTLSLVGKLEKR

>Burkholderia_thailandensis_E264_T6SS-1
MNQQTAAAQSTGAQVGTESSLLDEIVEKSKVAKSESEHARAKDLIGELVNQVLDGTVVVSDNLSATIDARVAELDRLISSQLSAVMHAAEFQRLESTWRGLDY
LVKESNTGSTIKIKALHAPKRDLVRDFKNATEFDQSALFKKVYEEEFGTFGGSPFGVLVGDYEISRQPEDQYFIEQMSHVAAAAHAPFIASAAPELLGLESFADL
GKPRDLGKVFDTVEYAKWKSFRDSEDSRYVGLTLPRFLGRLPFNPKDGAIAESFNFVEDVDGTDHDKYLWCNASWAFAARLTAAFDDFGWCAAIRGVEGGG
LVEDLPTHTFKTDDGEIALKCPTEIAITDRREKELSDLGFIPLVHCKNSDYAAFFAAQSVQKPKKYSTDSANANAVLSAQLQYIFSVSRIAHYLKAMMRDKIGSF
ASAQNVETFLNRWISQYVLLDDDATQEQKAQFPLREASVQVSEIPGKPGAYRSVAFLRPHFQLDELSISLRLVADLPKPANS

>Burkholderia_thailandensis_E264_T6SS-4
MQQLESSAEKVVVDQNNVNEDLKDILRRSFRPRTNEAAEAVQNAVETLLTYARRSRVVVREDVAQTIEQLVAELDKKISEQLTLVLHNKRFQSLEGAWRGLHY
LVSNTDTSENLKIRYLNISKADLGKTLRRFKGVVWDQSPIFKMIYEQEYGQFGGEPFGCLIGDFYFDHSMQDVSILTEMSKISAAAHAPFIAAAAPGLLQMDDW
SELSNPRDVSKIFTATEYAFWRRLRESNDSRYLALTLPRFLARVPYGPKTQPVEEFGFEEKVDPNRAEDFCWANSAYAMGANITRAFKTYGWCTKIRGVESGGA
VEVLPKFVLPSQDREVDLHCPTEIAISDRREHELSESGLMPLVYRKNSDTAAFIGAKTVHRPAIYEDDDATANSNLSSRLPYIFATCRFAHYLKCIVRDKIGSFKS
AEDTQRWLNDWLMNYVDGDPSISSEVTKSQRPLSAAEVVVDEIPENPGYYRAQFFLRPHFQLEGLTVSLRLVSKLPSTKQEVTT

>Campylobacter_jejuni_strain_WP2202
MANTKAVDMPIIEQIMEKSKYSKTDESYSIAKRGVAEFISEIVKSDNAEEKINKFALDEMIAHIDYLLSKQMDEVLHNEEFQKLESTWRGLRFLVERTDFNENIK
IDLFDIRKEEALEDFENNPDITQSVVYKNIYSSEYGQFGGEPVGAIIGDYQLGSASPDMTFLNKMASIAAMSHSPFLTSFGPKFFGLDDYSELANIQDLQGLLEGP
QYTRWRTFRENEDSKYTGLLVTRFLARSPYDPEENPIKSFNYKENVHASHNHLLWANSSYTFCTRLTESFAKYRWCGNIIGPKSGGTVKDLPTYLYENFGTIQS
KIPTEVLITDRREYELAEAGFITLTLRRDSNNAAFFSANSPLKPKLFQNTPEGKEAETNYRLGTQLPYIFLISRLAHYLKVLQREEIGSWKERSDIENGLNEWIRQ
YISDQENPPSEVRSRRPFRAAQVKVSDIPGEPGWYKIGLSVRPHFKYMGGNFELSLVGKLDKE

>Citrobacter_rodentium_DBS100
MATQAQNHKQAAPQATTQNDFNALLTREFKPKSEQAKSAVEMAVKTLAEQALSTSITMADDAYKNIAAIIAEIDLKLSEQINLILHHEEFQRLESAWRGLHYLV
NNTETDEKLKLRFMDISKDDLRRNMKRYKGIAWDQSPLFKQIYEEEYGQLGGEPYGCLVADYHFDHSAPDVDLLSSIGKVAASAHMPFITGASPSVMQMDSW
QELANPRDLTKIFTQNLEYAAWNSLRQSEDSRYIGLAMPRFLARLPYGINTNPVDNFNFEEDTDGANHSKYVWANAAYAMAVNINRSFKHYGWCTMIRGVES
GGVVEDLPCHTFPTDDGGVDMKCPTEIAISDRREAELAKNGFIPLVHRKNTDYAAFIGAQSLQKPAEYYDPDATANANLSARLPYLFACSRFAHYLKCIVRDKI
GSFKEREDMQRWLNNWVMNYVDGDPANSSQETKARRPLAAAEVVVEDVEGNPGYYQAKFFLRPHFQLEGLTVSLRMVAKLPSLKDVA

>Edwardsiella_ictaluri_93-146
MSEQNLPESAAVESAAPEDSLLDSIIAETRMARSDLERERARDLLGEWVSEVLSGTVTVSGDVLASIEARIAQIDALLSAQLSTIMHEPAFQKLEGSWRGLHYL
VHQSETGTGLKIRMLNVSRKDLIRDFKSAAEFDQSALFKKVYEEEYGTFGGAPFAAMIGDYEFSNHPEDLFLLEEISHVAAAAHAPFLSAASAGMFGLDSLTEL
SIPRDLAKGFDTVEYAKWKSLRQSEDARYIALALPHVLGRLPYGAATVPVEAFNFEEDVNGKEHGKYLWLNAAYALGTRLTQAYAKYGWCAAIRGVEGGGL
VEGLPAHTFTTDDGEVELKCPTEVAITDRREKELADLGFIALTHCKGTDYAAFFSTQSAQKPKEYDSDSANANARIACQLQYIMATSRFAHYLKSMVRDKLGS
FMSRSECEYFLNQWISNYVVGSDDAGQDIKAKYPLREARIDVSDIPGKPGFYKAVAYLKPHFQLEGLTASLRLVADLPPPAQG

>Edwardsiella_tarda_EIB202
MSEQNLPESSAAESAALEGGLLDSIIAETRMARSDLEKARARDLLGEWVSEVLSGTVTVSGDVLASIEARIAQIDALLSAQLSAIMHEPAFQKLEGSWRGLHYL
VHQSETGTGLKIRMLNVSRADLIRDFKSAAEFDQSALFKKVYEEEYGTFGGAPFAAMIGDYEFSNHPEDLFLLEEISHVAAAAHAPFLSAASAGMFGLDSLTEL
SIPRDLAKGFDTVEYAKWKSLRQSEDARYIALALPHVLGRLPYGATTVPVESFNFEENVSGKEHGKYLWLNAAYALGTRLTQAYAKYGWCAAIRGAEGGGLV
EGLPAHTFTTDDGEVELKCPTEVAITDRREKELAELGFIALTHCKGTDYAAFFSTQSVQKPKEYDSDSANANARISCQLQYIMATSRFAHYLKSMVRDKIGSFM
SRSECEYFLNQWISNYVVGSDDAGQDIKAKYPLREARIDVSDIPGKPGFYKAVAYLKPHFQLEGLTASLRLVADLPPPAQG

>Edwardsiella_tarda_PPD130/91
MSEQNLPESSAAESAALEGGLLDSIIAETRMARSDLEKARARDLLGEWVSEVLSGTVTVSGDVLASIEARIAQIDALLSAQLSAIMHEPAFQKLEGSWRGLHYL

VHQSETGTGLKIRMLNVSRADLIRDFKSAAEFDQSALFKKVYEEEYGTFGGAPFAAMIGDYEFSNHPEDLFLLEEISHVAAAAHAPFLSAASAGMFGLDSLTEL
SIPRDLAKGFDTVEYAKWKSLRQSEDARYIALALPHVLGRLPYGATTVPVESFNFEENVSGKEHGKYLWLNAAYALGTRLTQAYAKYGWCAAIRGAEGGGLV
EGLPAHTFTTDDGEVELKCPTEVAITDRREKELAELGFIALTHCKGTDYAAFFSTQSVQKPKEYDSDSANANARISCQLQYIMATSRFAHYLKSMVRDKIGSFM
SRSECEYFLNQWISNYVVGSDDAGQDIKAKYPLREARIDVSDIPGKPGFYKAVAYLKPHFQLEGLTASLRLVADLPPPAQG

>Enterobacter_cloacae_strain_DSM_30054_T6SS-2
MSVNTENGSAQGQTTVLEKESVYASLFDKINLTPATSLGDINAFLDDAALSDAPAGERLTAAMQVFMDCIRKSGQPVEKLDKTLIDHHIAELDFQISRQLDAVM
HHAEFQKVESLWRGLKQLVDNTDYRQNVKTEILDVSKDDLRQDFEDAPELIQSGLYWHTYTAEYDTPGGEPIGSVISAYEFDASPQDVALLRNISKVSAAAHM
PFIGAVGPKFFLKESMEEVAAIKDIGNYFDRAEYIKWKSFRDTDDARYIGLVMPRVLGRLPYGPDTVPVRSFNYVEQVKGPDHEKYLWTSASFSFASNMVKSFI
NNGWCVQIRGPQAGGAVKDLPIHLYDLGTGNQVKIPSEVMIPETREFEFANLGFIPLSYYKNRDYACFFSANSAQKPALYDTADATANSRINARLPYIFLLSRIAH
YLKLIQRENIGTTKDRRLLELELNTWVRSLVTEMTDPGDELQASHPLRDAKVVVEDIEDNPGFFRVKLFAVPHFQVEGMDVNLSLVSQMPKAKAKA

>Enterobacter_cloacae_strain_DSM_30054_T6SS-1
MSNQTQQHEQQAGQAFSQDEFSALLNKEFRPKTDQARSAVESAVKTLAQQALENTVTFSNDTYRTIQNLIAGIDEQLSQQVNQIIHHEEFQKLESAWRGLSYL
VNNTETDEMLKIRFMSISKQELGRTLKRYKGVGWDQSPIFKKIYEQEYGQFGGEPFGCIVGDYYFDHSPQDVELLGEMARIGSAAHCPFITGTAPGVMQMESW
QELANPRDLTKIFQNTEYAAWRSLRESEDARYLGLVMPRFLSRLPYGIRTNPVDSFDFEEQTDGANHNSYAWANAAYAMAANINRSFKEYGWCTSIRGVESGG
AVENLPCHTFPSDDGGVDMKCPTEIAISDRREAELAKNGFMPLVHRKNSDFAAFIGAQSLQKPAEYHDPDATANARLASRLPYLFACCRFAHYLKCIVRDKIGS
FREREEMERWLNDWVMNYVDGDPANSSQETKSRKPLAAAEVQVQEIEDNPGYYAAKFFLRPHYQLEGLTVSLRLVSKLPSLKTKDA

>Enterobacter_cloacae_subsp._cloacae_ATCC_13047_T6SS-2
MLMSVNTENGSAQGQTTVLEKESVYASLFDKINLTPATSLGDINAFLDDAALSDAPAGERLTAAMQVFMDCIRKSGQPVEKLDKTLIDHHIAELDFQISRQLDA
VMHHAEFQKVESLWRGLKQLVDNTDYRQNVKTEILDVSKDDLRQDFEDAPELIQSGLYWHTYTAEYDTPGGEPIGSVISAYEFDASPQDVALLRNISKVSAAA
HMPFIGAVGPKFFLKESMEEVAAIKDIGNYFDRAEYIKWKSFRDTDDARYIGLVMPRVLGRLPYGPDTVPVRSFNYVEQVKGPDHEKYLWTSASFSFASNMVK
SFINNGWCVQIRGPQAGGAVKDLPIHLYDLGTGNQVKIPSEVMIPETREFEFANLGFIPLSYYKNRDYACFFSANSAQKPALYDTADATANSRINARLPYIFLLSRI
AHYLKLIQRENIGTTKDRRLLELELNTWVRSLVTEMTDPGDELQASHPLRDAKVVVEDIEDNPGFFRVKLFAVPHFQVEGMDVNLSLVSQMPKAKAKA

>Enterobacter_cloacae_subsp._cloacae_ATCC_13047_T6SS-1
MSNQTQQHEQQAGQAFSQDEFSALLNKEFRPKTDQARSAVESAVKTLAQQALENTVTFSNDTYRTIQNLIAGIDEQLSQQVNQIIHHEEFQKLESAWRGLSYL
VNNTETDEMLKIRFMSISKQELGRTLKRYKGVGWDQSPIFKKIYEQEYGQFGGEPFGCIVGDYYFDHSPQDVELLGEMARIGSAAHCPFITGTAPGVMQMESW
QELANPRDLTKIFQNTEYAAWRSLRESEDARYLGLVMPRFLSRLPYGIRTNPVDSFDFEEQTDGANHNSYAWANAAYAMAANINRSFKEYGWCTSIRGVESGG
AVENLPCHTFPSDDGGVDMKCPTEIAISDRREAELAKNGFMPLVHRKNSDFAAFIGAQSLQKPAEYHDPDATANARLASRLPYLFACCRFAHYLKCIVRDKIGS
FREREEMERWLNDWVMNYVDGDPANSSQETKSRKPLAAAEVQVQEIEDNPGYYAAKFFLRPHYQLEGLTVSLRLVSKLPSLKTKDA

>Enterobacter_sp._DSM_30060
MSNQTQQHDQQAGQAFSQDEFSALLNKEFRPKTDQARSAVESAVKTLAQQALENTVTFSNDTYRTIQNLIAGIDEQLSQQVNQIIHHEEFQKLESAWRGLSYL
VNNTETDEMLKIRFMSISKQELGRTLKRYKGVGWDQSPIFKKIYEQEYGQFGGEPFGCIVGDYYFDHSPQDVELLGEMARIGSAAHCPFITGTAPGVMQMESW
QELANPRDLTKIFQNTEYAAWRSLRESEDARYLGLVMPRFLSRLPYGIRTNPVDSFDFEEQTDGANHNSYSWANAAYAMAANINRSFKEYGWCTSIRGVESGG
AVENLPCHTFPSDDGGVDMKCPTEIAISDRREAELAKNGFMPLVHRKNSDFAAFIGAQSLQKPAEYHDPDATANARLASRLPYLFACCRFAHYLKCIVRDKIGS
FREREEMERWLNDWVMNYVDGDPANSSQETKSRKPLAAAEVQVQEIEDNPGYYAAKFFLRPHYQLEGLTVSLRLVSKLPSLKTKDA

>Escherichia_coli_042_T6SS-1
MLMSVQKEKNVAESVVSEAHAGDSVYASLFEKINLSPVSALSALDIWQDPQAMSEASADERLTAGMQVFMECLAKAGTQVEKLDKALIDHHIAELDYQISRQ
LDAVLHHPEFQKVESLWRGVKSLVDKTDFRRNVKIELLDLSKDDLRQDFEDAPEIIQSGLYLQTYVAEYDTPGGEPIAALVSAWEFDASAQDVALLKNISRVAA
SAHMPFIGSVGPAFFQKETMEEVAAIKDIGNHFERAEYIKWNAFRETDDARYIGLVMPRVLGRLPYGPDTVPVRSFNYVEEVKGPDHHKYLWTNASFAFAAN
MVRSFVTNGWCVQIRGPQAGGAVQDLPIHLYDLGTGNQVKIPSEVMIPETREFEFANLGFIPLSYYKNRDYACFFSANSAQKPALYDTPDATANSRINARLPYIF
LLSRIAHYLKIIQRENIGTTKDRRLLELELNNWIRGLVTEMTDPGDELQASHPLRDGKVVVEDIEDNPGFFRVKLYAVPHFQVEGMDVSLSLVSQMPKAKA

>Escherichia_coli_042_T6SS-2

MTVASTLGLNETQYATDDCLEEIINNTRAVRQDSEKTRFKLQINNFLAEVASGSLVINSDLIGSIEKRIADIDKLMSEQLSLIMHATEFQKIESAWTGLYKLVQASV
TENVKYTVLHCTKKELLKDFKSASDFDQSVLFKNIYESEYGTFGGTPYSAFVGDFYFDNTPQDIDLLEHISHVAASAHAPFLSAIAPGMLSMSSFSELPYPRDLA
KLFETTDYARWRSFRQTDDSRYVGLTLPQSLGRIPYGMKTIPAETFNFEEHISEDNSGKDYLWVNTAFELACRIVDAFEEYGWCAAIRGVEGGGLVKSLPAYNY
VSHTGERLLQCPTEVAISDRREKELSDLGFIPLVYCKGTDFAAFFAVQSVNKARLYNTDQANANAKLSSQLQYILATSRFAHYLKVIVRDKVGSFMSRTECQTY
LQNWIMQYVVASDNAGQETKARYPLREASVEVIEVPGSPGNYRAIAWIKPHFQLEGLSMSLRLVADLPSSVS

>Escherichia_coli17-2_T6SS-1

MLMSVQKEKNVAESVVSEAHAGDSVYASLFEKINLSPVSALSALDIWQDPQAMSEASADERLTAGMQVFMECLAKAGTQVEKLDKALIDHHIAELDYQISRQ
LDAVLHHPEFQKVESLWRGVKSLVDKTDFRRNVKIELLDLSKDDLRQDFEDAPEIIQSGLYLQTYVAEYDTPGGEPIAALVSAWEFDASAQDVALLKNISRVAA
SAHMPFIGSVGPAFFQKETMEEVAAIKDIGNHFERAEYIKWNAFRETDDARYIGLVMPRVLGRLPYGPDTVPVRSFNYVEEVKGPDHHKYLWTNASFAFAAN
MVRSFVTNGWCVQIRGPQAGGAVQDLPIHLYDLGTGNQVKIPSEVMIPETREFEFANLGFIPLSYYKNRDYACFFSANSAQKPALYDTPDATANSRINARLPYIF
LLSRIAHYLKIIQRENIGTTKDRRLLELELNNWIRGLVTEMTDPGDELQASHPLRDGKVVVEDIEDNPGFFRVKLYAVPHFQVEGMDVSLSLVSQMPKAKA

>Escherichia_coli17-2_T6SS-2

MTVASTLGLNETQYATDDCLEEIINNTRAVRQDSEKTRFKLQINNFLAEVASGSLVINSDLIGSIEKRIADIDKLMSEQLSLIMHATEFQKIESAWTGLYKLVQASV
TENVKYTVLHCTKKELLKDFKSASDFDQSVLFKNIYESEYGTFGGTPYSAFVGDFYFDNTPQDIDLLEHISHVAASAHAPFLSAIAPGMLSMSSFSELPYPRDLA
KLFETTDYARWRSFRQTDDSRYVGLTLPQSLGRIPYGMKTIPAETFNFEEHISEDNSGKDYLWVNTAFELACRIVDAFEEYGWCAAIRGVEGGGLVKSLPAYNY
VSHTGERLLQCPTEVAISDRREKELSDLGFIPLVYCKGTDFAAFFAVQSVNKARLYNTDQANANAKLSSQLQYILATSRFAHYLKVIVRDKVGSFMSRTECQTY
LQNWIMQYVVASDNAGQETKARYPLREASVEVIEVPGSPGNYRAIAWIKPHFQLEGLSMSLRLVADLPSSVS

>Escherichia_coli_CFT073

MSVQQEHATSETATLTTTESGGVYQSLFDKINLTPVSSIQEIDLWQNSETLADASPDERVTAAIHVLLSCLAKSGEDVVKLDKSLLDFHIDDLDQKISKQLDAVM
HHPEFQKVESLWRGTWFVVQRTDFRKNVRIELLDISKEHLRQDFDDSPEIIQSGLYRHTYIQEYDTPGGEPVASLISSYEFDNSPQDIALLRNISRVSAASHMPFIG
SVGPKFFLKNSMEEVAAIKDIGNYFDRAEYIKWKSFRDTDDSRYVGLVMPRVLGRLPYGPDTVPVRSFNYVEEVKGPDHEKYLWTNASFAFAANMVKSFVNN
GWCVQIRGPQAGGAVADLPIHLYDLGTGNQVKIPSEVMIPETREFEFANLGFIPLSYYKNRDYACFFSANSAQKPALYDTADATANSRINARLPYIFLLSRIAHYL
KIIQRENIGTTKDRRVLELELNTWIRTLVTEMTDPGDELQASHPLRDGKVIVEDIEDNPGFFRVRLFAVPHFQIEGMDVNLSLVSQMPKAKA

>Escherichia_coli_DE719

MYIQAVTGQFGGEPVAAVIGNFAFKNTTPDMKLLKYISQVSAMAHSPFLSSVSSEFFGLDSWTELPGIKEPGAIFEGPAYSRWRALRESEDSRYLGLTAPRFLLR
HPYSPDENPVKTFRYHEDVSQSHESYLWGNTSFLLAANLAESFAKYRWCPNIIGPSSGGAVKDLPVHLYESMGQMQAKIPTEVLITDRREYELAEEGFITLTMR
KGSDNACFFSANSVQKPKTFPKTPEGKAAETNYKLGTQLPYLFVISRLAHYIKVIQREQLGSWKERSDLERELNTWIRQYVADQENPPAEVRSHRPLRQAKIEV
LDVDGEPGWYQVAISVRPHFKYMGASFDLSLVGRLDKE*

>Escherichia_coli_KTE165

MSVQQEHATSETATLTTTESGGVYQSLFDKINLTPVSSIQEIDLWQNSETLADASPDERVTAAIHVLLSCLAKSGENVVKLDKSLLDFHIDDLDQKISKQLDAVM
HHPEFQKVESLWRGTWFVVQRTDFRKNVRIELLDISKEHLRQDFDDSPEIIQSGLYRHTYIQEYDTPGGEPVASLISSYEFDNSPQDIALLRNISRVSAASHMPFIG
SVGPKFFLKNSMEEVAAIKDIGNYFDRAEYIKWKSFRDTDDSRYVGLVMPRVLGRLPYGPDTVPVRSFNYVEEVKGPDHDKYLWTNASFAFAANMVKSFVN
NGWCVQIRGPQAGGAVADLPIHLYDLGTGNQVKIPSEVMIPETREFEFANLGFIPLSYYKNRDYACFFSANSAQKPALYDTADATANSRINARLPYIFLLSRIAHY
LKIIQRENIGTTKDRRVLELELNTWIRTLVTEMTDPGDELQASHPLRDGKVIVEDIEDNPGFFRVRLFAVPHFQIEGMDVNLSLVSQMPKAKA

>Escherichia_coli_PCN033

MSLQEEELVSSHAGQTDQASSLLDQIMAQTRIQPGSEGYDVARQGVTAFIASILQSTASAEPVNKLAVDSMIADIDERISRQMDVIIHAPAFQQVESFWRSLKTM
VDRVDFRENIKVNVLHVTKQELLEDFEFAPEIIQSGFYKHVYSSGFGQFGGEPIAAVLGAYEFKNTAPDMKLLQYVSAVGAMAHAPFLSSVSPEFMGLNSWTE
LPNIKDLYAIFEGPAYTKWRALRDSEDSRYLGLTAPRFLLRQPYSPTDNPVKNFNYHEDVSRNHEDYLWGNTAWMLACNVADSFAKYRWCPNIIGPQSGGAVK

DLPVHLFETMGQIQAKIPTEVLVTDRREFELAEEGFITLTMRKDSDNAAFFSANSVQKPKHFPGKDAETNYKLGTQLPYLFIINRLAHYIKVLQREQLGSWKER
SDLERELNTWIRQYVADQENPPADVRSRKPLRAARVEVMDVEGEPGWYQVALSVRPHFKFMGANFELSLVGRLDRE

>Escherichia_coli_RS218
MYIQAVTGQFGGEPVAAVIGNFAFKNTTPDMKLLKYISQVSAMAHSPFLSSVSSEFFGLDSWTELPGIKEPGAIFEGPAYSRWRALRESEDSRYLGLTAPRFLLR
HPYSPDENPVKTFRYHEDVSQSHESYLWGNTSFLLAANLAESFAKYRWCPNIIGPSSGGAVKDLPVHLYESMGQMQAKIPTEVLITDRREYELAEEGFITLTMR
KGSDNACFFSANSVQKPKTFPKTPEGKAAETNYKLGTQLPYLFVISRLAHYIKVIQREQLGSWKERSDLERELNTWIRQYVADQENPPAEVRSHRPLRQAKIEV
LDVDGEPGWYQVAISVRPHFKYMGASFDLSLVGRLDKE*

>Escherichia_coli_SEPT362
MSLQEEELVSSHAGQPEQASSLLDQIMAQTRIQPGSEGYDVARQGVTAFIASILQSTASAEPVNKLAVDSMIADIDERISRQMDVIIHAPAFQQVESFWRSLKTM
VDRVDFRENIKVNVLHVTKQELLEDFEFAPEIIQSGFYKHVYSSGFGQFGGEPIAAVLGAYEFKNTAPDMKLLQYVSTVGAMAHAPFLSSVSPEFMGLNSWTE
LPNIKDLYAIFEGPAYTKWRALRDSEDSRYLGLTAPRFLLRQPYSPTDNPVKNFNYYEDVSQNHEDYLWGNTAWMLACNIADSFAKYRWCPNIIGPQSGGAVK
DLPVHLFETMGQIQAKIPTEVLVTDRREFELAEEGFITLTMRKDSDNAAFFSANSVQKPKHFPGKDAETNYKLGTQLPYLFIINRLAHYIKVLQREQLGSWKER
SDLERELNTWIRQYVADQENPPADVRSRKPLRAAKVEVMDVEGEPGWYQVALSVRPHFKFMGANFELSLVGRLDRE

>Helicobacter_hepaticus_ATCC_51449
MSGASANTQTSIKELSIIDSIMQTSRYSKEDESYNIAKMGVVEFITEIVKTDSAENKINKYTLDEMIAHIDDIISRQMDEILHNEQIQQLESTWRGLYFLVERTNFQ
ENIKINLFDVTKQEALEDFEANPDITTATLYKRIYSSEYGQFGGEPVGAILGDYALNASTPDMNFLSKMSSIAAMSHAPFLTSMSAGFFGLDNYAELPKIQDLKA
LLEGPQYVKWRTFRENEDSKYAGLLVTRFLTRSPYEPQENPIKKFNYKENVHNSHNHLLWGNTIYAFATRLTDSFANYRWCGNIIGPKAGGAVKDLPTYIYESF
GTTQSKIPTEVLITDRCEYELAESGFIAFTLRRDSNNAVFFSANAALKPKIFPNTPEGKEAETNYRLGTQLPYIFLVSRLAHYLKVLQREEIGSWKERADIENGLN
EWMRQYVSDQENPPAEVRSRRPFRGAKVLVSEIEGEAGWYRINLNVRPHFKFMGANFELSLVGKLDRE*

>Klebsiella_pneumoniae_strain_Kp52.145
MSVTTENAPVQGQTTLQENSAGEGVYASLFEKINLTPASRLGDINDFLDDAALSEAPAAERLTAAMQVFMERIRQSGQRVEKLDKTLIDHHIAELDFQISRQLD
AVMHHQEFQQVESLWRGLKQLVDNTDYRQNVKTEILDVAKDDLRQDFEDAPELIQSGLYWHTYTAEYDTPGGEPIGSVISAYEFDASPQDVALLRNISRVSAA
AHMPFIGAVGPAFFLKETMEEVAAIKDIGNYFDRAEYIRWKAFRETDDARYIGLVMPRVLGRLPYGPDTVPVRSFNYVEQVKGPDHEKYLWTSAAFSFASNMV
KSFVNNGWCVQIRGPQAGGAVKDLPIHLYDLGTGNQVKIPSEVMIPETREFEFASLGFIPLSYYKNRDYACFFSANSAQKPALYDTADATANSRINARLPYIFLLS
RIAHYLKMIQRENIGTTKDRRLLELNTWVRSLVTEMTDPGDELQASHPLHDASVVVEDIEDNPGFFRVKLYAVPHFQVEGMDVNLSLVSQMPKAKA

>Klebsiella_pneumoniae_subsp._pneumoniae_HS11286
MSVTTENAPVQGQTTLQENRAGEGVYASLFEKINLTPASRLGDINDFLDDAALSEAPAAERLTAAMQVFMERIRQSGQRVEKLDKTLIDHHIAELDFQISRQLD
AVMHHQEFQQVESLWRGLKQLVDNTDYRQNVKTEILDVAKDDLRQDFEDAPELIQSGMYWHTYTAEYDTPGGEPIGSVISAYEFDASPQDVALLRNISRVSAA
AHMPFIGAVGPAFFLKETMEEVAAIKDIGNYFDRAEYIRWKAFRETDDARYIGLVMPRVLGRLPYGPDTVPVRSFNYVEQVKGPDHEKYLWTSAAFSFASNMV
KSFVNNGWCVQIRGPQAGGAVKDLPIHLYDLGTGNQVKIPSEVMIPETREFEFASLGFIPLSYYKNRDYACFFSANSAQKPALYDTADATANSRINARLPYIFLLS
RIAHYLKMIQRENIGTTKDRRLLELELNTWVRSLVTEMTDPGDELQASHPLRDASVVVEDIEDNPGFFRVKLYAVPHFQVEGMDVNLSLVSQMPKAKA

>Klebsiella_pneumoniae_subsp._pneumoniae_NTUH-K2044
MSVTTENAPVQGQTTLQENSAGEGVYASLFEKINLTPASRLGDINDFLDDAALSEAPAAERLTAAMQVFMERIRQSGQRVEKLDKTLIDHHIAELDFQISRQLD
AVMHHQEFQQVESLWRGLKQLVDNTDYRQNVKTEILDVAKDDLRQDFEDAPELIQSGLYWHTYTAEYDTPGGEPIGSVISAYEFDASPQDVALLRNISRVSAA
AHMPFIGAVGPAFFLKETMEEVAAIKDIGNYFDRAEYIRWKAFRETDDARYIGLVMPRVLGRLPYGPDTVPVRSFNYVEQVKGPDHEKYLWTSAAFSFASNMV
KSFVNNGWCVQIRGPQAGGAVKDLPIHLYDLGTGNQVKIPSEVMIPETREFEFASLGFIPLSYYKNRDYACFFSANSAQKPALYDTADATANSRINARLPYIFLLS
RIAHYLKMIQRENIGTTKDRRLLELELNTWVRSLVTEMTDPGDELQASHPLRDASVVVEDIEDNPGFFRVKLYAVPHFQVEGMDVNLSLVSQMPKAKA

>Kosakonia_oryzae_strain_KO348_T6SS-2
MLMSVQNESVATGESVVLQGTQAGGVYASLFEKINLNPVTTLSALDIWQDAQAMSDATADERLTAGMQVFLECLTKSDSKVEKLDRNLIDHHIAELDYQISRQ

LDAVMHHEAFQAVESLWCGLKSLVDKTDFRQNVKIELLDLSKDDLRQDFEDSPEIIQSGLYKHTYIDEYDTPGGEPIAALISAYEFDASAQDVALLRNISKVSAA
AHMPFIGSAGPKFFLKDAMADVAAIKDIGNYFDRAEYIKWKSFRETDDSRYIGLVMPRVLGRLPYGPDTVPVRSFNYVEEVKGPDHDKYLWTNASFAFAANM
VRSFINNGWCVQIRGPQAGGAVQDLPIHLYDLGTGNQVKIPSEVMIPETREFEFANLGFIPLSYYKNRDYACFFSANSTQKPALYDTADATANSRINARLPYIFLL
SRIAHYLKLIQRENIGTTKDRRLLELELNTWVRSLVTEMTDPGDELQASHPLRDAKVVVEDIDDNPGFFRVKLYAIPHFQVEGMDVNLSLVSQMPKAKS

>Kosakonia_oryzae_strain_KO348_T6SS-1
MSNPSQQQELQQGAQAFSQDEFSALLSKEFRPKTDQARSAVESAVKTLAQQALENTVTFSSDTYRTIQNLIAGIDEKLSQQINQIIHHEDFQKLESAWRGLSHLV
NNTETDEMLKIRFMSISKQELGRNLKRYKGVGWDQSPLFKKIYEEEYGQFGGEPFGCLVGDYYFDHSPQDVELLGEMARIGAAAHCPFITGTAPSVMQMESW
QELANPRDLTKIFQNTEYAAWRSLRESEDARYLGLVMPRFLSRLPYGIRTNPVDSFDFEEETDGANHNNYSWANAAYAMATNINRSFKEYGWCTSIRGVESGG
AVENLPCHTFPSDDGGVDMKCPTEIAISDRREAELAKNGFMPLIHRKNSDFAAFIGAQSLQKPMEYHDADATANARLASRLPYLFACCRFAHYLKCIVRDKIGS
FRERDEMERWLNDWVMNYVDGDPANSSQETKARKPLAAAEVQVQEIEDNPGYYAAKFFLRPHYQLEGLTVSLRLVSKLPSLKTKEA

>Methylomonas_methanica_MC09
MAELDAQAQQGATETLEFDDFSALLSKEFKPGTEAANQVNTAVSTLAQFALQDVSKISDDAIKSIQSIIASLDEKISEQLNLVLHHPDFQQLEGAWRGLHYLVN
NTETDEMLKIKVFNVSKKELGKTLKKFKGTAWDQSPLFKKLYEEEYGTFGGEPFGCLVGDYHFDHSPPDVELLGEMAKIAAASHTPFLSGVAPSVLQMDSWS
ELANPRDLTKIFQTPEYAAWRSLRDSDDSRYLGLAMPRFLSRLPYGAKTDPVDEFDFEEDTSGADSTKYTWSNAAYAMAVNINRSFKLYGWCSRIRGIESGGA
VEGLPVHTFPTDDGGVDMKCPTEIAITDRREAELAKSGFMPLIHKKNSDFAAFIGAQSLQKPAEYDDPDATANANLAARLPYLFATCRFAHYLKCMVRDKVGS
FKERDDMEKWLTSWINNYVDPNPAMSDELTKSRKPLAAAEVVVEEVEGNPGYYSSKFFLRPHYQLEGLTVSLRLVSKLPSGKGG

>Myxococcus_xanthus_DK_1622
MANETQTQKSTGVANDASLSLLDEILSEAKLKPKDEGYDVAKRGVQAFITEMLAPNRSEERVDKALVDAMIAEIDKRLSSQVNEILHAKEFQKLESSWRSLKF
MVDRTDFRENTRVEMLNASKEDLQKDFEDAPEVTKSGLYKLVYSNEYGVFGGKPYGIISANYDFNVGPQDMELLRKCASVAAMAHAPFIGNAAPEVFGEESF
LKLPDLKDLKSLFEGPQYARWHSFRESEDARYVGLALPRFLLRLPYGEKTVPVKAFNFTEDVVGHHERYLWGHASVALTSRVADSFAKFRWSPNIIGPQSGGA
VENLPLHQYEAMGEIQTKKIPTEVMLTERREFELSEEGFIGLVFRKDSDNAAFFSANSTQKPRFFGNTPEGKAAETNYRLGTQLPYMFIMTRLAHYIKVLQREQI
GSWKEKSDLERELNHWLSQYISDMDDPAPAVRSRRPLRAARVVVEDVEGQPGWYRCSLQVRPHFKYMGASFTLSLVGKLDKE

>Pantoea_ananatis_LMG_2665
MNQTSQQPQSQGGASFSQDEFSALLNKEFRPKSDQAREAVESAVKTLAQQALENTVTMSSDAYRTIQALIAEIDEKLSQQVNQIIHHEEFQKLESAWRGLSYLV
NNTETDEMLKIRFMSISKQELSRILKRYKGVGWDQSPLFKKIYEQEYGQFGGEPFGCLVGDYYFDHSPQDVELLGEMARIGAAAHCPFITGTAPSVMQMESW
QELANPRDLTKIFQNTEYAAWRSLRESEDARYLGLVMPRFLSRLPYGIRTNPVDSFDFEEQTDGSDHSNYSWTNAAYAMAANINRSFKEYGWCTSIRGVESGG
AVEDLPCHTFPSDDGGVDMKCPTEIAISDRREAELAKNGFMPLVHRKNSDFAAFIGAQSLQKPAEYHDADATANARLAARLPYLFACCRFAHYLKCIVRDKIG
SFRERDEMERWLNDWVMNYVDGDPANSSQETKSRKPLAAAEVNVEEQEDNPGYYAAKFFLRPHYQLEGLTVSLRLVSKLPSLKSNEN

>Paraburkholderia_phymatum_STM815_T6SS-3
MAKQHAQTAVAGVATGSDFTQLLNQEFRPKTQQAREAVELAVQTLAEQALRQSATISDDAYKSIAAIIAQIDHKLSEQINLILHHDDYQKLESAWRGLHHLVSN
TETDERLKIRFMDISKDELRRTMKRYKGLAWDQSPLFKQIYEEEYGQLGGEPYGCLVADYYFDHTPPDVDLLGSIAKISAAAHTPFISGASPAVLQMESWQELA
NPRDLTKIFTQNLEYAPWNSLRNTEDARYIGLAMPRFLARLPYGAKTNPVDEFDFEEDTAGSDHRHYAWSNAAYAMGVNINRSFKLYGWCSLIRGVESGGTV
ENLPCHTFPTDDGGVDMKCPTEIAISDRREAELSKNGFIPLIHRKNTDHATFIGAQSMQKPAEYHDADATANANLSARLPYLFACSRFAHYLKCIVRDKIGAFKE
REDMQRWLNEWIMNYVDADPANSSQETKARRPLAAAEVLVEEVEGNPGYYQAKFFLRPHFQLEGLTVSLRLVAKLPSIKEAA

>Paraburkholderia_phymatum_STM815_T6SS-b
MASNTAAGGAARETQTLDASAGAELSLLERIVHEGNMAVEPSQSGYAKKLIGQLASQILDEGMRTSPDKSVVAMINERVAEIDKLLSDQLNAIMHDEAFQALE
ASWTGLHDMVYGTETGPQLKLRLLNVTKKELLKDLETAVDHDMSTLFKKIYEEEYGTFGGAPYSLLIGDYSFGRHPQDIALLERISKVAAAAHAPFIASAAPG
LFDLKSFTDLGVTRDLSKTFESAELAAWRSFRDSEDSRYVSLVLPSYAARLPYGAKTKPVENFNFEEDVDGTDHGKYLWANSAYQLGLRITDAFAKYSWATAI
RGVEGGGKVDNMVAHTYKTGEGDIVLKCPTEVTITDRREKELNDLGFIALVNSKGSNFATFFGGQTTNRPKVYNKDAANANAQLSARLPYVLAASRFAHYM
KVIMRDKVGSFQSRSDIESYLNNWVADYVLINPSASHAQKARFPLGEARVDVTEVPGKPGAYRATCFLKPHFQMEELTASIRLVAELPAAAA

>Pectobacterium_atrosepticum_SCRI1043

MSIVEEQVQAGATSASGSLLDDIMAQARISPVDEGYSVAKQGIAALVANILDSGNAAEPVNKALVDSMIVELDKKLSKQIDVILHAKELQELESSWRSMKLLID
RTDFRENIKLLVLHATKEELLEDFEFAPEISQSGFYKHVYSSGYGQFGGQPIGGVIGDYALTQSSPDIKLMQYVSAVGAMAHAPFISSVAPTFFGVDRFTDLPSIK
DLKSVFEGPAYTKWRSLRESEDGRYLGLTAPRFLARLPYDPVENPIKGFNYKEDISTDHEHYLWGNTAYLMGTALTDSFAKYRWCPNIIGPQSGGAITDLPVHV
YEAMGQLQAKIPTEVLITDRREYEMAEEGFITLTMRKDSDNAAFFSANSVQKPKVFPNTKEGKEAETNYKLGTQLPYMFIINRLAHYIKVLQREQIGSWKERQ
DLERELNTWIKQYIADQENPPADVRSRRPLRAAQIKVLDVEGEPGWYQVTMSVRPHFKYMGANFELSLVGRLDKE

>Pectobacterium_brasiliense_1692

MSIVEEQVQAGATSASGSLLDDIMAQARISPVDEGYSVAKQGIAALVANILDSGNAAEPVNKALVDSMIVELDKKLSKQIDVILHAKELQELESSWRSMKLLID
RTDFRENIKLLVLHATKEELLEDFEFAPEISQSGFYKHVYSSGYGQFGGQPIGGVIGDYALTQSSPDIKLMQYVSAVGAMAHAPFISSVAPTFFGVDRFTDLPSIK
DLKSVFEGPAYTKWRSLRESEDGRYLGLTAPRFLARLPYDPVENPIKGFNYKEDISTDHEHYLWGNTAYLMGTALTDSFAKYRWCPNIIGPQSGGAITDLPVHV
YEAMGQLQAKIPTEVLITDRREYEMAEEGFITLTMRKDSDNAAFFSANSVQKPKVFPNTKEGKEAETNYKLGTQLPYMFIINRLAHYIKVLQREQIGSWKERQ
DLERELNTWIKQYIADQENPPADVRSRRPLRAAQIKVLDVEGEPGWYQVTMSVRPHFKYMGANFELSLVGRLDKE

>Photorhabdus_laumondii_subsp._laumondii_TTO1

MSLHEESSTTLTPGSTTSLLDEIMSQARMTPENDGYHIAKQGVAAFISSILDTGTNEEPVNKLVVDRMIVELDKTLSTQVDEIMHAKEFQELESSWRSLKLLVDR
TDFRENIKINIIHATKEELLEDFEFSPEIIQSGLYKHVYSTGYGQFGGEPVAAVIGNYAFSNSSPDIKLMQYVSAVGAMAHAPFLSSVAPDFFGISSFTELPAIKDLK
SVFEGPAHTKWRALRESEDSRYLGLTTPRFLLRLPYSTVENPIKNFNYQEDVSKDHEHFLWGNTAYLLATCLTDSFAKYRWCPNIIGPQSGGTVSDLPVHLFEA
MGQIQAKIPTEVLVTDRREFELAEEGFITLTMRKGSDNAAFFSANSVQKPKVFPNTREGKMAETNYKLGTQLPYMFIINRLAHYIKVLQREQIGSWKERQDLE
RELNTWLKQYIADQENPPTDVRSRRPLRSAQIQVLDVEGDPGWYQVAMQVRPHFKYMGASFELSLVGRLDKE

>Proteus_mirabilis_BB2000

MSLNAEAQEQQAPASTGSLLDEIMAQSRMSPETEAYDIAKQGVAAFISNIFASESEDQQINRLLIDKMLVELDNKLSTQVDEILHAPKFQEIEASWRSLKLLVDR
TDFRENIKINILHATKEELLEDFEFSPEIVQSGFYQHVYSSGYGQFGGEPVATVIGNYAFNNTAPDMKLMQYVSTVGAMAHAPFLSSVSPNFFGINSYAELPAIKD
LKSVFEGPAHTKWRSLREAEDSRYLGLTAPRFLLRLPYSPTENPVKMFNYSENVKQDHEHYLWGNTAFLLASCINDSFAKYRWCPNIIGPQSGGTVNDLPVHIF
EAMGELQTKIPTEVLVTDRREFELAEEGFITLTMRKGSDNAAFFSANSVQKPKVFPNTPEGKLAETNYKLGTQLPYMFIINRLAHYIKVLQREQIGSWKERQDL
ERELNIWLKQYVADQENPPADVRSKRPLRSAEIKVLDVDGDPGWYQVAIQVRPHFKYMGANFELSLVGRLDKE

>Proteus_mirabilis_HI4320

MSLNAEAQEQQAPASTGSLLDEIMAQSRMSPETEAYDIAKQGVAAFISNIFASESEDQQINRLLIDKMLVELDNKLSTQVDEILHAPKFQEIEASWRSLKLLVDR
TDFRENIKINILHATKEELLEDFEFSPEIVQSGFYQHVYSSGYGQFGGEPVATVIGNYAFNNTAPDMKLMQYVSTVGAMAHAPFLSSVSPNFFGINSYAELPAIKD
LKSVFEGPAHTKWRSLREAEDSRYLGLTAPRFLLRLPYSPTENPVKMFNYSENVKQDHEHYLWGNTAFLLASCINDSFAKYRWCPNIIGPQSGGTVNDLPVHIF
EAMGELQTKIPTEVLVTDRREFELAEEGFITLTMRKGSDNAAFFSANSVQKPKVFPNTPEGKLAETNYKLGTQLPYMFIINRLAHYIKVLQREQIGSWKERQDL
ERELNIWLKQYVADQENPPADVRSKRPLRSAEIKVLDVDGDPGWYQVAIQVRPHFKYMGANFELSLVGRLDKE

>Pseudomonas_aeruginosa_PAK_H2-T6SS

MSTSAAQQGRNQNGEYNILDSIIAETRLSPDDEAYDIAKRGVSAFIEELLKPQNDGEPVKKAMVDRMIAEIDAKLSTQMDEILHHPDFQALESAWRGLQLLVD
RTNFRENIKIEILSVSKQDLLDDFEDSPEVMQSGLYKHIYTAEYGQFGGQPVGAIIANYYMSPSSPDVKLMQYVSSVACMAHAPFVAAAGPKFFGLESFTGLPD
LKDLKDHFEGPQFAKWQSFRQSEDSRYVALTVPRFLLRTPYDPEENPVKSFAYKETVANSHEHYLWGNTAYAFGTKLTDSFAKYRWCPNIIGPQSGGAVEDLPL
HHFESMGEIETKIPTEVLVSDRREYELAEEGFISLTMRKGSDNAAFFSASSVQKPKFFGISAEGKAAELNYKLGTQLPYMMIVNRLAHYLKVLQREQLGSWKE
RTDLELELNKWIRQYVADQENPSAEVRGRRPLRAAQIIVSDVEGEPGWYRVSLNVRPHFKYMGADFTLSLVGKLDKE

>Pseudomonas_aeruginosa_PAK_H1-T6SS

MAELSTENLAQGQTTTEQTSEFASLLLQEFKPKTERAREAVETAVRTLAEHALEQTSLISNDAIKSIESIIAAIDAKLTAQVNLIMHHADFQQLESAWRGLHYLVN
NTETDEQLKIRVLNISKPELHKTLKKFKGTTWDQSPIFKKLYEEEYGQFGGEPYGCLVGDYYFDQSPPDVELLGEMAKISAAMHAPFISAASPTVMGMGSWQE

LSNPRDLTKIFTTPEYAGWRSLRESEDSRYIGLTMPRFLARLPYGAKTDPVEEFAFEEETDGADSSKYAWANSAYAMAVNINRSFKLYGWCSRIRGVESGGEVQ
GLPAHTFPTDDGGVDMKCPTEIAISDRREAELAKNGFMPLLHKKNTDFAAFIGAQSLQKPAEYDDPDATANANLAARLPYLFATCRFAHYLKCIVRDKIGSFKE
KDEMQRWLQDWILNYVDGDPAHSTETTKAQHPLAAAEVVVEEVEGNPGYYNSKFFLRPHYQLEGLTVSLRLVSKLPSAKEA

>Pseudomonas_aeruginosa_PAK_H3-T6SS
MPKSSAAEQSGESSTQTLSLLDEIIAKGRMAHDDSQQDYARDMLAEFATQVLDEGMAVDKDTVAMINDRISQIDALISDQLNQIIHHPELQKLEASWRGLHQL
VSNTETSARLKLRLLNVGKNELQNDLEKAVEFDQSALFKKIYEEEYGTFGGHPFSLLIGDFTFGRHPQDIGLLEKLSNVAAAAHAPFIAAASPRLFDMNSFTEL
AVPRDLTKIFESLELIKWRAFRESEDSRYVSLVLPNFLLRLPYGPETRPVEGMNYVEDVNGSDHSKYLWGNAAWVLAQRITEAFAKYGWCAAIRGAEGGGAV
EGLPAHTFRTSSGDLSLKCPTEVAITDRREKELNDLGFISLCHKKNSDVAVFFGGQTTNKARLYNTNEANANARLSAMLPYVLAASRFAHYLKVIMRDKVGSF
MTRDNVQTYLNNWIADYVLINDNAPQEIKAQYPLREARVDVSEVAGKPGAYRATVFLRPHFQLEELSASIRLVANLPPPVAA

>Pseudomonas_aeruginosa_PAO1_H2-T6SS
MSTSAAQQGRNQNGEYNILDSIIAETRLSPDDEAYDIAKRGVSAFIEELLKPQNDGEPVKKAMVDRMIAEIDAKLSTQMDEILHHPDFQALESAWRGLQLLVD
RTNFRENIKIEILSVSKQDLLDDFEDSPEVMQSGLYKHIYTAEYGQFGGQPVGAIIANYYMSPSSPDVKLMQYVSSVACMAHAPFVAAAGPKFFGLESFTGLPD
LKDLKDHFEGPQFAKWQSFRQSEDSRYVALTVPRFLLRTPYDPEENPVKSFAYKETVANSHEHYLWGNTAYAFGTKLTDSFAKYRWCPNIIGPQSGGAVEDLPL
HHFESMGEIETKIPTEVLVSDRREYELAEEGFISLTMRKGSDNAAFFSASSVQKPKFFGISAEGKAAELNYKLGTQLPYMMIVNRLAHYLKVLQREQLGSWKE
RTDLELELNKWIRQYVADQENPSAEVRGRRPLRAAQIIVSDVEGEPGWYRVSLNVRPHFKYMGADFTLSLVGKLDKE

>Pseudomonas_aeruginosa_PAO1_H1-T6SS
MAELSTENLAQGQTTTEQTSEFASLLLQEFKPKTERAREAVETAVRTLAEHALEQTSLISNDAIKSIESIIAALDAKLTAQVNLIMHHADFQQLESAWRGLHYLV
NNTETDEQLKIRVLNISKPELHKTLKKFKGTTWDQSPIFKKLYEEEYGQFGGEPYGCLVGDYYFDQSPPDVELLGEMAKISAAMHAPFISAASPTVMGMGSWQ
ELSNPRDLTKIFTTPEYAGWRSLRESEDSRYIGLTMPRFLARLPYGAKTDPVEEFAFEEETDGADSSKYAWANSAYAMAVNINRSFKLYGWCSRIRGVESGGEV
QGLPAHTFPTDDGGVDMKCPTEIAISDRREAELAKNGFMPLLHKKNTDFAAFIGAQSLQKPAEYDDPDATANANLAARLPYLFATCRFAHYLKCIVRDKIGSFK
EKDEMQRWLQDWILNYVDGDPAHSTETTKAQHPLAAAEVVVEEVEGNPGYYNSKFFLRPHYQLEGLTVSLRLVSKLPSAKEA

>Pseudomonas_aeruginosa_PAO1_H3-T6SS
MPKSSAAEQSGESSTQTLSLLDEIIAKGRMAHDDSQQDYARDMLAEFATQVLDEGMAVDKDTVAMINDRISQIDALISDQLNQIIHHPELQKLEASWRGLHQL
VSNTETSARLKLRLLNVGKNELQNDLEKAVEFDQSALFKKIYEEEYGTFGGHPFSLLIGDFTFGRHPQDIGLLEKLSNVAAAAHAPFIAAASPRLFDMNSFTEL
AVPRDLTKIFESLELIKWRAFRESEDSRYVSLVLPNFLLRLPYGPETRPVEGMNYVEDVNGTDHSKYLWGNAAWVLAQRITEAFAKYGWCAAIRGAEGGGAV
EGLPAHTFRTSSGDLSLKCPTEVAITDRREKELNDLGFISLCHKKNSDVAVFFGGQTTNKARLYNTNEANANARLSAMLPYVLAASRFAHYLKVIMRDKVGSF
MTRDNVQTYLNNWIADYVLINDNAPQEIKAQYPLREARVDVSEVAGKPGAYRATVFLRPHFQLEELSASIRLVANLPPPVAA

>Pseudomonas_aeruginosa_UCBPP-PA14_H2-T6SS
MSTSAAQQGRNQNGEYNILDSIIAETRLSPDDEAYDIAKRGVSAFIEELLKPQNDGEPVKKAMVDRMIAEIDAKLSTQMDEILHHPDFQALESAWRGLQLLVD
RTNFRENIKIEILSVSKQDLLDDFEDSPEVMQSGLYKHIYTAEYGQFGGQPVGAIIANYYMSPSSPDVKLMQYVSSVACMAHAPFVAAAGPKFFGLESFTGLPD
LKDLKDHFEGPQFAKWQSFRQSEDSRYVALTVPRFLLRTPYDPEENPVKSFAYKETVANSHEHYLWGNTAYAFGTKLTDSFAKYRWCPNIIGPQSGGAVEDLPL
HHFESMGEIETKIPTEVLVSDRREYELAEEGFISLTMRKGSDNAAFFSASSVQKPKFFGISAEGKAAELNYKLGTQLPYMMIVNRLAHYLKVLQREQLGSWKE
RTDLELELNKWIRQYVADQENPSAEVRGRRPLRAAQIIVSDVEGEPGWYRVSLNVRPHFKYMGADFTLSLVGKLDKE

>Pseudomonas_aeruginosa_UCBPP-PA14_H1-T6SS
MAELSTENLAQGQTTTEQTSEFASLLLQEFKPKTERAREAVETAVRTLAEHALEQTSLISNDAIKSIESIIAALDAKLTAQVNLIMHHADFQQLESAWRGLHYLV
NNTETDEQLKIRVLNISKPELHKTLKKFKGTTWDQSPIFKKLYEEEYGQFGGEPYGCLVGDYYFDQSPPDVELLGEMAKISAAMHAPFISAASPTVMGMGSWQ
ELSNPRDLTKIFTTPEYAGWRSLRESEDSRYIGLTMPRFLARLPYGAKTDPVEEFAFEEETDGADSSKYAWANSAYAMAVNINRSFKLYGWCSRIRGVESGGEV
QGLPAHTFPTDDGGVDMKCPTEIAISDRREAELAKNGFMPLLHKKNTDFAAFIGAQSLQKPAEYDDPDATANANLAARLPYLFATCRFAHYLKCIVRDKIGSFK
EKDEMQRWLQDWILNYVDGDPAHSTETTKAQHPLAAAEVVVEEVEGNPGYYNSKFFLRPHYQLEGLTVSLRLVSKLPSAKEA

>Pseudomonas_aeruginosa_UCBPP-PA14_H3-T6SS

MPKSSAAEQSGESSTQTLSLLDEIIAKGRMAHDDSQQDYARDMLAEFATQVLDEGMAVDKDTVAMINDRISQIDALISDQLNQIIHHPELQKLEASWRGLHQL
VSNTETSARLKLRLLNVGKNELQNDLEKAVEFDQSALFKKIYEEEYGTFGGQPFSLLIGDFTFGRHPQDIGLLEKLSNVAAAAHAPFIAAASPRLFDMNSFTEL
AVPRDLTKIFESLELIKWRAFRESEDSRYVSLVLPNFLLRLPYGPETRPVEGMNYVEDVNGTDHSKYLWGNAAWVLAQRITEAFAKYGWCAAIRGAEGGGAV
EGLPAHTFRTSSGDLSLKCPTEVAITDRREKELNDLGFISLCHKKNSDVAVFFGGQTTNKARLYNTNEANANARLSAMLPYVLAASRFAHYLKVIMRDKVGSF
MTRDNVQTYLNNWIADYVLINDNAPQEIKAQYPLREARVDVSEVAGKPGAYRATVFLRPHFQLEELSASIRLVANLPPPVAA

>Pseudomonas_fluorescens_F113_F1-T6SS

MTDNTAREGVQNLGATQETSEFASLLLQEFKPKTERAREAVETAVRTLAEQALAQTDLVSNDAIKSIESIIAAIDAKLTAQVNQVIHHPDFQQLESAWRGLHYLV
NNTESDEQLKIRVLNVSKTDLHKTLKKFKGTAWDQSPIFKKMYEEEYGQFGGEPYGCLVGDYYFDQSPPDVELLGELSKVCAAMHSPFIAAASPTVMGMGS
WQELSNPRDLTKIFTTPEYAGWRSLRESEDSRYIGLTMPRFLARLPYGAKTDPVEAFAFEENTDGADSSKYTWANAAYAMAVNINRSFKHFGWCSRIRGVESG
GEVENLPAHTFPTDDGGVDMKCPTEIAISDRREAELAKNGFMPLLHKKNTDFAAFIGAQSLQKPAEYDDPDATANANLAARLPYLFATCRFAHYLKCIVRDKI
GSFKEKDEMQRWLQDWILNYVDGDPAHSTETTKAQHPLAAAEVIVEEVEGNPGYYNSKFYLRPHYQLEGLTVSLRLVSKLPSAKGA

>Pseudomonas_fluorescens_F113_F3-T6SS

MPASSSAQTSESTTQTLSLLDKIIAEGRMAHDDSQQDYARDMLAEFATQVLDEGMAIDKDTVAMINDRISQIDQLISAQLNEVLHHPDLQKLEASWRGLHLLV
QNTETSTRLKLRLLNVTQKELQNDLEKAVEFDQSALFKKIYEEEYGTFGGHPFSLLVGDYTFGRHPQDIGLLEKLSNVAAAAHAPFIAAASPRLFDMTSFTELAI
PRDLSKIFESQELIKWRAFRESEDSRYVSLVLPHFLLRLPYGPDTLPVEGINYVEDVNGTDHSKYLWGQELIKWRAFRESEDSRYVSLVLPHFLLRLPYGPDTLP
VEGINYVEDVNGTDHSKYLWGNAAWTLAQRITEAFAKYGWCAAIRGAEGGGAVEGLPAHTFRTSSGDLSLKCPTEVAITDRREKELNDLGFISLCHKKNSDV
AVFFGGQTTNKAKLYNTNEANANARISAMLPYVLAASRFAHYLKVIMRDKVGSFMTRDNVQTYLNNWIADYVLINDNAPQEIKAQYPLREARVDVTEVAGK
PGAYKATVFLRPHFQLEELTASIRLVATLPPPVAA

>Pseudomonas_protegens_CHA0

MTELMRDNQAQPGATEQASEFASLLLQEFKPKTERAREAVETAVRTLAEQALAQTDLVSNDAIKSIESIIAAIDAKLTAQVNQVIHHPEFQKLESAWRGLHYLV
NNTETDEQLKIRVLNISKPELHKTLKKFKGTAWDQSPLFKKMYEEEYGQFGGEPYGCLVGDYYFDQSPPDVELLGELSKVCAAMHSPFIAAASPTVMGMGSW
QELSNPRDLTKIFTTPEYAGWRSLRESEDSRYIGLTMPRFLARLPYGAKTDPVEAFAFEENTDGADSSKYTWANAAYAMAVNINRSFKHFGWCSRIRGVESGGE
VENLPAHTFPTDDGGVDMKCPTEIAISDRREAELAKNGFMPLLHKKNTDFAAFIGAQSLQKPAEYDDPDATANANLAARLPYLFATCRFAHYLKCIVRDKIGSF
KEKDEMQRWLQDWILNYVDGDPAHSTETTKAQHPLAAAEVVVEEVEGNPGYYNSKFFLRPHYQLEGLTVSLRLVSKLPSAKGA

>Pseudomonas_protegens_Pf-5

MTELMRDNQAQPGATEQASEFASLLLQEFKPKTERAREAVETAVRTLAEQALAQTDLVSNDAIKSIESIIAAIDAKLTAQVNQVIHHPEFQKLESAWRGLHYLV
NNTETDEQLKIRVLNISKPELHKTLKKFKGTAWDQSPLFKKMYEEEYGQFGGEPYGCLVGDYYFDQSPPDVELLGELSKVCAAMHSPFIAAASPTVMGMGSW
QELSNPRDLTKIFTTPEYAGWRSLRESEDSRYIGLTMPRFLARLPYGAKTDPVEAFAFEENTDGADSSKYTWANAAYAMAVNINRSFKHFGWCSRIRGVESGGE
VENLPAHTFPTDDGGVDMKCPTEIAISDRREAELAKNGFMPLLHKKNTDFAAFIGAQSLQKPAEYDDPDATANANLAARLPYLFATCRFAHYLKCIVRDKIGSF
KEKDEMQRWLQDWILNYVDGDPAHSTETTKAQHPLAAAEVVVEEVEGNPGYYNSKFFLRPHYQLEGLTVSLRLVSKLPSAKGA

>Pseudomonas_putida_KT2440_K3-T6SS

MPKPSSTAITTDATADTLSNTTLLDQIMAETKLVPAQEGYQIARQGVSAFITEILKSNDPDQLINKHRADQMIAEIDRMLGKQMDAILHQPEFQQLESSWRSLKL
LVDRTDFRENIKLEILHVSKEDLLDDFENAADVTCSGIYKHVYTAGYGQFGGEPVAAMVGNYNFGPSSPDIKLLSYMASVGAMSHAPFLAAPAPEFFNLNSFE
ELPNLKEIKDLFAGPRHAKWRAFRESEDARHVALTGPRFMLRSAYHPQEQPIESFNYNEDIAGQHDNYLWGNSAFLLASCINDSFARYRWCPNIIGPQSGGAVE
DLPVHLYESLGQLQAKIPTEVLISDRKEFELAEEGFIALTMRKDSDNAAFFSANSVQKPKHFPKTPEGLQAQTNYKLGTQLPYLFIVNRLAHYIKVLQREQIGS
WKERRDLESELNKWIKQYVADQENPSADVRSRRPLRAARIEVSDVAGDPGWYQVSLAVRPHFKYMGANFEISLVGRLDTQ

>Pseudomonas_putida_KT2440_K2-T6SS

MPKPSSTAITTDATADTLSNTTLLDQIMAETKLVPAQEGYQIARQGVSAFITEILKSNDPDQLINKHRADQMIAEIDRMLGKQMDAILHQPEFQQLESSWRSLKL
LVDRTDFRENIKLEILHVSKEDLLDDFENAADVTCSGIYKHVYTAGYGQFGGEPVAAMVGNYNFGPSSPDIKLLSYMASVGAMSHAPFLAAPAPEFFNLNSFE

ELPNLKEIKDLFAGPRHAKWRAFRESEDARHVALTGPRFMLRSAYHPQEQPIESFNYNEDIAGQHDNYLWGNSAFLLASCINDSFARYRWCPNIIGPQSGGAVE
DLPVHLYESLGQLQAKIPTEVLISDRKEFELAEEGFIALTMRKDSDNAAFFSANSVQKPKHFPKTPEGLQAQTNYKLGTQLPYLFIVNRLAHYIKVLQREQIGS
WKERRDLESELNKWIKQYVADQENPSADVRSRRPLRAARIEVSDVAGDPGWYQVSLAVRPHFKYMGANFEISLVGRLDTQ

>Pseudomonas_putida_KT2440_K1-T6SS
MTDKQTAPQQAGDLVEVENFAPQPSLLDSIISQSRVARSDTERNRTRDLIGELVNQVLEGEMTPSKDLIAVLDSRIAEIDAMLSEQMNEIMHAREFQQLEASWR
GLKYQVDQTETSTTLKIHLLNASKKDLVRDLKAASEFDQSALFKKVYEEEYGTFGGAPFGMLIGDYEFNRNPEDMYLLEEISHVAAAAHAPFISAASSELFGW
DSFTEMSGPRDLAKIFDTVEYAKWKSFRASEDSRYVGLTLPHVLGRLPYGPDTTPVEEFNFVESVDGRDHNKYLWMNAAYALGTRVTDAFARYGWCVAIRGV
EGGGLVEGLPTHTFKTDDGEIALKCPTEIAITDRREKELSDLGFIPLVHCKGTDYAAFFGTQSAQKQKQYNTDIANANARLSAQLQYIFATSRIAHYMKAIMRD
KIGSFASRMDVERFLNQWLASYVLLDDTASQEAKAKFPLREARAEVFEVPGKPGVYKAVTYLRPHYQLDELTASLRLVAELPQGARG

>Pseudomonas_sp._JY-Q
MTDKQTAPQQAGDLVEVENFAPQPSLLDSIISQSRVARSDTERNRTRDLIGELVNQVLEGEMTPSKDLIAVLDSRIAEIDAMLSEQMNEIMHAREFQQLEASWR
GLKYQVDQTETSTTLKIHLLNASKKDLVRDLKAASEFDQSALFKKVYEEEYGTFGGAPFGMLIGDYEFNRNPEDMYLLEEISHVAAAAHAPFISAASSELFGW
DSFTEMSGPRDLAKIFDTVEYAKWKSFRASEDSRYVGLTLPHVLGRLPYGPDTTPVEEFNFVESVDGRDHNKYLWMNAAYALGTRVTDAFARYGWCVAIRGV
EGGGLVEGLPTHTFKTDDGEIALKCPTEIAITDRREKELSDLGFIPLVHCKGTDYAAFFGTQSAQKQKQYNTDIANANARLSAQLQYIFATSRIAHYMKAIMRD
KIGSFASRMDVERFLNQWLASYVLLDDTASQEAKAKFPLREARAEVFEVPGKPGVYKAVTYLRPHYQLDELTASLRLVAELPQGARG

>Pseudomonas_syringae_pv._actinidiae_str._Shaanxi_M228
MAKSPSAESQGSEGNTQTLTLLDKIIAEGRMAHDDSQQGYARDMLAEFATQVLDEGMAIDKDTVAMINERIGKIDSLISDQLNEILHHEDVQKLEASWRGLHA
LVKNTETGTRLKLRLLNVSQKELQTDLEKAVEFDQSALFKKIYEEEYGTFGGHPFSVLVGDFTFGRHPQDIGLLEKLSSVAAAAHAPFIAAASPRLFDMSSFTEL
SVPRDLAKVFESQELIKWRSFRESEDSRYVSLVLPHYLLRLPYGPDTNPVEGMNFVENTTGTDHSKYLWGNAAWALTQRITEAYAKYGWCAAIRGAEGGGAV
EGLPAHTFRTSSGDLSLKCPTEVAITDRREKELNDLGFISLCHKKNSDVAVFFGGQSTNKPRLYNTNEANANARISAMLPYVLAASRFAHYLKVIMRDKIGSFM
TRDNVQTYLNNWIADYVLINDNAPQEIKAQYPLREARVDVTEVPGKPGAYRATVFLRPHFQLEELTASIRLVATLPPPVAA

>Pseudomonas_syringae_pv._syringae_B728a
MTDKQPVTQQDSALVEVENFAPQSSLLDSIISESRVARSETERTRTRDLIGELVAQVLEGEMTPSKDLIAVLDARIAEIDSMLSEQMNEIMHAREFQQLEASWRG
LKYQVDQTETSTTLKIHLLNASKKDLVRDLKASSEFDQSALFKKIYEEEYGTFGGAPFGMLLGDYEFNRSPEDMYLLEEISHVAAAAHAPFISAASAELFGWDS
FTDMAGPRDLAKIFDTVEYAKWKSFRASEDSRYVGLTLPHVLGRLPYGPDTTPVEEFNFVESVDGRDHNKYLWMNAAYALGTRVTDAFSRYGWCVAIRGVE
GGGLVEGLPTHTFKTDDGEIALKCPTEIAITDRREKELSDLGFIPLVHCKGTDYAAFFGTQSTQKQKQYNTDIANANARLSAQLQYIFATSRIAHYMKAIMRDKI
GSFASRKDVELFLNKWLSSYVLLDDTASQEAKAKFPLREARAEVFEVPGKPGVYKAVTYLRPHYQLDELTASLRLVAELPQSTRG

>Pseudomonas_syringae_pv._tomato_str._DC3000
MPSNSSAKQKSVETSEHSILDSIIAQTSLTRDDEAYDIAKRGVSAFIEELLKPQNNGEPVKKAMVDRMIAEIDAKLSRQMDEILHHPDFQSLESSWRGLQLLVDR
TNFRENIKIEILNVSKDDLLEDFEDSPEVMQSGLYKHIYTAEYGQFGGQPVGAIIANYFMSPSSPDVKLMQYVSSVACMSHAPFIAAAGPKFFGLESFTGLPDLK
DLKDHFEGPQFAKWQSFRQSEDSRYVGLTVPRFLLRNPYDPQENPVKSFVYKETVANSHEHYLWGNTAYAFGTKLTDSFAKFRWCPNIIGPQSGGAVEDLPLH
HFESMGEIETKIPTEVLVSDRREYELAEEGFISLTMRKGSDNAAFFSASSVQKPKFFGNSAEGKAAELNYKLGTQLPYMMIVNRLAHYLKVLQREQLGSWKER
TDLELELNKWIRQYVADQENPSAEVRGRRPLRAAQITVSDVEGEPGWYRVSLNVRPHFKYMGADFTLSLVGKLDKE

>Ralstonia_solanacearum_GMI1000
MLDNQSAALAQSAPAAESVNLLDEIVEQSRVAKSDAEHARAKDIIGELVNQVLQGTVVVSDNLSATLDARVAELDRLISSQLSAVMHTPEFQKLEGTWRGLSY
LVKETQTSTMLKIKVLNATKRDLVRDFKTAIDFDQSALFKKVYEEEFGTFGGAPFGTLIGDFEVTRQPEDVYFIEQMSHVAAAAHAPFIASSSPELFGLESFADL
GKPRDLAKVFDTVEYAKWKSFRESEDSRYVGLTMPRFLGRLPYNPKDGTTVEGFNFVEDVDGTDHSKYLWCNAAWAFGARLTAAFEDFGWCAAIRGVEGG
GLVEDLPTHTFKTDDGEIALKCPTEIAITDRREKELSDLGFIPLVHCKNSDYAAFFAAQSVQRPKKYDTDSANANAVLSAQLQYIFSVSRVAHYLKAMMRDKIG
SFASAKNVETFLNRWISQYVLLDDNATQEQKAQFPLREASIQVSEVPGKPGTYRSVAFLRPHFQLDELSISLRLVADLPKPTNS

>Rhizobium_etli_bv._mimosae_str._Mim1

MTAETQLQTKEDVGFAPDENLLSKVVAATRQTEPDRAQDLLRTLTDQALKGTVTYDRNLTITLNNAIAEIDKAISGQLAAIMQSPEFTKLEGAWRGLNYLVKN
SETSANLKIRVLNASKRDVAKDLAKAVEFDQSRLFKSVYEDEFGTPGGEPMGALIGDYEFDNSFDDVQLLQGVSMIAAAAFAPFISAASPRMFGFEDFRELAKP
RDLEKIFETVEYAKWRSLRDSDDSRFVTLAMPRVLARMPYGPKTSPIEEFNFDETGGSKTGELSHGSYCWMNATYVMGTRLTEAFSKNGWCTAIRGAENGGK
VEGLPMHIFSSDDGDLDLKCPTEVGITDRRDAELGKLGFLPLCHYKNTDYAVFFGAQTAHKPKLYDRPEATANAAVSARLPYMMATSRFAHYLKVMGRDKIG
SFMEAEDCEAWLNRWISNYVNANDDAGEESRAKYPLREAKVTVQEIPGRPGAYNAVAWMRPWLQMEELTTSLRMVARIPSKN

>Rhizobium_leguminosarum_bv._trifolii

MTAETQLQTKEDVGFAPDENLLSKVVAATRQTEPDRAQDLLRTLTDQALKGTVTYDRNLTITLNNAIAEIDKAISGQLAAIMQSPEFTKLEGAWRGLNYLVKN
SETSANLKIRVLNASKRDVAKDLAKAVEFDQSRLFKSVYEDEFGTPGGEPMGALIGDYEFDNSFDDVQLLQGVSMIAAAAFAPFISAASPRMFGFEDFRELAKP
RDLEKIFETVEYAKWRSLRDSDDSRFVTLAMPRVLARMPYGPKTSPIEEFNFDETGGSKTGELSHGSYCWMNATYVMGTRLTEAFSKNGWCTAIRGAENGGK
VEGLPMHIFSSDDGDLDLKCPTEVGITDRRDAELGKLGFLPLCHYKNTDYAVFFGAQTAHKPKLYDRPEATANAAVSARLPYMMATSRFAHYLKVMGRDKIG
SFMEAEDCEAWLNRWISNYVNANDDAGEESRAKYPLREAKVTVQEIPGRPGAYNAVAWMRPWLQMEELTTSLRMVARIPSKN

>Salmonella_entericasub_sp._enterica_serovar_Dublinstr._CT_02021853

MANSNMQATDAVAQDTASASGEFDALLNQAFRPKTTQAAKAVEAAVQTLAQQALANTITVSDDAYKSISAIIAQIDFKLTEQIKLILQHPDWQKLESSWRGME
HLVYNTETDEKLKIRFMNLSKDELRRNMKRYKGIAWDQSPMFKKLYEAEYGQLGGEPYGCIIADYYFDHTPPDVDLLGSIAKVAASAHAPFIAGASPSVLQM
DSWQELANPRDLTKIVTQNLEYAPWNSLRASEDSRYIGLTMPRFLARLPYGAKTNPVDEFDFEEDADGSDHTKYVWSNAAYAMGVNINRSFKHYGWCTLIRG
VESGGAVENLPCHTFPTDDGGVDMKCPTEIAISDRREAELAKNGFIPLIHRKNSDYAAFIGAQSLQKPQEYYDPDATANANLSARLPYLFACSRFAHFLKCIVRD
KIGSFKEREDMQRWLNEWIMNYVDADPVNSSQETKARRPLAAAEVVVEEVEGNPGYYDAKFFLRPHFQLEGLTGSLRLVTKLPSVKQGNA

>Salmonella_enterica_subsp._enterica_serovar_Gallinarum_str_.287/91

MSLQEEELVSTRTEQGSHAPSLLDEIMAQTRIQPGADGYDIARQGVTAFIASLLQSTSSAEPVNKLAVDSMIADIDARISRQMDAIIHAPQFQQLESFWRSMKAL
VDRVDFRENIKVHVLHVTKAELLEDFEFAPEITQSGFYKHVYSSGFGQFGGEPIAAVLGAYEFKNTTPDMKLLNYVSAVGAMAHAPFLSSVSPEFMGLKSWTE
LPNIKDLYAIFEGPAYTKWRALRDSEDSRYLGLTAPRFLLRQPYSATENPIKSFNYAEDVSHSHEHYLWGNTAWMLASNIADSFAKYRWAPNIIGPQSGGAVKDL
PVHLYESMGQIQAKIPTEVLITDRREFELAEEGFITLTMRKNSDNAAFFSANSVQKPKHFPGKDAETNYKLGTQLPYLFIINRLAHYIKVLQREQLGSWKERSDL
ERELNIWVRQYVADQENPPADVRSRKPLRGAKVEVLDVDGEPGWYQVAISARPHFKYMGASFELSLVGRLDK

>Salmonella_enterica_subsp._enterica_serovar_Typhimurium

MANSNMQATDAVAQDTASASGEFDALLNQAFRPKTTQAAKAVEAAVQTLANTITVSDDAYKSISAIIAQIDFKLTEQIKLILQHPDWQKLESSWRGMEHLVYN
TETDEKLKIRFMNLSKDELRRNMKRYKGIAWDQSPMFKKLYEAEYGQLGGEPYGCIIADYYFDHTPPDVDLLGSIAKVAASAHAPFIAGASPSVLQMDSWQE
LANPRDLTKIVTQNLEYAPWNSLRASEDSRYIGLTMPRFLARLPYGAKTNPVDEFDFEEDADGSDHTKYVWSNAAYAMGVNINRSFKHYGWCTLIRGVESGG
AVENLPCHTFPTDDGGVDMKCPTEIAISDRREAELAKNGFIPLIHRKNSDYAAFIGAQSLQKPQEYYDPDATANANLSARLPYLFACSRFAHFLKCIVRDKIGSF
KEREDMQRWLNEWIMNYVDADPVNSSQETKARRPLAAAEVVVEEVEGNPGYYDAKFFLRPHFQLEGLTGSLRLVTKLPSVKQGNA

>Salmonella_enterica_subsp._enterica_serovar_Typhimurium_str._14028S

MANSNMQATDAVAQDTASASGEFDALLNQAFRPKTTQAAKAVEAAVQTLANTITVSDDAYKSISAIIAQIDFKLTEQIKLILQHPDWQKLESSWRGMEHLVYN
TETDEKLKIRFMNLSKDELRRNMKRYKGIAWDQSPMFKKLYEAEYGQLGGEPYGCIIADYYFDHTPPDVDLLGSIAKVAASAHAPFIAGASPSVLQMDSWQE
LANPRDLTKIVTQNLEYAPWNSLRASEDSRYIGLTMPRFLARLPYGAKTNPVDEFDFEEDADGSDHTKYVWSNAAYAMGVNINRSFKHYGWCTLIRGVESGG
AVENLPCHTFPTDDGGVDMKCPTEIAISDRREAELAKNGFIPLIHRKNSDYAAFIGAQSLQKPQEYYDPDATANANLSARLPYLFACSRFAHFLKCIVRDKIGSF
KEREDMQRWLNEWIMNYVDADPVNSSQETKARRPLAAAEVVVEEVEGNPGYYDAKFFLRPHFQLEGLTGSLRLVTKLPSVKQGNA

>Salmonella_enterica_subsp._enterica_serovar_Typhimurium_str._LT2

MANSNMQATDAVAQDTASASGEFDALLNQAFRPKTTQAAKAVEAAVQTLAQQALANTITVSDDAYKSISAIIAQIDFKLTEQIKLILQHPDWQKLESSWRGME
HLVYNTETDEKLKIRFMNLSKDELRRNMKRYKGIAWDQSPMFKKLYEAEYGQLGGEPYGCIIADYYFDHTPPDVDLLGSIAKVAASAHAPFIAGASPSVLQM
DSWQELANPRDLTKIVTQNLEYAPWNSLRASEDSRYIGLTMPRFLARLPYGAKTNPVDEFDFEEDADGSDHTKYVWSNAAYAMGVNINRSFKHYGWCTLIRG

VESGGAVENLPCHTFPTDDGGVDMKCPTEIAISDRREAELAKNGFIPLIHRKNSDYAAFIGAQSLQKPQEYYDPDATANANLSARLPYLFACSRFAHFLKCIVRD
KIGSFKEREDMQRWLNEWIMNYVDADPVNSSQETKARRPLAAAEVVVEEVEGNPGYYDAKFFLRPHFQLEGLTGSLRLVTKLPSVKQGNA

>Salmonella_enterica_subsp._enterica_serovar_Typhimurium_str._SL1344
MANSNMQATDAVAQDTASASGEFDALLNQAFRPKTTQAAKAVEAAVQTLANTITVSDDAYKSISAIIAQIDFKLTEQIKLILQHPDWQKLESSWRGMEHLVYN
TETDEKLKIRFMNLSKDELRRNMKRYKGIAWDQSPMFKKLYEAEYGQLGGEPYGCIIADYYFDHTPPDVDLLGSIAKVAASAHAPFIAGASPSVLQMDSWQE
LANPRDLTKIVTQNLEYAPWNSLRASEDSRYIGLTMPRFLARLPYGAKTNPVDEFDFEEDADGSDHTKYVWSNAAYAMGVNINRSFKHYGWCTLIRGVESGG
AVENLPCHTFPTDDGGVDMKCPTEIAISDRREAELAKNGFIPLIHRKNSDYAAFIGAQSLQKPQEYYDPDATANANLSARLPYLFACSRFAHFLKCIVRDKIGSF
KEREDMQRWLNEWIMNYVDADPVNSSQETKARRPLAAAEVVVEEVEGNPGYYDAKFFLRPHFQLEGLTGSLRLVTKLPSVKQGNA

>Salmonella_enterica_subsp._enterica_serovar_Typhimurium_strain_ATCC_14028
MANSNMQATDAVAQDTASASGEFDALLNQAFRPKTTQAAKAVEAAVQTLANTITVSDDAYKSISAIIAQIDFKLTEQIKLILQHPDWQKLESSWRGMEHLVYN
TETDEKLKIRFMNLSKDELRRNMKRYKGIAWDQSPMFKKLYEAEYGQLGGEPYGCIIADYYFDHTPPDVDLLGSIAKVAASAHAPFIAGASPSVLQMDSWQE
LANPRDLTKIVTQNLEYAPWNSLRASEDSRYIGLTMPRFLARLPYGAKTNPVDEFDFEEDADGSDHTKYVWSNAAYAMGVNINRSFKHYGWCTLIRGVESGG
AVENLPCHTFPTDDGGVDMKCPTEIAISDRREAELAKNGFIPLIHRKNSDYAAFIGAQSLQKPQEYYDPDATANANLSARLPYLFACSRFAHFLKCIVRDKIGSF
KEREDMQRWLNEWIMNYVDADPVNSSQETKARRPLAAAEVVVEEVEGNPGYYDAKFFLRPHFQLEGLTGSLRLVTKLPSVKQGNA

>Serratia_marcescens_RM66262
MSNSPQQQNALQTTETFSSDEFSALLNKEFRPKTDQAKEAVENAVKTLAQQALENTVTVSSDAYRTIQALIAEIDEKLSQQINQIIHHDDFQKLEGAWHGLHYL
VNNSETDEMLKIRFMSISKQELGRTLKRYKGVGWDQSPIFKKVYEEEYGQFGGEPFGCLVGDYYFDHSPQDVELLGEMAKIGAASHCPFIAGTAPSVMQMES
WQELSNPRDLTKIFQNTEYAAWRSLRESEDARYLGLVMPRFLARLPYGIRTNPVDEFDFEEDTDGATHGNYTWTNAAYAMAANINRSFKEFGWCTAIRGVESG
GAVENLPCHTFPSDDGGVDMKCPTEIAISDRREAELAKNGFMPLVHRKNSDFAAFIGAQSLQKPAEYYDADASANAQLSARLPYLFACCRFAHYLKCIVRDKI
GSFRERDDMERWLNDWIMNYVDGDPANSSQETKSRKPLAAAEVQVEEIEDNPGYYSAKFFLRPHYQLEGLTVSLRLVSKLPSLKQNDAS

>Serratia_marcescens_strain_KZ11_T6SS-2
MSNSPQQQNALQTTETFSSDEFSALLNKEFRPKTDQAKEAVENAVKTLAQQALENTVTVSSDAYRTIQALIAEIDEKLSQQINQIIHHDDFQKLEGAWHGLHYL
VNNSETDEMLKIRFMSISKQELGRTLKRYKGVGWDQSPIFKKIYEEEYGQFGGEPFGCLVGDYYFDHSPQDVELLGEMAKIGAASHCPFIAGTAPSVMQMESW
QELSNPRDLTKIFQNTEYAAWRSLRESEDARYLGLVMPRFLARLPYGIRTNPVDEFDFEEDTDGATHGNYTWTNAAYAMAANINRSFKEFGWCTAIRGVESGG
AVENLPCHTFPSDDGGVDMKCPTEIAISDRREAELAKNGFMPLVHRKNSDFAAFIGAQSLQKPAEYYDADASANAQLSARLPYLFACCRFAHYLKCIVRDKIG
SFRERDDMERWLNDWIMNYVDGDPANSSQETKSRKPLAAAEVQVEEIEDNPGYYSAKFFLRPHYQLEGLTVSLRLVSKLPSLKQNDAS

>Serratia_marcescens_strain_KZ11_T6SS-1
MRARQRRYSAKQQATAVAEVETQDTENELDGLLKRSFRPRTNEASEAVRRAIGTLSEYANKGKVKVSQDVVLTIESLIAQIDEQLSQQMNNILHHKEFQKLESA
WQGLSYLVDNTNVSETLKIRVLNISQDELTRNLRRYRGSAWDQSPVFKQIYEQEYGQFGGEPFGCIIGDFEFDHSPMSVTLLTELAKISAASHCPFISAASPSLLQ
MSKWNELGNPRDIGKIFTTPEYASWRRLRESNDSRYLVLTMPRFLSRLPYGAKTNPIEEFAFEESVRPDVDDDFSWANSAYAMGVNINRAFHEYGWCSKIRGIE
SGGSVEELPAYAFPSDEGGYELTCPTEVAISDRREQELSDAGFLPLVYRKHSDFAAFIGSCTMHAPAKYEDPDATANAKLSSRLPYIFATCRFAHYLKCIVRDKIG
SFRSRDDMQLWLNDWLMNYVDGDPSVSTEATKARRPLAAAEVRVEDVEDDPGYYRAHFYLRPHYQLEGMTVSLRLVSKLPSAKKDGSR

>Serratia_marcescens_subsp._marcescens_Db11
MSNSPQQQNALQTTETFSSDEFSALLNKEFRPKTDQAKEAVENAVKTLAQQALENTVTVSSDAYRTIQALIAEIDEKLSQQINQIIHHDDFQKLEGAWHGLHYL
VNNSETDEMLKIRFMSISKQELGRTLKRYKGVGWDQSPIFKKVYEEEYGQFGGEPFGCLVGDYYFDHSPQDVELLGEMAKIGAASHCPFIAGTAPSVMQMES
WQELSNPRDLTKIFQNTEYAAWRSLRESEDARYLGLVMPRFLARLPYGIRTNPVDEFDFEEDTDGATHGNYTWTNAAYAMAANINRSFKEFGWCTAIRGVESG
GAVENLPCHTFPSDDGGVDMKCPTEIAISDRREAELAKNGFMPLVHRKNSDFAAFIGAQSLQKPAEYYDADASANAQLSARLPYLFACCRFAHYLKCIVRDKI
GSFRERDDMERWLNDWIMNYVDGDPANSSQETKSRKPLAAAEVQVEEIEDNPGYYSAKFFLRPHYQLEGLTVSLRLVSKLPSLKQNDAS

>Serratia_sp._FS14

MRARQRRYSAKQQATAVAEVETQDTENELDGLLKRSFRPRTNEASEAVRRAIGTLSEYANKGKVKVSQDVVLTIESLIAQIDEQLSQQMNNILHHKEFQKLESA
WQGLSYLVDNTNVSETLKIRVLNISQDELTRNLRRYRGSAWDQSPVFKQIYEQEYGQFGGEPFGCIIGDFEFDHSPMSVTLLTELAKISAASHCPFISAASPSLLQ
MSKWNELGNPRDIGKIFTTPEYASWRRLRESNDSRYLVLTMPRFLSRLPYGAKTNPIEEFAFEESVRPDVDDDFSWANSAYAMGVNINRAFHEYGWCSKIRGIE
SGGSVEELPAYAFPSDEGGYELTCPTEVAISDRREQELSDAGFLPLVYRKHSDFAAFIGSCTMHAPAKYEDPDATANAKLSSRLPYIFATCRFAHYLKCIVRDKIG
SFRSRDDMQLWLNDWLMNYVDGDPSVSTEATKARRPLAAAEVRVEDVEDDPGYYRAHFYLRPHYQLEGMTVSLRLVSKLPSAKKDGSR

>Vibrio_alginolyticus_12G01_T6SS-1
MSAEAQAPEQEAALVESGSLLDSILNETRLKPSDEGFDVAKRGVEAFISELLSSSNTEKVDQSLVDLMISEIDQKLSKQVDAILHNEEVQAIESTWRGLKYLVDH
TDFRENIQIELISAKKDEVLDDFEDAPEVVKSGLYKQIYTREYGQFGGKPVGAVICDYAMSASSPDIKLMEYMANVGAMSHAPFITSASAKFFGLDSYEELPNL
KDLKSVFEGPQYAKWRGLREHEDARYLGLCTSRFMLRTPYSVEDNPIKAFDYDEQVTDSHDNYLWGNSAYAMASKISESFAKYRWCPNIIGPQSGGTVQDLP
VYNFEAMGQIETKIPTEILVSDRREYELAEEGFIALTMRKGSDNAAFFSANSVQKPKVFANTPEGKQAEMNYKLGTQLPYMFIINRLAHYIKVLQREQIGSWKE
RSDLEIELNKWIRQYVSDQENPPAEVRGRRPLRAAKVEVSDVEGDPGWYKVSMSVRPHFKYMGASFDLSLVGKLDQ

>Vibrio_alginolyticus_12G01_T6SS-2
MTTEAEKQLDPTVNEQTTSFLDQAIGATKQTEASRAEELIKTLAEEAMKGTVTWNKNLTVTFREAINLLDRQISDQLSEVMHHPELQKLEGSWRGLNYLVMN
SETSSTLKVRMMSMTKKELHKDLSKAVEFDQSQIFKKVYESEFGSAGGEPYGALIGDYEFTNHPEDIESLRLMSNVAASGFSPFLSAASPALFGFDEWTELSKP
RDLDKVFESLEYAQWRSFRESADSRFVSLTMPKVLARLPYGQATAPVEEFGFEEFEVDPVSGIAVNAEHNDYCWMNSSYVLGVKLTDAFSKYGFCTAIRGAEG
GGRVDNLPTHFFMSDDGDPDMKCPTEIGITDRREAELGKLGFLPLCHYKNTNYAVFFGAQTCQKPANHESPEATANAAISARLPYMMATSRFAHYLKVMARD
KIGSFMEAEDVESWLNRWILGYVNASEGGGQEIRAKYPLADARVQVKEIPGAPGSYNAVAWLKPWLQMEELTTSLRLVAKIPQSGG

>Vibrio_anguillarum_strain_MHK3
MSTTEAVLERQSATQGSLLDEIMAQTRIAPTEEGYDVAKKGVAAFIENLIGSNQTEEPVNKSLVDQMLVELDKKISAQMDEILHEERFQQMESSWRGLKLFIDR
TDFRENNKVDLLHVTKGELLEDFEFAPETTQSGLYKHVYSSGYGQFGGEPTGAIIGNFAFTPSTPDMKLLQYMGALGAMAHAPFISSVGPEFFGIDSFEELPNIK
DLKSTFESPKYTKWRSLRESEDARYLGLTAPRFLLRVPYDPTENPIKSFNYVESVSASHEHYLWGNTAFAFATRLTDSFAKYRWCPNIIGPQSGGAVEDLPVHVF
ESMGALQSKIPTEVLVTDRKEFELAEEGFIALTMRKGSDNAAFFSANSIQKPKIFPNTKEGKEAETNYKLGTQLPYMMIINRLAHYVKVLQREQIGAWKERQD
LERELNGWIKQYVADQENPPADVRSRRPLRAAKIEVMDVEGEPGWYQVSLSVRPHFKYMGANFELSLVGRLDQA

>Vibrio_cholerae_2740-80
MSTTEKVLERPQLAQGSLLDEIMAQTRIAPSEEGYDIAKKGVAAFIENLMGSQHSAEPVNKSLVDQMLVELDKKISAQMDEILHNSQFQAMESAWRGLKLFV
DRTDFRENNKVEILHVTKDELLEDFEFAPETAQSGLYKHVYSAGYGQFGGEPVGAIIGNYAFTPSTPDMKLLQYMGALGAMAHAPFISSVGPEFFGIDSFEELP
NIKDLKSTFESPKYTKWRSLRESEDARYLGLTAPRFLLRVPYDPIENPVKSFNYAENVSASHEHYLWGNTAFAFATRLTDSFAKYRWCPNIIGPQSGGAVEDLPV
HVFESMGALRFLLRVPYDPIENPVKSFNYAENVSASHEHYLWGNTAFAFATRLTDSFAKYRWCPNIIGPQSGGAVEDLPVHVFESMGALQSKIPTEVLITDRKEF
ELAEEGFIALTMRKGSDNAAFFSANSIQKPKVFPNTKEGKEAETNYKLGTQLPYMMIINRLAHYVKVLQREQIGAWKERQDLERELNSWIKQYVADQENPPA
DVRSRRPLRAARIEVMDVEGNPGWYQVSLSVRPHFKYMGANFELSLVGRLDQA

>Vibrio_cholerae_C6706
MSTTEKVLERPQLAQGSLLDEIMAQTRIAPSEEGYDIAKKGVAAFIENLMGSQHSAEPVNKSLVDQMLVELDKKISAQMDEILHNSQFQAMESAWRGLKLFV
DRTDFRENNKVEILHVTKDELLEDFEFAPETAQSGLYKHVYSAGYGQFGGEPVGAIIGNYAFTPSTPDMKLLQYMGALGAMAHAPFISSVGPEFFGIDSFEELP
NIKDLKSTFESPKYTKWRSLRESEDARYLGLTAPRFLLRVPYDPIENPVKSFNYAENVSASHEHYLWGNTAFAFATRLTDSFAKYRWCPNIIGPQSGGAVEDLPV
HVFESMGALQSKIPTEVLITDRKEFELAEEGFIALTMRKGSDNAAFFSANSIQKPKVFPNTKEGKEAETNYKLGTQLPYMMIINRLAHYVKVLQREQIGAWKE
RQDLERELNSWIKQYVADQENPPADVRSRRPLRAARIEVMDVEGNPGWYQVSLSVRPHFKYMGANFELSLVGRLDQA

>Vibrio_cholerae_DL4211
MSTTEKVLERPQLAQGSLLDEIMAQTRIAPSEEGYDIAKKGVAAFIENLMGSQHSAEPVNKSLVDQMLVELDKKISAQMDEILHNSQFQAMESAWRGLKLFV
DRTDFRENNKVEILHVTKDELLEDFEFAPETAQSGLYKHVYSAGYGQFGGEPVGAIIGNYAFTPSTPDMKLLQYMGALGAMAHAPFISSVGPEFFGIDSFEELP
NIKDLKSTFESPKYTKWRSLRESEDARYLGLTAPRFLLRVPYDPIENPVKSFNYAENVSASHEHYLWGNTAFAFATRLTDSFAKYRWCPNIIGPQSGGAVEDLPV

HVFESMGALQSKIPTEVLITDRKEFELAEEGFIALTMRKGSDNAAFFSANSIQKPKVFPNTKEGKEAETNYKLGTQLPYMMIINRLAHYVKVLQREQIGAWKE
RQDLERELNSWIKQYVADQENPPADVRSRRPLRAARIEVMDVEGNPGWYQVSLSVRPHFKYMGANFELSLVGRLDQA

>Vibrio_cholerae_DL4215
MSTTEKVLERPQLAQGSLLDEIMAQTRIAPSEEGYDIAKKGVAAFIENLMGSQHSAEPVNKSLVDQMLVELDKKISAQMDEILHNSQFQAMESAWRGLKLFV
DRTDFRENNKVEILHVTKDELLEDFEFAPETAQSGLYKHVYSAGYGQFGGEPVGAIIGNYAFTPSTPDMKLLQYMGALGAMAHAPFISSVGPEFFGIDSFEELP
NIKDLKSTFESPKYTKWRSLRESEDARYLGLTAPRFLLRVPYDPIENPVKSFNYAENVSASHEHYLWGNTAFAFATRLTDSFAKYRWCPNIIGPQSGGAVEDLPV
HVFESMGALQSKIPTEVLITDRKEFELAEEGFIALTMRKGSDNAAFFSANSIQKPKVFPNTKEGKEAETNYKLGTQLPYMMIINRLAHYVKVLQREQIGAWKE
RQDLERELNSWIKQYVADQENPPADVRSRRPLRAARIEVMDVEGNPGWYQVSLSVRPHFKYMGANFELSLVGRLDQA

>Vibrio_cholerae_O1_biovar_El_Tor_str._N16961
MSTTEKVLERPQLAQGSLLDEIMAQTRIAPSEEGYDIAKKGVAAFIENLMGSQHSAEPVNKSLVDQMLVELDKKISAQMDEILHNSQFQAMESAWRGLKLFV
DRTDFRENNKVEILHVTKDELLEDFEFAPETAQSGLYKHVYSAGYGQFGGEPVGAIIGNYAFTPSTPDMKLLQYMGALGAMAHAPFISSVGPEFFGIDSFEELP
NIKDLKSTFESPKYTKWRSLRESEDARYLGLTAPRFLLRVPYDPIENPVKSFNYAENVSASHEHYLWGNTAFAFATRLTDSFAKYRWCPNIIGPQSGGAVEDLPV
HVFESMGALQSKIPTEVLITDRKEFELAEEGFIALTMRKGSDNAAFFSANSIQKPKVFPNTKEGKEAETNYKLGTQLPYMMIINRLAHYVKVLQREQIGAWKE
RQDLERELNSWIKQYVADQENPPADVRSRRPLRAARIEVMDVEGNPGWYQVSLSVRPHFKYMGANFELSLVGRLDQA

>Vibrio_cholerae_strain_A1552
MSTTEKVLERPQLAQGSLLDEIMAQTRIAPSEEGYDIAKKGVAAFIENLMGSQHSAEPVNKSLVDQMLVELDKKISAQMDEILHNSQFQAMESAWRGLKLFV
DRTDFRENNKVEILHVTKDELLEDFEFAPETAQSGLYKHVYSAGYGQFGGEPVGAIIGNYAFTPSTPDMKLLQYMGALGAMAHAPFISSVGPEFFGIDSFEELP
NIKDLKSTFESPKYTKWRSLRESEDARYLGLTAPRFLLRVPYDPIENPVKSFNYAENVSASHEHYLWGNTAFAFATRLTDSFAKYRWCPNIIGPQSGGAVEDLPV
HVFESMGALQSKIPTEVLITDRKEFELAEEGFIALTMRKGSDNAAFFSANSIQKPKVFPNTKEGKEAETNYKLGTQLPYMMIINRLAHYVKVLQREQIGAWKE
RQDLERELNSWIKQYVADQENPPADVRSRRPLRAARIEVMDVEGNPGWYQVSLSVRPHFKYMGANFELSLVGRLDQA

>Vibrio_cholerae_V52
MSTTEKVLERPQLAQGSLLDEIMAQTRIAPSEEGYDIAKKGVAAFIENLMGSQHSAEPVNKSLVDQMLVELDKKISAQMDEILHNSQFQAMESAWRGLKLFV
DRTDFRENNKVEILHVTKDELLEDFEFAPETAQSGLYKHVYSAGYGQFGGEPVGAIIGNYAFTPSTPDMKLLQYMGALGAMAHAPFISSVGPEFFGIDSFEELP
NIKDLKSTFESPKYTKWRSLRESEDARYLGLTAPRFLLRVPYDPIENPVKSFNYAENVSASHEHYLWGNTAFAFATRLTDSFAKYRWCPNIIGPQSGGAVEDLPV
HVFESMGALQSKIPTEVLITDRKEFELAEEGFIALTMRKGSDNAAFFSANSIQKPKVFPNTKEGKEAETNYKLGTQLPYMMIINRLAHYVKVLQREQIGAWKE
RQDLERELNSWIKQYVADQENPPADVRSRRPLRAARIEVMDVEGNPGWYQVSLSVRPHFKYMGANFELSLVGRLDQA

>Vibrio_coralliilyticus_OCN008_T6SS-1
MTAEAQAPEQEAALAESGSLLDSILNETRLKPSDEGFDVAKRGVEAFIGELLSNNSAEKVDQSLVDLMIGEIDQKLSKQVDAILHNEEVQAIESTWRGLKYLVD
HTDFRENILIELISAKKDEMLDDFEDAPEVVKSGLYKQIYTREYGQFGGKPVGAVICDYQLTSSSPDIKLMEYMANVGAMSHAPFITSASSKFFGLDSYEELPN
MKDLKSVFEGPQYTKWRGLREHEDSRYLGLCTSRFMLRNPYSVEDNPIKAFDYDELVTDDHNNFLWGNSAYAMASKISESFAKYRWCPNIIGPQSGGAVFDLP
VYNYESMGQIETKIPTEILVSDRREFELAEEGFIALTMRKGSDNAAFFSANSIQKPKVYANTPEGKNAEMNYKLGTQLPYMFIINRLAHYIKVLQREQIGSWKE
RSDLEIELNKWIRQYVSDQENPPAEVRGRRPLRAAKVEVSDVEGDPGWYKVSMSVRPHFKYMGASFDLSLVGKLDQ

>Vibrio_coralliilyticus_OCN008_T6SS-2
MSTEAENQTQTEAAEGSLSFLDRAIEATTQTPADTTKELFSVLAEQALSGTVTWDKNVTKTIENAISEIDKKLSKQLSEVMQQKDLQKLEGSWRGLQKLVKES
ELGRDLKIKMVDFSQEELLDQFEDAPAIDRSPLFNAVYQGEFGTAGGEPYGTFIGDYEFSAKDEDVALLRYMGEVAAACHAPFVAAANAQMFEFNDFTTFDEG
KPVAAGFDSPAYAAWNAFRESDDARYVTLTLPRTLARLPYGAKGLGTELFDYEELGTDMDGNPNPENNDQLVWSNAAYDLGLKMTQAYTASGWCTSIRGLD
NGGKVENLPNLTYKTEAGDLVQQCPTEVNLTDEREKELSDLGFLPLVHYKNSNYGVFIGGQTTQKPKTYTDPDATANAAISARLPYIMASSRIAHYLKVMGRD
KLGSNLEAPDIQRELQLWIDQYTNAGAIGNEQRAKTPLCESRIEVVEQPGRPGSYSAVAHLRPWLQLEELTTSVRMVAKIPG

>Vibrio_fluvialis_85003_T6SS-2

MSTTETVLERPQLAEGSLLDEIMAQTRIAPSEEGYDVAKKGVAAFIENLIGGNQLDEAVNKSLVDQMLVELDKKISAQMDEILHDEKYQQMESAWRGLKLFV
DRTDFRENNKVDLLHVTKEELLEDFEFAPETTQSGLYKHVYSSGYGQFGGEPTGAIIGNFAFTPSTPDMKLLQYMGALGAMAHAPFISSVGPQFFGIDSFEELP
NIKDIKSTFESPKYTKWRALRESEDARYIGLTAPRFLLRVPYDPTENPIKSFNYSENVSQSHEHYLWGNTAFAFATRLTDSFAKYRWCPNIIGPQSGGAVEDLPVH
VFESMGALQSKIPTEVLITDRKEFELAEEGFIALTMRKGSDNAAFFSANSIQKAKIFPNTKEGKEAETNYKLGTQLPYMMIINRLAHYVKVLQREQIGAWKERQ
DLERELNAWIKQFVADQENPPADVRSRRPLRAAKIEVMDVEGNPGWYQVSLAVRPHFKYMGANFELSLVGRLDQA

>Vibrio_fluvialis_85003_T6SS-1
MSTELENQVSPEAGTESLSLLDRAIAATNQTPADQTKELFSVLAEQALSGTVTWDKNLTKTIENAIAEIDKQMSRQLSAIMQQDDFRRLEGSWRGLNKLVTESE
TGQSLKIKLVDFTKDELIEQFEDAPAVDRSPLFDALYQKEFGTAGGEPYGALIGDYTFSHKDEDVALMRYMGETAAASHAPFIAAANPEMFEFDSFETFNEGKP
VASGFDSPAYASWNAFRESDDSRYVVLTLPQTLARLPYGDKGLGTKSFAYEELATDADGNPKPKNNAQLVWSNAAYELGLKMTQAHTQFGWCTAIRGLDNG
GKVENLPNLTYKSEAGDLMQQCPIEVNLTDEREKELSDLGFLPLVHYKNTNYGVFIGSQTAQKPKTYTDPDATGNAAISARLPYIMASSRIAHYLKVMGRDML
GSNLEAADVQKDLQTWIDQYTNSGAVGNAERSKTPLCESRIQVVEQPGRPGAYSAVAHLRPWLQLEELTTSVRMVTKIPG

>Vibrio_parahaemolyticus_BB22OP_T6SS-1
MSAEAQAPEQEAALVESGSLLDSILNETRLKPSDEGFDVAKRGVEAFISELLSSSNTEKVDQSLVDLMISEIDQKLSKQVDAILHNEEVQAIESTWRGLKYLVDH
TDFRENIQIELISAKKDEVLDDFEDAPEVVKSGLYKQIYTREYGQFGGKPVGAVICDYNMSASSPDIKLMEYMANVGAMSHAPFITSASAKFFGLDSYEELPNL
KDLKSVFEGPQYTKWRGLREHEDARYLGLCTSRFMLRTPYSVEDNPIKAFDYDEHVTDSHDNYLWGNSAYAMASKISESFAKYRWCPNIIGPQSGGSVHDLP
VYNFEAMGQIETKIPTEILVSDRREYELAEEGFIALTMRKGSDNAAFFSANSVQKPKVFANTPEGKQAEMNYKLGTQLPYMFIINRLAHYIKVLQREQIGSWKE
RSDLEIELNKWIRQYVSDQENPPAEVRGRRPLRAAKVEVSDVEGDPGWYKVSMSVRPHFKYMGASFDLSLVGKLDQ

>Vibrio_parahaemolyticus_BB22OP_T6SS-2
MTTQVEKQLDPTVGEQTTSFLEQAIGATKQTEASRAEELIKTLTEEAMKGTVSWNKNLTVTFREAINLLDRQISEQLSEVMHHPELQKLEGSWRGLNYLVMNS
ETSSTLKIRMISITKKELHKDLSKAVEFDQSQIFKKVYESEFGSAGGEPYGALIGDYEFTNHPEDIESLRLMSNVAASGFSPFLSAASPALFGFDEWTELSKPRDL
DKVFESLEYAQWRSFRESADSRFVSLTMPKVLARLPYGQATSPVEAFGFEEFDVDPVSGIAVNADHNDYCWMNSSYVLGVKLTDAFSKYGFCTAIRGAEGGG
RVDNLPTHFFMSDDGDPDMKCPTEIGITDRREAELGKLGFLPLCHYKNTNYAVFFGAQTCQKPANHESPEVAANAAISARLPYMMATSRFAHYLKVMARDKI
GSFMEAEDVESWLNRWILGYVNASEGGGQEIRAKYPLADARVQVKEIPGSPGSYNAVAWLKPWLQMEELTTSLRLVAKIPQSGG

>Vibrio_parahaemolyticus_RIMD_2210633_T6SS-1
MSAEAQAPEQEAALVESGSLLDSILNETRLKPSDEGFDVAKRGVEAFISELLSSSNTEKVDQSLVDLMISEIDQKLSKQVDAILHNEEVQAIESTWRGLKYLVDH
TDFRENIQIELISAKKDEVLDDFEDAPEVVKSGLYKQIYTREYGQFGGKPVGAVICDYNMSASSPDIKLMEYMANVGAMSHAPFITSASAKFFGLDSYEELPNL
KDLKSVFEGPQYTKWRGLREHEDARYLGLCTSRFMLRTPYSVEDNPIKAFDYDEHVTDSHDNYLWGNSAYAMASKISESFAKYRWCPNIIGPQSGGSVHDLP
VYNFEAMGQIETKIPTEILVSDRREYELAEEGFIALTMRKGSDNAAFFSANSVQKPKVFANTPEGKQAEMNYKLGTQLPYMFIINRLAHYIKVLQREQIGSWKE
RSDLEIELNKWIRQYVSDQENPPAEVRGRRPLRAAKVEVSDVEGDPGWYKVSMSVRPHFKYMGASFDLSLVGKLDQ

>Vibrio_parahaemolyticus_RIMD_2210633_T6SS-2
MTTQVEKQLDPTVGEQTTSFLEQAIGATKQTEASRAEELIKTLTEEAMKGTVSWNKNLTVTFREAINLLDRQISEQLSEVMHHPELQKLEGSWRGLNYLVMNS
ETSSTLKIRMISITKKELHKDLSKAVEFDQSQIFKKVYESEFGSAGGEPYGALIGDYAQWRSFRESADSRFVSLTMPKVLARLPYGQATSPVEAFGFEEFDVDPV
SGIAVNADHNDYCWMNSSYVLGVKLTDAFSKYGFCTAIRGAEGGGRVDNLPTHFFMSDDGDPDMKCPTPYMMATSRFAHYLKVMARDKIGSFMEAEDVES
WLNRWILGYVNASEGGGQEIRAKYPLADARVQVKEIPGSPGSYNAVAWLKPWLQMEELTTSLRLVAKIPQSGG

>Vibrio_parahaemolyticus_strain_12-009A/1335_T6SS-1
MSAEAQAPEQEAALVESGSLLDSILNETRLKPSDEGFDVAKRGVEAFISELLSSSNTEKVDQSLVDLMISEIDQKLSKQVDAILHNEEVQAIESTWRGLKYLVDH
TDFRENIQIELISAKKDEVLDDFEDAPEVVKSGLYKQIYTREYGQFGGKPVGAVICDYNMSASSPDIKLMEYMANVGAMSHAPFITSASAKFFGLDSYEELPNL
KDLKSVFEGPQYTKWRGLREHEDARYLGLCTSRFMLRTPYSVEDNPIKAFDYDEHVTDSHDNYLWGNSAYAMASKISESFAKYRWCPNIIGPQSGGSVHDLP
VYNFEAMGQIEMKIPTEILVSDRREYELAEEGFIALTMRKGSDNAAFFSANSVQKPKVFANTPEGKQAEMNYKLGTQLPYMFIINRLAHYIKVLQREQIGSWK
ERSDLEIELNKWIRQYVSDQENPPAEVRGRRPLRAAKVEVSDVEGDPGWYKVSMSVRPHFKYMGASFDLSLVGKLDQ

>Vibrio_parahaemolyticus_strain_12-009A/1335_T6SS-2

MTTQVEKQLDPTVGEQTTSFLEQAIGATKQTEASRAEELIKTLTEEAMKGTVSWNKNLTVTFREAINLLDRQISEQLSEVMHHPELQKLEGSWRGLNYLVMNS
ETSSTLKIRMISITKKELHKDLSKAVEFDQSQIFKKVYESEFGSAGGEPYGALIGDYEFTNHPEDIESLRLMSNVAASGFSPFLSAASPALFGFDEWTELSKPRDL
DKVFESLEYAQWRSFRESADSRFVSLTMPKVLARLPYGQATSPVEAFGFEEFDVDPVSGIAVNADHNDYCWMNSSYVLGVKLTDAFSKYGFCTAIRGAEGGG
RVDNLPTHFFMSDDGDPDMKCPTEIGITDRREAELGKLGFLPLCHYKNTNYAVFFGAQTCQKPANHESPEVAANAAISARLPYMMATSRFAHYLKVMARDKI
GSFMEAEDVESWLNRWILGYVNASEGGGQEIRAKYPLADARVQVKEIPGSPGSYNAVAWLKPWLQMEELTTSLRLVAKIPQSGG

>Vibrio_parahaemolyticus_strain12-297/B

MSAEAQAPEQEAALVESGSLLDSILNETRLKPSDEGFDVAKRGVEAFISELLSSSNTEKVDQSLVDLMISEIDQKLSKQVDAILHNEEVQAIESTWRGLKYLVDH
TDFRENIQIELISAKKDEVLDDFEDAPEVVKSGLYKQIYTREYGQFGGKPVGAVICDYNMSASSPDIKLMEYMANVGAMSHAPFITSASAKFFGLDSYEELPNL
KDLKSVFEGPQYTKWRGLREHEDARYLGLCTSRFMLRTPYSVEDNPIKAFDYDEHVTDSHDNYLWGNSAYAMASKISESFAKYRWCPNIIGPQSGGSVHDLP
VYNFEAMGQIEMKIPTEILVSDRREYELAEEGFIALTMRKGSDNAAFFSANSVQKPKVFANTPEGKQAEMNYKLGTQLPYMFIINRLAHYIKVLQREQIGSWK
ERSDLEIELNKWIRQYVSDQENPPAEVRGRRPLRAAKVEVSDVEGDPGWYKVSMSVRPHFKYMGASFDLSLVGKLDQ

>Vibrio_parahaemolyticus_strain_D4_T6SS-1

MSAEAQAPEQEAALVESGSLLDSILNETRLKPSDEGFDVAKRGVEAFISELLSSSNTEKVDQSLVDLMISEIDQKLSKQVDAILHNEEVQAIESTWRGLKYLVDH
TDFRENIQIELISAKKDEVLDDFEDAPEVVKSGLYKQIYTREYGQFGGKPVGAVICDYNMSASSPDIKLMEYMANVGAMSHAPFITSASAKFFGLDSYEELPNL
KDLKSVFEGPQYTKWRGLREHEDARYLGLCTSRFMLRTPYSVEDNPIKAFDYDEHVTDSHDNYLWGNSAYAMASKISESFAKYRWCPNIIGPQSGGSVHDLP
VYNFEAMGQIETKIPTEILVSDRREYELAEEGFIALTMRKGSDNAAFFSANSVQKPKVFANTPEGKQAEMNYKLGTQLPYMFIINRLAHYIKVLQREQIGSWKE
RSDLEIELNKWIRQYVSDQENPPAEVRGRRPLRAAKVEVSDVEGDPGWYKVSMSVRPHFKYMGASFDLSLVGKLDQ

>Vibrio_parahaemolyticus_strain_D4_T6SS-2

MTTQLETQLDPTVEGQSYSFLEQAIGATKQTEASRAEELIKTLTEEALKGTVTWNKNLTVTFREAITALDKKISEQLSAVMHHPELQKLEGTWRGLNYLVMNS
ETSSTLKIRMVSMTKKELHKDLSKAVEFDQSQVFKKVYESEFGSAGGEPYGAIIGDYEFTNHPEDIETLRLMSNVAAAGFSPFLSAASPALFGFDEWTELSKPR
DLEKVFESLEYTQWRAFRESQDSRFVSLTMPKVLARLPYGQATSPVEEFGFEEFSIDETLGIAVNAQHNDYCWMNSSYVLGAKLTDAFAKYGFCTAIRGAEGG
GRVDNLPTHYFMSDDGDPDMKCPTEIGITDRRESELGKLGFLPLCHYKNTNYAVFFGAQTCQKPAKYDSPDATANAAISARLPYMMATSRFAHYLKVMARDK
IGSFMEAEDVESWLNRWILGYVNASEGGGQEIRAKYPLADARVQVREVPGAPGSYNAVAWLKPWLQMEELTTSLRLVAKIPQSGG

>Vibrio_parahaemolyticus_strain_T9109

MSAEAQAPEQEAALVESGSLLDSILNETRLKPSDEGFDVAKRGVEAFISELLSSSNTEKVDQSLVDLMISEIDQKLSKQVDAILHNEEVQAIESTWRGLKYLVDH
TDFRENIQIELISAKKDEVLDDFEDAPEVVKSGLYKQIYTREYGQFGGKPVGAVICDYNMSASSPDIKLMEYMANVGAMSHAPFITSASAKFFGLDSYEELPNL
KDLKSVFEGPQYTKWRGLREHEDARYLGLCTSRFMLRTPYSVEDNPIKAFDYDEHVTDSHDNYLWGNSAYAMASKISESFAKYRWCPNIIGPQSGGSVHDLP
VYNFEAMGQIETKIPTEILVSDRREYELAEEGFIALTMRKGSDNAAFFSANSVQKPKVFANTPEGKQAEMNYKLGTQLPYMFIINRLAHYIKVLQREQIGSWKE
RSDLEIELNKWIRQYVSDQENPPAEVRGRRPLRAAKVEVSDVEGDPGWYKVSMSVRPHFKYMGASFDLSLVGKLDQ

>Vibrio_proteolyticus_NBRC_13287

MTAEAQAPEQEAALAESGSLLDSILNETRLKPSDEGFDVAKRGVEAFIGELLSNSASEKVDQSLVDLMIAEIDQKLSTQVDAILHNEEVQAIESTWRGLKYLVD
HTDFRENIQIELISAKKDEVLDDFEDAPEVVKSGLYKQIYTREYGQFGGKPVGAVICDYQLSAATPDIKLMEYMANVGAMSHAPFITSASSKFFGLDSYEELPN
LKDLKAVFEGPQYAKWRGLREHEDARYLGLCTSRFMLRNPYSVEDNPIKAFDYNEMVTDNHDHFLWGNSAYAMASKISESFAKYRWCPNIIGPQSGGAVFDL
PVYNYESMGQIETKIPTEILVSDRREFELAEEGFVALTMRKGSDNAAFFSANSIQKPKVFPNTPEGKQAEMNYKLGTQLPYMFIINRLAHYIKVLQREQIGSWKE
RSDLEIELNKWIRQYVSDQENPPAEVRGRRPLRAAKVEVSDVEGDPGWYRVSMSVRPHFKYMGASFDLSLVGKLDQN

>Vibrio_vulnificus_106-2A_T6SS-1

MSTTETVQERPQLLQGGLLDEIMAQTRIAPSEEGYDIAKKGVAAFIENLIDSKHSADPVNKSLVDQMLVELDKKISSQMDEILHDSKFQAMESSWRGLKLFVD
RTDFRENNKVEIIHVTKEELLEDFEFAPETSQSGLYKHVYSAGYGQFGGEPVGAMIGNYVFTPSTPDMKLLQYMGALGAMAHAPFISSVGPEFFGIDSFEELPN

VKDLKSTFESPKYTKWRSIRESEDARYIGLTAPRFLLRVPYDPTENPVKSFNYEEGVSASHEHYLWGNTAFAFATRLTDSFAKYRWCPNIIGPQSGGAVEDLPVH
VFESMGALQSKIPTEVLITDRKEFELAEEGFIALTMRKGSDNATFFSANSIQKPKIFPNTKEGKEAETNYKLGTQLPYMMIINRLAHYVKVLQREQIGAWKERQ
DLERELNSWIKQYVADQENPPADVRSRRPLRAARIEVMDVEGNPGWYQVSLSVRPHFKYMGANFELSLVGRLDQA
G

>Vibrio_vulnificus_106-2A_T6SS-2
MTTQLETQLDPTVEGQSYSFLEQAIGATKQTEASRAEELIKTLTEEALKGTVTWNKNLTVTFREAITALDKKISEQLSAVMHHPELQKLEGTWRGLNYLVMNS
ETSSTLKIRMVSMTKKELHKDLSKAVEFDQSQVFKKVYESEFGSAGGEPYGAIIGDYEFTNHPEDIETLRLMSNVAAAGFSPFLSAASPALFGFDEWTELSKPR
DLEKVFESLEYTQWRAFRESQDSRFVSLTMPKVLARLPYGQATSPVEEFGFEEFSIDETLGIAVNAQHNDYCWMNSSYVLGAKLTDAFAKYGFCTAIRGAEGG
GRVDNLPTHYFMSDDGDPDMKCPTEIGITDRRESELGKLGFLPLCHYKNTNYAVFFGAQTCQKPAKYDSPDATANAAISARLPYMMATSRFAHYLKVMARDK
IGSFMEAEDVESWLNRWILGYVNASEGGGQEIRAKYPLADARVQVREVPGAPGSYNAVAWLKPWLQMEELTTSLRLVAKIPQSGG

>Xanthomonas_oryzae_pv._oryzae_PXO99A
MAAEVSSATHNSAPAALAETGLLDQIVEQSKVAKSETEHQRAKDIISELAKEVLEGTVVVSDNLNLTLDARIAEIDRIISEQLSAVMHAPEFQKLESSWRGLHY
MCEQTSTGVTMKIKVFNSPQRDLVRDFKSAIDFDQSALFKKVYEEEFGTFGGAPFGALVGDYYIGRQPEDMYFVEQMSHVAAAAHAPFISAASESMFGLETFT
DLGKPRDLAKVFDTVEYAKWKSFRESEDSRYVGLTMPRFLGRLPYNPKDGTTVEGFNFVEDVDAADHSKYLWCNTAYAMATRLTKAFEDYGWCAAIRGVE
GGGLVEDLPTHTFRTDDGEVALKCPTEIAITDRREKELSDLGFIPLVHCKNTDYAAFFGAQSAQKPKKYNTDAANANAILSAQLQYMFAVSRIAHYLKAMMRE
KIGSFASARNVEDFLNRWISQYVLLDDNAGQEAKAQYPLREASVQVSEVPGKPGVYRAVAFIRPHFQLDELSVSLRLVAELPAGTKKA

>Xanthomonas_oryzae_pv._oryzicola_strain_GX01
MVAEVSSATHNSAPAALAETGLLDQIVEQSKVAKSETEHQRAKDIISELAKEVLEGTVVVSDNLNLTLDARIAEIDRIISEQLSAVMHAPEFQKLESSWRGLHYM
CEQTSTGVTMKIKVFNSPQRDLVRDFKSAIDFDQSALFKKVYEEEFGTFGGAPFGALVGDYYIGRQPEDMYFVEQMSHVAAAAHAPFISAASESMFGLETFTD
LGKPRDLAKVFDTVEYAKWKSFRESEDSRYVGLTMPRFLGRLPYNPKDGTTVEGFNFVEDVDAADHSKYLWCNTAYAMATRLTKAFEDYGWCAAIRGVEG
GGLVEDLPTHTFRTDDGEVALKCPTEIAITDRREKELSDLGFIPLVHCKNTDYAAFFGAQSAQKPKKYNTNAANANAILSAQLQYMFAVSRIAHYLKAMMREK
IGSFASARNVEDFLNRWISQYVLLDDNAGQEAKAQYPLREASVQVSEVPGKPGVYRAVAFIRPHFQLDELSVSLRLVAELPAGTKKA

>Xenorhabdus_bovienii_SS-2004_T6SS-1
MAQHEENSTALASGATTSLLDEIMSQARITPENDGYYIAKQGIAAFISSILDTGANEEPINKLVVDKMIVELDKKLSEQMDEIMHAKPFQELESSWRSLKILVDR
TDFRENIKINIIHATKDEMLEDFEFSPEIIQSGFYKHVYSTGYGQFGGEPVAAVIGNYAFTNSSPDIKLMQYVSAVGAMAHSPFLSSVSPEFFGVNSFTDLPAIKDL
KSVFEGPSHTKWRALRESEDSRYLGLTAPRFLLRLPYSSVENPIKNFNYQENVSRDHEHFLWGNTAYLMASCLTDSFAKYRWCPNIIGPQSGGTVSDLPVHLYE
AMGQVQAKIPTEVLITDRREFELAEEGFITLTMRKGSDNAAFFSANSVQKPKVFPNTREGKIAETNYKLGTQLPYMFIINRLAHYIKVLQREQIGSWKERQDLE
RELNMWLKQYIADQENPPTDVRSRRPLRAAEIKVLDVEGDPGWYQVAMQVRPHFKYMGASFELSLVGRLDKE

>XenorhabdusbovieniiSS-2004_T6SS-2
MSTTEQETEFQQEQLSKEYTPDELSVLLNKEFRPQSDQARSAVESAVTTLAQQALENTVTISGDTYRTIQALITEIDKKLSQQINHILHHDEFQALEGAWRGLHY
LVNNTETDEMLKIRVMCISKKELGSTLKRFKGVSWDQSPVFKKIYEAEYGQFGGEPYGCLIGDYYFDHTAPDVELLREMAQISAAAHCPFLSGASPSVMQME
CWQELSNPRDLTKVFQNTEYAPWHSLRESEDVRYLGLTLPRFLSRLPYGNKTNPVDNFNFEEDTEGPTHSNYTWANAAYAMAVNINRSFKLYGWCTSIHGVES
GGTVENLPCYTFPTDDGGVDIKCPTEIAISDRREAELAKNGFIPLLHRKNTDMAAFIGAQSLQKPSEYYDHDATANANLSARLPYLFACCRFAHYLKCIVRDKI
GTFQERNDMENWLNNWIINYVDGDPTNSSAGTKARKPLAAAEVQVEEIPSNPGYYHAKFFLRPHYQLEGLTVSLRLVSRLPSLKST

>Yersinia_pestis_CO92
MQKAQLNDTPDINVDLAENELDDILKRSFRPRTDEASATVRRAIGTLAAYANKGQVKVNRDVVLTIESLVAEIDEKLSDQMNLILHHKEFQKLESAWRGLSYL
VDNTDANETLKIRVLNISQDELGKTLRRYRGSAWDQSPIFKQVYEHEYGQFGGEPFGCMIGDYEFDHSPQSVALLTELAKVAAAAHCPFITSSSPSIMQMNNWR
ELGNPRDIGKIFTTPEYAPWRRLRESNDSRYLVLTLPRFLSRLPYGAKNHPIDDFAFEEVVSPHASEDFTWANAAYAMGVNINRAFNEYGWCSRIRGIESGGSVE
ELPAYAFPSDEGGYELTCPTELAISDRRENELSNAGFLPLVYRKNSDFAAFIGSCTLHAPANYDDPDATANARLSARLPYIFATCRFAHYLKCIVRDKIGSFRSRD

DMQLWLNDWVMNYVDGDPSISTEATKAKRPLAAAEVQVEDVEDDPGFYRAHFYLRPHYQLEGMTVSLRLVSKLPSNKESEKK

>Yersinia_pseudotuberculosis_IP_31758
MNLILHHKEFQKLESAWRGLSYLVDNTDANETLKIRVLNISQDELGKTLRRYRGSAWDQSPIFKQVYEHEYGQFGGEPFGMQKAQLNDTPDINVDLAENELD
DILKRSFRPRTDEASATVRRAIGTLAAYANKGQVKVNRDVVLTIESLVAEIDEKLSDQCMIGDYEFDHSPQSVALLTELAKVAAAAHCPFITSSSPSIMQMNNWR
ELGNPRDIGKIFTTPEYAPWRRLRESNDSRYLVLTLPRFLSRLPYGAKNHPIDDFAFEEVVSPHASEDFTWANAAYAMGVNINRAFNEYGWCSRIRGIESGGSVE
ELPAYAFPSDEGGYELTCPTELAISDRRENELSNAGFLPLVYRKNSDFAAFIGSCTLHAPANYDDPDATANARLSARLPYIFATCRFAHYLKCIVRDKIGSFRSRD
DMQLWLNDWVMNYVDGDPSISTEATKAKRPLAAAEVQVEDVEDDPGFYRAHFYLRPHYQLEGMTVSLRLVSKLPSNKESEKK

>Yersinia_pseudotuberculosis_YPIII_T6SS-4
MQKAQLNDTPDINVDLAENELDDILKRSFRPRTDEASATVRRAIGTLAAYANKGQVKVNRDVVLTIESLVAEIDEKLSDQMNLILHHKEFQKLESAWRGLSYL
VDNTDANETLKIRVLNISQDELGKTLRRYRGSAWDQSPIFKQVYEHEYGQFGGEPFGCMIGDYEFDHSPQSVALLTELAKVAAAAHCPFITSSSPSIMQMNNWR
ELGNPRDIGKIFTTPEYAPWRRLRESNDSRYLVLTLPRFLSRLPYGAKNHPIDDFAFEEVVSPHASEDFTWANAAYAMGVNINRAFNEYGWCSRIRGIESGGSVE
ELPAYAFPSDEGGYELTCPTELAISDRRENELSNAGFLPLVYRKNSDFAAFIGSCTLHAPANYDDPDATANARLSARLPYIFATCRFAHYLKCIVRDKIGSFRSRD
DMQLWLNDWVMNYVDGDPSISTEATKAKRPLAAAEVQVEDVEDDPGFYRAHFYLRPHYQLEGMTVSLRLVSKLPSNKESEKK

>Yersinia_pseudotuberculosis_YPIII_T6SS-3
MSTQDAKTKQATVASATEYSDFNTLLTKEFKAKSEQTIAAVEGAVKTLAEQALANSVTVSDDAYKSIASIIAEIDRKLSDQINLILHHQEFQSLESAWRGLHYLV
NNTETDEKLRLRFLDISKDELRRNMKRYKGVAWDQSPLFKKIYEEEYGQLGGEPYGCLVADYYFDHTAPDVDLLASIGKVAASAHVPFITGAAPSVLQMESW
QELSNPRDLTKIFTQNLEYAAWNSLRQSEDSRYIGLAMPRFLSRLPYGINTNPVDAFHFEETTDGADHSKYVWSNAAYAMAVNINRSFKEYGWCTLIRGVESG
GVVEGLPCHTFPTDDGGIDMKCPTEIAISDRREAELAKNGFIPLVHRKNTDYAAFIGAQSLQKPAEYYDSDATANANLSARLPYLFACSRFAHYLKCIVRDKIGS
FKERDEMQRWLNDWVMNYVDGDPANSSLETKARRPLAAAEVIVEEVEGNPGYYQAKFFLRPHFQLEGLTVSLRMVAKLPSLKDVA
